# Poorly expressed alleles of several human immunoglobulin heavy chain variable (IGHV) genes are common in the human population

**DOI:** 10.1101/2020.09.05.284257

**Authors:** Mats Ohlin

**Author notes:** Correspondence should be addressed to: Mats Ohlin, Dept. of Immunotechnology, Lund University, Medicon Village building 406, S-22381 Lund, Sweden.

## Abstract

Extensive diversity has been identified in the human heavy chain immunoglobulin locus, including allelic variation, gene duplication, and insertion/deletion events. Several genes have been suggested to be deleted in many haplotypes. Such findings have commonly been based on inference of germline repertoire from data sets covering antibody heavy chain encoding transcripts. The inference process operate under conditions that may limit identification of genes transcribed at low levels. The presence of rare transcripts that would indicate the presence of poorly expressed alleles in haplotypes that otherwise appear to have deleted these genes has now been assessed. Alleles IGHV1-2*05, IGHV1-3*02, IGHV4-4*01, and IGHV7-4-1*01 were all identified as being expressed at very low levels from multiple haplotypes, haplotypes that by inference often appeared not to express these genes at all. These alleles harbor unusual sequence variants that may compromise the functionality of the encoded products. Transcripts of two of these alleles to a large degree do not encode a functional product, suggesting that these alleles might be non-functional. It is proposed that the functionality status of immunoglobulin genes should also include assessment of their ability to encode functional protein products.

## INTRODUCTION

The specificity-defining variable domains of human immunoglobulin heavy chains are encoded by genes (IGHV, IGHD, and IGHJ) located on chromosome 14. Several alleles have been associated with most of the IGHV genes and substantial differences between subjects exist with respect to which genes/alleles that are available to mount an immune response. Such genes are described in the IMGT database (1) that is employed by a range of bioinformatics tools to define the genes and the downstream hypermutation processes involved in establishment and evolution of particular antibodies. Such processes require precise definition of the germline IGHV, IGHD and IGHJ genes that have been used to generate the gene encoding an antibody of interest. This analysis is complicated by the fact that germline gene databases are incomplete and even contain allelic sequences in error (2). Extensive efforts are in place to describe new germline gene alleles and to define immunoglobulin genotypes/haplotypes of multiple subjects. Genomic sequencing, in particular the introduction of long-read sequence technology, will likely offer important insight in this field of research in the future (3). Germline gene inference based of the information content of next generation sequencing (NGS) data sets has, however, emerged as an important approach to define personalized germline gene allele repertoires available to individuals to generate their antibody responses (4–8). These approaches allows for better gene/allele assignment and tracking of hypermutation pathways that have resulted in antibody sequences of the subject under investigation. The germline gene/allele repertoire of large numbers of individuals has now been described by use of this approach. The VDJbase server (https://www.vdjbase.org/) (9) allows public access to such processed information. The fact that numerous subjects carry different alleles of IGHJ allows further refinement and validation of gene assignments through haplotyping (10–13) and a deeper understanding of antibody-encoding genes that can be generated through gene rearrangement and somatic hypermutation. That approach has furthermore enabled identification of gene deletion and insertion events and other complex genetic events only some of which had been previously characterized (14). It is possible to readily identify haplotypes that carry either the IGHV1-8/IGHV3-9 or the IGHV3-64D/IGHV5-10-1 genes, or those that have incorporated well-recognized insertions close to e.g. IGHV3-30, IGHV3-43, IGHV1-69, and IGHV2-70 genes. It has however also been possible to identify allele usage bias and mosaic patters of deleted genes in a large set of subjects (10), as well as functional deletions of large parts of the IGHV locus (10,11).

Several genes including IGHV1-3, IGHV4-4, and IGHV7-4-1, and to some extent also IGHV1-2, proximal to the IGHD gene locus were shown to be subject to a lack of perceived expression from one or both haplotypes of many subject under investigation (10). This effect may be caused by an absence of the gene in question, or complete silencing of gene transcription/expression, but also by filtering strategies in inference programs that in the interest of analysis specificity sacrifice the ability of the tool to detect alleles expressed at low levels, or at low levels compared to other alleles of the same gene that are present in the genotype. In this study, evidence of alleles of these four genes is identified, alleles that typically escape detection by inference tools as they are transcribed at levels much lower than other common alleles of the same gene.

## MATERIAL AND METHODS

### Immunoglobulin transcriptome data sets

100 immunoglobulin variable domain-encoding transcriptome data sets of project PRJEB26509 were downloaded from the European Nucleotide Archive. All data sets except ERR2567273, and ERR2567275 were used for the present analysis. These data sets represent immunoglobulin IgM and IgD heavy-chain encoding transcriptomes and light chain-encoding transcriptomes of sorted (CD19^+^, CD27^-^, IgD^+^, IgA^-^, and IgG^-^) human naïve B-cells derived from healthy subjects and subjects with celiac disease in Norway (10). The libraries had been generated using 5’-RACE technology and sequenced on the MiSeq sequencing platform. 94 of the data sets, including 33 sets that could be further studied using haplotyping based on IGHJ6 heterozygosity, were used in a past study of deletion patterns in IGHV repertoires (10). 96 of these data sets, including 34 sets that could be further studied using haplotyping based on IGHJ6 heterozygosity, are also described in VDJbase (https://www.vdjbase.org/) (9) as study P1.

Data sets of IgG and IgA-encoding transcriptomes of peripheral blood and bone marrow of two subjects (donors 2 and 4) (15), previously shown to carry the IGHV7-4-1*01 but not the IGHV7-4-1*02 allele (11,13), were used to illustrate evolution of IGHV7-4-1*01 during somatic hypermutation and selection. Pre-processing of the data as well as subsequent IgDiscover-based germline gene inference and IMGT/HighV-QUEST analysis of these data sets have been previously described (11,15).

### Germline gene inference

Heavy chain immunoglobulin germline genes were inferred from NGS data sets of project PRJEB26509 using IgDiscover 0.11 (6). Inference (1 iteration) was performed using settings summarized in Supplementary Methods. A set (Supplementary Methods) of all alleles of human germline heavy chain genes that had at least one allele considered to be functional by the international ImMunoGeneTics information system^®^ (IMGT) was downloaded from http://www.imgt.org and used as the starting database for the inference process. Bases of IGHV genes are numbered according to the standard IMGT numbering system (16).

### IMGT/HighV-QUEST analysis

Immunoglobulin germline gene assignment was performed using IMGT/HighV-QUEST (17) on data sets that could be haplotyped based on association to different alleles of IGHJ6 (10). The assay was performed using IMGT/V-QUEST program version 3.5.18 or 3.5.19, and IMGT/V-QUEST reference directory release 202011-3 or 202031-2. Reads that were unequivocally assigned to a single germline gene allele were used for analysis of allele expression levels, functionality, and length of the CDR3-encoding part of rearranged genes.

### Structures of antibodies with an origin in IGHV7-4-1

The structures of five antibodies with an origin in IGHV7-4-1 for which high resolution (<2.5 Å) structures as defined in IMGT/3Dstructure-DB and IMGT/2Dstructure-DB (18) were investigated. Coordinates for PDB entries 4D9Q, 4EOW, 5CGY, 5ZMJ, and 6B5R were downloaded from Protein Data Bank (https://www.rcsb.org/) and visualized using PyMOL 2.0.5 (The PyMOL Molecular Graphics System, Schrödinger, LLC).

## RESULTS

### Germline allele IGHV1-2*05 is expressed at low levels compared to other common alleles of this gene

In a past study using the TIgGER inference tool, it was suggested that IGHV1-2 was deleted in at least 7/66 haplotypes of 33 subjects (DOI: 10.1038/s41467-019-08489-3). VDJbase (9) report common occurrences of IGHV1-2*02 (n=66), IGHV1-2*04 (n=63), and IGHV1-2*06 (n=18), and one case of IGHV1-2*07, among the 96 subjects of the study as documented in the database, but no cases of IGHV1-2*01 and IGHV1-2*05. Furthermore, the gene is reported to be deleted in 6/68 haplotypes of 34 subjects in VDJbase (Supplementary Figure 1). When reanalyzing publicly available data sets of this study (PRJEB26509) with the IgDiscover tool the common expression of alleles IGHV1-2*02, IGHV1-2*04, and IGHV1-2*06 in these subjects was confirmed. IGHV1-2*07, a recently recognized allele that was not present in the database used to initiate the inference process, was not inferred by IgDiscover but in-depth examination of the reads associated with IGHV1-2 in data set ERR2567201 confirmed the presence of this allele in one of the haplotypes of this subject (data not shown). Neither IGHV1-2*01 nor IGHV1-2*05 were reported by IgDiscover to be present in the final inferred genotypes of any of these subjects (Table 1, Supplementary Figure 2). IGHV1-2*05 was however implicated as being present at low levels in five data sets although the inference was not maintained following the final filtering step (Table 1). Genomic data suggests that SNPs relevant for the presence of IGHV1-2*05 in European populations has been identified (Supplementary Figure 3). This prompted further detailed assessment of the available data.

**Table 1.** Alleles of IGHV1-2, IGHV1-3, IGHV4-4, and IGHV7-4-1 of 35 haplotypable (based on heterozygocity of IGHJ6) data sets used for the present analysis. Inferred alleles as defined by IgDiscover in the V_expressed file (Supplementary Figure 2) and alleles that are scored by IMGT/HighV-QUEST are shown.

|  | Data set (run accession number) | Reads assigned by IMGT/HighV-QUEST to IGHV after preprocessing of data set | IGHV1-2 |  | IGHV1-3 |  | IGHV4-4 |  | IGHV7-4-1 |  |
| --- | --- | --- | --- | --- | --- | --- | --- | --- | --- | --- |
|  |  |  | IgDiscover | High/V-Quest | IgDiscover | High/V-Quest | IgDiscover | High/V-Quest | IgDiscover | High/V-Quest |
| 1 | ERR2567187 | 225406 | *04<sup>†</sup> | *04, *05 | *01 | *01 | *02 | *01, *02 | *02 | *01, *02 |
| 2 | ERR2567189 | 224871 | *02<sup>¶</sup> | *02, *04 | *01 | *01, *02 | *02, *07 | *02, *07 |  | *01 |
| 3 | ERR2567192 | 212646 | *02, *04 | *02, *04 | *01 | *01 | *02 | *02 | *01 | *01 |
| 4 | ERR2567199 | 240716 | *02, *04 | *02, *04 | *01 | *01, *02 | *02, *07 | *02, *07 | *01 | *01 |
| 5 | ERR2567200 | 180844 | *02, *04 | *02, *04 | *01 | *01, *02 | *02, *07 | *02, *07 | *01 | *01 |
| 6 | ERR2567201 | 215909 | *02 | *02, *07 |  | *02 | *07 | *07 |  |  |
| 7 | ERR2567204 | 265106 | *04 *06 | *04 *06 | *01 | *01 | *02 | *02 | *02 | *01, *02 |
| 8 | ERR2567206 | 262069 | *06 | *05, *06 | *01 | *01 | *02 | *01, *02 | *02 | *02 |
| 9 | ERR2567213 | 211280 | *02 | *02 |  | *02 | *07 | *07 |  |  |
| 10 | ERR2567214 | 276041 | *04 | *04 | *01 | *01 | *02 | *02 | *01 | *01 |
| 11 | ERR2567215 | 220448 | *02, *04 | *02, *04 | *01 | *01, *02 | *02, *07 | *02, *07 | *01 | *01 |
| 12 | ERR2567217 | 182276 | *02 | *02 |  | *02 | *07 | *07 |  |  |
| 13 | ERR2567220 | 151436 | *02<sup>¶</sup> | *02, *04 | *01 | *01, *02 | *02, *07 | *02, *07 | *01 | *01 |
| 14 | ERR2567221 | 164821 | *02<sup>¶</sup> | *02, *04 | *01 | *01 | *02, *07 | *02, *07 |  |  |
| 15 | ERR2567223 | 156379 | *02, *06 | *02, *06 | *01 | *01, *02 | *02, *07 | *02, *07 | *02 | *02 |
| 16 | ERR2567226 | 117706 | *02 | *02 |  | *02 | *02, *07 | *02, *07 |  | *01 |
| 17 | ERR2567230 | 183580 | *02 | *02 | *02 | *02 | *02, *07 | *02, *07 |  | *01 |
| 18 | ERR2567231 | 240347 | *02<sup>†</sup> | *02, *05 | *01 | *01, *02 | *07 | *01, *07 | *02 | *02 |
| 19 | ERR2567232 | 192854 | *02, *06 | *02, *06 | *01 | *01, *02 | *02, *07 | *02, *07 | *02 | *02 |
| 20 | ERR2567240 | 149185 | *02 | *02 |  | *02 | *07 | *07 |  |  |
| 21 | ERR2567242 | 348123 | *02<sup>¶</sup> | *02, *04 | *01 | *01, *02 | *02, *07 | *02, *07 |  | *01 |
| 22 | ERR2567243 | 253195 | *04<sup>†</sup> | *04, *05 | *01 | *01 | *02 | *01, *02 | *02 | *01, *02 |
|  |  |  | IgDiscover | High/V-Quest | IgDiscover | High/V-Quest | IgDiscover | High/V-Quest | IgDiscover | High/V-Quest |
| 23 | ERR2567246 | 223410 | *04 | *04 | *01 | *01 | *02 | *02 | *01 | *01 |
| 24 | ERR2567249 | 223137 | *04<sup>†</sup> | *04, *05 | *01 | *01 | *02 | *01, *02 | *02 | *01, *02 |
| 25 | ERR2567254 | 241355 | *04 | *04 | *01 | *01 | *02 | *02 | *01 | *01 |
| 26 | ERR2567259 | 189273 | *02<sup>†</sup> | *02, *05 | *01 | *01, *02 | *07 | *01, *07 | *02 | *02 |
| 27 | ERR2567261 | 308875 | *04 | *04 | *01 | *01 | *02 | *02 | *01 | *01 |
| 28 | ERR2567263 | 312067 | *02 | *02 | *02 | *02 | *07 | *07 |  |  |
| 29 | ERR2567264 | 226845 | *02, *04 | *02, *04 | *01 | *01, *02 | *02 | *02 | *01 | *01 |
| 30 | ERR2567265 | 194775 | *02, *04 | *02, *04 | *01 | *01, *02 | *02, *07 | *02, *07 | *01 | *01 |
| 31 | ERR2567266 | 321732 | *04 | *04 | *01 | *01 | *02, *07 | *02, *07 |  | *01 |
| 32 | ERR2567271 | 168997 | *06<sup>¶</sup> | *04 *06 | *01 | *01 | *02 | *02 | *02 | *01, *02 |
| 33 | ERR2567274 | 184738 | *04 | *04 | *01 | *01 | *02 | *02 |  | *01 |
| 34 | ERR2567276 | 148621 | *02, *06 | *02, *06 | *01 | *01, *02 | *02, *07 | *02, *07 | *02 | *02 |
| 35 | ERR2567277 | 193732 | *02 | *02, *04 | *01 | *01, *02 | *02, *07 | *02, *07 | *01 | *01 |

<sup>†</sup> IgDiscover tentatively inferred IGHV1-2\*05 in this data set but the allele did not feature in the more strictly filtered V\_expressed output.
<sup>¶</sup> IgDiscover tentatively inferred IGHV1-2\*04 in this data set but the allele did not feature in the more strictly filtered V\_expressed output.

Data sets representing the IGHV transcriptome of the 35 subjects, for which haplotyping based on expression of two different alleles of IGHJ6 was possible, were subjected to IMGT/HighV-QUEST analysis. Transcripts derived from rearrangements involving IGHV1-2*05 were present in one haplotype in 6 subjects but always at a level (0.092±0.017% per haplotype of all IGHV-encoding reads) substantially lower than those of IGHV1-2*02 (2.28±0.52% (present in 29 haplotypes)), IGHV1-2*04 (0.63±0.22% (present in 28 haplotypes)), IGHV1-2*06 (1.86±0.8% (present in 6 haplotypes)), or IGHV1-2*07 (2.50% (present in 1 haplotype)). The relatively low levels of transcripts of IGHV1-2*04 in some data sets had resulted in its exclusion from the final genotype proposed by IgDiscover (Table 1) through the standard filtering feature used in this study. If both haplotypes of IGHV1-2 were occupied by alleles other than IGHV1-2*05, only very few reads (0.0015±0.0026%) associated to this allele were observed. Thus, reads associated to IGHV1-2*05 were rarely identified in these data sets as a consequence of antibody evolution, or PCR or sequencing errors of reads originating from rearrangements involving other alleles of IGHV1-2. The frequency of such background reads were, however, about 10-fold higher in subjects that harbored IGHV1-2*06 (the allele of IGHV1-2 that is most similar to IGHV1-2*05) in the genotype (0.0062±0.003%)%, as compared to those that only had other alleles of IGHV1-2 (0.0005±0.0007%), illustrating the effect that other alleles may have on the analysis of rarely expressed genes/alleles. Although the number of reads of IGHV1-2*05 was low, it was in all cases observed that reads encoding IGHV1-2*05 were primarily associated to the IGHJ6-defined haplotype not associated to reads of the other allele found in the subject under investigation (Figure 1A), further supporting the validity of the observation of these reads with an origin in rearrangements that involved IGHV1-2*05. Altogether, these observations support the presence of IGHV1-2*05 in 6/70 haplotypes although the allele had not been inferred by TIgGER or IgDiscover. The presence of IGHV1-2*05 in these genotypes, argues that the gene had not been deleted in any of the investigated haplotypes.

**Figure 1.**
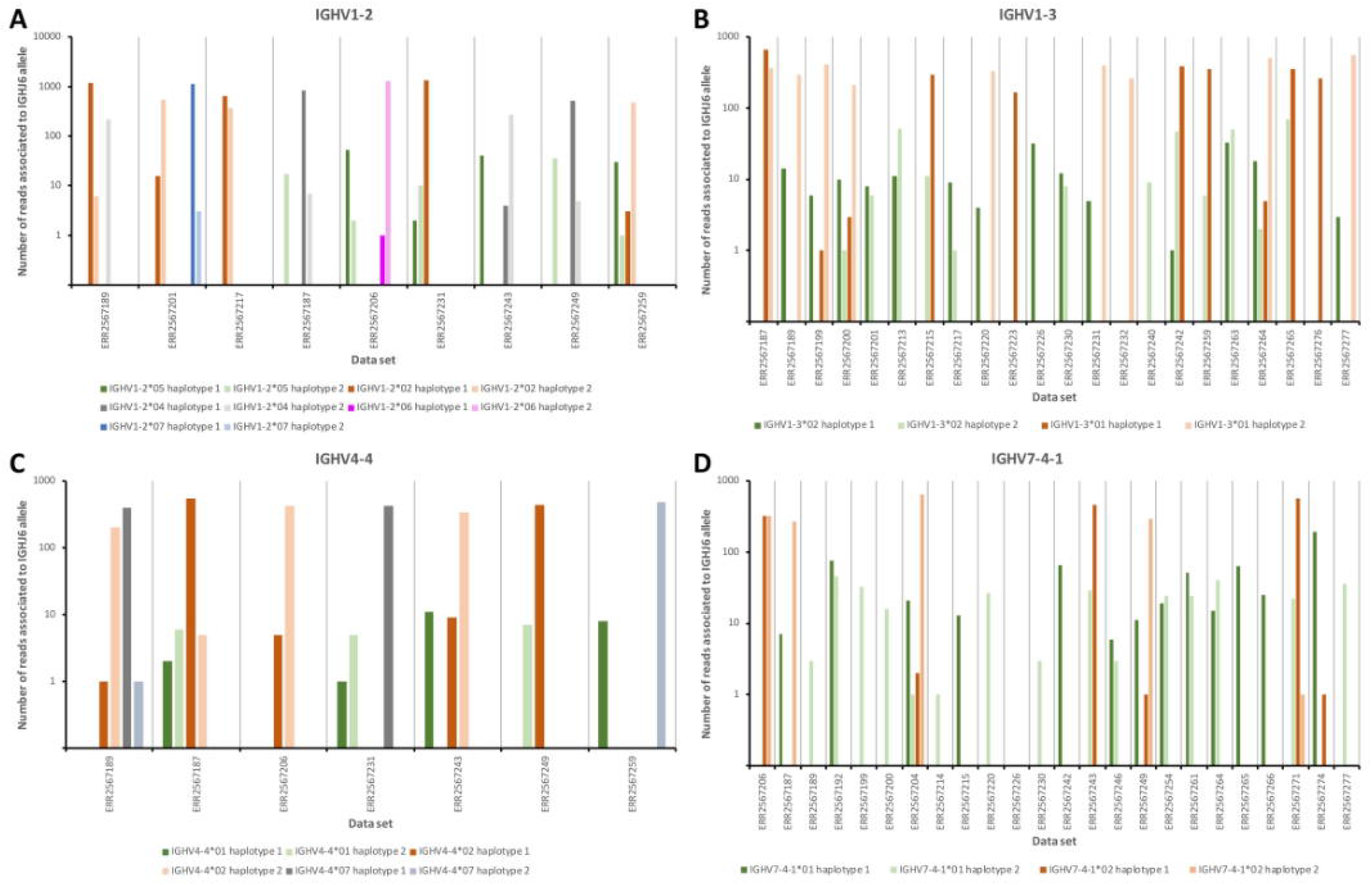
The number of reads of alleles likely present in the germline repertoire of 35 subjects (Table 1) associated to the two different alleles of IGHJ6 of the genotype. **A.** 3 data sets that express 1-2 alleles of IGHV1-2 other than IGHV1-2*05 (left part of panel), and six data sets that express IGHV1-2*05. **B.** One data sets that is homozygous for IGHV1-3*01 (left part of panel), and 21 data sets that express IGHV1-3*02, 19 which also contain such reads associated to IGHJ6. **C.** One data sets that expresses two different alleles of IGHV4-4 other than IGHV4-4*01 (left part of panel), and six data sets that express IGHV4-4*01, five which also contain such reads associated to IGHJ6. **D.** One data sets that is homozygous for IGHV7-4-1*02 (left part of panel), and 23 data sets that express IGHV7-4-1*01, 22 which also contain such reads associated to IGHJ6. In all cases, haplotype 1 represents the haplotype with the IGHJ6 allele with the lowest alphanumeric name in the data set in question (in all cases but one (ERR2567242) this allele is IGHJ6*02). Only reads that by IMGT/HighV-QUEST analysis were uniquely associated to a single IGHV allele and a single allele of IGHJ6 were used in the calculation to generate this illustration.

IGHV1-2*05 is, as observed above, expressed at low levels. 68% of the reads derived from IGHV1-2*05 were, however, considered to be productive by IMGT/HighV-QUEST. The IGHV-IGHD-IGHV rearrangements were mostly in-frame (Figure 2A). These observations suggest that many reads derived from IGHV1-2*05 are able to encode a functional product.

**Figure 2.**
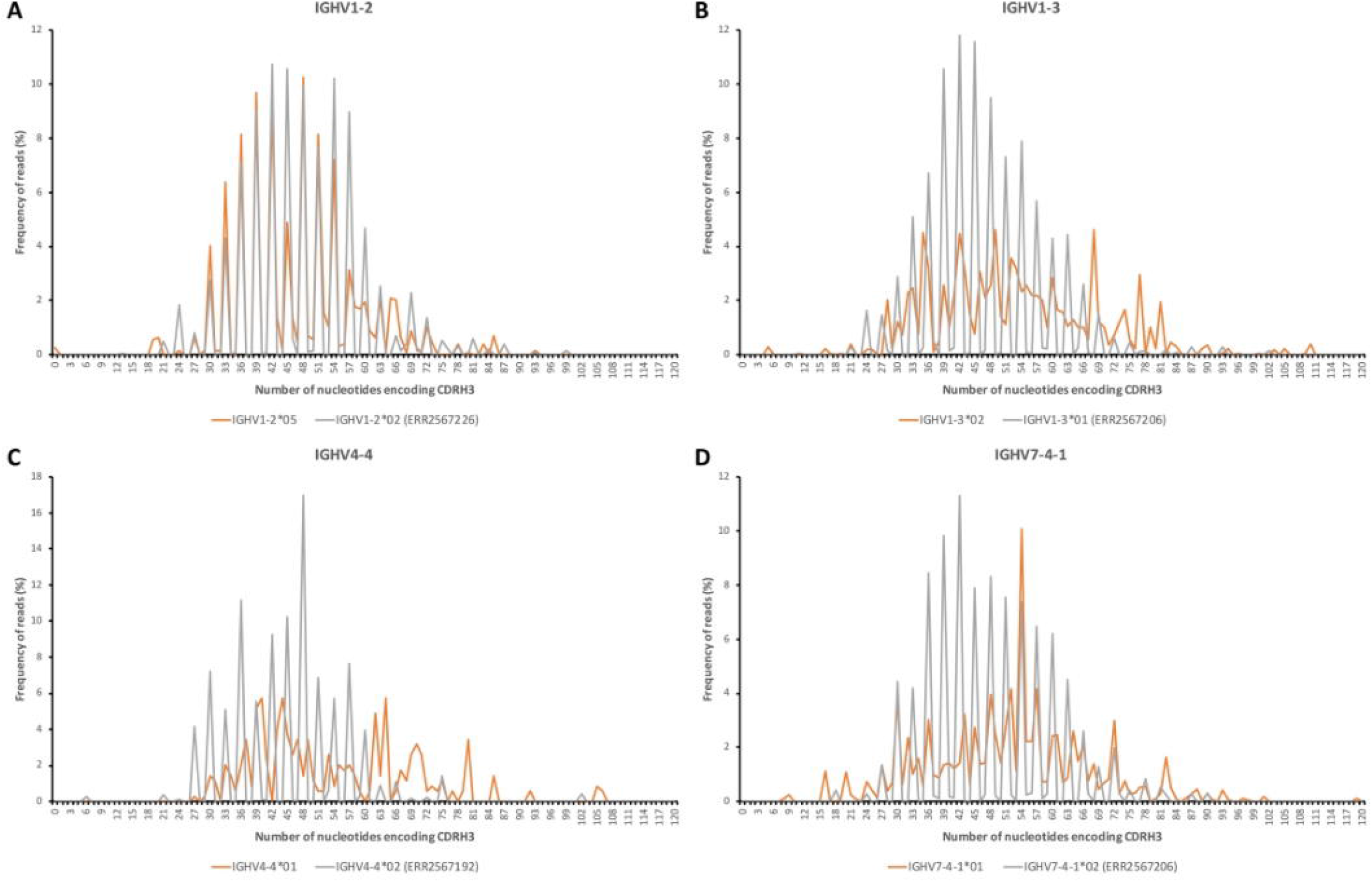
The number of nucleotides of the CDR3-encoding part of reads derived from IGHV1-2*05 (A), IGHV1-3*02 (B), IGHV4-4*01 (C), and IGHV7-4-1*01 (D) extracted from all donor that use these genes. The distribution of lengths of bases are compared to those of IGHV1-2*02 (data set ERR2567226), IGHV1-3*01 (data set ERR2567206), IGHV4-4*02 (data set ERR2567192), and IGHV7-4-1*02 (data set ERR2567206), respectively.

### Germline allele IGHV1-3*02 is expressed at low levels compared to another common allele of this gene

In a past study using the TIgGER inference tool, it was suggested that IGHV1-3 was deleted in 26/66 haplotypes of 33 subjects for which haplotypes had been determined (10). VDJbase indicated that the gene was present as IGHV1-3*01 or IGHV1-3*01 T35A (now officially recognized as IGHV1-3*05) in 77/96 subjects of this dataset. No instances of IGHV1-3*02 had been inferred. The Ensembl database (19) however suggests that bases 6, 12, 167, 208, 291, and 296, bases indicative of IGHV1-3*02, are all present at a frequency of about 40% in many populations, including in European populations (Supplementary Figure 4). Reanalysis of publicly available sequencing data sets by IgDiscover indicated that IGHV1-3*02 was present in the genotype of 11/98 subjects (in all cases without simultaneous detection of IGHV1-3*01). Further analysis, using IMGT/HighV-QUEST, of the 35 datasets that could be haplotyped using haplotype-defining allelic variation in IGHJ6 indicated that 12/35 datasets were homozygous with respect to IGHV1-3*01. Reads derived from rearrangements involving IGHV1-3*01 were present at a frequency of 1.78±0.15% in these subjects. The frequency of reads assigned to IGHV1-3*02 in these data sets was, as expected, very low, 0.0007±0.0009%. Thus, reads associated to IGHV1-3*02 had not been artificially generated in these data sets through antibody evolution, or through PCR or sequencing errors of reads originating from rearrangements involving IGHV1-3*01. Subjects (n=16) that had IGHV1-3*01 associated to only one of its haplotypes expressed such transcripts at a frequency of 0.90±0.22%. Reads associated to IGHV1-3*02 were found in 14 of these subjects at a frequency of 0.035±0.017%, i.e. a level of reads 50-fold higher that observed in subjects that were homozygous for IGHV1-3*01 and thus could not express IGHV1-3*02 unless the gene would have been duplicated. Subjects (n=7) that did not express antibody-encoding genes derived from IGHV1-3*01 (frequency 0.002±0.002% of all reads), also expressed sequences derived from IGHV1-3*02 (0.065±0.19%). In all subjects but two these transcripts were associated to both haplotypes (Figure 1B), although the numbers of transcripts useful for haplotype analysis was, given the number of sequences in these data set, very low. Altogether, IGHV1-3*01 was associated to 40/70 haplotypes while IGHV1-3*02 was associated to at least 26/70 haplotypes. Only 4/70 haplotypes (defined by reads associated to either allele of IGHJ6) could not be identified in any reads, possibly suggesting a deletion of IGHV1-3.

In contrast to reads derived from genes rearranged from IGHV1-3*01 that were mostly (for instance 83% and 84% in ERR2567187 and ERR2567204, respectively) productive, reads derived from transcripts with an origin in IGHV1-3*02 were commonly non-productive. Of 1797 reads that also contained a detectable IGHJ sequence, only 14% were, based on the nucleotide sequence alone, considered to be productive. Many rearrangements were not in frame as evidenced by the length of the sequence that encoded CDR3 (Figure 2B). Thus most observed transcripts derived from IGHV1-3*02 cannot encode a functional product.

### Germline allele IGHV4-4*01 is expressed at low levels compared to other common alleles of this gene

In a past study using the TIgGER inference tool, it was suggested that IGHV4-4 was deleted in 18/94 genotypes and at least 29/66 haplotypes of 33 subjects (10). However, a recent analysis of the material, as presented in VDJbase, suggests that IGHV4-4 is not absent in any of the genotypes of this set (IGHV4-4*02 and IGHV4-4*07 are found in 78/96 and 63/96 genotypes, respectively). However, it is reported to be absent in 6/68 haplotypes of haplotypable genomes, while IGHV4-4*02 and IGHV4-4*07 are found in 36/68 and 26/68 haplotypes, respectively (Supplementary Figure 1). IGHV4-4*01 is not suggested by VDJbase to be expressed in any of the 96 subject for which data is available in the database. In the present study, IGHV4-4*01 was not inferred by IgDiscover in any of the 98 data sets. IGHV4-4*02 and IGHV4-4*07 were found in 37 and 27 haplotypes of 35 subjects that could be haplotyped based on differential expression of alleles of IGHJ6 (Table 1; Supplementary Figure 2). In subjects with alleles of IGHV4-4*02 and/or IGHV4-4*07 in both haplotypes the frequency of reads assigned by IMGT/HighV-QUEST to IGHV4-4*01 was, as expected, very low (0.0007±0.0007%). When assessing individual reads from the 6 subjects that did not carry IGHV4-4*02 or IGHV4-4*07 on one of its haplotypes, it was possible to find evidence of transcripts of IGHV4-4*01 at a statistically significant higher level (0.026±0.07%) than if both sites of IGHV4-4 were occupied by any of the other alleles (p=0.0001 (Mann-Whitney onesided test)). In 5/6 subjects a few reads with an origin in a rearrangement utilizing IGHV4-4*01 associated to IGHJ6 were found. These appropriately associated to the allele of IGHJ6 that was not used by the other allele of IGHV4-4 present in the genome (Figure 1C). The frequency of transcripts derived from IGHV4-4*01 when found in the genome was, however, estimated to be 38-fold and 37-fold lower than the frequency of transcripts derived from a single copy of IGHV4-4*02 and IGHV4-4*07, respectively. Collectively the data suggests that IGHV4-4*01 is present in many haplotypes that do not express other common alleles of this gene but that its level of expression is low.

Only 7% of reads assigned to IGHV4-4*01 were considered by IMGT/HighV-QUEST to represent productive sequences, with a large number of sequences having out-of frame IGHV-IGHD-IGHJ rearrangements (Figure 2C). Thus most observed transcripts derived from IGHV4-4*01 cannot encode a functional product.

Alleles of IGHV4-4, IGHV4-59, and IGHV4-61 are in many cases similar and their exact location in the genotype may not always be known. One such example is IGHV4-59*08 that often appears to reside in gene IGHV4-61 (20). The inferred genotype of these genes were identified by IGHJ6-based haplotyping in the six samples that encoded IGHV4-4*01. In all cases, alleles of IGHV4-59 and IGHV4-61 were assigned to both haplotypes of these subjects (data not shown). It is thus hypothesized that IGHV4-4*01 was indeed located to gene IGHV4-4 on one of each subjects’ haplotypes.

Sequence features of IGHV4-4*01 at base 46 (C) and 308 (G) differentiate this allele from most alleles of IGHV4-4, IGHV4-59 and IGHV4-61 (Supplementary Figure 5). SNP analysis suggest that these variants are present at a frequency of 3-4% in European populations and at a higher frequency in many other populations (Supplementary Figure 5). Altogether, this data lend support to the likely identification of IGHV4-4*01 in large datasets like the one investigated here.

### Germline allele IGHV7-4-1*01 is expressed at low levels compared to another common allele of this gene

In a past study using the TIgGER inference tool, it was suggested that IGHV7-4-1 was deleted in 60/94 genotypes, and at least 49/66 (74%) haplotypes of 33 subjects (10). A recent analysis of the material, as presented in VDJbase, suggests that IGHV7-4-1 is deleted in 56/96 genotypes including in 19/34 genotypes that can be haplotyped based on heterozygosity of IGHJ6 (Supplementary Figure 1). Furthermore, VDJbase data shows that IGHV7-4-1*02 is present in 37/96 genotypes including 13 that can be haplotyped (in two of these cases (I15 and I92) the evidence for the presence of this allele shows very low confidence), while IGHV7-4-1*01 is present in only 9/96 genotypes, two of which can be haplotyped. This contrast to SNP analysis that suggests that the allele-defining variant of IGHV7-4-1*01 is found at a larger frequency in most populations (Supplementary Figure 6). When reanalyzing these publicly available, haplotypable data sets of project PRJEB26509 (n=35) with IgDiscover, IGHV7-4-1*02 was inferred at high frequency in 11 transcriptomes, but also IGHV7-4-1*01 at low frequency in 12 transcriptomes (Table 1; Supplementary Figure 2). IGHV7-4-1*01 was only inferred in transcriptomes that did not simultaneously express IGHV7-4-1*02.

To better understand the occurrence of IGHV7-4-1*01, data sets were individually analyzed using IMGT High/V-QUEST. One sample (ERR2567206) encoded transcripts derived from IGHV7-4-1*02 from both haplotypes. The level of transcripts perceived as originating from IGHV7-4-1*01 in this samples was as expected very low (0.0015% of all reads). 10 samples carried IGHV7-4-1*02 on one of its haplotypes. 11 samples that expressed genes derived from IGHV7-4-1*02 from none or one of its haplotypes expressed similarly low levels (0.001±0.001%) of IGHV7-4-1*01-derived transcripts. Higher levels (0.056±0.051%) of IGHV7-4-1*01-derived transcripts were seen in 23/35 samples. The frequency of transcripts of IGHV7-4-1*02 from a single copy of the allele was, however, significantly higher (p<0.0001 (Mann-Whitney one-sided test)) and determined to represent 0.96±0.32% of all reads. Although the number of reads was low, IGHV7-4-1*01 was suggested to associate to at least 28/70 haplotypes (Figure 1D). Such reads could be assigned to one or both haplotypes in a subject except in one data set as no reads were associated to IGHJ6. Given the low number of reads associated to IGHV7-4-1*01 it is conceivable that additional haplotypes may carry the allele although no rearrangements to IGHJ6 were found in the transcriptome to confirm such an association. The frequency of IGHV7-4-1/haplotype, taking both IGHV7-4-1*01 and IGHV7-4-1*02 into account, was thus at least 57%. If IGHV7-4-1*02 was expressed in the same sample as IGHV7-4-1*01, the two alleles were as expected, by IGHJ6-based haplotyping, not primarily associated to the same haplotype whenever rearrangements of IGHV7-4-1*01 associated to IGHJ6 were actually found in the data (Figure 1D). Among the 34 samples, only 6 (17%) showed no evidence of expression of either IGHV7-4-1*01 or IGHV7-4-1*02. Altogether, in-depth analysis of the underlying data identify expression of IGHV7-4-1*01 in multiple samples, expression that is not always readily detected by inference technology.

Assessment by IMGT/HighV-QUEST of the perceived functionality of rearranged sequences derived from IGHV7-4-1*01 suggested that 31% of reads that also contained a IGHJ-derived sequence were perceived as being productive and featured an in-frame IGHV-IGHD-IGHJ rearrangement (Figure 2D). To further assess the somatic evolution of rearrangements derived from IGHV7-4-1*01, NGS data sets of IgA and IgG repertoires of two subjects known to express IGHV7-4-1*01 but not IGHV7-4-1*02 (11,15) were investigated. These two genes differ in only one base in the part of the gene that encode the final protein product. IGHV7-4-1*01 encodes an unusual, likely surface-exposed, cysteine residue at position 92 (C92), a residue located far from an antibody’s paratope (Supplementary Figure 7). The number of reads and clones with an origin in IGHV7-4-1*01 was low, but it was observed that all productive rearrangements of these data sets (Supplementary Figure 8) had mutated the germline-encoded C92 to a range of other residues (primarily serine, proline, tyrosine, histidine, leucine, pheylalanine, and asparagine). There thus appears to be a strong driving force to replace the unusual cysteine residue of IGHV7-4-1*01 with another residue during somatic hypermutation and selection, despite the fact that codon 92 does not carry a RGYW/WRCY (21) or a WA/TA mutational hotspot (22).

### IGHV1-2*05 and IGHV4-4*01 are expressed on the same haplotype

Both IGHV1-2*05 and IGHV4-4*01 were each identified in six genotypes (Table 1, Figure 1A, C). Despite their low overall frequency, these alleles were in all cases found in the very same genotypes. An analysis of the haplotypes confirmed that these alleles are present together on the same haplotype in these six genotypes (Figure 1A, C). The most likely conserved order of genes/alleles in all the six haplotypes of the present study that carry these poorly expressed alleles was: IGHV6-1*01 – IGHV1-2*05 – IGHV1-3*01 – IGHV4-4*01 – IGHV7-4-1*02. The entire associated IGHV locus of these haplotypes were, however, not identical as this set of alleles were associated to alleles like IGHV3-11*01, IGHV3-11*05, or IGHV3-11*06; IGHV3-48*01, IGHV3-48*02, or IGHV3-48*04; IGHV3-49*03, IGHV3-49*04, or IGHV3-49*05; IGHV1-69*01 and IGHV1-69*06, IGHV1-69*02, IGHV1-69*04, or IGHV1-69*10.

## DISCUSSION

Different subjects have access to highly different sets of alleles of germline genes to encode the naïve antibody population they may use to mount humoral immune responses against the hostile environment. Such differences may translate into differences in their ability to raise antibodies against some epitopes (23). Efficient genomic sequencing of the immunoglobulin heavy and light chain loci have been particularly challenging and only lately been facilitated by long read sequencing technology. Consequently, our understanding of the complexity of immunoglobulin germline gene repertoires is limited. Fortunately, computational germline gene inference using information contained in large NGS data sets has allowed us to describe personal germline repertoires of many subjects, and to identify novel immunoglobulin germline gene alleles that were previously not defined (5,10,11,24). A number of such new alleles have been inferred, reviewed, and documented using a standardized procedure (25). However, the immunoglobulin germline gene rearrangement process, somatic hypermutation, and PCR and sequencing artefacts complicate germline gene inference. Consequently, such tools commonly operate under a set of restrictions, to enhance the specificity of the algorithm at the cost of sensitivity. For instance inferred alleles that are expressed at much lower levels than other alleles of the same gene in the genotype will commonly be removed during the process.

It has been established by genomic sequencing that gene insertion and deletion events as well as more complex events has generated highly different human immunoglobulin haplotypes. Recently the genotypes of almost 100 subjects were defined by an inference process (10). Importantly, haplotypic analysis could be performed on many of these data sets and multiple deletion events could be defined. As inference cannot efficiently detect alleles that are poorly expressed, an in-depth analysis of the such sequences was now performed with a focus on genes located in proximity to the IGHD locus, genes that had been suggested to be deleted in several haplotypes (10). Such analysis demonstrated the existence of reads providing evidence of alleles expressed at a level much lower than other alleles of the same gene. Such reads were not present in transcriptomes from genotypes that were known to have other alleles populating these gene sites. The existence of such alleles were in several cases supported by SNPs described through population studies. Such studies are commonly based on short read sequencing, a technology that suffers from substantial caveats in particular in relation to sequencing of immunoglobulin loci (26). Nevertheless, it is reassuring that SNP analysis support the existence of these alleles in samples obtained in Europe. Altogether, the common presence of the IGHV1-2*05, IGHV1-3*02, IGHV4-4*01, and IGHV7-4-1*01 alleles in a substantial fractions of haplotypes is now well established despite the fact that their level of expression is low. The results suggests that deletion at the genomic level is less common than that envisaged from past analysis (10).

Although the alleles IGHV1-2*05, IGHV1-3*02, IGHV4-4*01, and IGHV7-4-1*01 are present at substantial frequencies in a population, they may, due to their low level of expression, have little impact on the diversity of the antibody repertoire. It is not known if their limited expression is a result of an impairment in their ability to participate in the gene rearrangement processes, or to encode a functional product. It is now hypothesized that several of these alleles have limited ability to form a functional protein following rearrangement. IGHV1-3*02 encodes products that carry residues S56, E70, and N99. Virtually all other germline genes of the IGHV1 subgroup encode products with I56 (or another hydrophobic residue), K70, and T99. Similarly IGHV1-2*05 encodes an atypical residue V100 in the domain’s lower core, IGHV7-4-1*01 encodes an atypical C92, and IGHV4-4*01 encodes atypical residues P16 and C103. With access two NGS data sets defining IgG and IgA repertoires of two subjects previously shown to carry IGHV7-4-1*01 but not the more commonly expressed IGHV7-4-1*02 in the genotype, it was possible to assess how the immune system approached evolution of the allele encoding an unusual cysteine at position 92, far from the antibodies paratope. Indeed, this residue was invariably substituted in productively rearranged and mutated reads. There thus seems to be a strong incentive to remove this unusual residue during the hypermutation process. It is hypothesize that C92 compromise the structure or stability of the protein product and that its substitution improves the functionality of the encoded product and thereby its ability to get an upper hand during the selection process. This agrees with past findings of evolution of IGHV1-18*01, IGHV1-8*01 and IGHV5-51*01 that all carry unusual residues in their framework regions, residues that are also substituted at a high frequency during somatic hypermutation (27).

Most reads of IGHV1-3*02 and IGHV4-4*01 originated from rearranged genes that would not encode a full length product, for instance as they carried out-of-frame rearrangements or stop codons. It is conceivable that the level of functionality of these alleles should be modified from Functional to ORF. It seems possible that many of the reads represent rearrangements that are merely passengers in cells that use a rearrangement made on the other chromosome to encode an antibody. Nevertheless, their existence highlights the fact that these alleles are present in the haplotypes of many subjects although specific measures have to be taken if they are to be detected by germline gene inference. Importantly, their presence may aid our understanding of the antibody hypermutation process as they may serve as controls that have not undergone selection based on retained or improved antigen-binding properties. Furthermore, as we now realize the low level of transcription of some of these alleles, it will be possible to design computational approaches to correctly assign reads derived from such alleles even in the presence of other, more highly expressed alleles, through an allele-specific filtering strategy.

In summary, IGHV1-2*05, IGHV1-3*02, IGHV4-4*01, and IGHV7-4-1*01 are commonly present in human genotypes but they are poorly expressed and at least two of them do not commonly encode functional products. The identification of these alleles in many data sets through inference requires that computational processes are properly adapted to that task.

## DATA AVAILABILITY

Raw sequence data is available from the European Nucleotide Archive as project PRJEB26509. TIgGER-calculated genotypes and haplotypes are available from VDJbase (www.vdjbase.org).

## Supplementary Figure Legends

**Supplementary Figure 1.** Visualization of genotypes and haplotypes of 34 of the samples of the P1 study, also assessed in the present analysis, as defined by TIgGER inference technology and as displayed on the VDJbase website (http://vdjbase.org) (9). Illustrations were retrieved in July, 2020. This information was provided under a CC0 (Creative Commons 0) license.

**Supplementary Figure 2.** Visualization of 35 genotypes of subjects for which haplotyping, based on heterozygosity of IGHJ6, is possible, as inferred by IgDiscover technology (6).

**Supplementary Figure 3.** Allelic variants of IGHV1-2 as defined by IMGT are illustrated. Variability of some of the positions of these genes in samples obtained in different geographical locations as illustrated by the ENSEMBL browser (release 101, August 2020) (19) is shown. Only bases 163, 223, and 299 (IMGT numbering nomenclature (16)) of this gene display frequencies of variation >1% in the 1000 Genomes Project. The variant (SNP rs12588974) at base 299, indicative of the IGHV1-2*01 or IGHV1-2*05 alleles is present at about 5% in European populations. Bases 233 and 234 (SNPs rs782139757 and rs1425538657), that separate these two alleles remains as T and G, respectively, at very high frequency in most populations suggesting that IGHV1-2*01 is not common in these populations (not shown). All sequence variants of the illustrations of SNPs are indicated as seen in the reversed strand, hence they are complementary to the base of the coding strand.

**Supplementary Figure 4.** Allelic variants of IGHV1-3 as defined by IMGT are illustrated. Variability of some of the positions of these genes in samples obtained in different geographical locations as illustrated by the ENSEMBL browser (release 101, August 2020) (19) is shown. Only bases 6, 12, 167, 208, 291 and 296 (IMGT numbering nomenclature (16)) of this gene display frequencies of variation >1% in the 1000 Genomes Project. Variants indicative of the IGHV1-3*02 allele are present at about 40%. All sequence variants of the illustrations of SNPs are indicated as seen in the reversed strand, hence they are complementary to the base of the coding strand.

**Supplementary Figure 5.** Allelic variants of IGHV4-4 as defined by IMGT are illustrated. Variability of some of the positions of these genes in samples obtained in different geographical locations as illustrated by the ENSEMBL browser (release 101, August 2020) (19) is shown. Analysis of this gene is complicated by extensive similarity with alleles of IGHV4-59 and IGHV4-61, alleles of which are also shown. A few of the positions of IGHV4-4 that display frequencies of variation >1% in all populations in the 1000 Genomes Project are shown. Note that variants at bases 46 and 308 (IMGT numbering nomenclature (16)), indicative of the IGHV4-4*01 allele are present at about 3-4% in European populations. All sequence variants of the illustrations of SNPs are indicated as seen in the reversed strand, hence they are complementary to the base of the coding strand.

**Supplementary Figure 6.** Allelic variants of IGHV7-4-1 as defined by IMGT are illustrated. Variability of some of the positions of these genes in samples obtained in different geographical locations as illustrated by the ENSEMBL browser (release 101, August 2020) (19) is shown. Sequence variation in base 274 (IMGT numbering nomenclature (16)) suggests that the base associated to IGHV7-4-1*01 is more common than the base associated to other alleles of this gene in most populations. All sequence variants of the illustrations of SNPs are indicated as seen in the reversed strand, hence they are complementary to the base of the coding strand.

**Supplementary Figure 7.** High resolution structures of five antibodies with a heavy chain variable domain encoded by a gene with an origin in IGHV7-4-1. Heavy chain CDR3 is shown at the top of each sequence in red. The side chain of residue 92 (in all these cases a serine), located far from the antibody binding site is shown in green (carbon) and red (oxygen). Structures include PDB entries 4D9Q, (A), 4EOW (B), 5CGY (C), 5ZMJ (D), and 6B5R (E).

**Supplementary Figure 8.** Translated sequences of productive IgA and IgG-encoding reads derived from NGS data sets of two subjects (donors 2 and 4) that both have access to IGHV7-4-1*01 but not IGHV7-4-1*02 to generate an antibody repertoire (11). The sequencing protocol (15) allowed for determination of the sequence from the end of framework 1 and extended into the first constant domain of the heavy chain. The sequences encoded by IGHV7-4-1*01 and IGHV7-4-1*02 are shown on top of the figure. Residue 92 is highlighted by an arrow.

