## Supplementary Figure 1 for "Poorly expressed alleles of several human immunoglobulin heavy chain variable (IGHV) genes are common in the human population"

**Supplementary Figure 1.** Visualization of genotypes and haplotypes of 34 of the samples of the P1 study, also assessed in the present analysis, as defined by TIgGER inference technology and as displayed on the VDJbase website (http://vdjbase.org) (9). Illustrations were retrieved in July, 2020. This information was provided under a CC0 (Creative Commons 0) license.

### Genotype and haplotype of VDJbase sample I10 (study P1)

(illustrations downloaded from VDJbase in July, 2020)

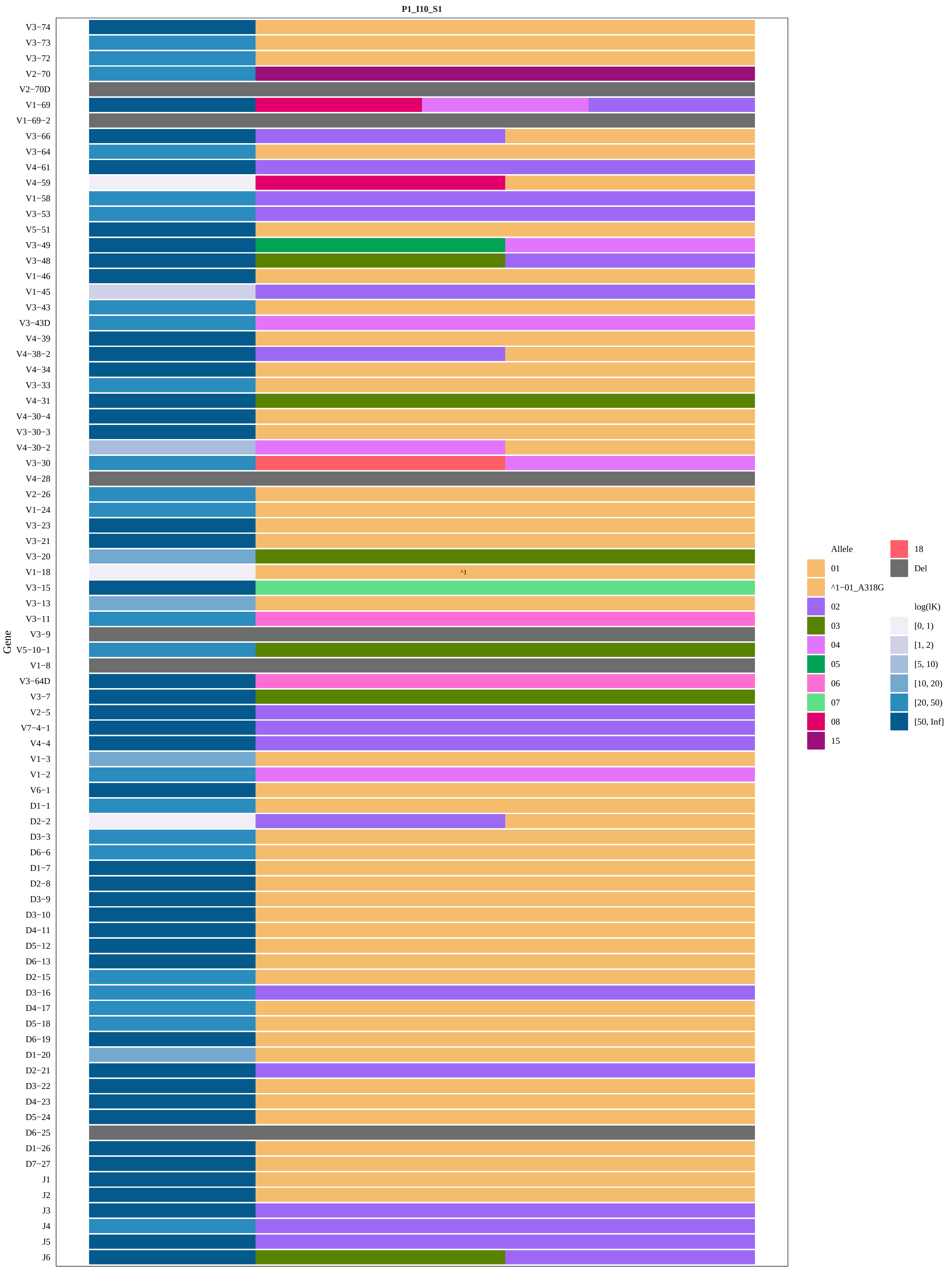
P1_I10_S1
Gene
V3−74
V3−73
V3−72
V2−70
V2−70D
V1−69
V1−69−2
V3−66
V3−64
V4−61
V4−59
V1−58
V3−53
V5−51
V3−49
V3−48
V1−46
V1−45
V3−43
V3−43D
V4−39
V4−38−2
V4−34
V3−33
V4−31
V4−30−4
V3−30−3
V4−30−2
V3−30
V4−28
V2−26
V1−24
V3−23
V3−21
V3−20
V1−18
V3−15
V3−13
V3−11
V3−9
V5−10−1
V1−8
V3−64D
V3−7
V2−5
V7−4−1
V4−4
V1−3
V1−2
V6−1
D1−1
D2−2
D3−3
D6−6
D1−7
D2−8
D3−9
D3−10
D4−11
D5−12
D6−13
D2−15
D3−16
D4−17
D5−18
D6−19
D1−20
D2−21
D3−22
D4−23
D5−24
D6−25
D1−26
D7−27
J1
J2
J3
J4
J5
J6
^1
Allele
01
^1−01_A318G
02
03
04
05
06
07
08
15
18
Del
log(lK)
[0, 1)
[1, 2)
[5, 10)
[10, 20)
[20, 50)
[50, Inf]

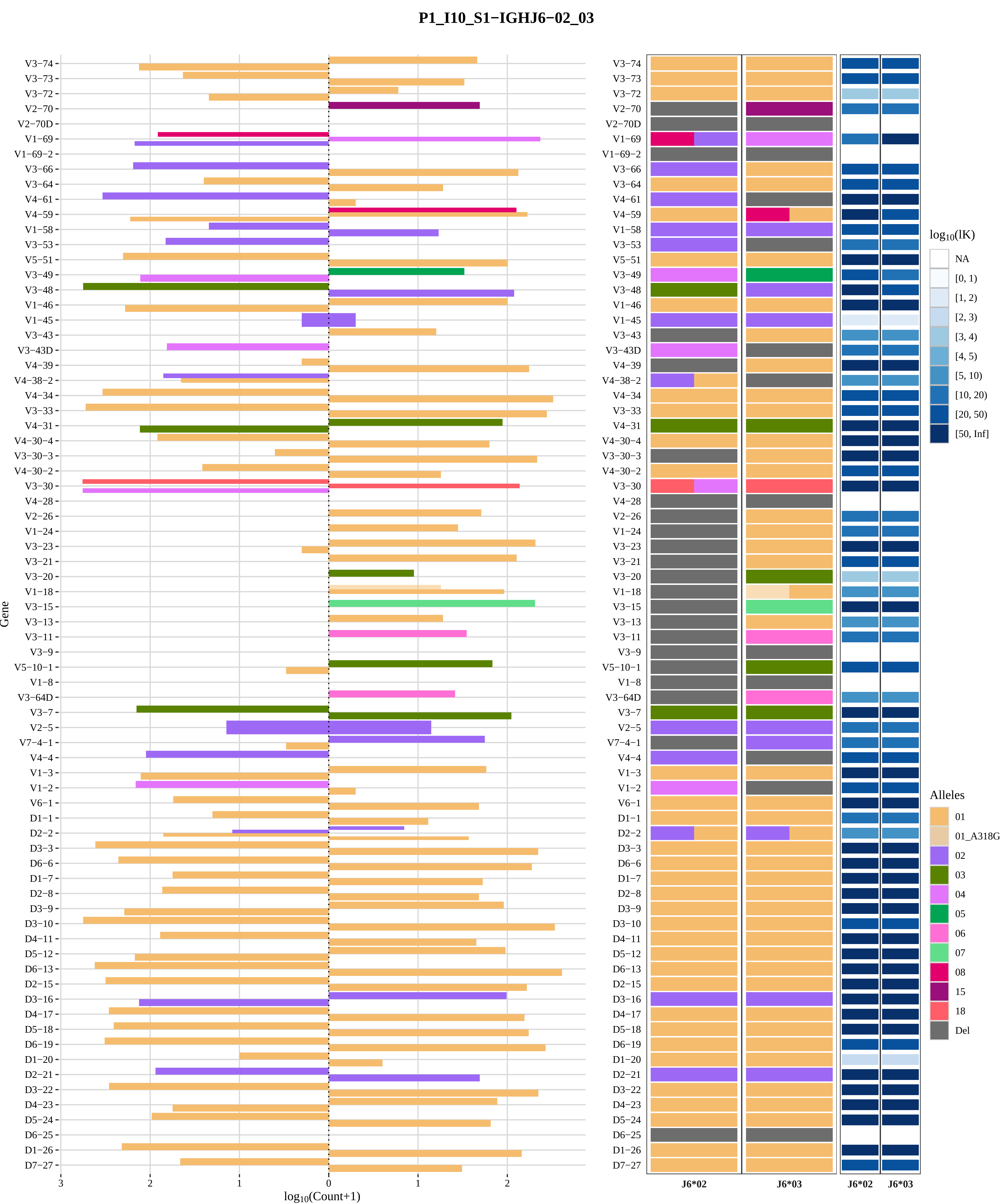
P1_I10_S1−IGHJ6−02_03
Gene
V3−74
V3−73
V3−72
V2−70
V2−70D
V1−69
V1−69−2
V3−66
V3−64
V4−61
V4−59
V1−58
V3−53
V5−51
V3−49
V3−48
V1−46
V1−45
V3−43
V3−43D
V4−39
V4−38−2
V4−34
V3−33
V4−31
V4−30−4
V3−30−3
V4−30−2
V3−30
V4−28
V2−26
V1−24
V3−23
V3−21
V3−20
V1−18
V3−15
V3−13
V3−11
V3−9
V5−10−1
V1−8
V3−64D
V3−7
V2−5
V7−4−1
V4−4
V1−3
V1−2
V6−1
D1−1
D2−2
D3−3
D6−6
D1−7
D2−8
D3−9
D3−10
D4−11
D5−12
D6−13
D2−15
D3−16
D4−17
D5−18
D6−19
D1−20
D2−21
D3−22
D4−23
D5−24
D6−25
D1−26
D7−27
3
2
1
0
1
2
log10(Count+1)
J6*02
J6*03
J6*02
J6*03
log10(lK)
NA
[0, 1)
[1, 2)
[2, 3)
[3, 4)
[4, 5)
[5, 10)
[10, 20)
[20, 50)
[50, Inf]
Alleles
01
01_A318G
02
03
04
05
06
07
08
15
18
Del

### Genotype and haplotype of VDJbase sample I12 (study P1)

(illustrations downloaded from VDJbase in July, 2020)

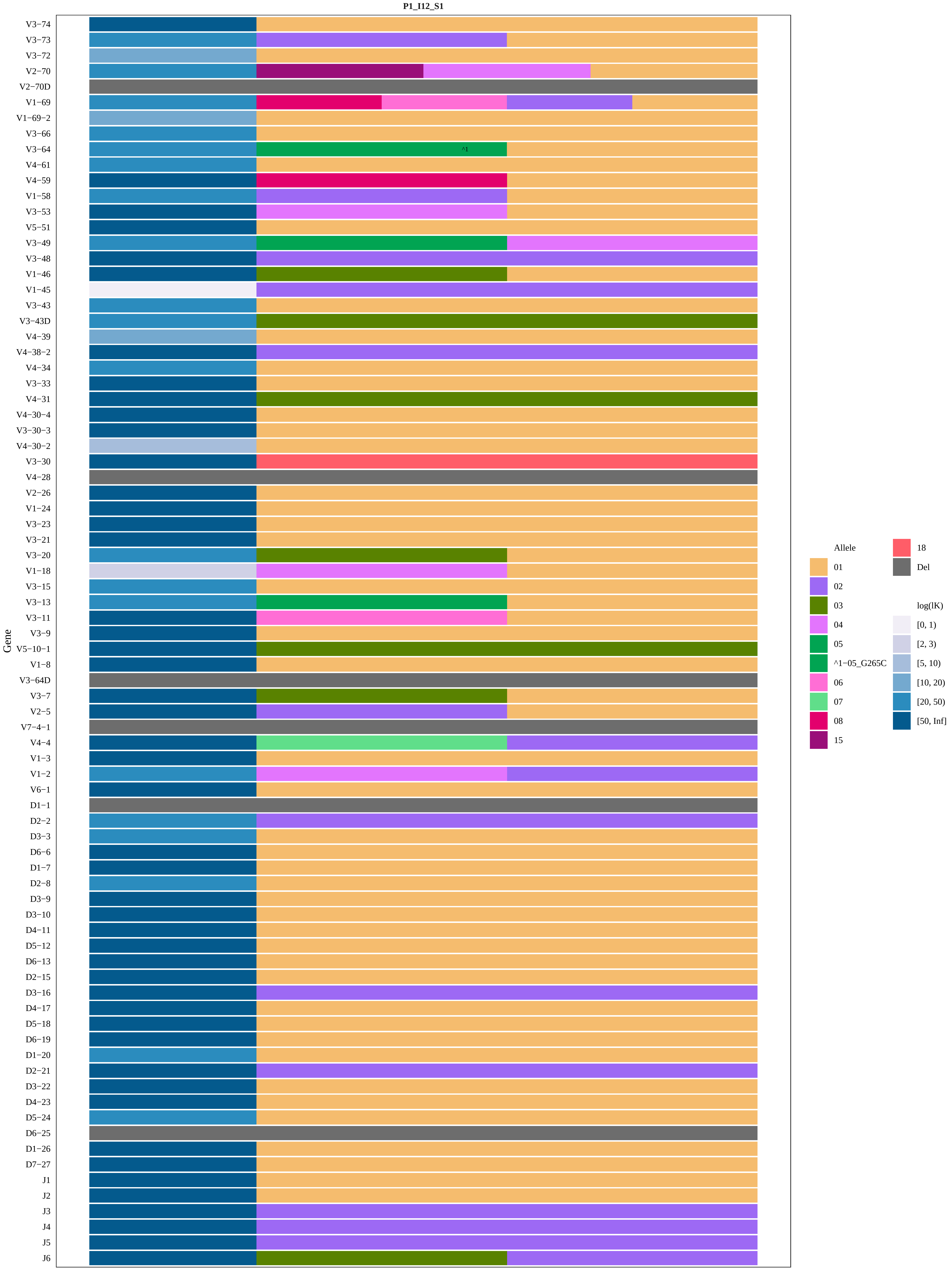

P1_I12_S1
Gene
V3-74
V3-73
V3-72
V2-70
V2-70D
V1-69
V1-69-2
V3-66
V3-64
V4-61
V4-59
V1-58
V3-53
V5-51
V3-49
V3-48
V1-46
V1-45
V3-43
V3-43D
V4-39
V4-38-2
V4-34
V3-33
V4-31
V4-30-4
V3-30-3
V4-30-2
V3-30
V4-28
V2-26
V1-24
V3-23
V3-21
V3-20
V1-18
V3-15
V3-13
V3-11
V3-9
V5-10-1
V1-8
V3-64D
V3-7
V2-5
V7-4-1
V4-4
V1-3
V1-2
V6-1
D1-1
D2-2
D3-3
D6-6
D1-7
D2-8
D3-9
D3-10
D4-11
D5-12
D6-13
D2-15
D3-16
D4-17
D5-18
D6-19
D1-20
D2-21
D3-22
D4-23
D5-24
D6-25
D1-26
D7-27
J1
J2
J3
J4
J5
J6
^1
Allele
01
02
03
04
05
^1-05_G265C
06
07
08
15
18
Del
log(lK)
[0, 1)
[2, 3)
[5, 10)
[10, 20)
[20, 50)
[50, Inf]

### P1_I12_S1−IGHJ6−02_03

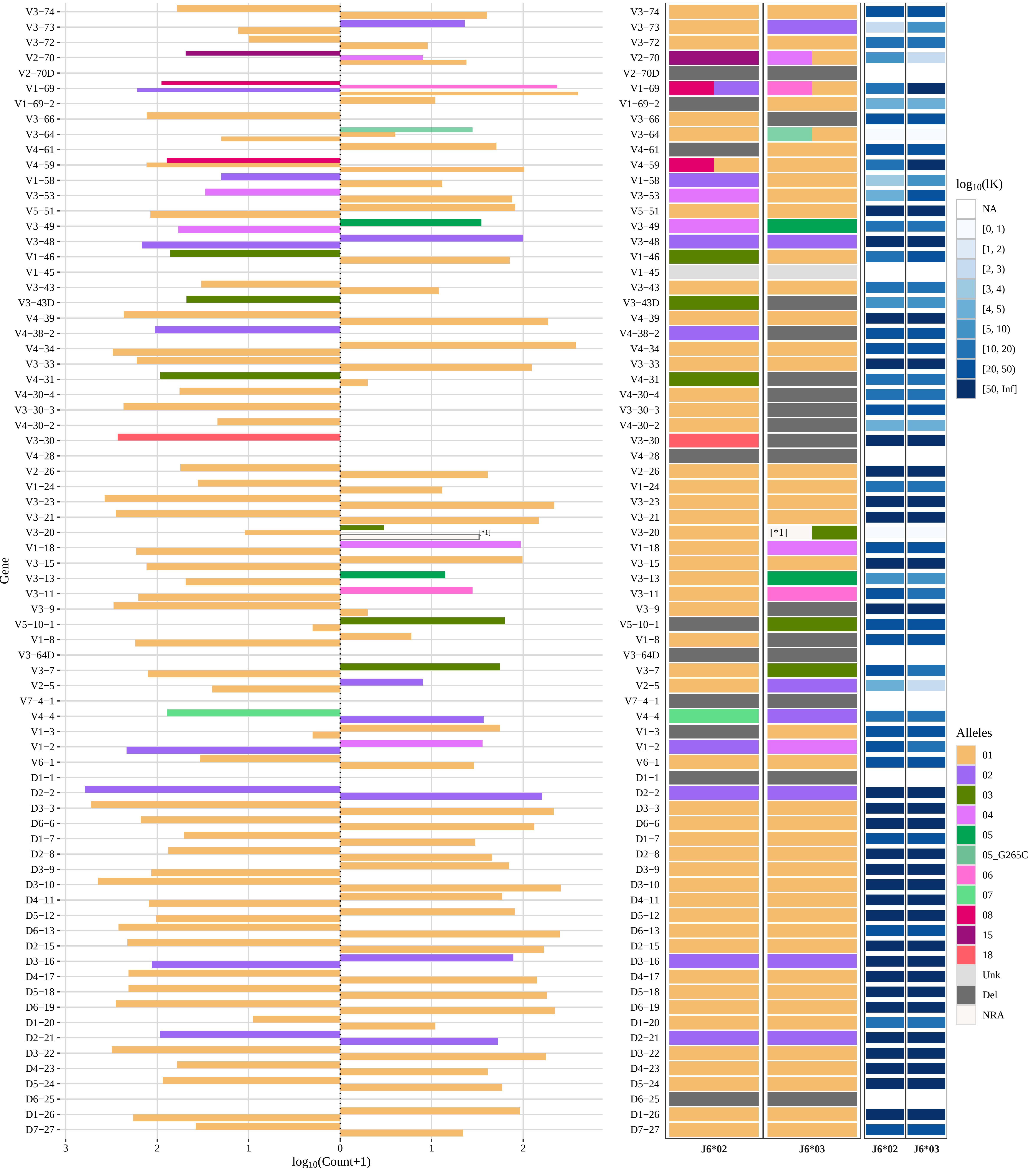


[*1] 01_03

### Genotype and haplotype of VDJbase sample I15 (study P1)

(illustrations downloaded from VDJbase in July, 2020)

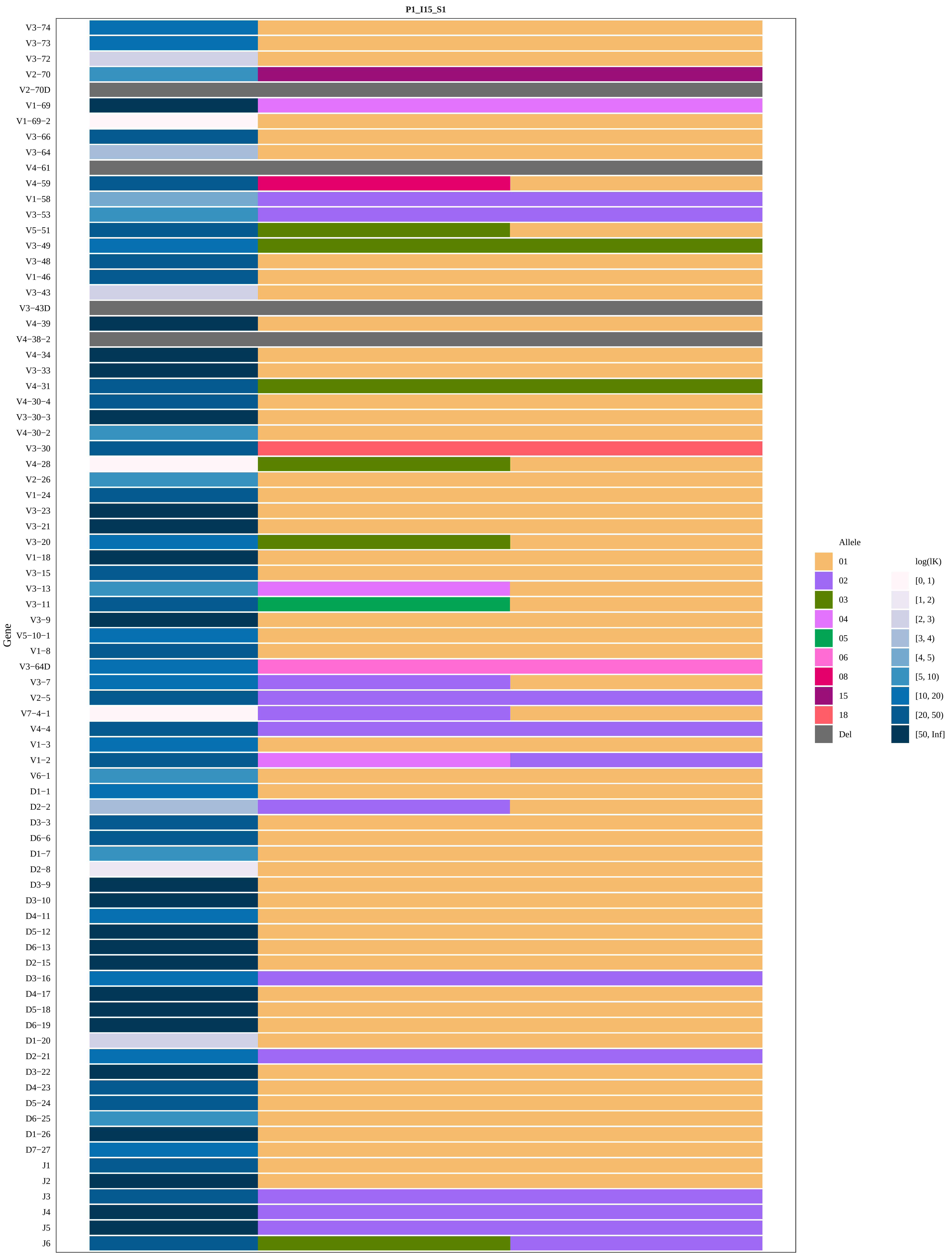

P1_I15_S1
Gene
V3−74
V3−73
V3−72
V2−70
V2−70D
V1−69
V1−69−2
V3−66
V3−64
V4−61
V4−59
V1−58
V3−53
V5−51
V3−49
V3−48
V1−46
V3−43
V3−43D
V4−39
V4−38−2
V4−34
V3−33
V4−31
V4−30−4
V3−30−3
V4−30−2
V3−30
V4−28
V2−26
V1−24
V3−23
V3−21
V3−20
V1−18
V3−15
V3−13
V3−11
V3−9
V5−10−1
V1−8
V3−64D
V3−7
V2−5
V7−4−1
V4−4
V1−3
V1−2
V6−1
D1−1
D2−2
D3−3
D6−6
D1−7
D2−8
D3−9
D3−10
D4−11
D5−12
D6−13
D2−15
D3−16
D4−17
D5−18
D6−19
D1−20
D2−21
D3−22
D4−23
D5−24
D6−25
D1−26
D7−27
J1
J2
J3
J4
J5
J6
Allele
01
02
03
04
05
06
08
15
18
Del
log(lK)
[0, 1)
[1, 2)
[2, 3)
[3, 4)
[4, 5)
[5, 10)
[10, 20)
[20, 50)
[50, Inf]

**P1_I15_S1−IGHJ6−02_03**

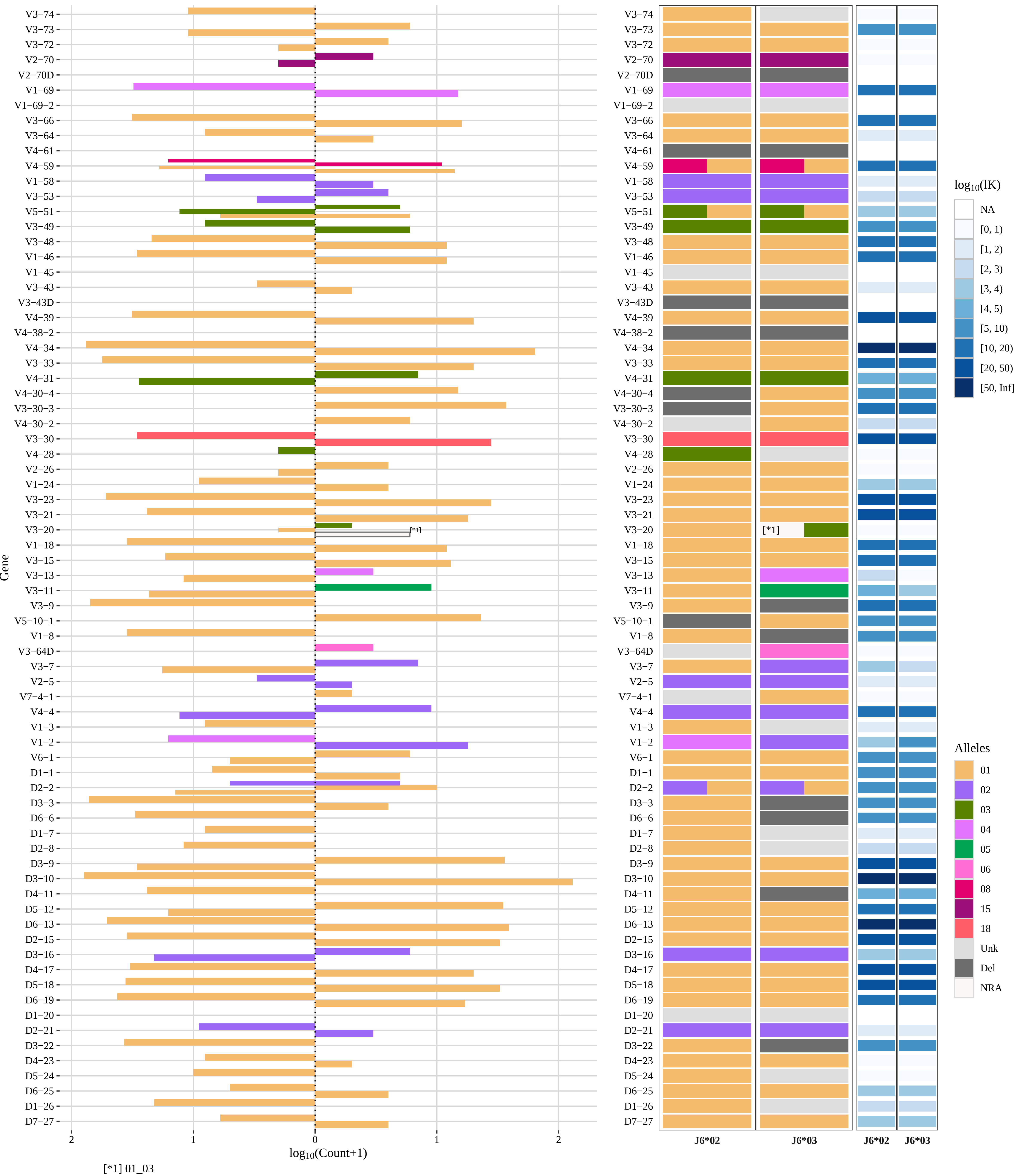

### Genotype and haplotype of VDJbase sample I22 (study P1)

(illustrations downloaded from VDJbase in July, 2020)

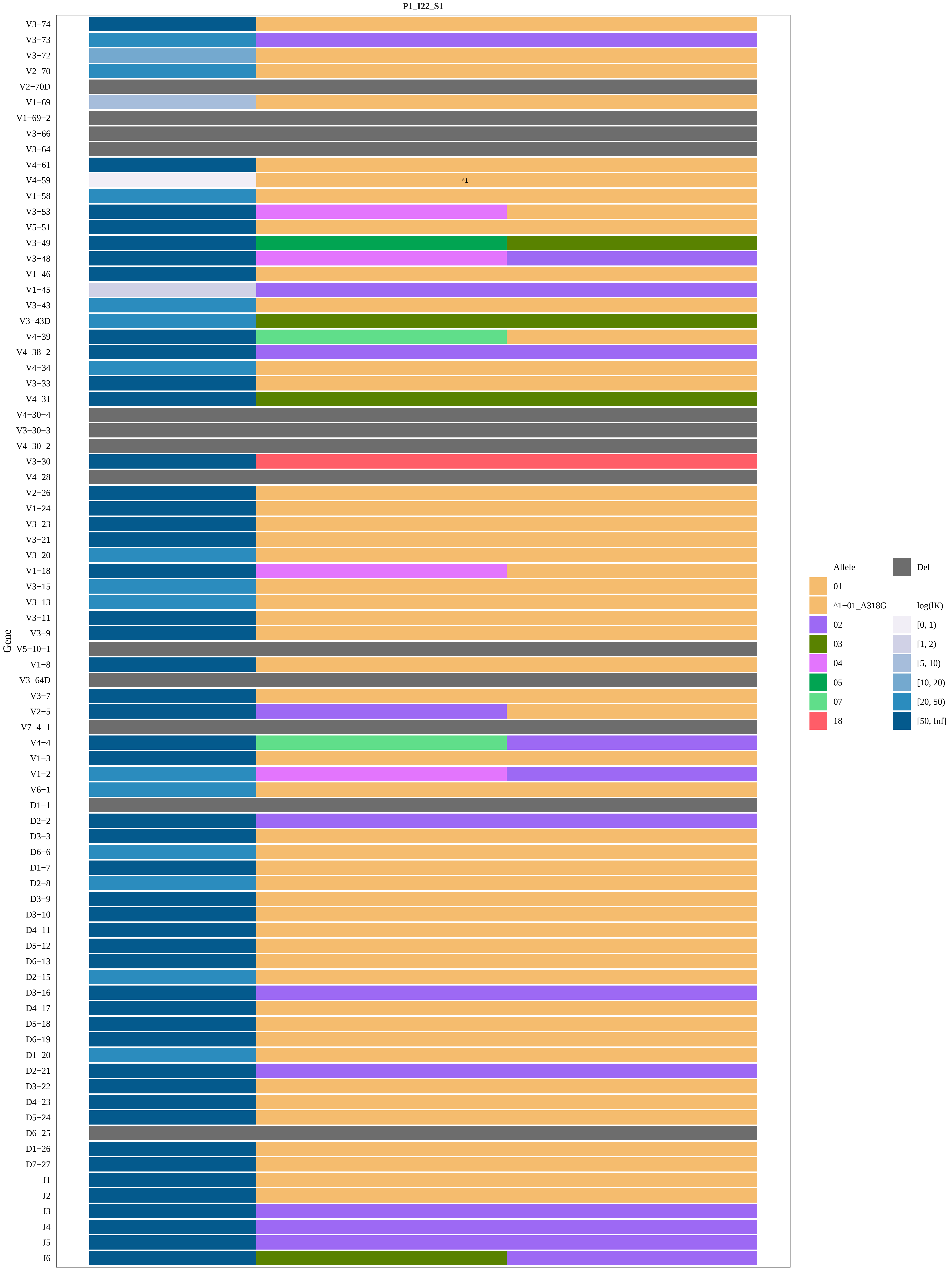
P1_I22_S1
Gene
V3−74
V3−73
V3−72
V2−70
V2−70D
V1−69
V1−69−2
V3−66
V3−64
V4−61
V4−59
^1
V1−58
V3−53
V5−51
V3−49
V3−48
V1−46
V1−45
V3−43
V3−43D
V4−39
V4−38−2
V4−34
V3−33
V4−31
V4−30−4
V3−30−3
V4−30−2
V3−30
V4−28
V2−26
V1−24
V3−23
V3−21
V3−20
V1−18
V3−15
V3−13
V3−11
V3−9
V5−10−1
V1−8
V3−64D
V3−7
V2−5
V7−4−1
V4−4
V1−3
V1−2
V6−1
D1−1
D2−2
D3−3
D6−6
D1−7
D2−8
D3−9
D3−10
D4−11
D5−12
D6−13
D2−15
D3−16
D4−17
D5−18
D6−19
D1−20
D2−21
D3−22
D4−23
D5−24
D6−25
D1−26
D7−27
J1
J2
J3
J4
J5
J6
Allele
01
^1−01_A318G
02
03
04
05
07
18
Del
log(lK)
[0, 1)
[1, 2)
[5, 10)
[10, 20)
[20, 50)
[50, Inf]

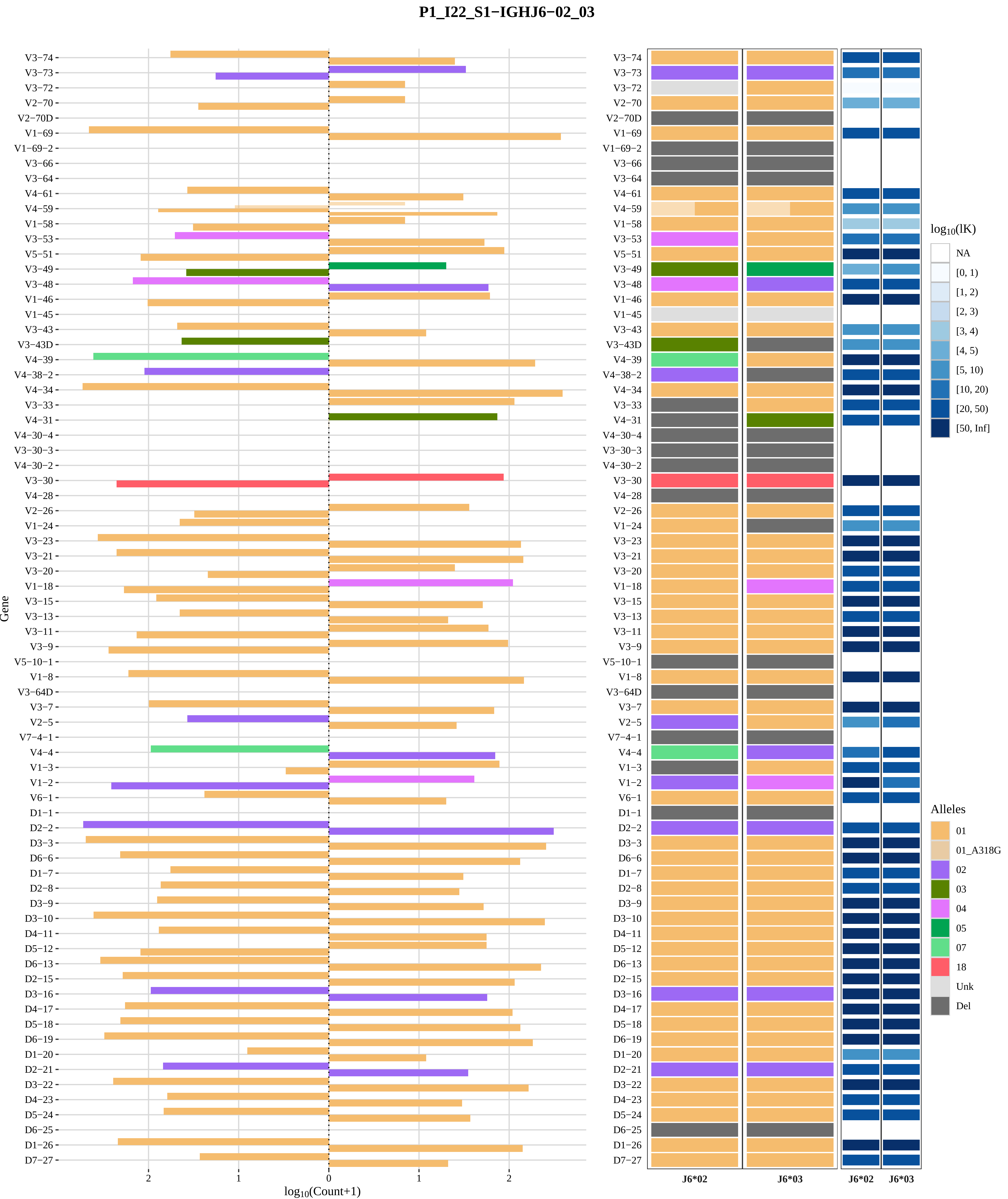

P1_I22_S1−IGHJ6−02_03
Gene
log10(Count+1)
2
1
0
1
2
V3−74
V3−73
V3−72
V2−70
V2−70D
V1−69
V1−69−2
V3−66
V3−64
V4−61
V4−59
V1−58
V3−53
V5−51
V3−49
V3−48
V1−46
V1−45
V3−43
V3−43D
V4−39
V4−38−2
V4−34
V3−33
V4−31
V4−30−4
V3−30−3
V4−30−2
V3−30
V4−28
V2−26
V1−24
V3−23
V3−21
V3−20
V1−18
V3−15
V3−13
V3−11
V3−9
V5−10−1
V1−8
V3−64D
V3−7
V2−5
V7−4−1
V4−4
V1−3
V1−2
V6−1
D1−1
D2−2
D3−3
D6−6
D1−7
D2−8
D3−9
D3−10
D4−11
D5−12
D6−13
D2−15
D3−16
D4−17
D5−18
D6−19
D1−20
D2−21
D3−22
D4−23
D5−24
D6−25
D1−26
D7−27
J6*02
J6*03
J6*02
J6*03
log10(lK)
NA
[0, 1)
[1, 2)
[2, 3)
[3, 4)
[4, 5)
[5, 10)
[10, 20)
[20, 50)
[50, Inf]
Alleles
01
01_A318G
02
03
04
05
07
18
Unk
Del

### Genotype and haplotype of VDJbase sample I23 (study P1)

(illustrations downloaded from VDJbase in July, 2020)

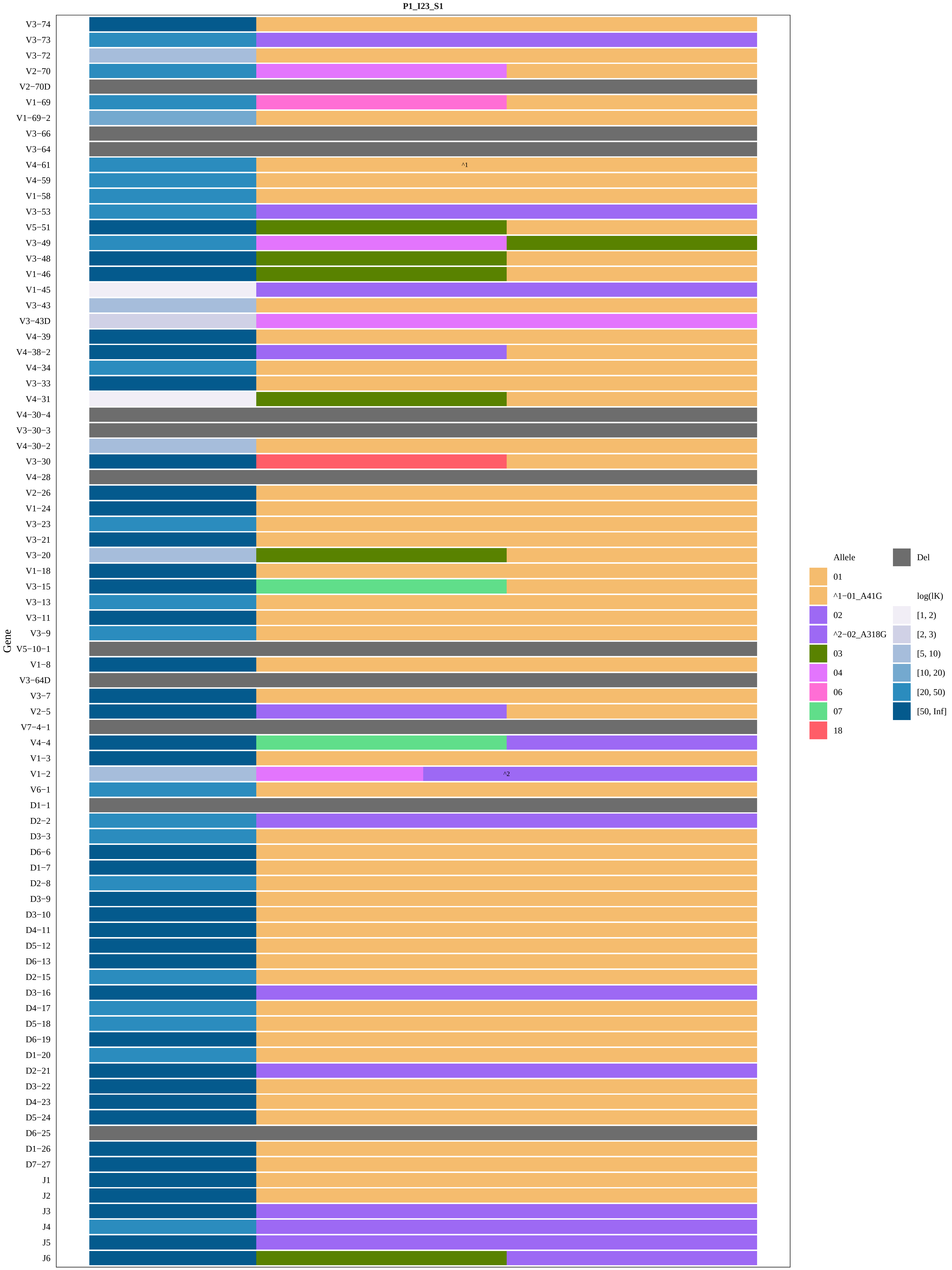
P1_I23_S1
Gene
V3−74
V3−73
V3−72
V2−70
V2−70D
V1−69
V1−69−2
V3−66
V3−64
V4−61
V4−59
V1−58
V3−53
V5−51
V3−49
V3−48
V1−46
V1−45
V3−43
V3−43D
V4−39
V4−38−2
V4−34
V3−33
V4−31
V4−30−4
V3−30−3
V4−30−2
V3−30
V4−28
V2−26
V1−24
V3−23
V3−21
V3−20
V1−18
V3−15
V3−13
V3−11
V3−9
V5−10−1
V1−8
V3−64D
V3−7
V2−5
V7−4−1
V4−4
V1−3
V1−2
V6−1
D1−1
D2−2
D3−3
D6−6
D1−7
D2−8
D3−9
D3−10
D4−11
D5−12
D6−13
D2−15
D3−16
D4−17
D5−18
D6−19
D1−20
D2−21
D3−22
D4−23
D5−24
D6−25
D1−26
D7−27
J1
J2
J3
J4
J5
J6
^1
^2
Allele
01
^1−01_A41G
02
^2−02_A318G
03
04
06
07
18
Del
log(lK)
[1, 2)
[2, 3)
[5, 10)
[10, 20)
[20, 50)
[50, Inf]

### P1_I23_S1−IGHJ6−02_03

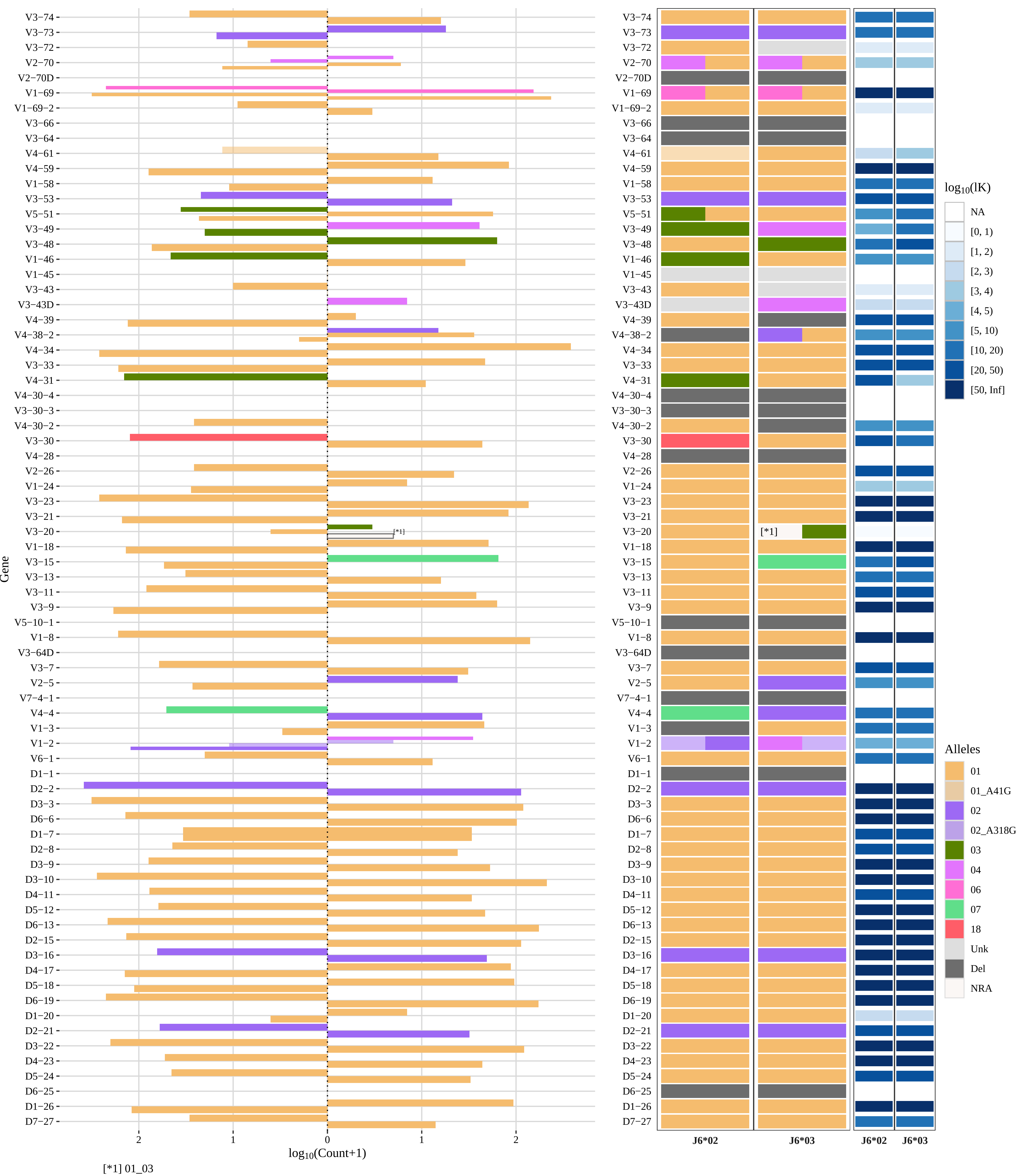

### Genotype and haplotype of VDJbase sample I24 (study P1)

(illustrations downloaded from VDJbase in July, 2020)

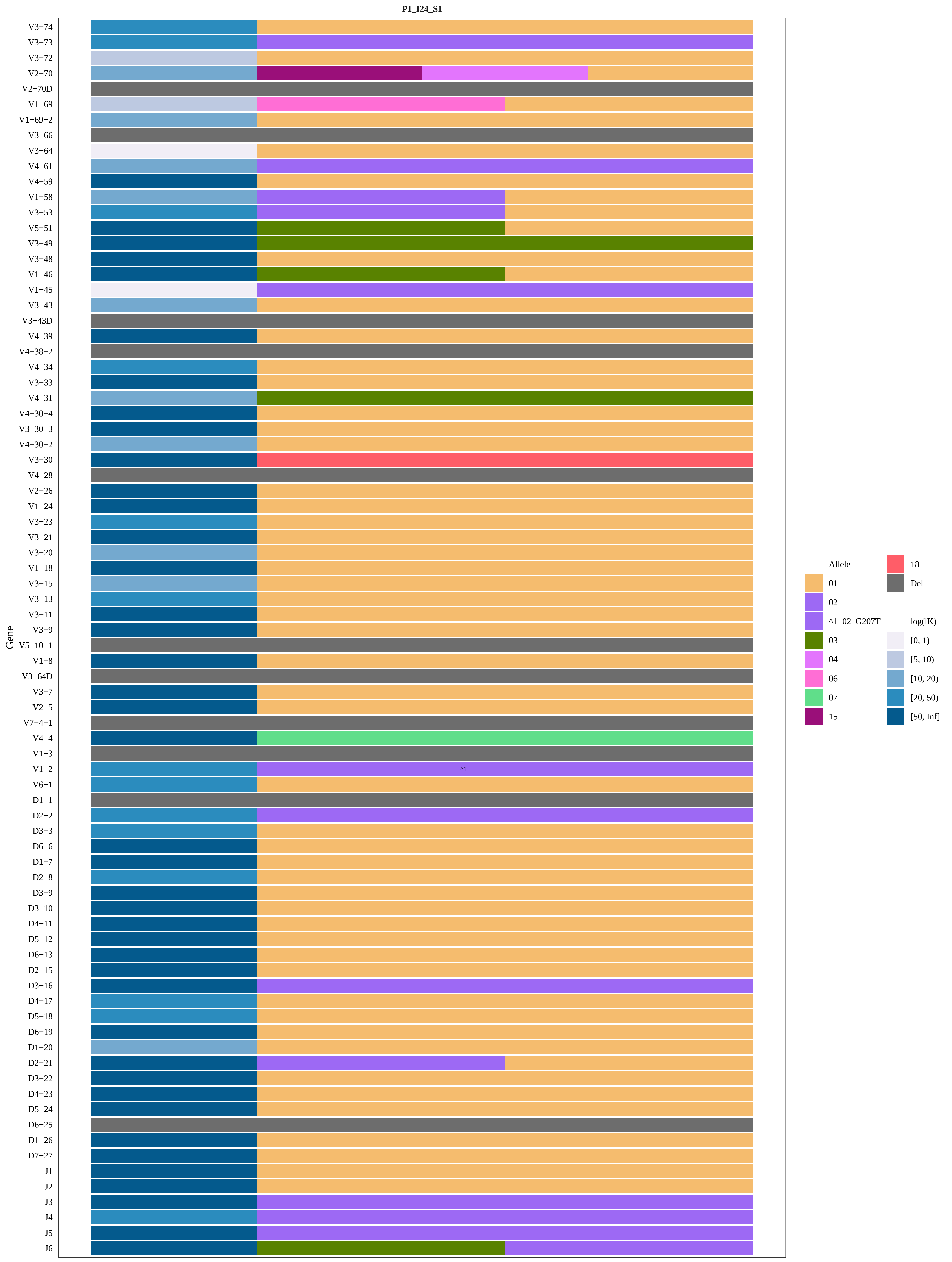
P1_I24_S1
Gene
V3−74
V3−73
V3−72
V2−70
V2−70D
V1−69
V1−69−2
V3−66
V3−64
V4−61
V4−59
V1−58
V3−53
V5−51
V3−49
V3−48
V1−46
V1−45
V3−43
V3−43D
V4−39
V4−38−2
V4−34
V3−33
V4−31
V4−30−4
V3−30−3
V4−30−2
V3−30
V4−28
V2−26
V1−24
V3−23
V3−21
V3−20
V1−18
V3−15
V3−13
V3−11
V3−9
V5−10−1
V1−8
V3−64D
V3−7
V2−5
V7−4−1
V4−4
V1−3
V1−2
V6−1
D1−1
D2−2
D3−3
D6−6
D1−7
D2−8
D3−9
D3−10
D4−11
D5−12
D6−13
D2−15
D3−16
D4−17
D5−18
D6−19
D1−20
D2−21
D3−22
D4−23
D5−24
D6−25
D1−26
D7−27
J1
J2
J3
J4
J5
J6
^1
Allele
01
02
^1−02_G207T
03
04
06
07
15
18
Del
log(lK)
[0, 1)
[5, 10)
[10, 20)
[20, 50)
[50, Inf]

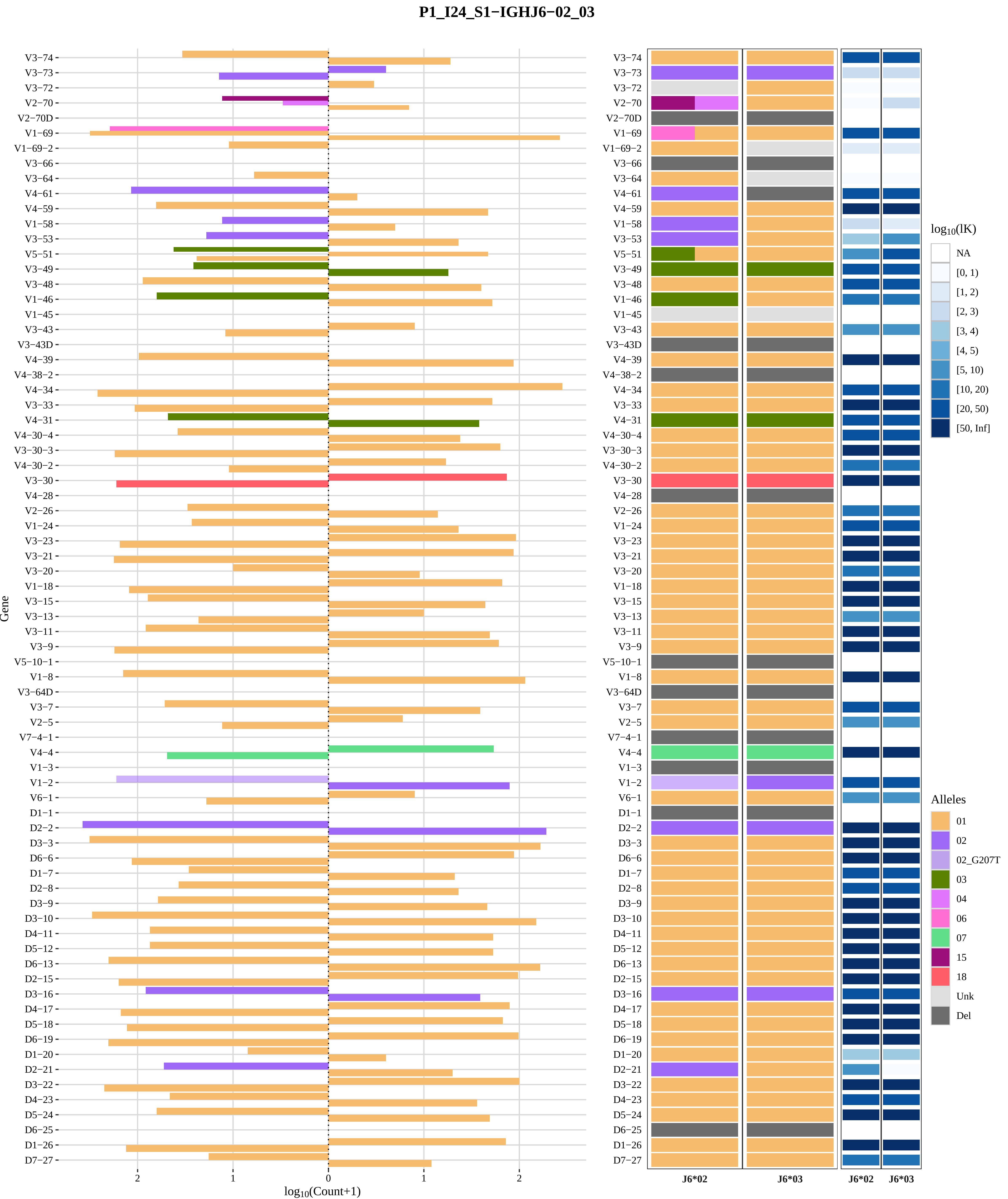
P1_I24_S1−IGHJ6−02_03
Gene
log10(Count+1)
2
1
0
1
2
V3−74
V3−73
V3−72
V2−70
V2−70D
V1−69
V1−69−2
V3−66
V3−64
V4−61
V4−59
V1−58
V3−53
V5−51
V3−49
V3−48
V1−46
V1−45
V3−43
V3−43D
V4−39
V4−38−2
V4−34
V3−33
V4−31
V4−30−4
V3−30−3
V4−30−2
V3−30
V4−28
V2−26
V1−24
V3−23
V3−21
V3−20
V1−18
V3−15
V3−13
V3−11
V3−9
V5−10−1
V1−8
V3−64D
V3−7
V2−5
V7−4−1
V4−4
V1−3
V1−2
V6−1
D1−1
D2−2
D3−3
D6−6
D1−7
D2−8
D3−9
D3−10
D4−11
D5−12
D6−13
D2−15
D3−16
D4−17
D5−18
D6−19
D1−20
D2−21
D3−22
D4−23
D5−24
D6−25
D1−26
D7−27
J6*02
J6*03
J6*02
J6*03
log10(lK)
NA
[0, 1)
[1, 2)
[2, 3)
[3, 4)
[4, 5)
[5, 10)
[10, 20)
[20, 50)
[50, Inf]
Alleles
01
02
02_G207T
03
04
06
07
15
18
Unk
Del

### Genotype and haplotype of VDJbase sample I27 (study P1)

(illustrations downloaded from VDJbase in July, 2020)

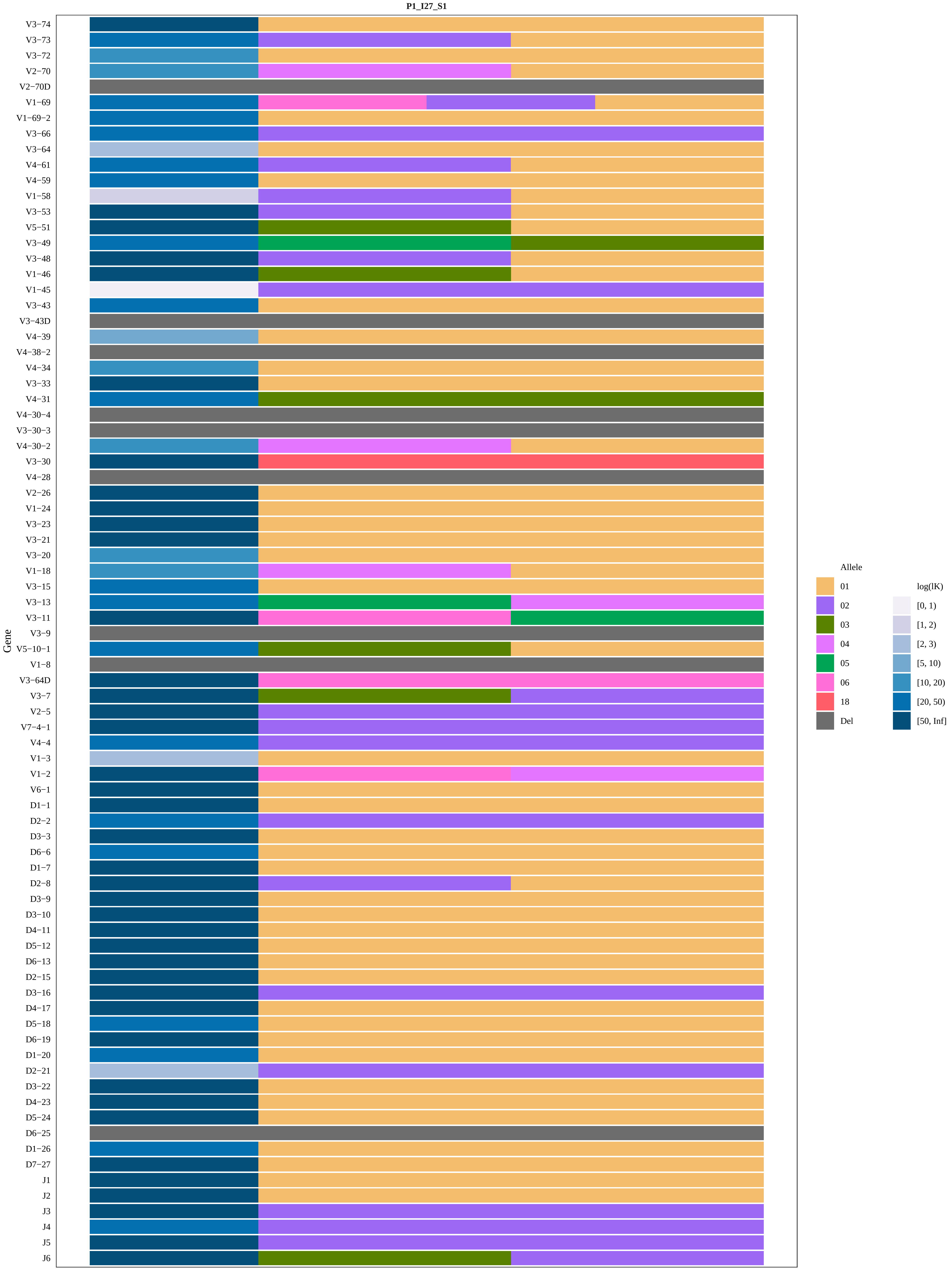
P1_I27_S1
Gene
V3−74
V3−73
V3−72
V2−70
V2−70D
V1−69
V1−69−2
V3−66
V3−64
V4−61
V4−59
V1−58
V3−53
V5−51
V3−49
V3−48
V1−46
V1−45
V3−43
V3−43D
V4−39
V4−38−2
V4−34
V3−33
V4−31
V4−30−4
V3−30−3
V4−30−2
V3−30
V4−28
V2−26
V1−24
V3−23
V3−21
V3−20
V1−18
V3−15
V3−13
V3−11
V3−9
V5−10−1
V1−8
V3−64D
V3−7
V2−5
V7−4−1
V4−4
V1−3
V1−2
V6−1
D1−1
D2−2
D3−3
D6−6
D1−7
D2−8
D3−9
D3−10
D4−11
D5−12
D6−13
D2−15
D3−16
D4−17
D5−18
D6−19
D1−20
D2−21
D3−22
D4−23
D5−24
D6−25
D1−26
D7−27
J1
J2
J3
J4
J5
J6
Allele
01
02
03
04
05
06
18
Del
log(lK)
[0, 1)
[1, 2)
[2, 3)
[5, 10)
[10, 20)
[20, 50)
[50, Inf]

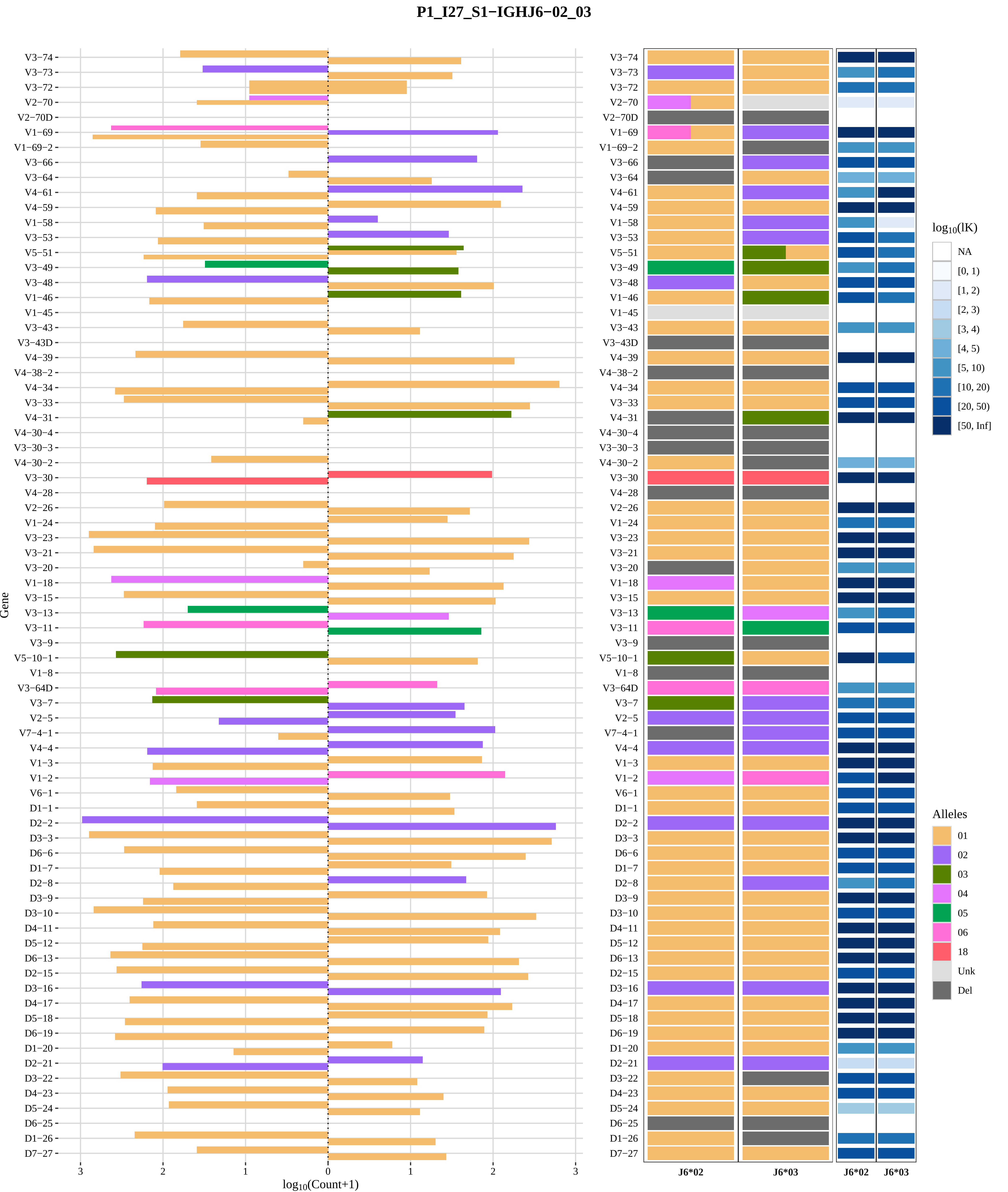

P1_I27_S1−IGHJ6−02_03
Gene
log10(Count+1)
J6*02
J6*03
log10(lK)
NA
[0, 1)
[1, 2)
[2, 3)
[3, 4)
[4, 5)
[5, 10)
[10, 20)
[20, 50)
[50, Inf]
Alleles
01
02
03
04
05
06
18
Unk
Del

### Genotype and haplotype of VDJbase sample I29 (study P1)

(illustrations downloaded from VDJbase in July, 2020)

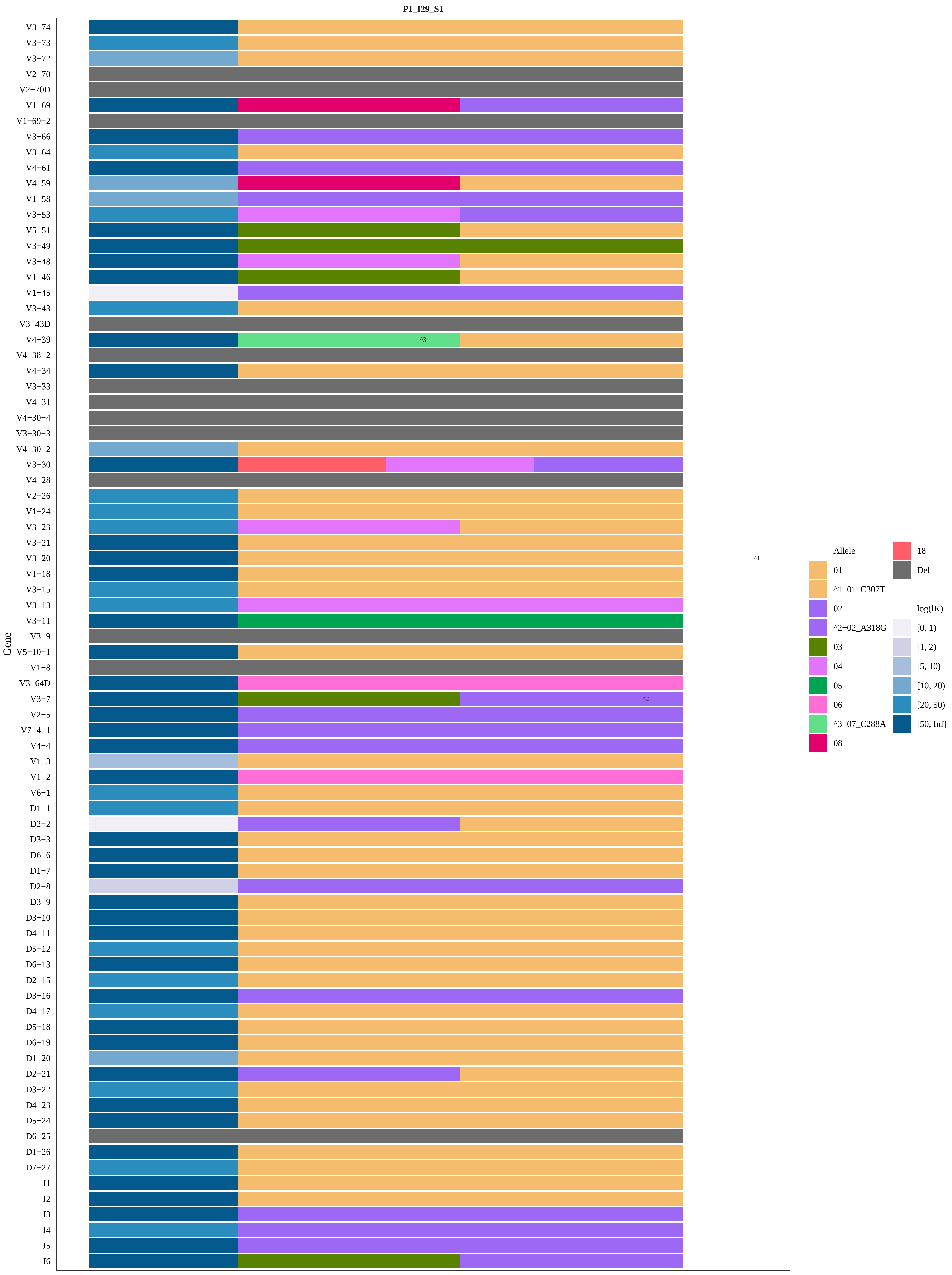
P1_I29_S1
Gene
V3−74
V3−73
V3−72
V2−70
V2−70D
V1−69
V1−69−2
V3−66
V3−64
V4−61
V4−59
V1−58
V3−53
V5−51
V3−49
V3−48
V1−46
V1−45
V3−43
V3−43D
V4−39
V4−38−2
V4−34
V3−33
V4−31
V4−30−4
V3−30−3
V4−30−2
V3−30
V4−28
V2−26
V1−24
V3−23
V3−21
V3−20
V1−18
V3−15
V3−13
V3−11
V3−9
V5−10−1
V1−8
V3−64D
V3−7
V2−5
V7−4−1
V4−4
V1−3
V1−2
V6−1
D1−1
D2−2
D3−3
D6−6
D1−7
D2−8
D3−9
D3−10
D4−11
D5−12
D6−13
D2−15
D3−16
D4−17
D5−18
D6−19
D1−20
D2−21
D3−22
D4−23
D5−24
D6−25
D1−26
D7−27
J1
J2
J3
J4
J5
J6
^3
^2
^1
Allele
01
^1−01_C307T
02
^2−02_A318G
03
04
05
06
^3−07_C288A
08
18
Del
log(lK)
[0, 1)
[1, 2)
[5, 10)
[10, 20)
[20, 50)
[50, Inf]

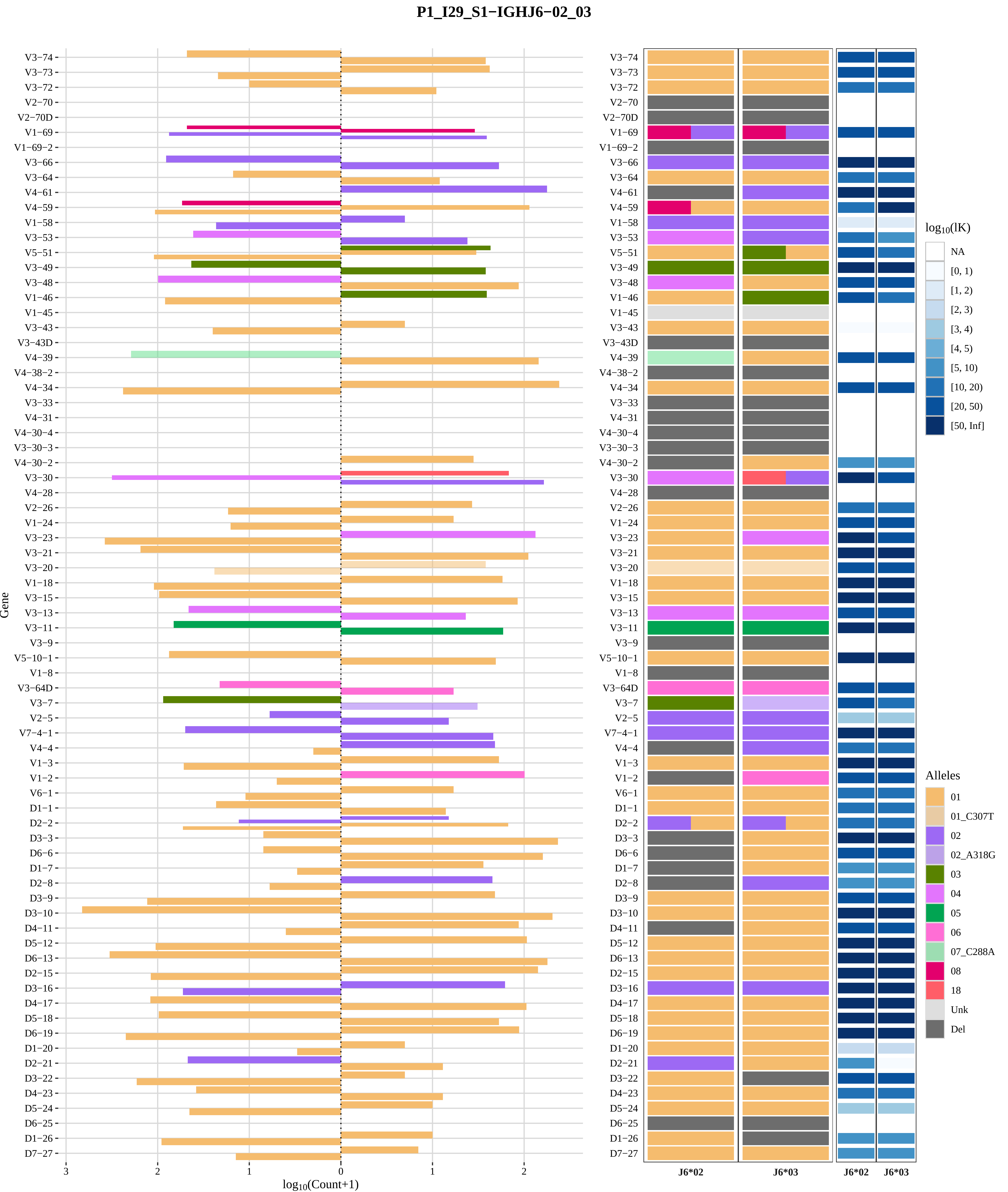
P1_I29_S1−IGHJ6−02_03
Gene
log10(Count+1)
V3−74
V3−73
V3−72
V2−70
V2−70D
V1−69
V1−69−2
V3−66
V3−64
V4−61
V4−59
V1−58
V3−53
V5−51
V3−49
V3−48
V1−46
V1−45
V3−43
V3−43D
V4−39
V4−38−2
V4−34
V3−33
V4−31
V4−30−4
V3−30−3
V4−30−2
V3−30
V4−28
V2−26
V1−24
V3−23
V3−21
V3−20
V1−18
V3−15
V3−13
V3−11
V3−9
V5−10−1
V1−8
V3−64D
V3−7
V2−5
V7−4−1
V4−4
V1−3
V1−2
V6−1
D1−1
D2−2
D3−3
D6−6
D1−7
D2−8
D3−9
D3−10
D4−11
D5−12
D6−13
D2−15
D3−16
D4−17
D5−18
D6−19
D1−20
D2−21
D3−22
D4−23
D5−24
D6−25
D1−26
D7−27
3
2
1
0
1
2
J6*02
J6*03
J6*02
J6*03
log10(lK)
NA
[0, 1)
[1, 2)
[2, 3)
[3, 4)
[4, 5)
[5, 10)
[10, 20)
[20, 50)
[50, Inf]
Alleles
01
01_C307T
02
02_A318G
03
04
05
06
07_C288A
08
18
Unk
Del

### Genotype and haplotype of VDJbase sample I33 (study P1)

(illustrations downloaded from VDJbase in July, 2020)

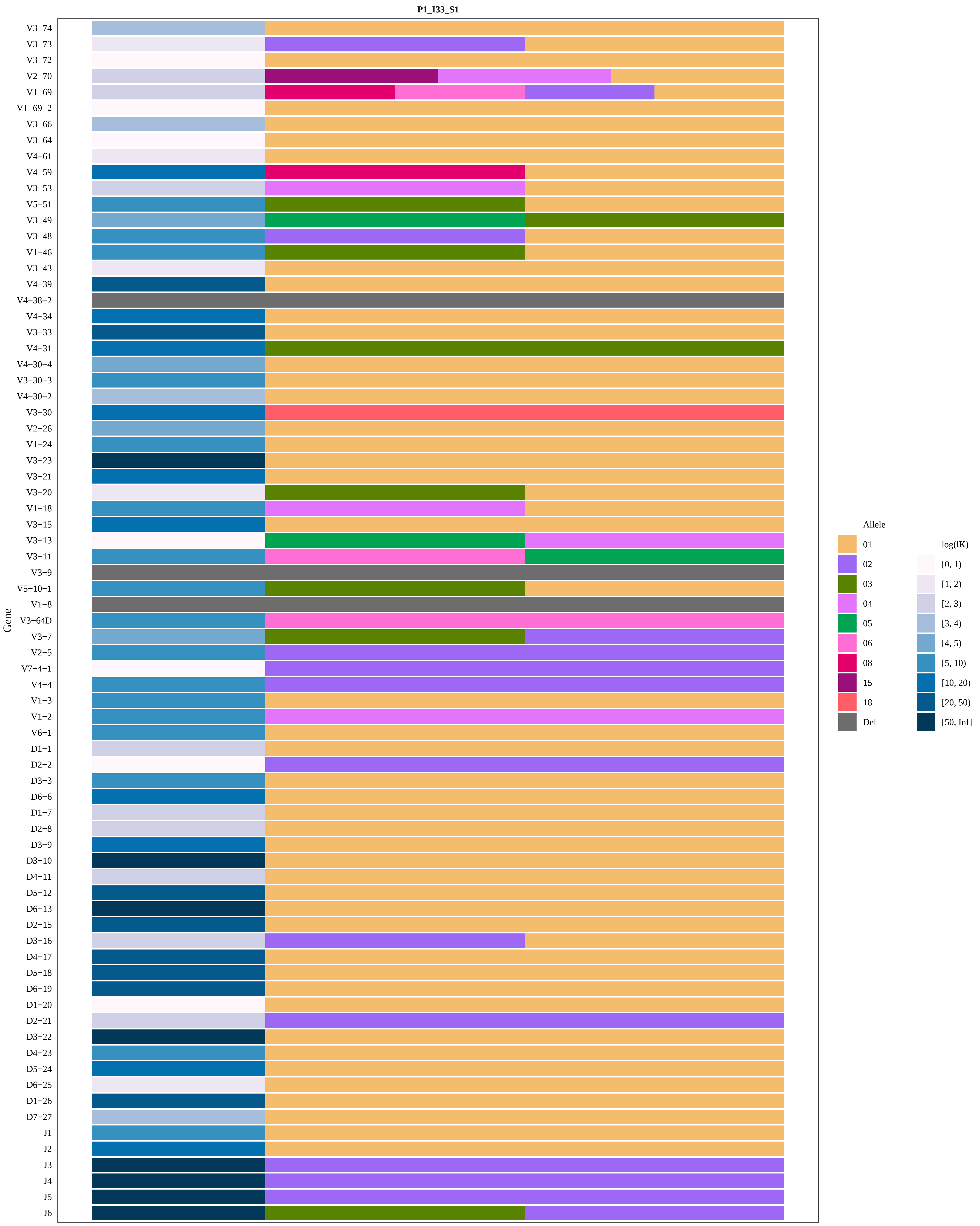

P1_I33_S1
V3−74
V3−73
V3−72
V2−70
V1−69
V1−69−2
V3−66
V3−64
V4−61
V4−59
V3−53
V5−51
V3−49
V3−48
V1−46
V3−43
V4−39
V4−38−2
V4−34
V3−33
V4−31
V4−30−4
V3−30−3
V4−30−2
V3−30
V2−26
V1−24
V3−23
V3−21
V3−20
V1−18
V3−15
V3−13
V3−11
V3−9
V5−10−1
V1−8
V3−64D
V3−7
V2−5
V7−4−1
V4−4
V1−3
V1−2
V6−1
D1−1
D2−2
D3−3
D6−6
D1−7
D2−8
D3−9
D3−10
D4−11
D5−12
D6−13
D2−15
D3−16
D4−17
D5−18
D6−19
D1−20
D2−21
D3−22
D4−23
D5−24
D6−25
D1−26
D7−27
J1
J2
J3
J4
J5
J6
Gene
Allele
01
02
03
04
05
06
08
15
18
Del
log(lK)
[0, 1)
[1, 2)
[2, 3)
[3, 4)
[4, 5)
[5, 10)
[10, 20)
[20, 50)
[50, Inf]

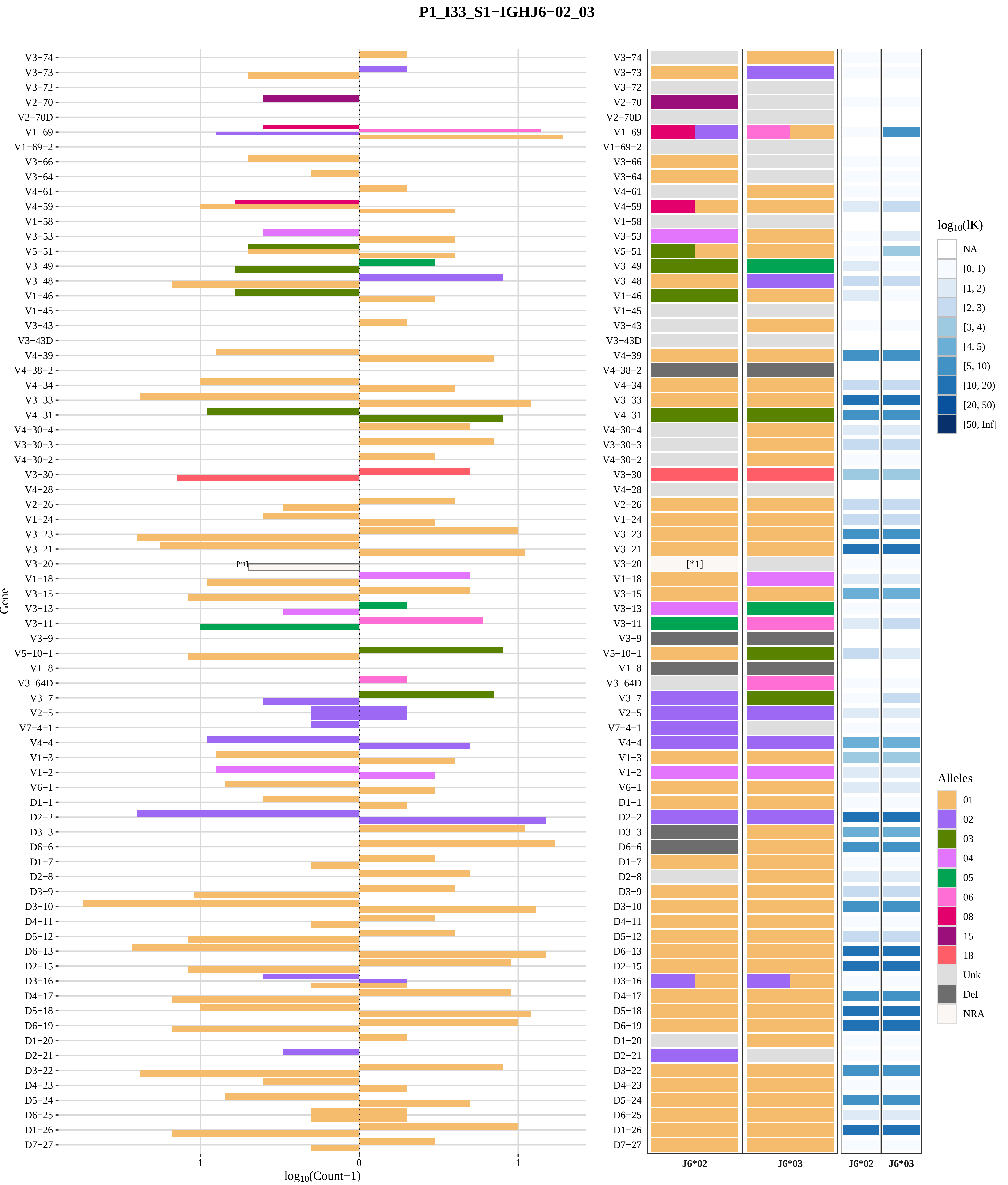
P1_I33_S1−IGHJ6−02_03
V3−74
V3−73
V3−72
V2−70
V2−70D
V1−69
V1−69−2
V3−66
V3−64
V4−61
V4−59
V1−58
V3−53
V5−51
V3−49
V3−48
V1−46
V1−45
V3−43
V3−43D
V4−39
V4−38−2
V4−34
V3−33
V4−31
V4−30−4
V3−30−3
V4−30−2
V3−30
V4−28
V2−26
V1−24
V3−23
V3−21
V3−20
V1−18
V3−15
V3−13
V3−11
V3−9
V5−10−1
V1−8
V3−64D
V3−7
V2−5
V7−4−1
V4−4
V1−3
V1−2
V6−1
D1−1
D2−2
D3−3
D6−6
D1−7
D2−8
D3−9
D3−10
D4−11
D5−12
D6−13
D2−15
D3−16
D4−17
D5−18
D6−19
D1−20
D2−21
D3−22
D4−23
D5−24
D6−25
D1−26
D7−27
Gene
[*1]
1
0
1
log10(Count+1)
J6*02
J6*03
J6*02
J6*03
log10(lK)
NA
[0, 1)
[1, 2)
[2, 3)
[3, 4)
[4, 5)
[5, 10)
[10, 20)
[20, 50)
[50, Inf]
Alleles
01
02
03
04
05
06
08
15
18
Unk
Del
NRA
[*1] 01_03

### Genotype and haplotype of VDJbase sample I39 (study P1)

(illustrations downloaded from VDJbase in July, 2020)

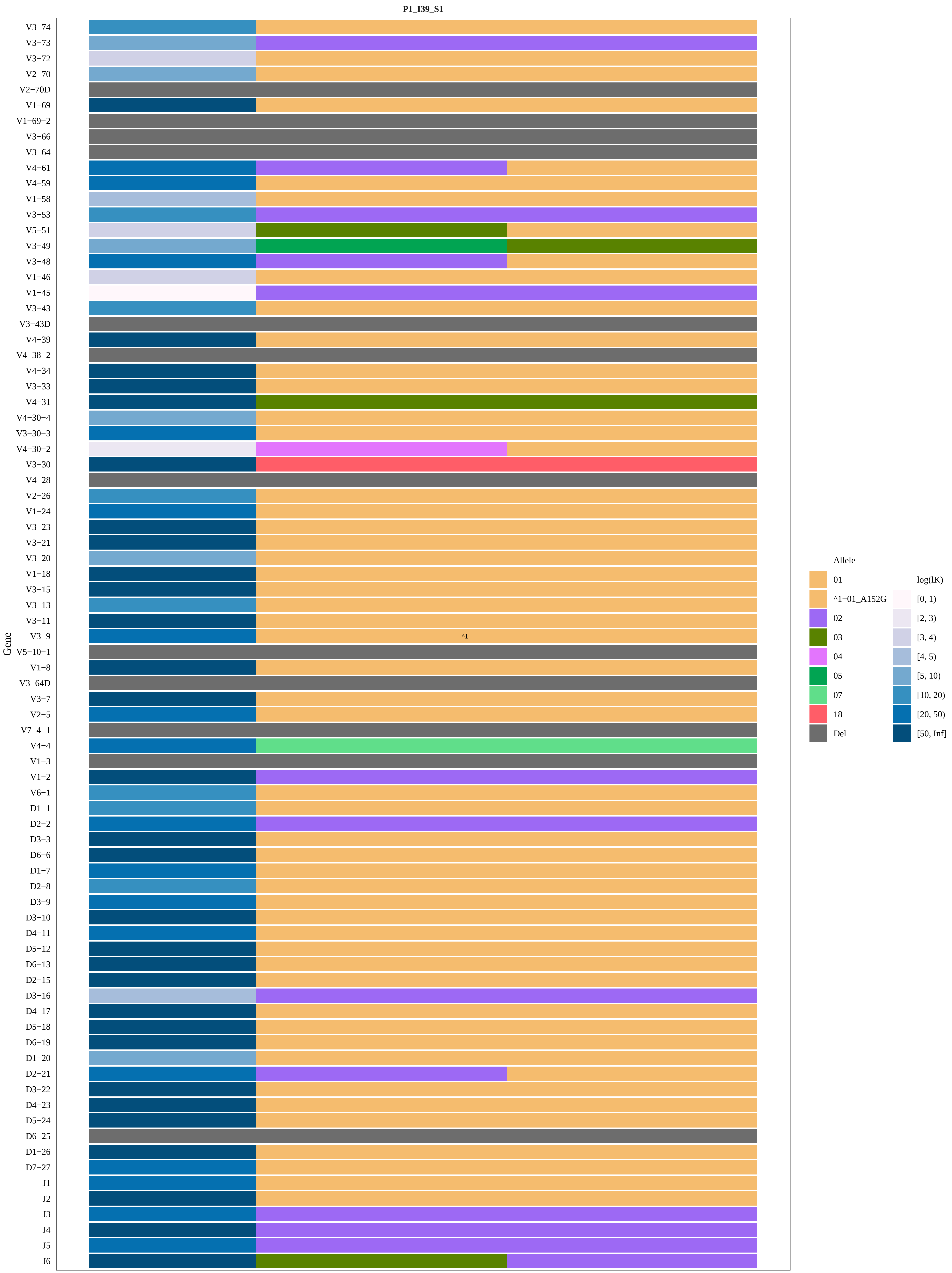
P1_I39_S1
Gene
V3−74
V3−73
V3−72
V2−70
V2−70D
V1−69
V1−69−2
V3−66
V3−64
V4−61
V4−59
V1−58
V3−53
V5−51
V3−49
V3−48
V1−46
V1−45
V3−43
V3−43D
V4−39
V4−38−2
V4−34
V3−33
V4−31
V4−30−4
V3−30−3
V4−30−2
V3−30
V4−28
V2−26
V1−24
V3−23
V3−21
V3−20
V1−18
V3−15
V3−13
V3−11
V3−9
V5−10−1
V1−8
V3−64D
V3−7
V2−5
V7−4−1
V4−4
V1−3
V1−2
V6−1
D1−1
D2−2
D3−3
D6−6
D1−7
D2−8
D3−9
D3−10
D4−11
D5−12
D6−13
D2−15
D3−16
D4−17
D5−18
D6−19
D1−20
D2−21
D3−22
D4−23
D5−24
D6−25
D1−26
D7−27
J1
J2
J3
J4
J5
J6
^1
Allele
01
^1−01_A152G
02
03
04
05
07
18
Del
log(lK)
[0, 1)
[2, 3)
[3, 4)
[4, 5)
[5, 10)
[10, 20)
[20, 50)
[50, Inf]

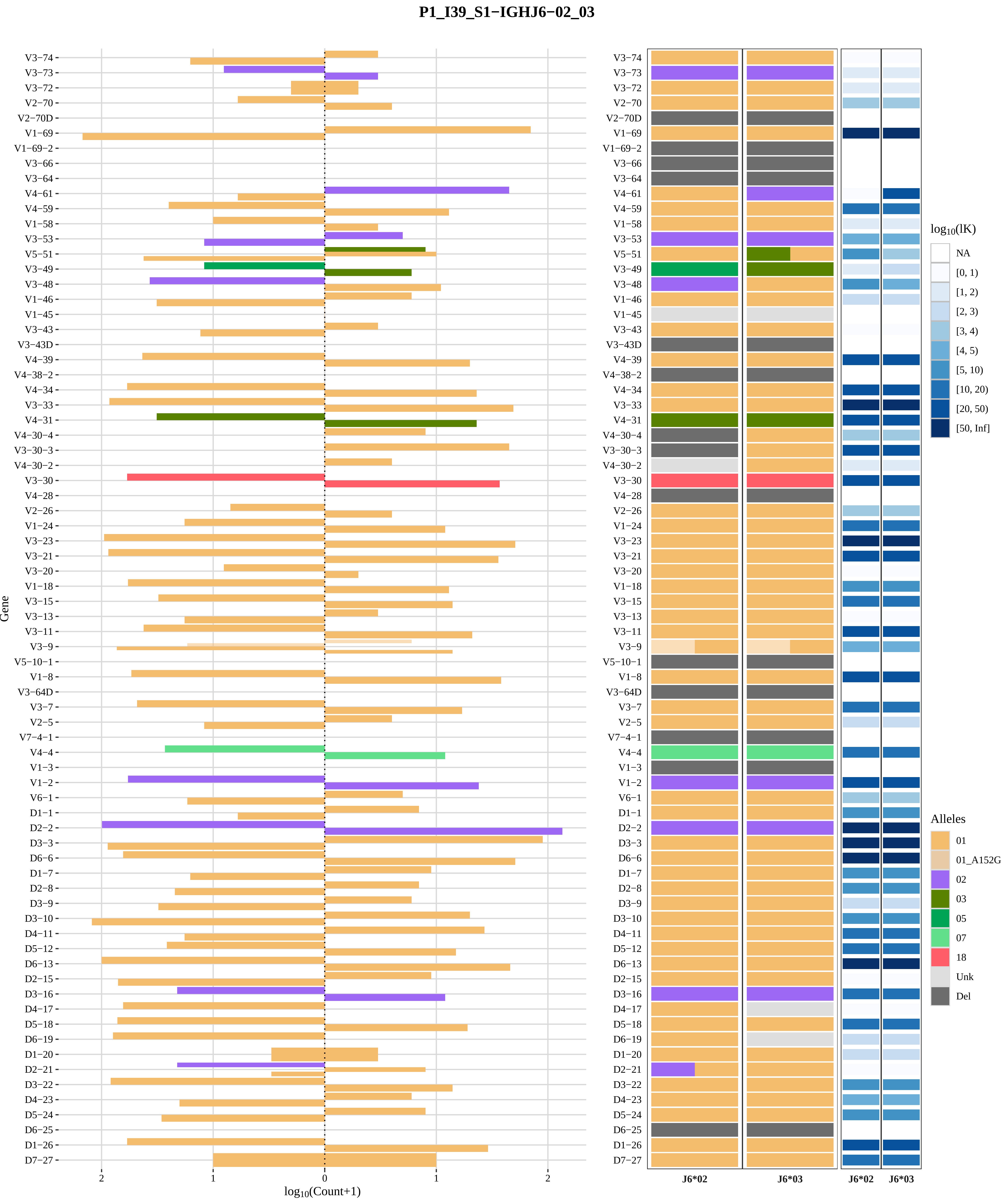
P1_I39_S1−IGHJ6−02_03
Gene
log10(Count+1)
2
1
0
1
2
V3−74
V3−73
V3−72
V2−70
V2−70D
V1−69
V1−69−2
V3−66
V3−64
V4−61
V4−59
V1−58
V3−53
V5−51
V3−49
V3−48
V1−46
V1−45
V3−43
V3−43D
V4−39
V4−38−2
V4−34
V3−33
V4−31
V4−30−4
V3−30−3
V4−30−2
V3−30
V4−28
V2−26
V1−24
V3−23
V3−21
V3−20
V1−18
V3−15
V3−13
V3−11
V3−9
V5−10−1
V1−8
V3−64D
V3−7
V2−5
V7−4−1
V4−4
V1−3
V1−2
V6−1
D1−1
D2−2
D3−3
D6−6
D1−7
D2−8
D3−9
D3−10
D4−11
D5−12
D6−13
D2−15
D3−16
D4−17
D5−18
D6−19
D1−20
D2−21
D3−22
D4−23
D5−24
D6−25
D1−26
D7−27
J6*02
J6*03
J6*02
J6*03
log10(lK)
NA
[0, 1)
[1, 2)
[2, 3)
[3, 4)
[4, 5)
[5, 10)
[10, 20)
[20, 50)
[50, Inf]
Alleles
01
01_A152G
02
03
05
07
18
Unk
Del

### Genotype and haplotype of VDJbase sample I40 (study P1)

(illustrations downloaded from VDJbase in July, 2020)

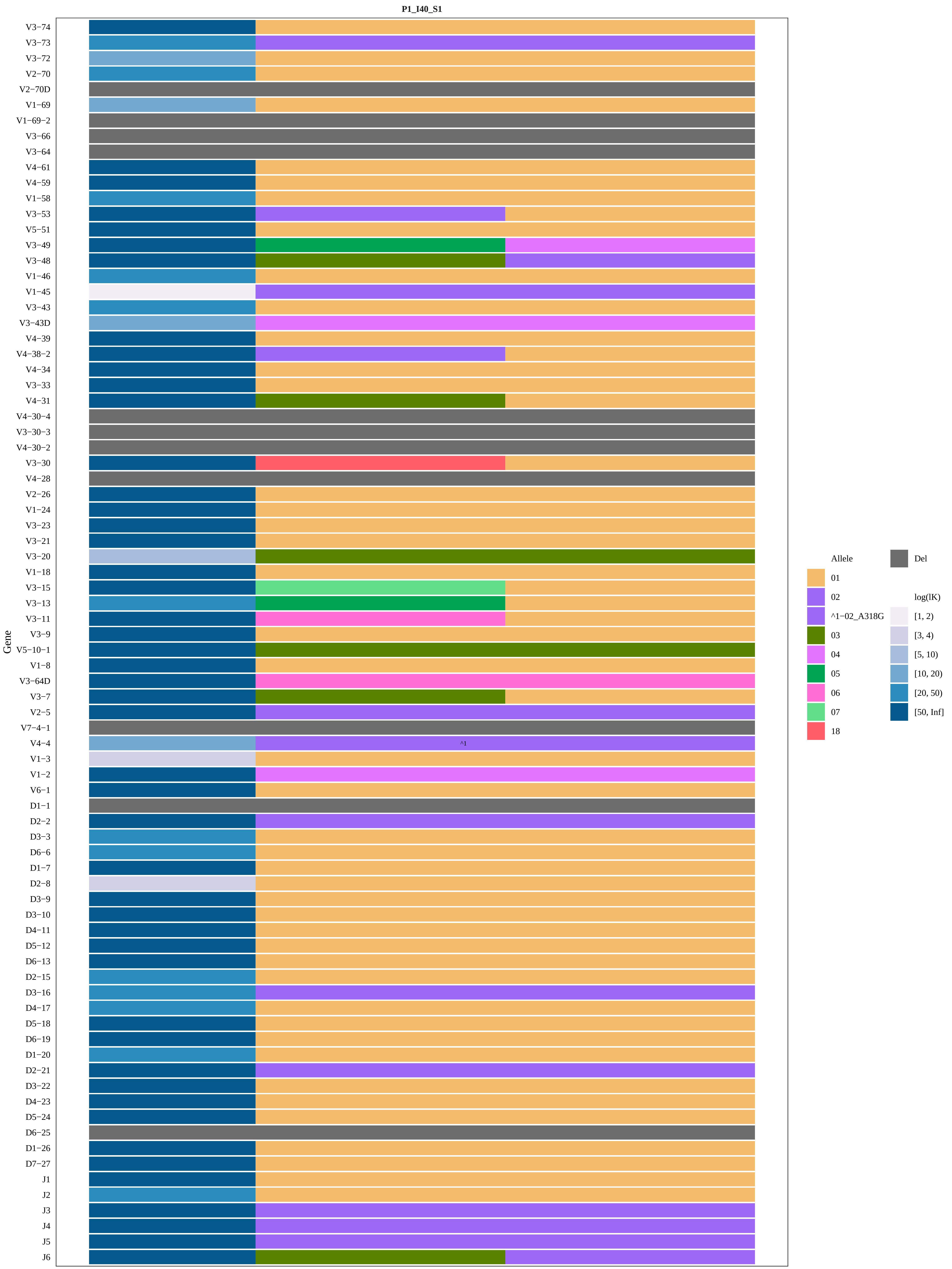
P1_I40_S1
Gene
V3−74
V3−73
V3−72
V2−70
V2−70D
V1−69
V1−69−2
V3−66
V3−64
V4−61
V4−59
V1−58
V3−53
V5−51
V3−49
V3−48
V1−46
V1−45
V3−43
V3−43D
V4−39
V4−38−2
V4−34
V3−33
V4−31
V4−30−4
V3−30−3
V4−30−2
V3−30
V4−28
V2−26
V1−24
V3−23
V3−21
V3−20
V1−18
V3−15
V3−13
V3−11
V3−9
V5−10−1
V1−8
V3−64D
V3−7
V2−5
V7−4−1
V4−4
V1−3
V1−2
V6−1
D1−1
D2−2
D3−3
D6−6
D1−7
D2−8
D3−9
D3−10
D4−11
D5−12
D6−13
D2−15
D3−16
D4−17
D5−18
D6−19
D1−20
D2−21
D3−22
D4−23
D5−24
D6−25
D1−26
D7−27
J1
J2
J3
J4
J5
J6
^1
Allele
01
02
^1−02_A318G
03
04
05
06
07
18
Del
log(lK)
[1, 2)
[3, 4)
[5, 10)
[10, 20)
[20, 50)
[50, Inf]

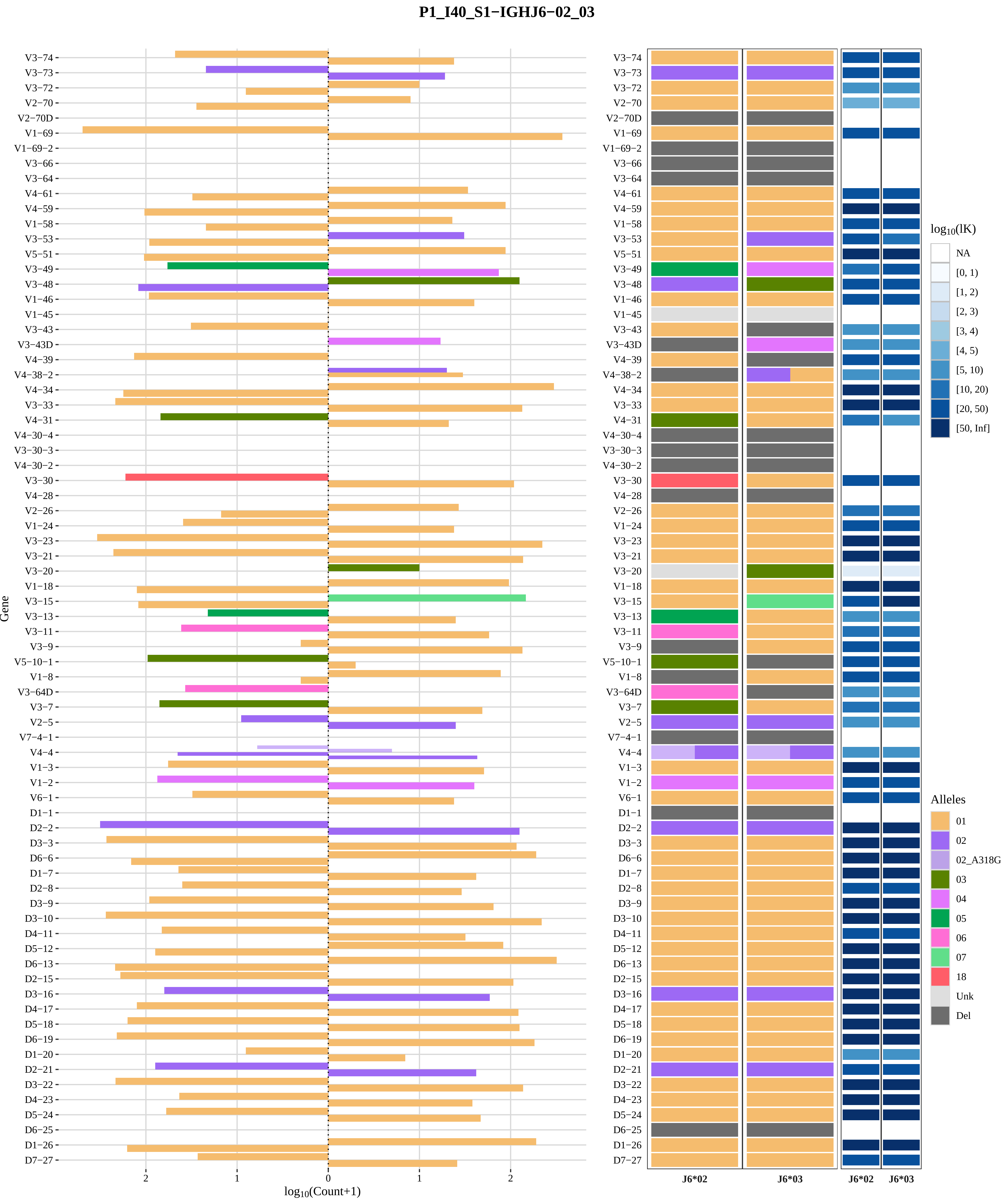

P1_I40_S1−IGHJ6−02_03
Gene
log10(Count+1)
V3−74
V3−73
V3−72
V2−70
V2−70D
V1−69
V1−69−2
V3−66
V3−64
V4−61
V4−59
V1−58
V3−53
V5−51
V3−49
V3−48
V1−46
V1−45
V3−43
V3−43D
V4−39
V4−38−2
V4−34
V3−33
V4−31
V4−30−4
V3−30−3
V4−30−2
V3−30
V4−28
V2−26
V1−24
V3−23
V3−21
V3−20
V1−18
V3−15
V3−13
V3−11
V3−9
V5−10−1
V1−8
V3−64D
V3−7
V2−5
V7−4−1
V4−4
V1−3
V1−2
V6−1
D1−1
D2−2
D3−3
D6−6
D1−7
D2−8
D3−9
D3−10
D4−11
D5−12
D6−13
D2−15
D3−16
D4−17
D5−18
D6−19
D1−20
D2−21
D3−22
D4−23
D5−24
D6−25
D1−26
D7−27
J6*02
J6*03
log10(lK)
NA
[0, 1)
[1, 2)
[2, 3)
[3, 4)
[4, 5)
[5, 10)
[10, 20)
[20, 50)
[50, Inf]
Alleles
01
02
02_A318G
03
04
05
06
07
18
Unk
Del

### Genotype and haplotype of VDJbase sample I41 (study P1)

(illustrations downloaded from VDJbase in July, 2020)

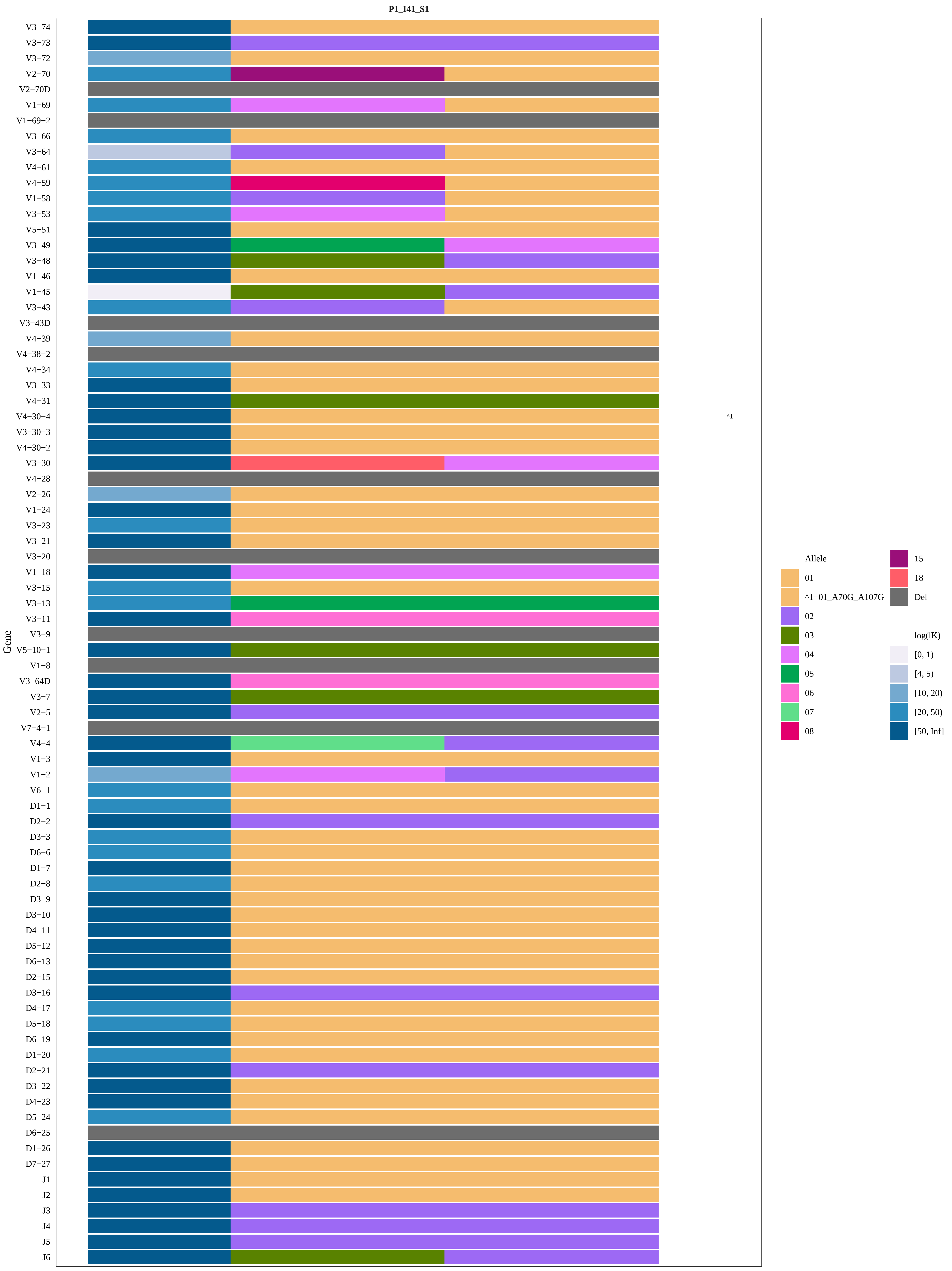

P1_I41_S1
Gene
V3−74
V3−73
V3−72
V2−70
V2−70D
V1−69
V1−69−2
V3−66
V3−64
V4−61
V4−59
V1−58
V3−53
V5−51
V3−49
V3−48
V1−46
V1−45
V3−43
V3−43D
V4−39
V4−38−2
V4−34
V3−33
V4−31
V4−30−4
V3−30−3
V4−30−2
V3−30
V4−28
V2−26
V1−24
V3−23
V3−21
V3−20
V1−18
V3−15
V3−13
V3−11
V3−9
V5−10−1
V1−8
V3−64D
V3−7
V2−5
V7−4−1
V4−4
V1−3
V1−2
V6−1
D1−1
D2−2
D3−3
D6−6
D1−7
D2−8
D3−9
D3−10
D4−11
D5−12
D6−13
D2−15
D3−16
D4−17
D5−18
D6−19
D1−20
D2−21
D3−22
D4−23
D5−24
D6−25
D1−26
D7−27
J1
J2
J3
J4
J5
J6
^1
Allele
01
^1−01_A70G_A107G
02
03
04
05
06
07
08
15
18
Del
log(lK)
[0, 1)
[4, 5)
[10, 20)
[20, 50)
[50, Inf]

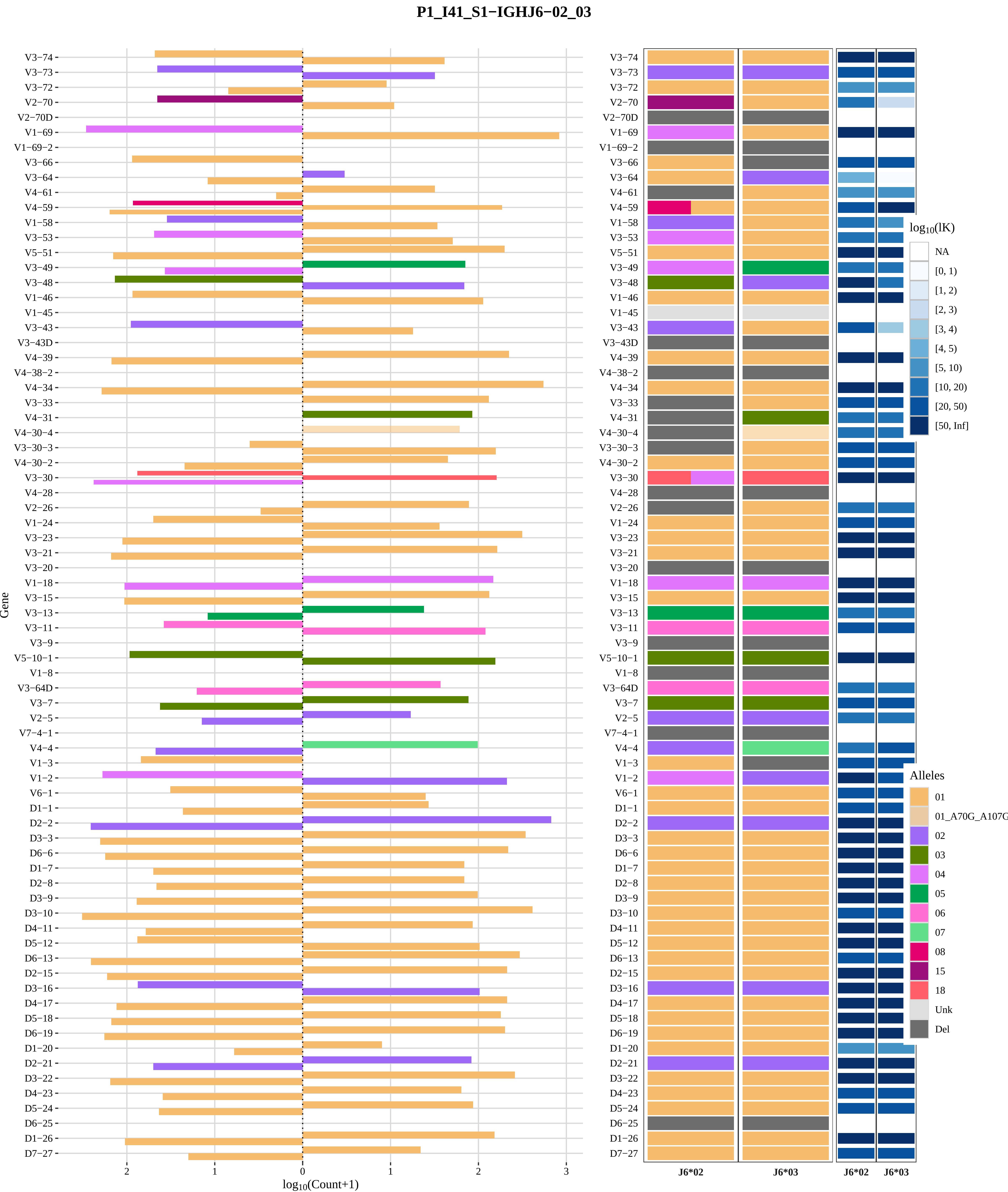

P1_I41_S1−IGHJ6−02_03
Gene
V3−74
V3−73
V3−72
V2−70
V2−70D
V1−69
V1−69−2
V3−66
V3−64
V4−61
V4−59
V1−58
V3−53
V5−51
V3−49
V3−48
V1−46
V1−45
V3−43
V3−43D
V4−39
V4−38−2
V4−34
V3−33
V4−31
V4−30−4
V3−30−3
V4−30−2
V3−30
V4−28
V2−26
V1−24
V3−23
V3−21
V3−20
V1−18
V3−15
V3−13
V3−11
V3−9
V5−10−1
V1−8
V3−64D
V3−7
V2−5
V7−4−1
V4−4
V1−3
V1−2
V6−1
D1−1
D2−2
D3−3
D6−6
D1−7
D2−8
D3−9
D3−10
D4−11
D5−12
D6−13
D2−15
D3−16
D4−17
D5−18
D6−19
D1−20
D2−21
D3−22
D4−23
D5−24
D6−25
D1−26
D7−27
2
1
0
1
2
3
log10(Count+1)
J6*02
J6*03
J6*02
J6*03
log10(lK)
NA
[0, 1)
[1, 2)
[2, 3)
[3, 4)
[4, 5)
[5, 10)
[10, 20)
[20, 50)
[50, Inf]
Alleles
01
01_A70G_A107G
02
03
04
05
06
07
08
15
18
Unk
Del

### Genotype and haplotype of VDJbase sample I43 (study P1)

(illustrations downloaded from VDJbase in July, 2020)

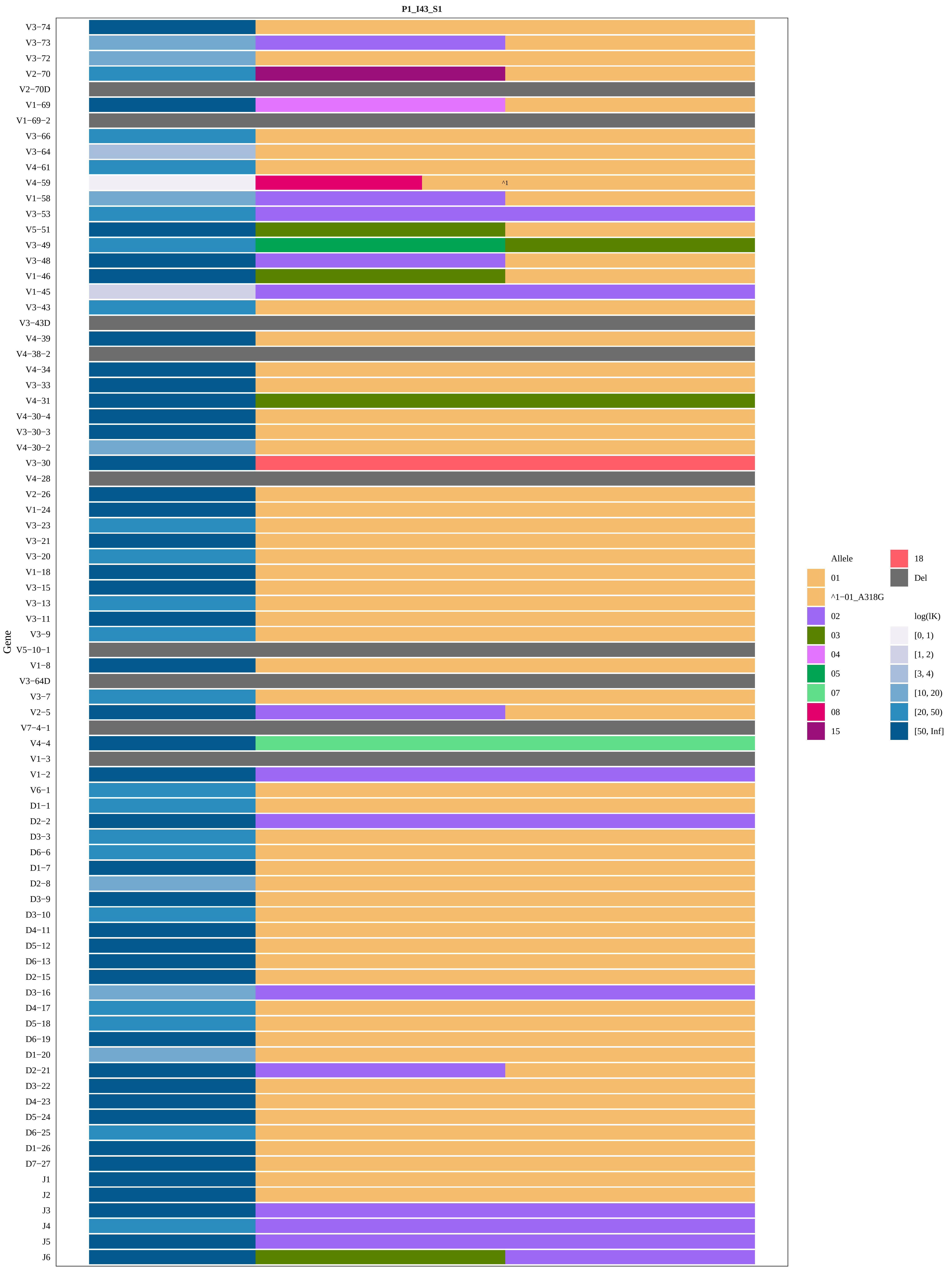
P1_I43_S1
Gene
V3−74
V3−73
V3−72
V2−70
V2−70D
V1−69
V1−69−2
V3−66
V3−64
V4−61
V4−59
V1−58
V3−53
V5−51
V3−49
V3−48
V1−46
V1−45
V3−43
V3−43D
V4−39
V4−38−2
V4−34
V3−33
V4−31
V4−30−4
V3−30−3
V4−30−2
V3−30
V4−28
V2−26
V1−24
V3−23
V3−21
V3−20
V1−18
V3−15
V3−13
V3−11
V3−9
V5−10−1
V1−8
V3−64D
V3−7
V2−5
V7−4−1
V4−4
V1−3
V1−2
V6−1
D1−1
D2−2
D3−3
D6−6
D1−7
D2−8
D3−9
D3−10
D4−11
D5−12
D6−13
D2−15
D3−16
D4−17
D5−18
D6−19
D1−20
D2−21
D3−22
D4−23
D5−24
D6−25
D1−26
D7−27
J1
J2
J3
J4
J5
J6
^1
Allele
01
^1−01_A318G
02
03
04
05
07
08
15
18
Del
log(lK)
[0, 1)
[1, 2)
[3, 4)
[10, 20)
[20, 50)
[50, Inf]

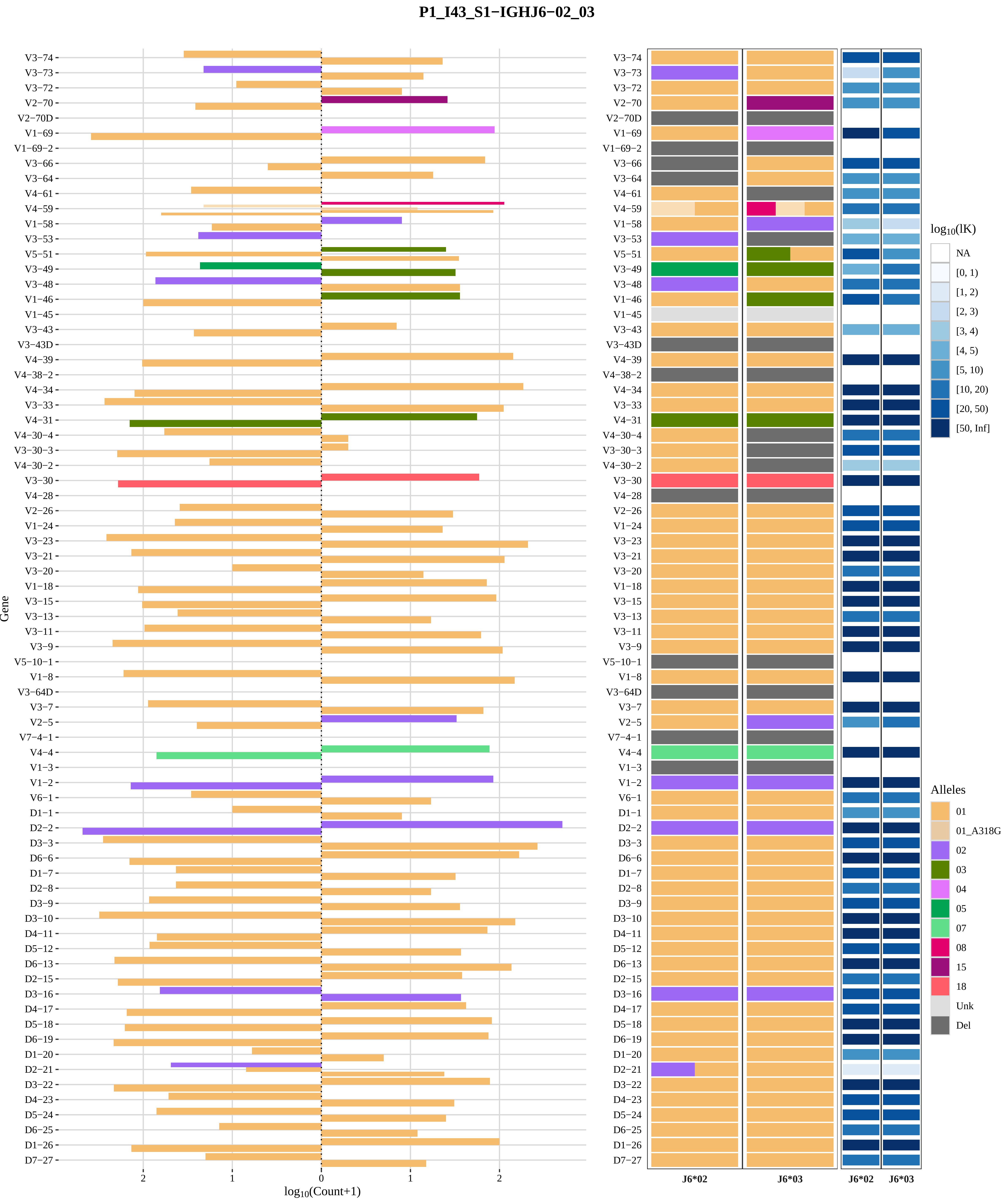

P1_I43_S1−IGHJ6−02_03
Gene
log10(Count+1)
J6*02
J6*03
log10(lK)
NA
[0, 1)
[1, 2)
[2, 3)
[3, 4)
[4, 5)
[5, 10)
[10, 20)
[20, 50)
[50, Inf]
Alleles
01
01_A318G
02
03
04
05
07
08
15
18
Unk
Del

### Genotype and haplotype of VDJbase sample I46 (study P1)

(illustrations downloaded from VDJbase in July, 2020)

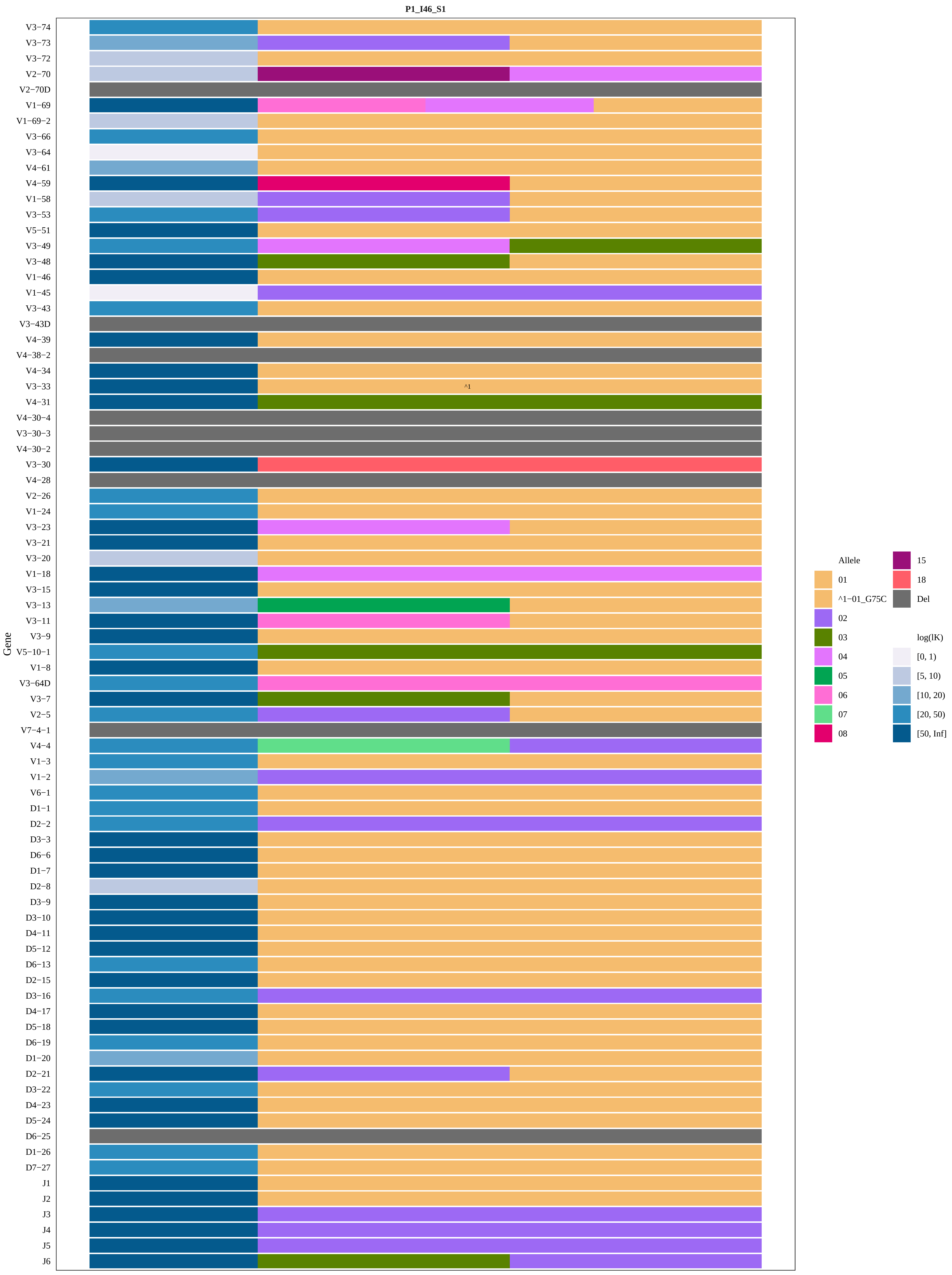

P1_I46_S1
Gene
V3−74
V3−73
V3−72
V2−70
V2−70D
V1−69
V1−69−2
V3−66
V3−64
V4−61
V4−59
V1−58
V3−53
V5−51
V3−49
V3−48
V1−46
V1−45
V3−43
V3−43D
V4−39
V4−38−2
V4−34
V3−33
^1
V4−31
V4−30−4
V3−30−3
V4−30−2
V3−30
V4−28
V2−26
V1−24
V3−23
V3−21
V3−20
V1−18
V3−15
V3−13
V3−11
V3−9
V5−10−1
V1−8
V3−64D
V3−7
V2−5
V7−4−1
V4−4
V1−3
V1−2
V6−1
D1−1
D2−2
D3−3
D6−6
D1−7
D2−8
D3−9
D3−10
D4−11
D5−12
D6−13
D2−15
D3−16
D4−17
D5−18
D6−19
D1−20
D2−21
D3−22
D4−23
D5−24
D6−25
D1−26
D7−27
J1
J2
J3
J4
J5
J6
Allele
01
^1−01_G75C
02
03
04
05
06
07
08
15
18
Del
log(lK)
[0, 1)
[5, 10)
[10, 20)
[20, 50)
[50, Inf]

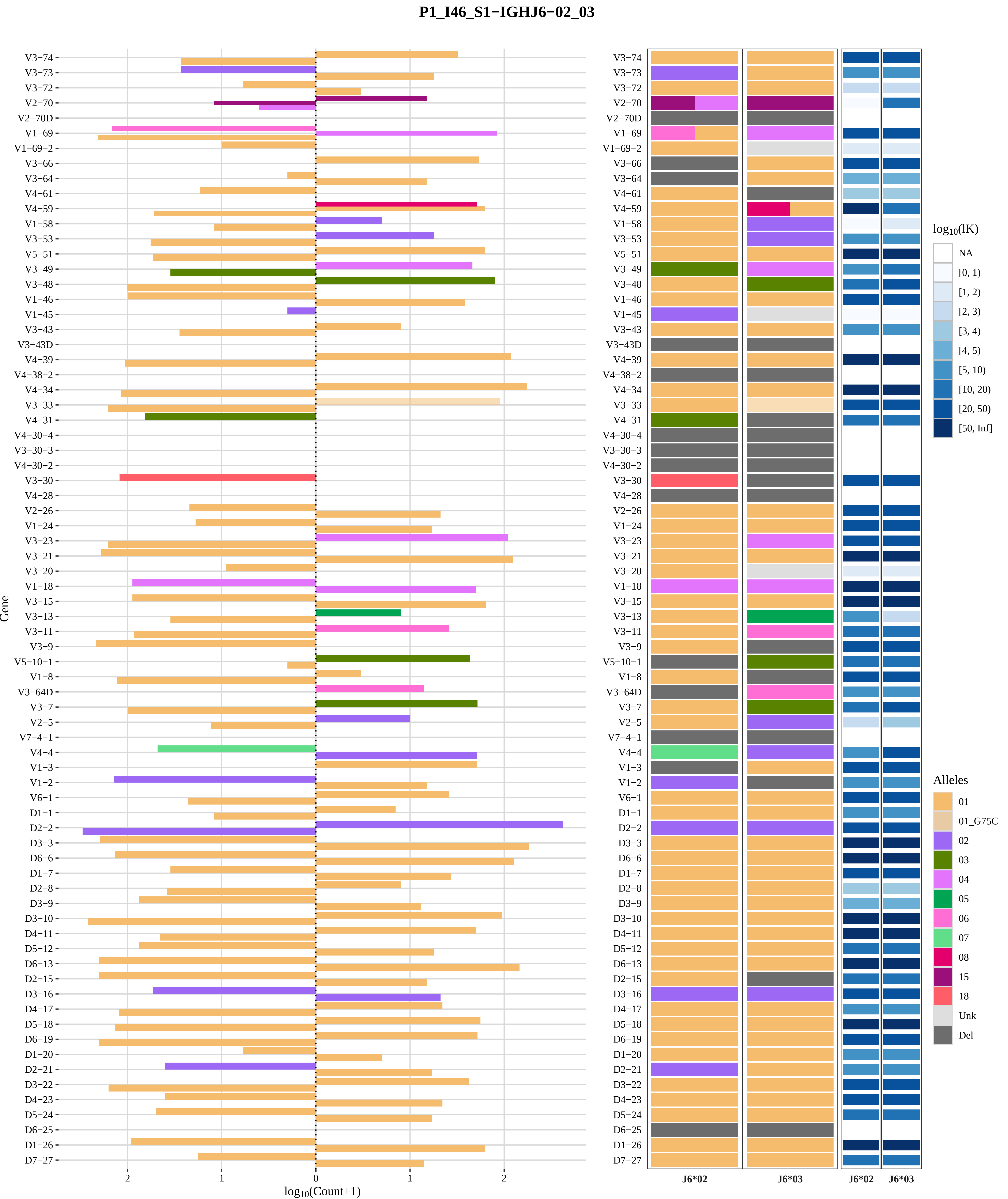
P1_I46_S1−IGHJ6−02_03
Gene
log10(Count+1)
J6*02
J6*03
log10(lK)
NA
[0, 1)
[1, 2)
[2, 3)
[3, 4)
[4, 5)
[5, 10)
[10, 20)
[20, 50)
[50, Inf]
Alleles
01
01_G75C
02
03
04
05
06
07
08
15
18
Unk
Del

### Genotype and haplotype of VDJbase sample I47 (study P1)

(illustrations downloaded from VDJbase in July, 2020)

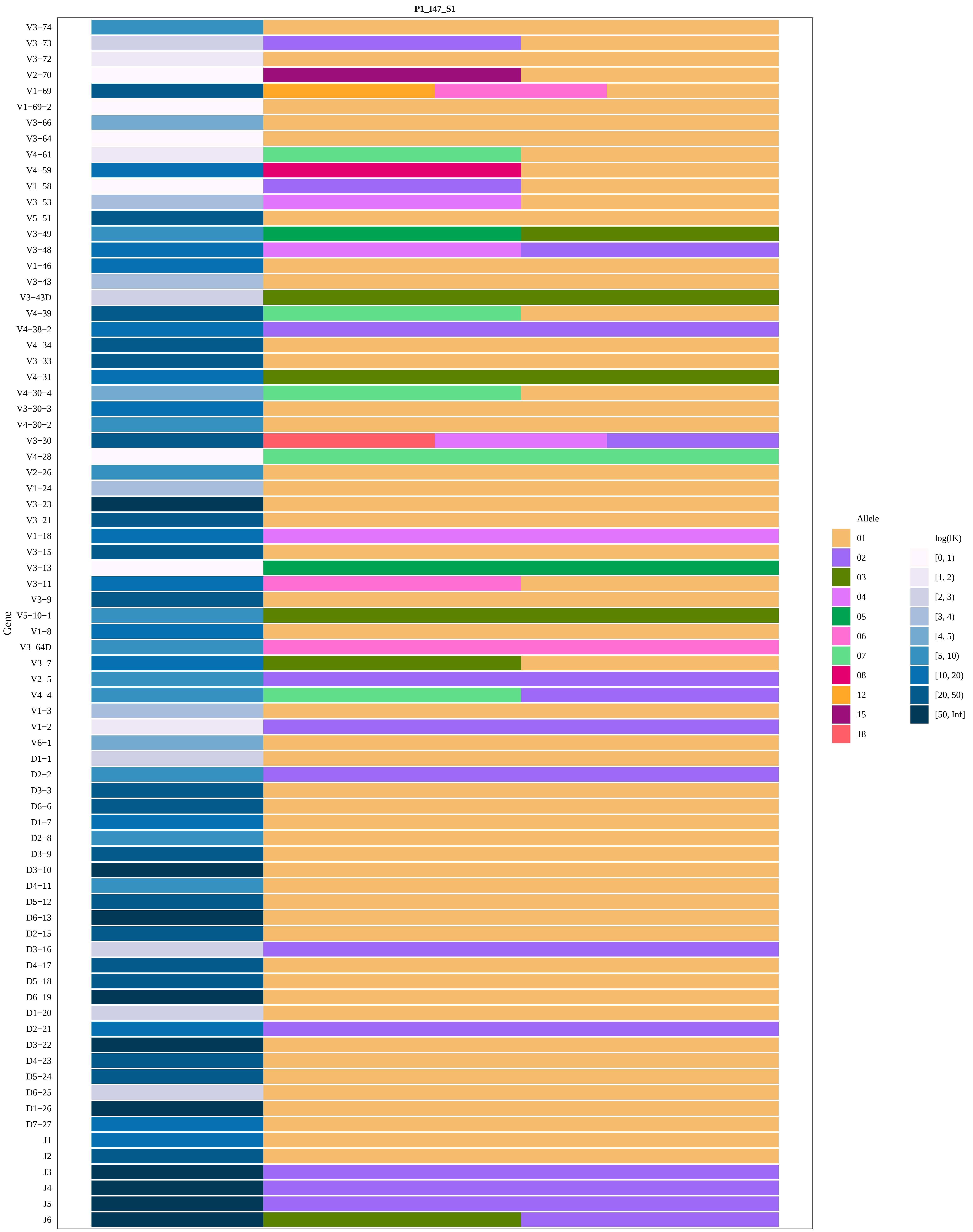

P1_I47_S1
Gene
V3−74
V3−73
V3−72
V2−70
V1−69
V1−69−2
V3−66
V3−64
V4−61
V4−59
V1−58
V3−53
V5−51
V3−49
V3−48
V1−46
V3−43
V3−43D
V4−39
V4−38−2
V4−34
V3−33
V4−31
V4−30−4
V3−30−3
V4−30−2
V3−30
V4−28
V2−26
V1−24
V3−23
V3−21
V1−18
V3−15
V3−13
V3−11
V3−9
V5−10−1
V1−8
V3−64D
V3−7
V2−5
V4−4
V1−3
V1−2
V6−1
D1−1
D2−2
D3−3
D6−6
D1−7
D2−8
D3−9
D3−10
D4−11
D5−12
D6−13
D2−15
D3−16
D4−17
D5−18
D6−19
D1−20
D2−21
D3−22
D4−23
D5−24
D6−25
D1−26
D7−27
J1
J2
J3
J4
J5
J6
Allele
01
02
03
04
05
06
07
08
12
15
18
log(lK)
[0, 1)
[1, 2)
[2, 3)
[3, 4)
[4, 5)
[5, 10)
[10, 20)
[20, 50)
[50, Inf]

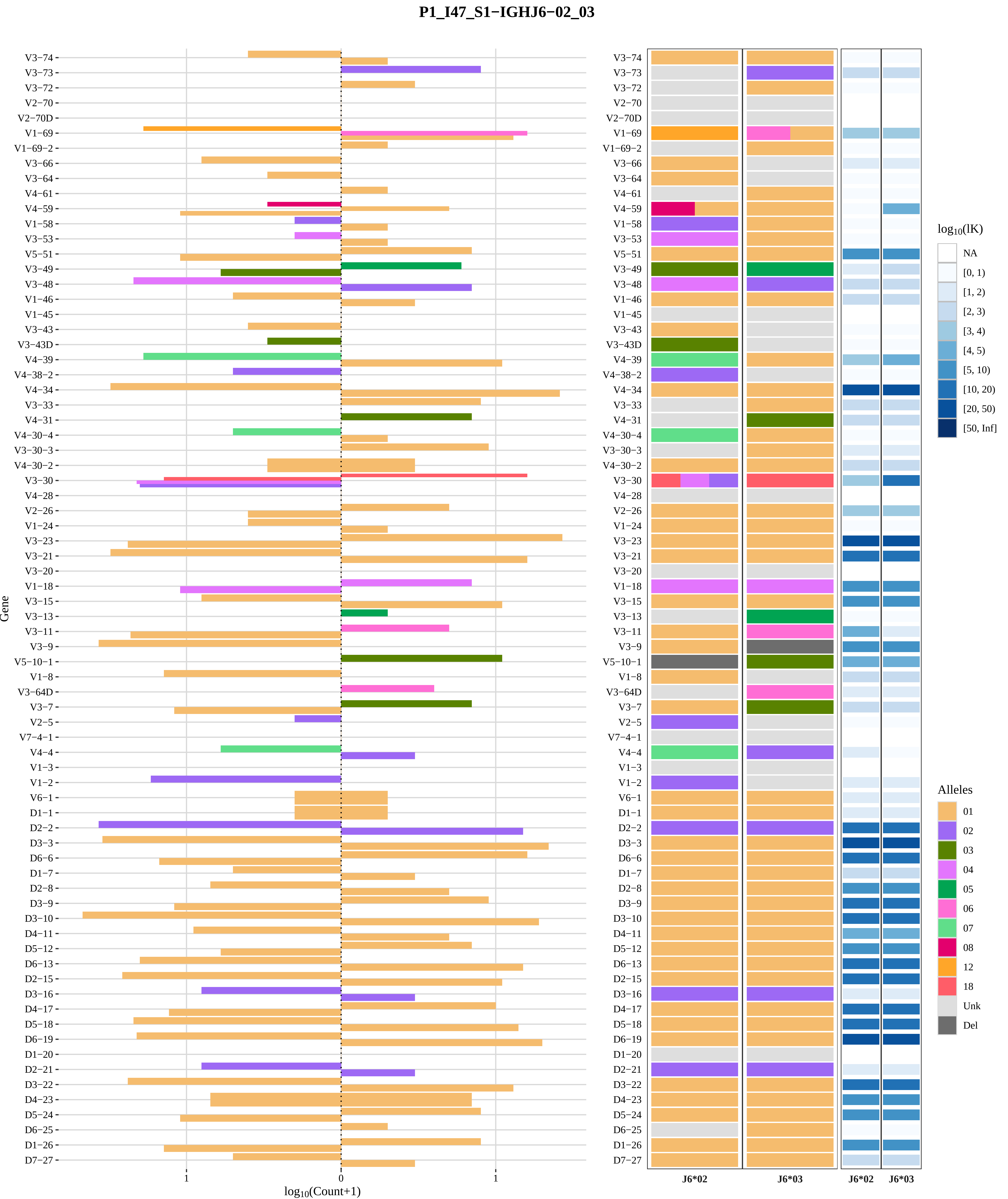

P1_I47_S1−IGHJ6−02_03
Gene
log10(Count+1)
J6*02
J6*03
log10(lK)
NA
[0, 1)
[1, 2)
[2, 3)
[3, 4)
[4, 5)
[5, 10)
[10, 20)
[20, 50)
[50, Inf]
Alleles
01
02
03
04
05
06
07
08
12
18
Unk
Del

### Genotype and haplotype of VDJbase sample I49 (study P1)

(illustrations downloaded from VDJbase in July, 2020)

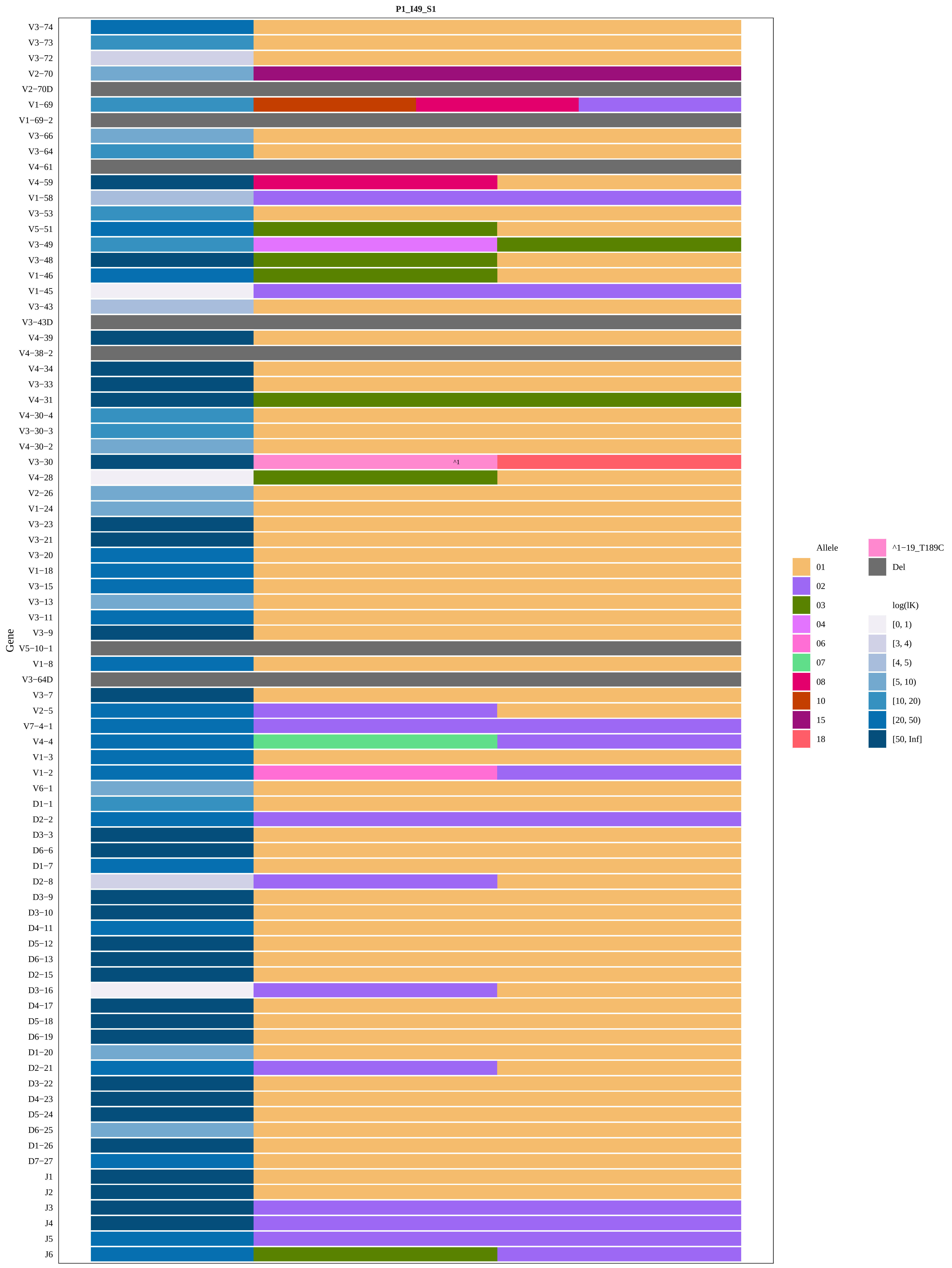
P1_I49_S1
Gene
V3−74
V3−73
V3−72
V2−70
V2−70D
V1−69
V1−69−2
V3−66
V3−64
V4−61
V4−59
V1−58
V3−53
V5−51
V3−49
V3−48
V1−46
V1−45
V3−43
V3−43D
V4−39
V4−38−2
V4−34
V3−33
V4−31
V4−30−4
V3−30−3
V4−30−2
V3−30
^1
V4−28
V2−26
V1−24
V3−23
V3−21
V3−20
V1−18
V3−15
V3−13
V3−11
V3−9
V5−10−1
V1−8
V3−64D
V3−7
V2−5
V7−4−1
V4−4
V1−3
V1−2
V6−1
D1−1
D2−2
D3−3
D6−6
D1−7
D2−8
D3−9
D3−10
D4−11
D5−12
D6−13
D2−15
D3−16
D4−17
D5−18
D6−19
D1−20
D2−21
D3−22
D4−23
D5−24
D6−25
D1−26
D7−27
J1
J2
J3
J4
J5
J6
Allele
01
02
03
04
06
07
08
10
15
18
^1−19_T189C
Del
log(lK)
[0, 1)
[3, 4)
[4, 5)
[5, 10)
[10, 20)
[20, 50)
[50, Inf]

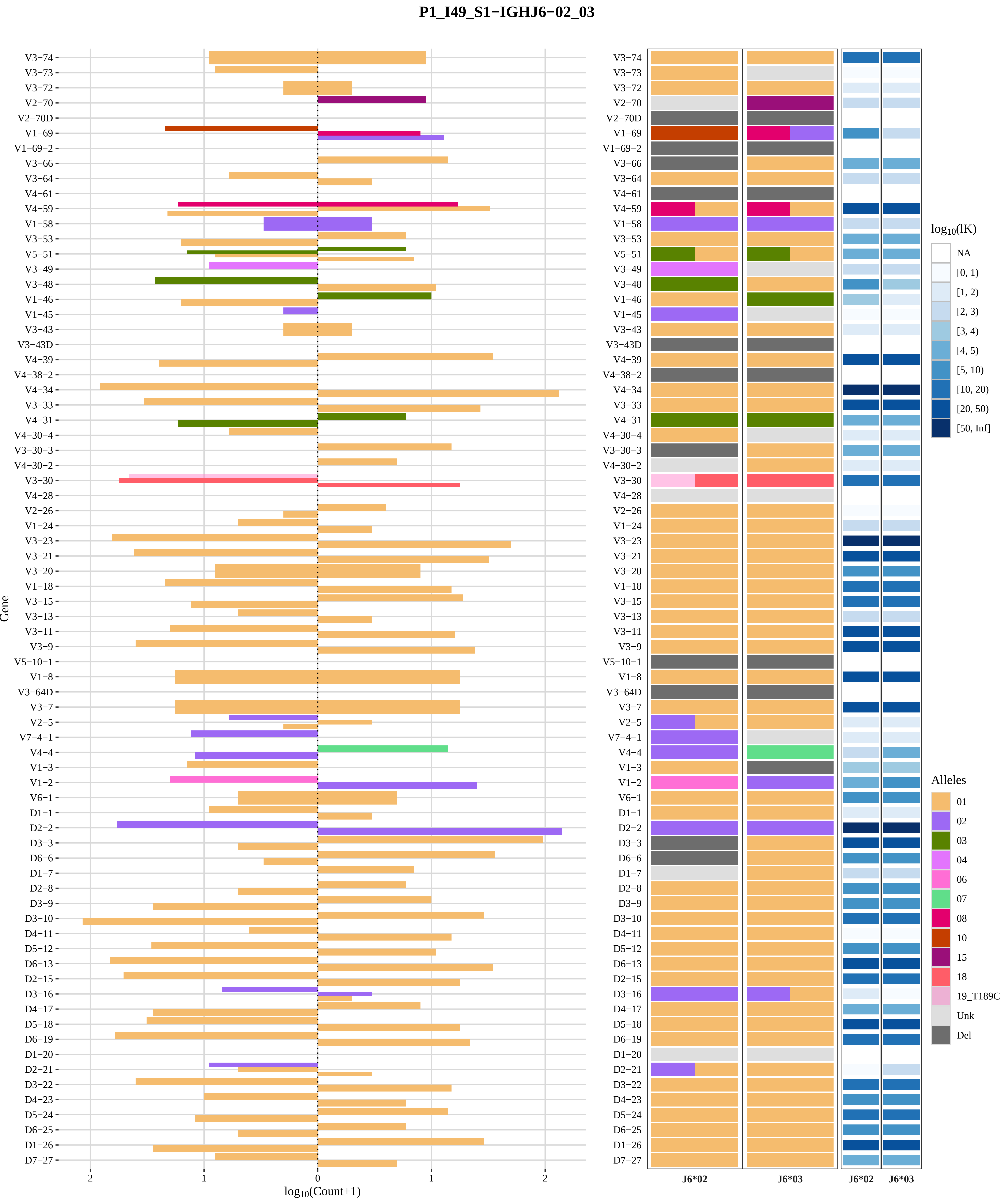
P1_I49_S1−IGHJ6−02_03
Gene
V3−74
V3−73
V3−72
V2−70
V2−70D
V1−69
V1−69−2
V3−66
V3−64
V4−61
V4−59
V1−58
V3−53
V5−51
V3−49
V3−48
V1−46
V1−45
V3−43
V3−43D
V4−39
V4−38−2
V4−34
V3−33
V4−31
V4−30−4
V3−30−3
V4−30−2
V3−30
V4−28
V2−26
V1−24
V3−23
V3−21
V3−20
V1−18
V3−15
V3−13
V3−11
V3−9
V5−10−1
V1−8
V3−64D
V3−7
V2−5
V7−4−1
V4−4
V1−3
V1−2
V6−1
D1−1
D2−2
D3−3
D6−6
D1−7
D2−8
D3−9
D3−10
D4−11
D5−12
D6−13
D2−15
D3−16
D4−17
D5−18
D6−19
D1−20
D2−21
D3−22
D4−23
D5−24
D6−25
D1−26
D7−27
2
1
0
1
2
log10(Count+1)
J6*02
J6*03
J6*02
J6*03
log10(lK)
NA
[0, 1)
[1, 2)
[2, 3)
[3, 4)
[4, 5)
[5, 10)
[10, 20)
[20, 50)
[50, Inf]
Alleles
01
02
03
04
06
07
08
10
15
18
19_T189C
Unk
Del

### Genotype and haplotype of VDJbase sample I52 (study P1)

(illustrations downloaded from VDJbase in July, 2020)

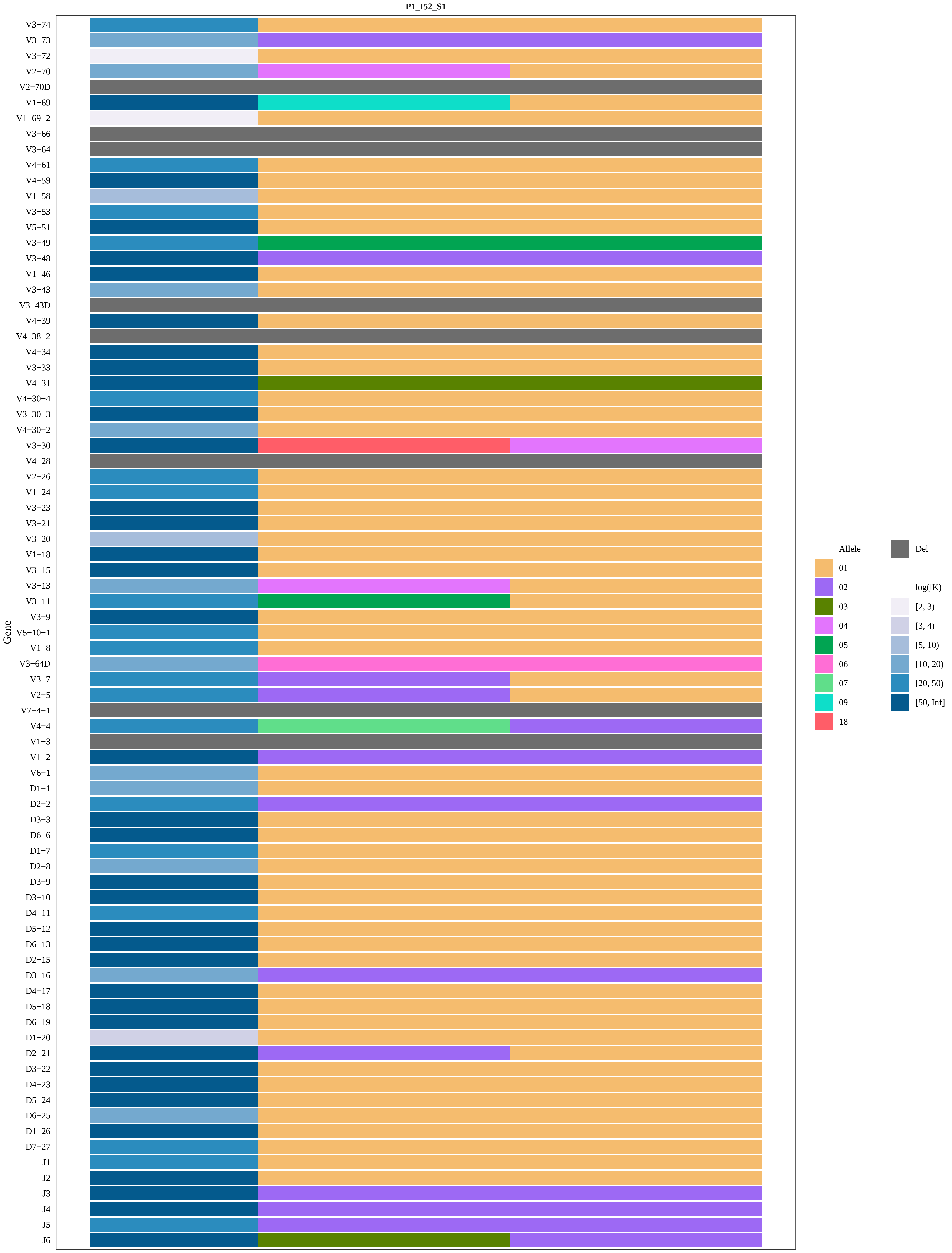
P1_I52_S1
Gene
V3−74
V3−73
V3−72
V2−70
V2−70D
V1−69
V1−69−2
V3−66
V3−64
V4−61
V4−59
V1−58
V3−53
V5−51
V3−49
V3−48
V1−46
V3−43
V3−43D
V4−39
V4−38−2
V4−34
V3−33
V4−31
V4−30−4
V3−30−3
V4−30−2
V3−30
V4−28
V2−26
V1−24
V3−23
V3−21
V3−20
V1−18
V3−15
V3−13
V3−11
V3−9
V5−10−1
V1−8
V3−64D
V3−7
V2−5
V7−4−1
V4−4
V1−3
V1−2
V6−1
D1−1
D2−2
D3−3
D6−6
D1−7
D2−8
D3−9
D3−10
D4−11
D5−12
D6−13
D2−15
D3−16
D4−17
D5−18
D6−19
D1−20
D2−21
D3−22
D4−23
D5−24
D6−25
D1−26
D7−27
J1
J2
J3
J4
J5
J6
Allele
01
02
03
04
05
06
07
09
18
Del
log(lK)
[2, 3)
[3, 4)
[5, 10)
[10, 20)
[20, 50)
[50, Inf]

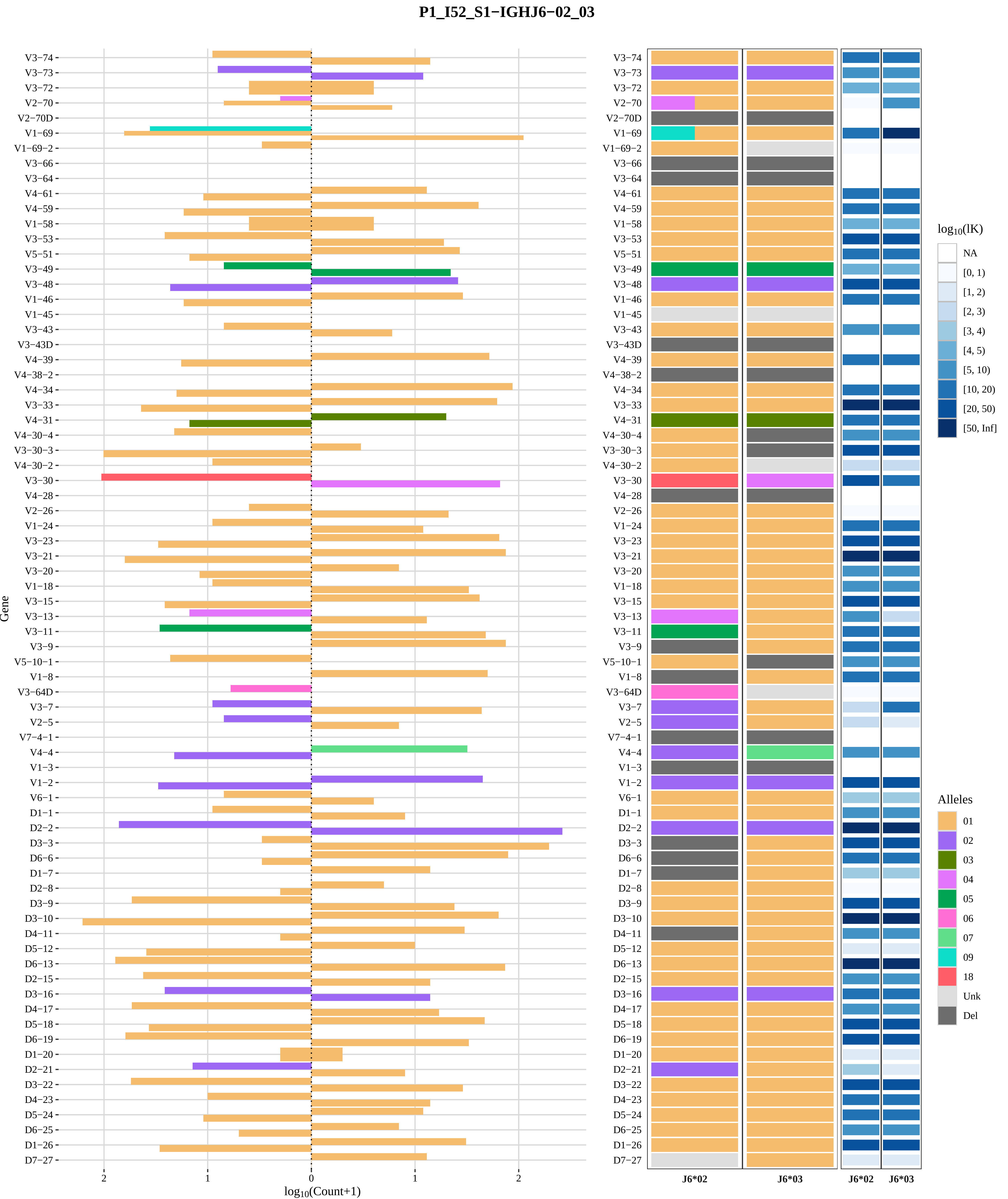
P1_I52_S1−IGHJ6−02_03
Gene
V3−74
V3−73
V3−72
V2−70
V2−70D
V1−69
V1−69−2
V3−66
V3−64
V4−61
V4−59
V1−58
V3−53
V5−51
V3−49
V3−48
V1−46
V1−45
V3−43
V3−43D
V4−39
V4−38−2
V4−34
V3−33
V4−31
V4−30−4
V3−30−3
V4−30−2
V3−30
V4−28
V2−26
V1−24
V3−23
V3−21
V3−20
V1−18
V3−15
V3−13
V3−11
V3−9
V5−10−1
V1−8
V3−64D
V3−7
V2−5
V7−4−1
V4−4
V1−3
V1−2
V6−1
D1−1
D2−2
D3−3
D6−6
D1−7
D2−8
D3−9
D3−10
D4−11
D5−12
D6−13
D2−15
D3−16
D4−17
D5−18
D6−19
D1−20
D2−21
D3−22
D4−23
D5−24
D6−25
D1−26
D7−27
2
1
0
1
2
log10(Count+1)
J6*02
J6*03
J6*02
J6*03
log10(lK)
NA
[0, 1)
[1, 2)
[2, 3)
[3, 4)
[4, 5)
[5, 10)
[10, 20)
[20, 50)
[50, Inf]
Alleles
01
02
03
04
05
06
07
09
18
Unk
Del

### Genotype and haplotype of VDJbase sample I56 (study P1)

(illustrations downloaded from VDJbase in July, 2020)

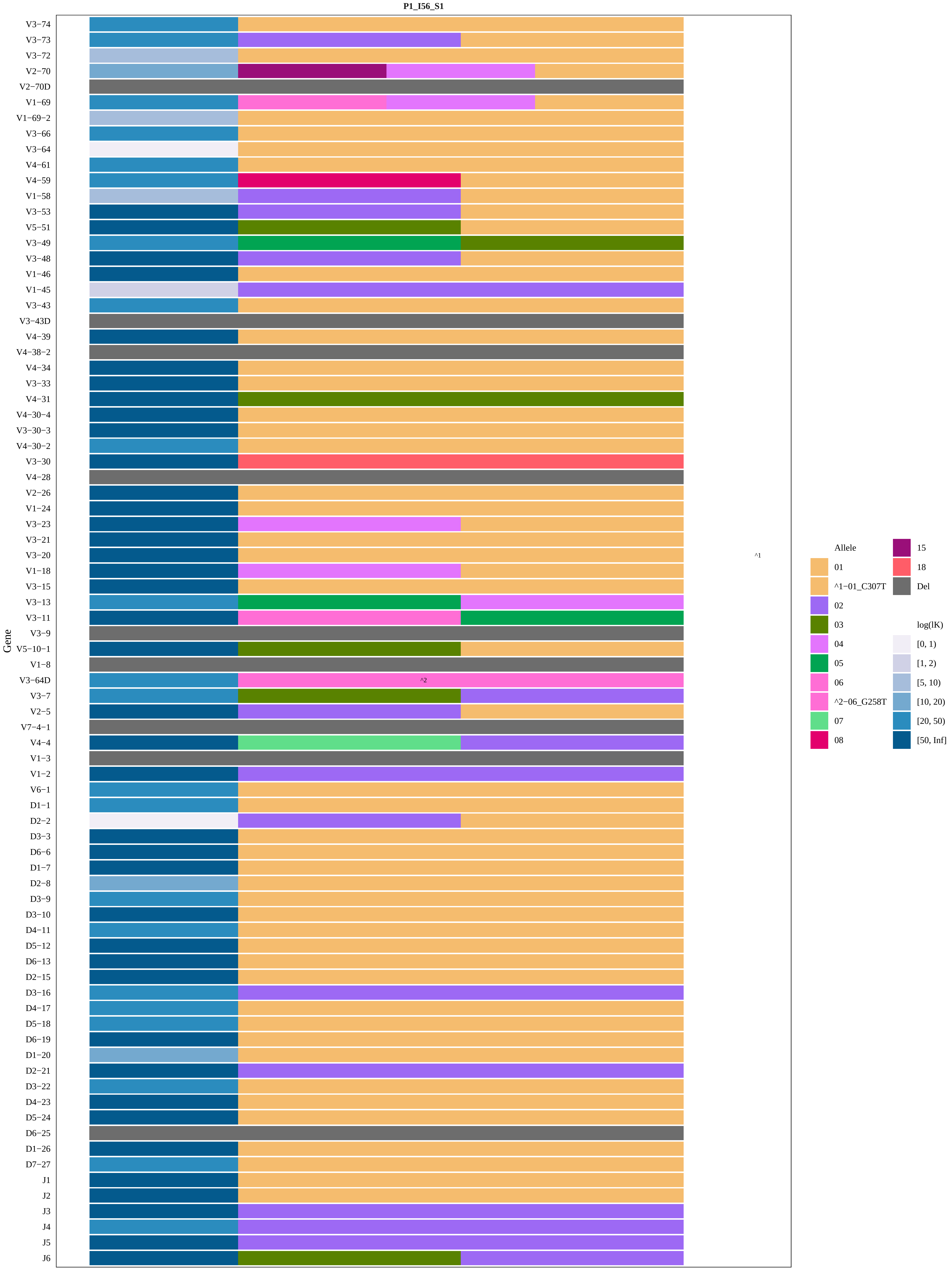

P1_I56_S1
Gene
V3−74
V3−73
V3−72
V2−70
V2−70D
V1−69
V1−69−2
V3−66
V3−64
V4−61
V4−59
V1−58
V3−53
V5−51
V3−49
V3−48
V1−46
V1−45
V3−43
V3−43D
V4−39
V4−38−2
V4−34
V3−33
V4−31
V4−30−4
V3−30−3
V4−30−2
V3−30
V4−28
V2−26
V1−24
V3−23
V3−21
V3−20
V1−18
V3−15
V3−13
V3−11
V3−9
V5−10−1
V1−8
V3−64D
V3−7
V2−5
V7−4−1
V4−4
V1−3
V1−2
V6−1
D1−1
D2−2
D3−3
D6−6
D1−7
D2−8
D3−9
D3−10
D4−11
D5−12
D6−13
D2−15
D3−16
D4−17
D5−18
D6−19
D1−20
D2−21
D3−22
D4−23
D5−24
D6−25
D1−26
D7−27
J1
J2
J3
J4
J5
J6
^2
^1
Allele
01
^1−01_C307T
02
03
04
05
06
^2−06_G258T
07
08
15
18
Del
log(lK)
[0, 1)
[1, 2)
[5, 10)
[10, 20)
[20, 50)
[50, Inf]

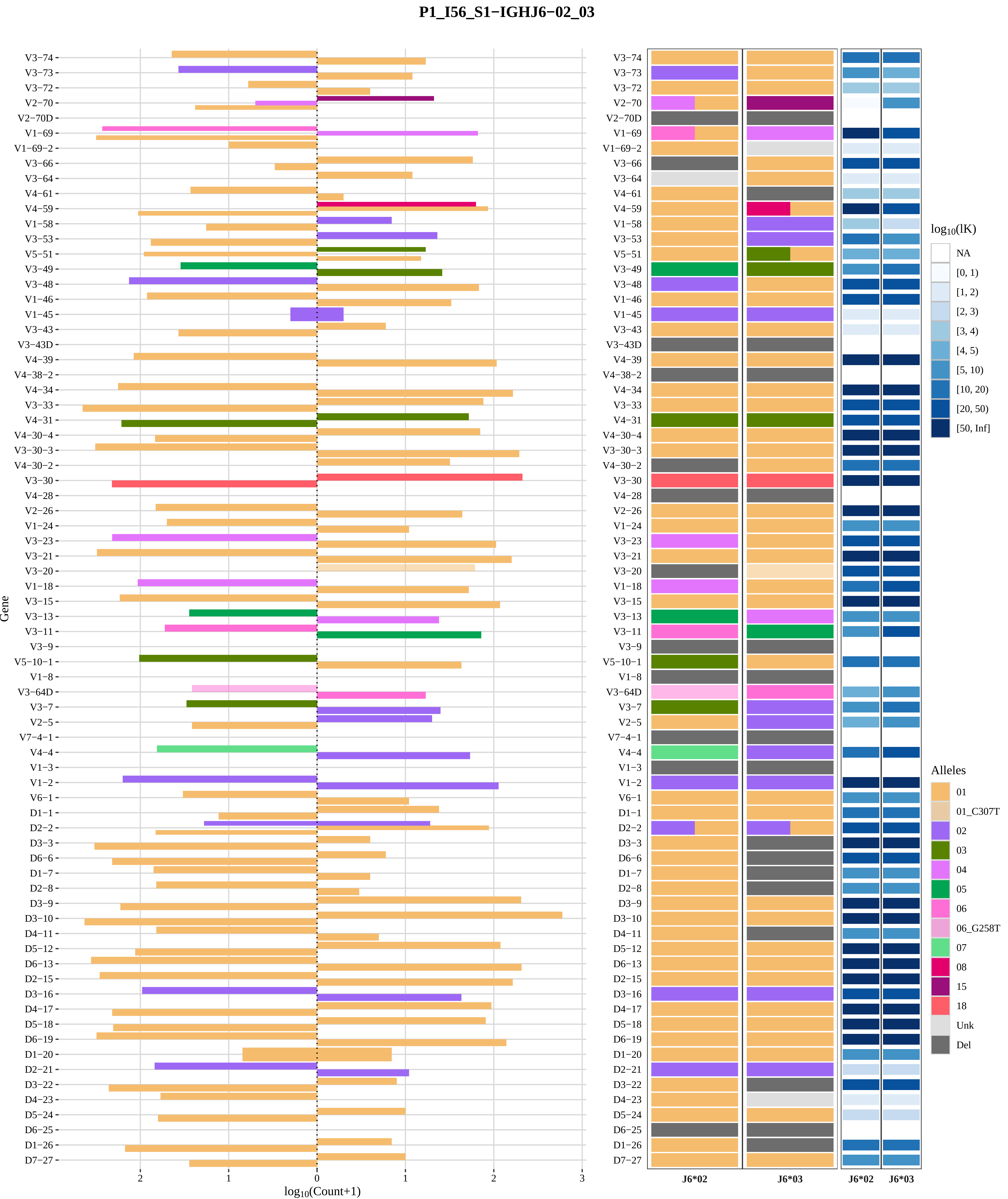

P1_I56_S1−IGHJ6−02_03
Gene
log10(Count+1)
J6*02
J6*03
log10(lK)
NA
[0, 1)
[1, 2)
[2, 3)
[3, 4)
[4, 5)
[5, 10)
[10, 20)
[20, 50)
[50, Inf]
Alleles
01
01_C307T
02
03
04
05
06
06_G258T
07
08
15
18
Unk
Del
V3−74
V3−73
V3−72
V2−70
V2−70D
V1−69
V1−69−2
V3−66
V3−64
V4−61
V4−59
V1−58
V3−53
V5−51
V3−49
V3−48
V1−46
V1−45
V3−43
V3−43D
V4−39
V4−38−2
V4−34
V3−33
V4−31
V4−30−4
V3−30−3
V4−30−2
V3−30
V4−28
V2−26
V1−24
V3−23
V3−21
V3−20
V1−18
V3−15
V3−13
V3−11
V3−9
V5−10−1
V1−8
V3−64D
V3−7
V2−5
V7−4−1
V4−4
V1−3
V1−2
V6−1
D1−1
D2−2
D3−3
D6−6
D1−7
D2−8
D3−9
D3−10
D4−11
D5−12
D6−13
D2−15
D3−16
D4−17
D5−18
D6−19
D1−20
D2−21
D3−22
D4−23
D5−24
D6−25
D1−26
D7−27

### Genotype and haplotype of VDJbase sample I57 (study P1)

(illustrations downloaded from VDJbase in July, 2020)

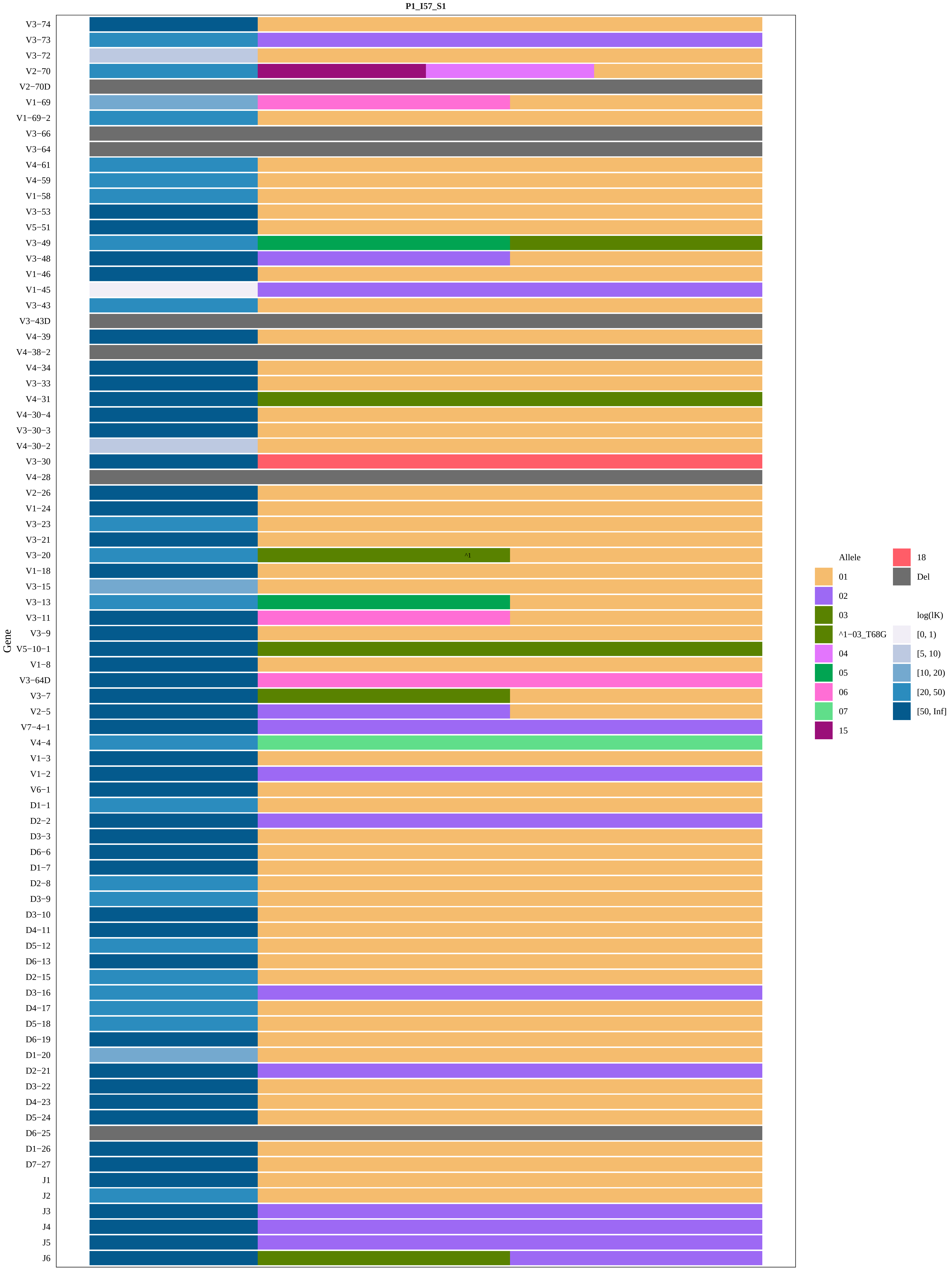
P1_I57_S1
Gene
V3−74
V3−73
V3−72
V2−70
V2−70D
V1−69
V1−69−2
V3−66
V3−64
V4−61
V4−59
V1−58
V3−53
V5−51
V3−49
V3−48
V1−46
V1−45
V3−43
V3−43D
V4−39
V4−38−2
V4−34
V3−33
V4−31
V4−30−4
V3−30−3
V4−30−2
V3−30
V4−28
V2−26
V1−24
V3−23
V3−21
V3−20
^1
V1−18
V3−15
V3−13
V3−11
V3−9
V5−10−1
V1−8
V3−64D
V3−7
V2−5
V7−4−1
V4−4
V1−3
V1−2
V6−1
D1−1
D2−2
D3−3
D6−6
D1−7
D2−8
D3−9
D3−10
D4−11
D5−12
D6−13
D2−15
D3−16
D4−17
D5−18
D6−19
D1−20
D2−21
D3−22
D4−23
D5−24
D6−25
D1−26
D7−27
J1
J2
J3
J4
J5
J6
Allele
01
02
03
^1−03_T68G
04
05
06
07
15
18
Del
log(lK)
[0, 1)
[5, 10)
[10, 20)
[20, 50)
[50, Inf]

**P1_I57_S1−IGHJ6−02_03**

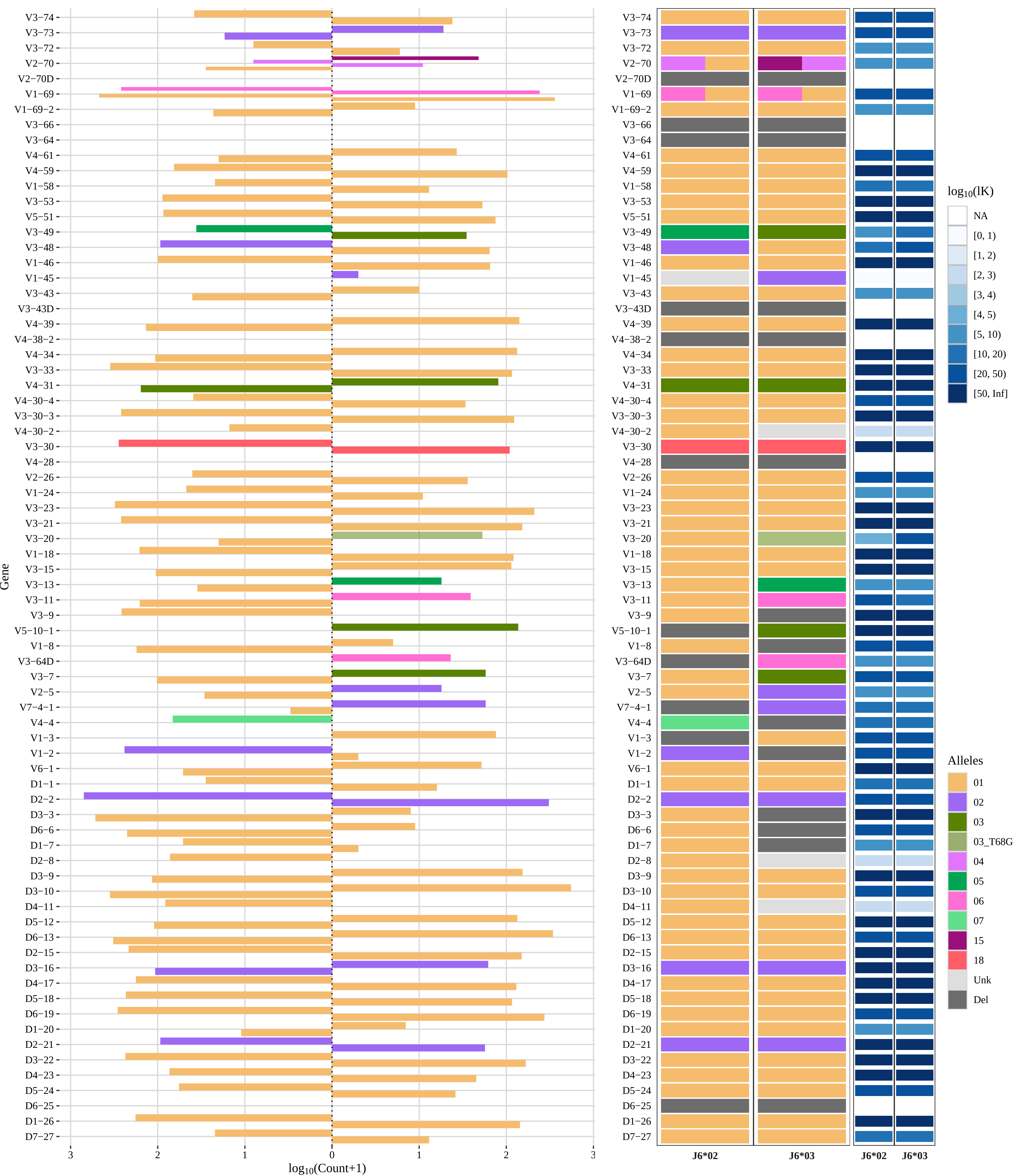

### Genotype and haplotype of VDJbase sample I58 (study P1)

(illustrations downloaded from VDJbase in July, 2020)

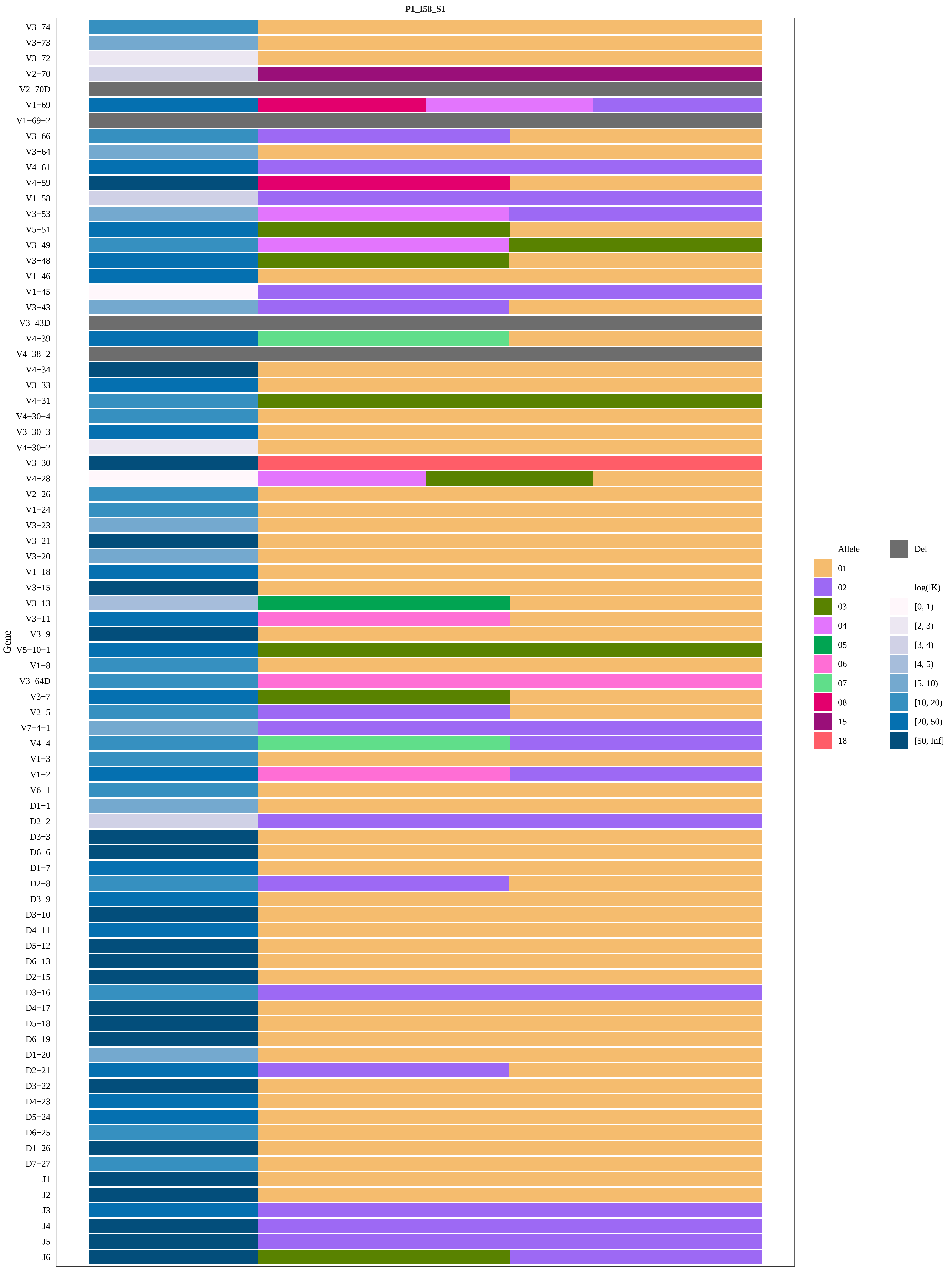
P1_I58_S1
Gene
V3−74
V3−73
V3−72
V2−70
V2−70D
V1−69
V1−69−2
V3−66
V3−64
V4−61
V4−59
V1−58
V3−53
V5−51
V3−49
V3−48
V1−46
V1−45
V3−43
V3−43D
V4−39
V4−38−2
V4−34
V3−33
V4−31
V4−30−4
V3−30−3
V4−30−2
V3−30
V4−28
V2−26
V1−24
V3−23
V3−21
V3−20
V1−18
V3−15
V3−13
V3−11
V3−9
V5−10−1
V1−8
V3−64D
V3−7
V2−5
V7−4−1
V4−4
V1−3
V1−2
V6−1
D1−1
D2−2
D3−3
D6−6
D1−7
D2−8
D3−9
D3−10
D4−11
D5−12
D6−13
D2−15
D3−16
D4−17
D5−18
D6−19
D1−20
D2−21
D3−22
D4−23
D5−24
D6−25
D1−26
D7−27
J1
J2
J3
J4
J5
J6
Allele
01
02
03
04
05
06
07
08
15
18
Del
log(lK)
[0, 1)
[2, 3)
[3, 4)
[4, 5)
[5, 10)
[10, 20)
[20, 50)
[50, Inf]

#### P1_I58_S1−IGHJ6−02_03

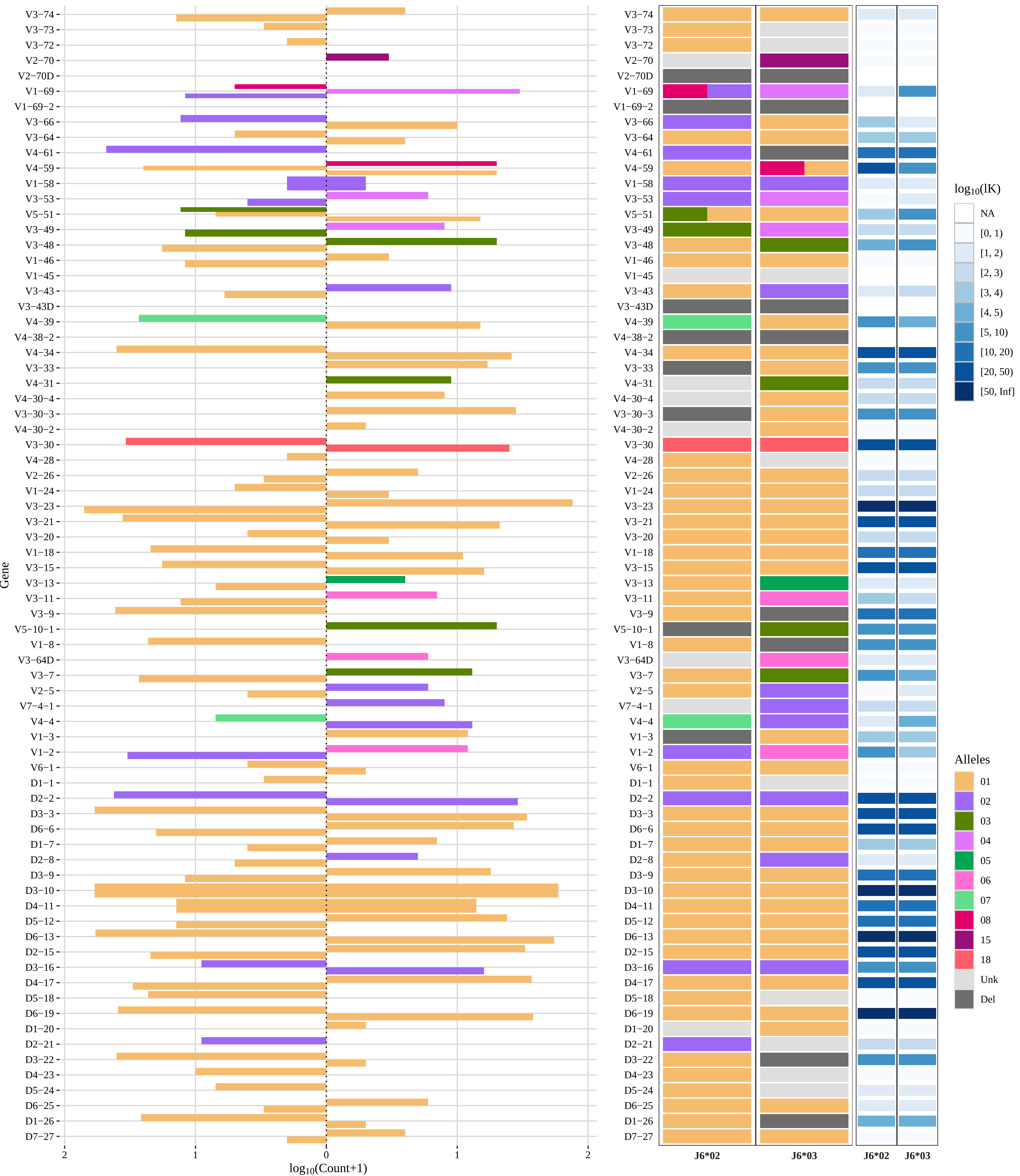

### Genotype and haplotype of VDJbase sample I67 (study P1)

(illustrations downloaded from VDJbase in July, 2020)

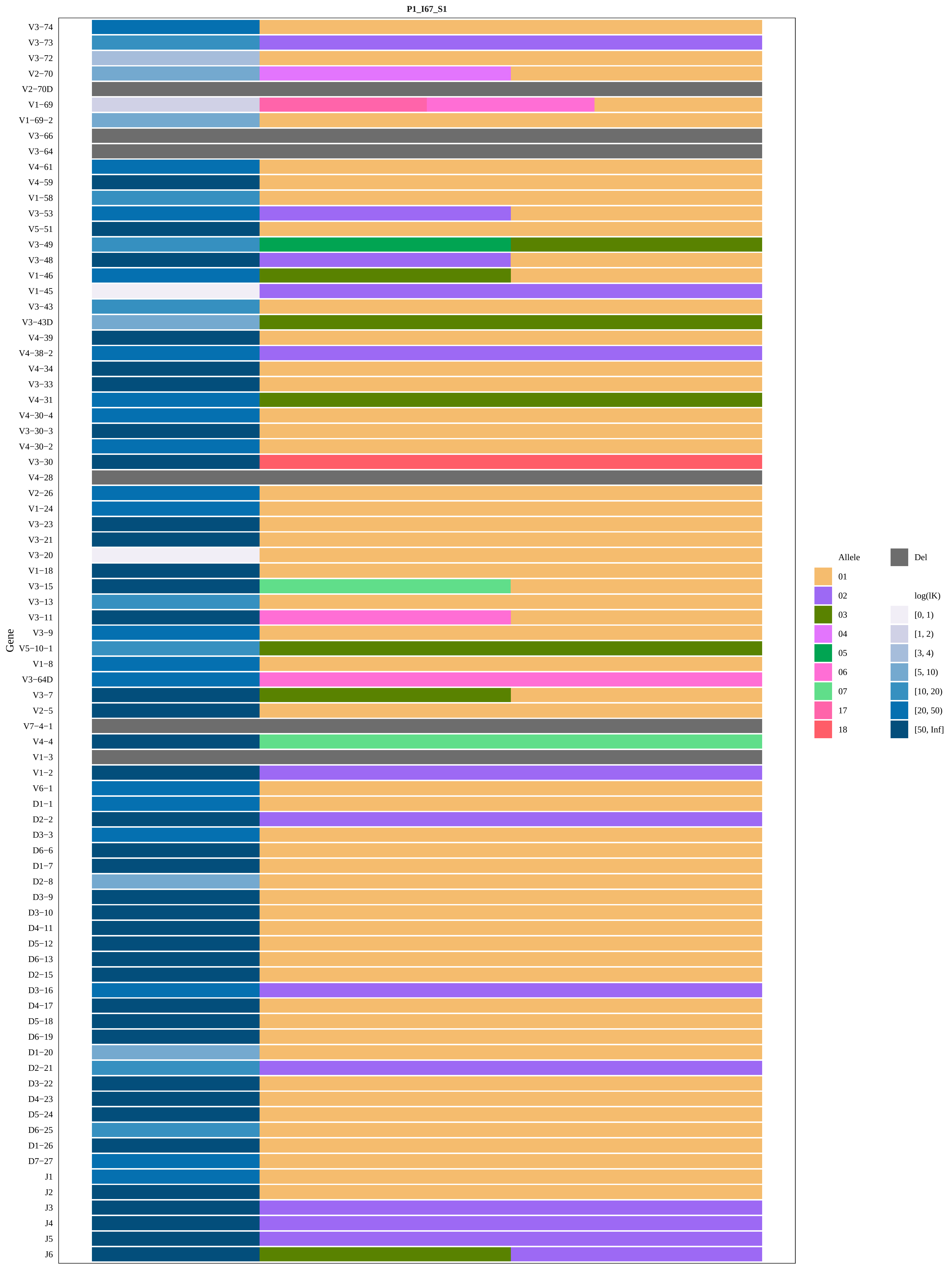

P1_I67_S1
Gene
V3−74
V3−73
V3−72
V2−70
V2−70D
V1−69
V1−69−2
V3−66
V3−64
V4−61
V4−59
V1−58
V3−53
V5−51
V3−49
V3−48
V1−46
V1−45
V3−43
V3−43D
V4−39
V4−38−2
V4−34
V3−33
V4−31
V4−30−4
V3−30−3
V4−30−2
V3−30
V4−28
V2−26
V1−24
V3−23
V3−21
V3−20
V1−18
V3−15
V3−13
V3−11
V3−9
V5−10−1
V1−8
V3−64D
V3−7
V2−5
V7−4−1
V4−4
V1−3
V1−2
V6−1
D1−1
D2−2
D3−3
D6−6
D1−7
D2−8
D3−9
D3−10
D4−11
D5−12
D6−13
D2−15
D3−16
D4−17
D5−18
D6−19
D1−20
D2−21
D3−22
D4−23
D5−24
D6−25
D1−26
D7−27
J1
J2
J3
J4
J5
J6
Allele
01
02
03
04
05
06
07
17
18
Del
log(lK)
[0, 1)
[1, 2)
[3, 4)
[5, 10)
[10, 20)
[20, 50)
[50, Inf]

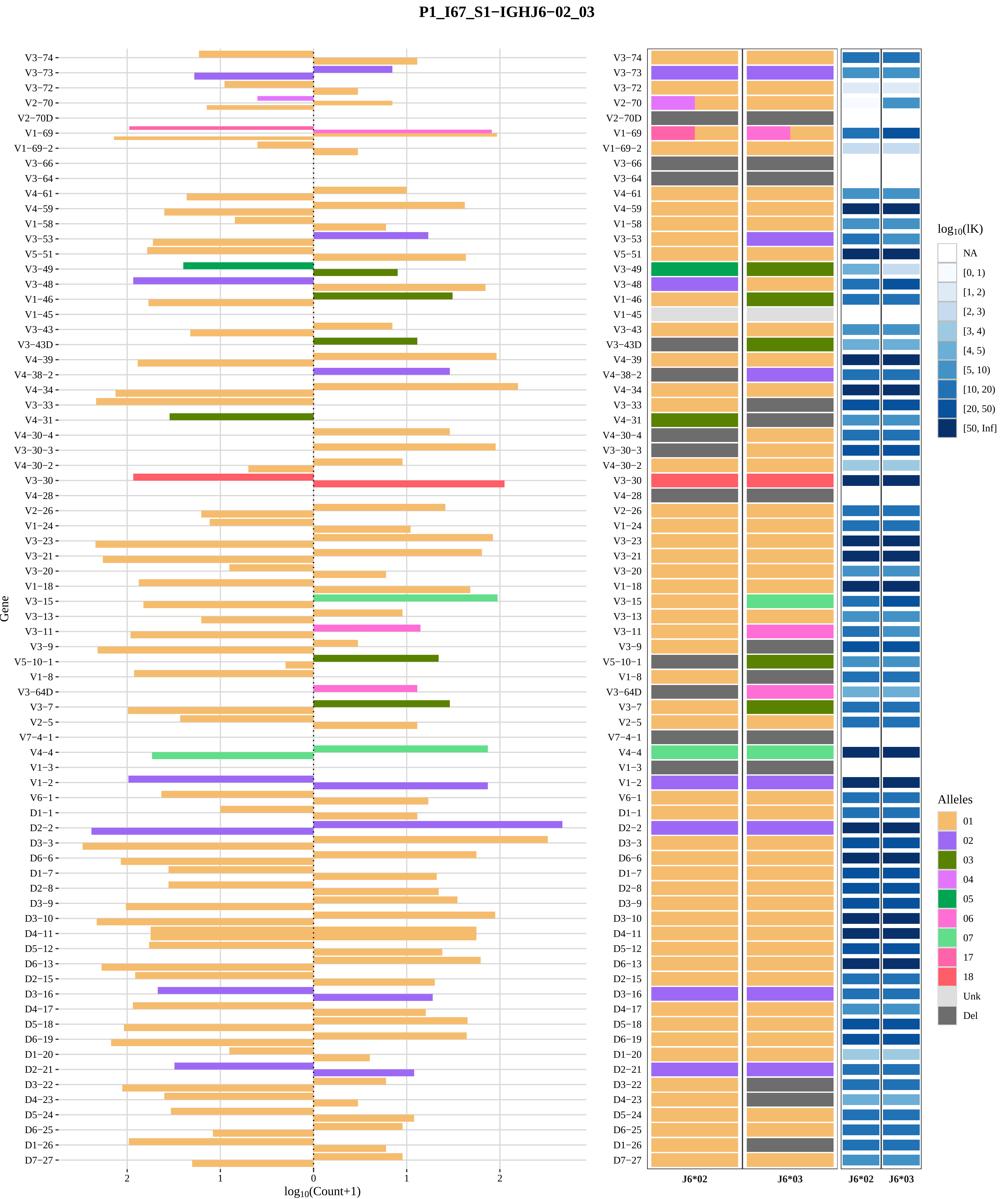
P1_I67_S1−IGHJ6−02_03
Gene
log10(Count+1)
J6*02
J6*03
log10(lK)
NA
[0, 1)
[1, 2)
[2, 3)
[3, 4)
[4, 5)
[5, 10)
[10, 20)
[20, 50)
[50, Inf]
Alleles
01
02
03
04
05
06
07
17
18
Unk
Del
V3−74
V3−73
V3−72
V2−70
V2−70D
V1−69
V1−69−2
V3−66
V3−64
V4−61
V4−59
V1−58
V3−53
V5−51
V3−49
V3−48
V1−46
V1−45
V3−43
V3−43D
V4−39
V4−38−2
V4−34
V3−33
V4−31
V4−30−4
V3−30−3
V4−30−2
V3−30
V4−28
V2−26
V1−24
V3−23
V3−21
V3−20
V1−18
V3−15
V3−13
V3−11
V3−9
V5−10−1
V1−8
V3−64D
V3−7
V2−5
V7−4−1
V4−4
V1−3
V1−2
V6−1
D1−1
D2−2
D3−3
D6−6
D1−7
D2−8
D3−9
D3−10
D4−11
D5−12
D6−13
D2−15
D3−16
D4−17
D5−18
D6−19
D1−20
D2−21
D3−22
D4−23
D5−24
D6−25
D1−26
D7−27

### Genotype and haplotype of VDJbase sample I69 (study P1)

(illustrations downloaded from VDJbase in July, 2020)

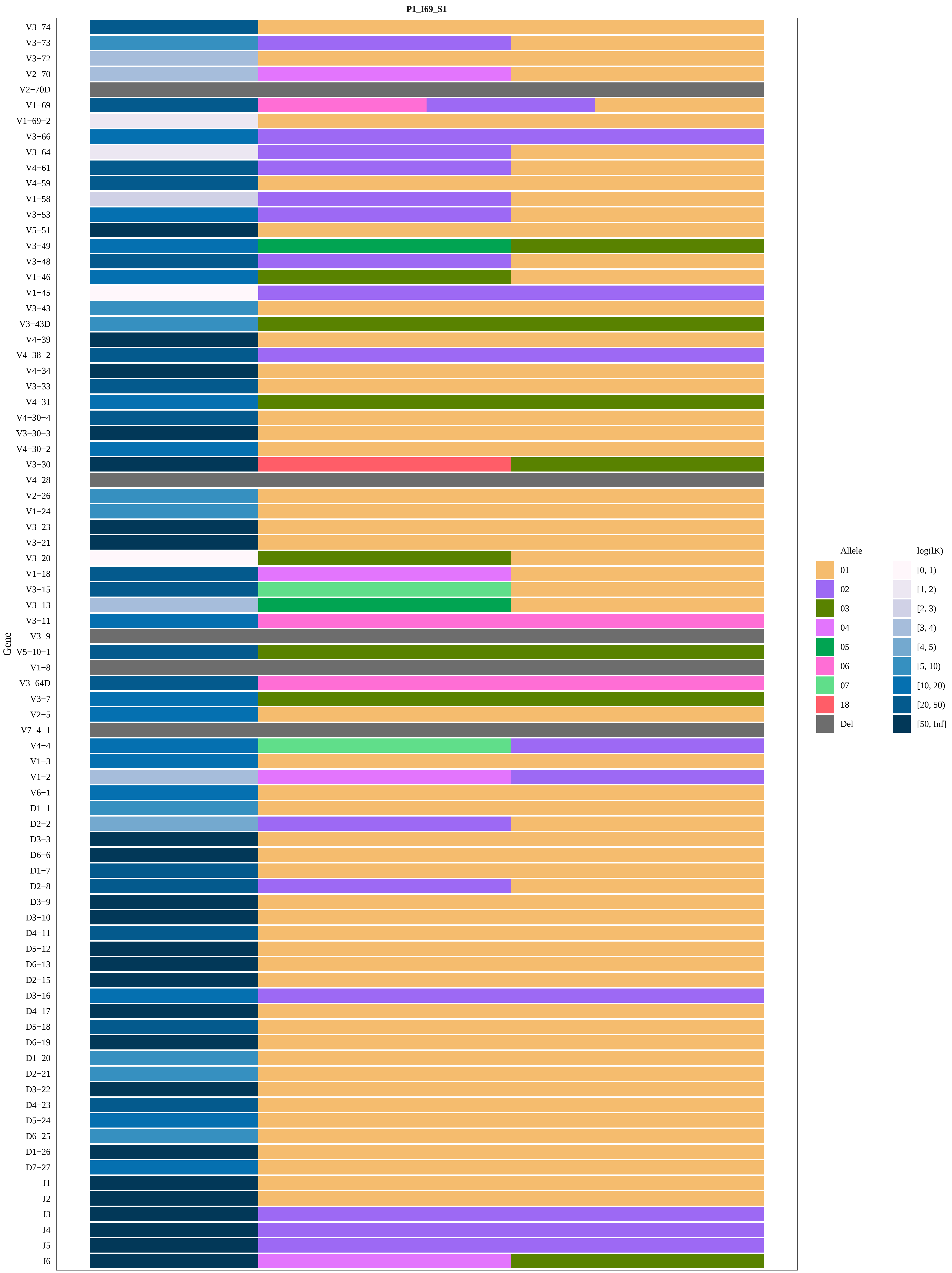
P1_I69_S1
Gene
V3−74
V3−73
V3−72
V2−70
V2−70D
V1−69
V1−69−2
V3−66
V3−64
V4−61
V4−59
V1−58
V3−53
V5−51
V3−49
V3−48
V1−46
V1−45
V3−43
V3−43D
V4−39
V4−38−2
V4−34
V3−33
V4−31
V4−30−4
V3−30−3
V4−30−2
V3−30
V4−28
V2−26
V1−24
V3−23
V3−21
V3−20
V1−18
V3−15
V3−13
V3−11
V3−9
V5−10−1
V1−8
V3−64D
V3−7
V2−5
V7−4−1
V4−4
V1−3
V1−2
V6−1
D1−1
D2−2
D3−3
D6−6
D1−7
D2−8
D3−9
D3−10
D4−11
D5−12
D6−13
D2−15
D3−16
D4−17
D5−18
D6−19
D1−20
D2−21
D3−22
D4−23
D5−24
D6−25
D1−26
D7−27
J1
J2
J3
J4
J5
J6
Allele
01
02
03
04
05
06
07
18
Del
log(lK)
[0, 1)
[1, 2)
[2, 3)
[3, 4)
[4, 5)
[5, 10)
[10, 20)
[20, 50)
[50, Inf]

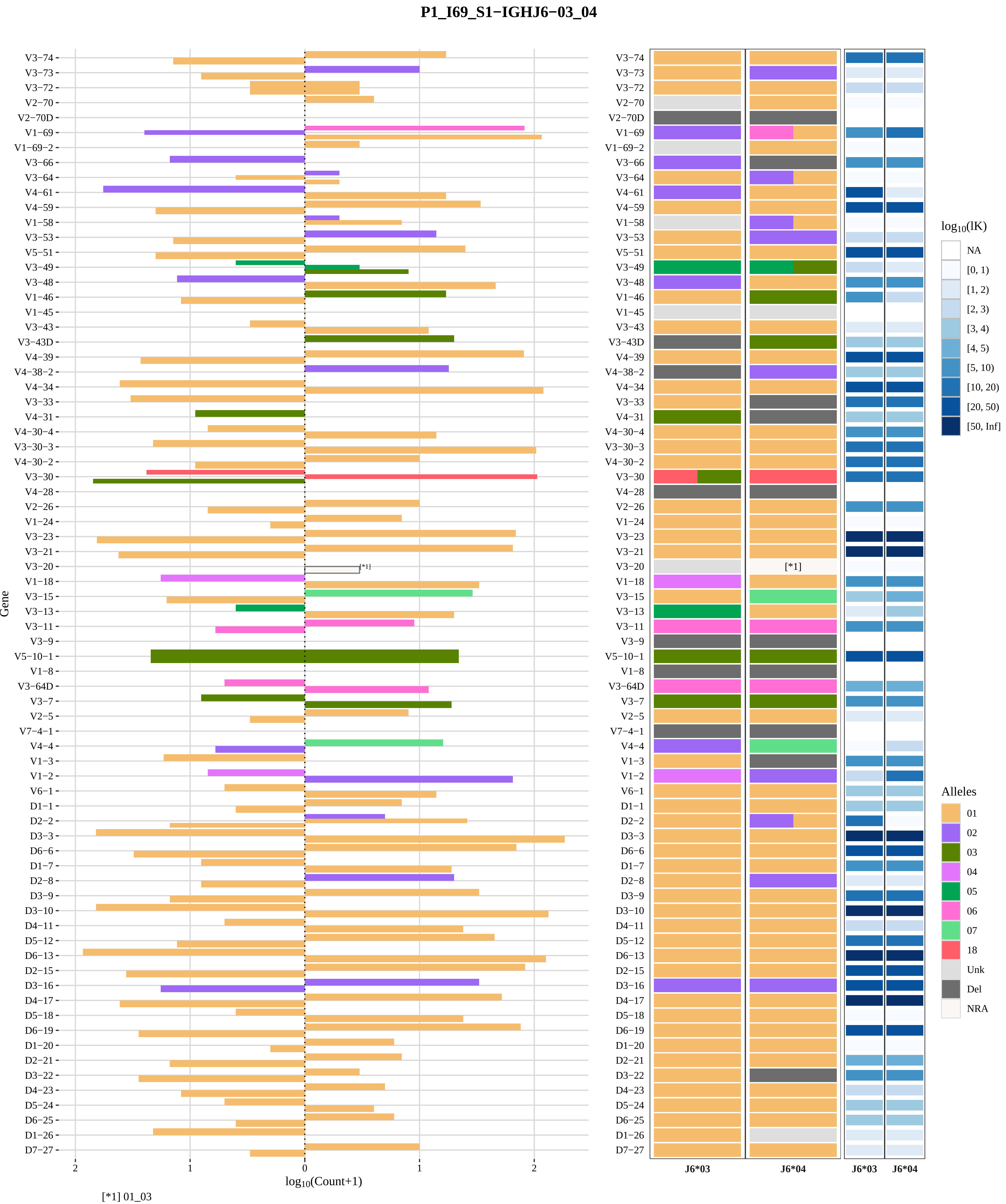

P1_I69_S1−IGHJ6−03_04
Gene
log10(Count+1)
[*1] 01_03
log10(lK)
NA
[0, 1)
[1, 2)
[2, 3)
[3, 4)
[4, 5)
[5, 10)
[10, 20)
[20, 50)
[50, Inf]
Alleles
01
02
03
04
05
06
07
18
Unk
Del
NRA
J6*03
J6*04
J6*03
J6*04

### Genotype and haplotype of VDJbase sample I70 (study P1)

(illustrations downloaded from VDJbase in July, 2020)

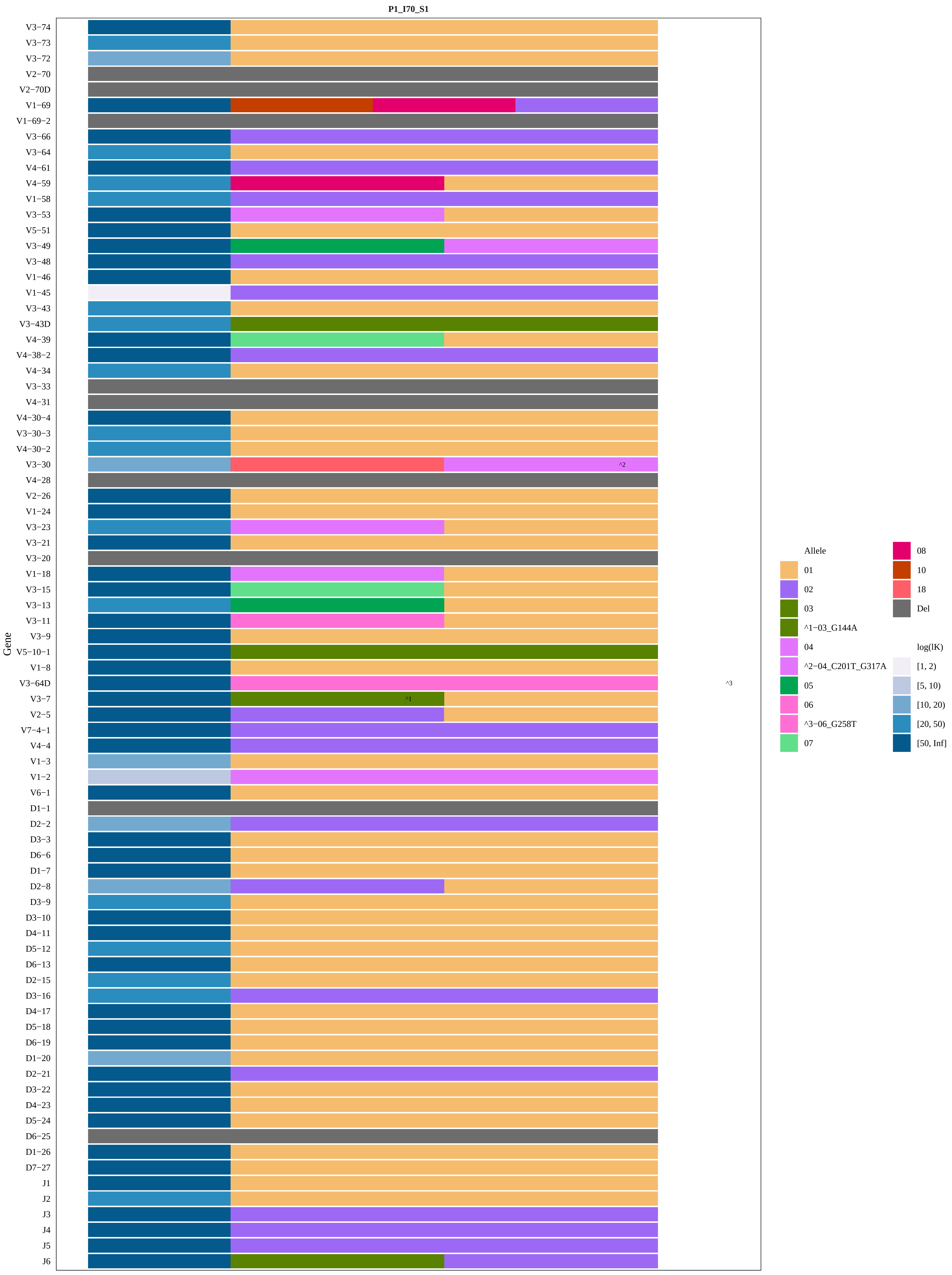
P1_I70_S1
Gene
V3−74
V3−73
V3−72
V2−70
V2−70D
V1−69
V1−69−2
V3−66
V3−64
V4−61
V4−59
V1−58
V3−53
V5−51
V3−49
V3−48
V1−46
V1−45
V3−43
V3−43D
V4−39
V4−38−2
V4−34
V3−33
V4−31
V4−30−4
V3−30−3
V4−30−2
V3−30
V4−28
V2−26
V1−24
V3−23
V3−21
V3−20
V1−18
V3−15
V3−13
V3−11
V3−9
V5−10−1
V1−8
V3−64D
V3−7
V2−5
V7−4−1
V4−4
V1−3
V1−2
V6−1
D1−1
D2−2
D3−3
D6−6
D1−7
D2−8
D3−9
D3−10
D4−11
D5−12
D6−13
D2−15
D3−16
D4−17
D5−18
D6−19
D1−20
D2−21
D3−22
D4−23
D5−24
D6−25
D1−26
D7−27
J1
J2
J3
J4
J5
J6
^1
^2
^3
Allele
01
02
03
^1−03_G144A
04
^2−04_C201T_G317A
05
06
^3−06_G258T
07
08
10
18
Del
log(lK)
[1, 2)
[5, 10)
[10, 20)
[20, 50)
[50, Inf]

### P1_I70_S1−IGHJ6−02_03

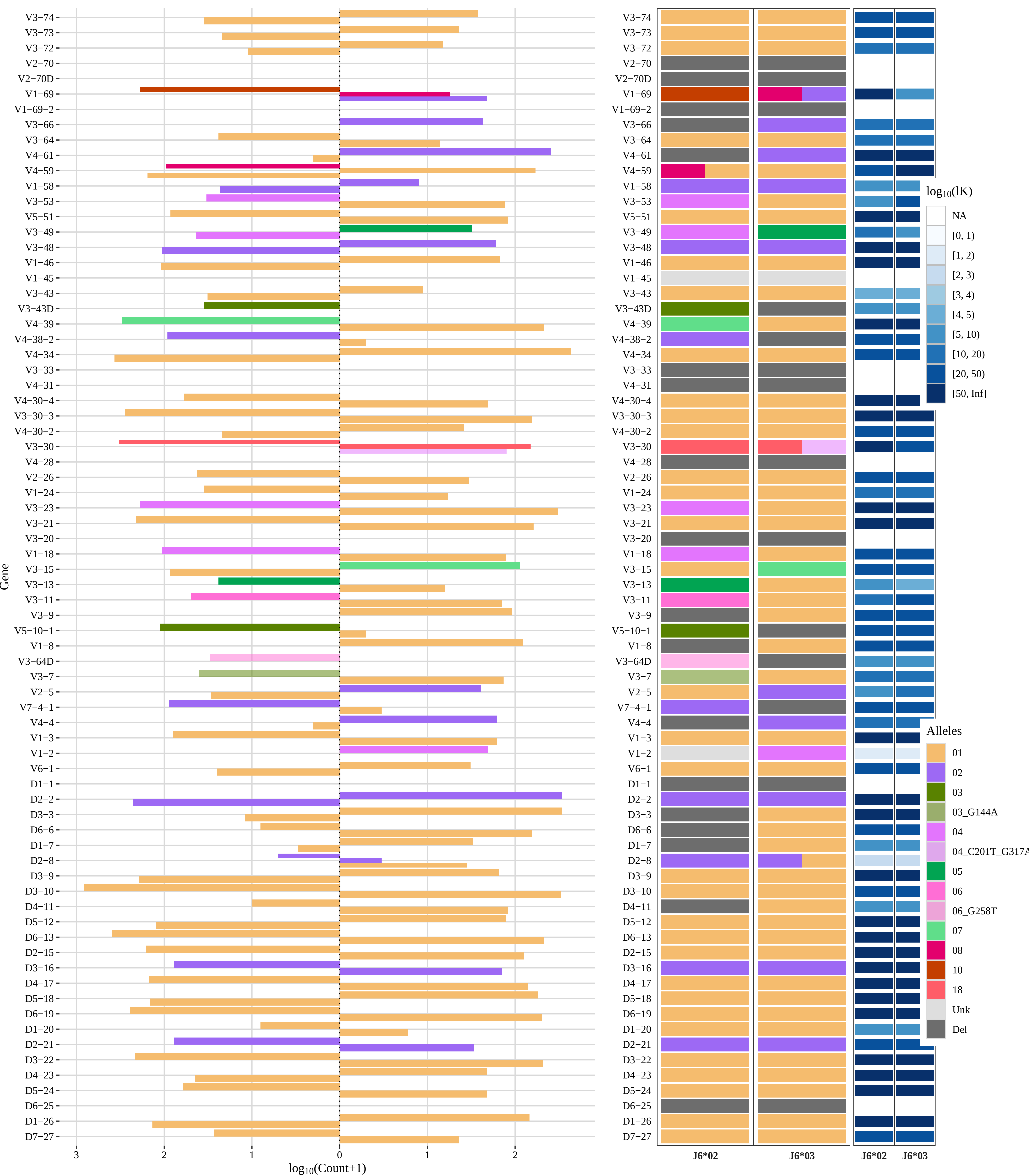

### Genotype and haplotype of VDJbase sample I73 (study P1)

(illustrations downloaded from VDJbase in July, 2020)

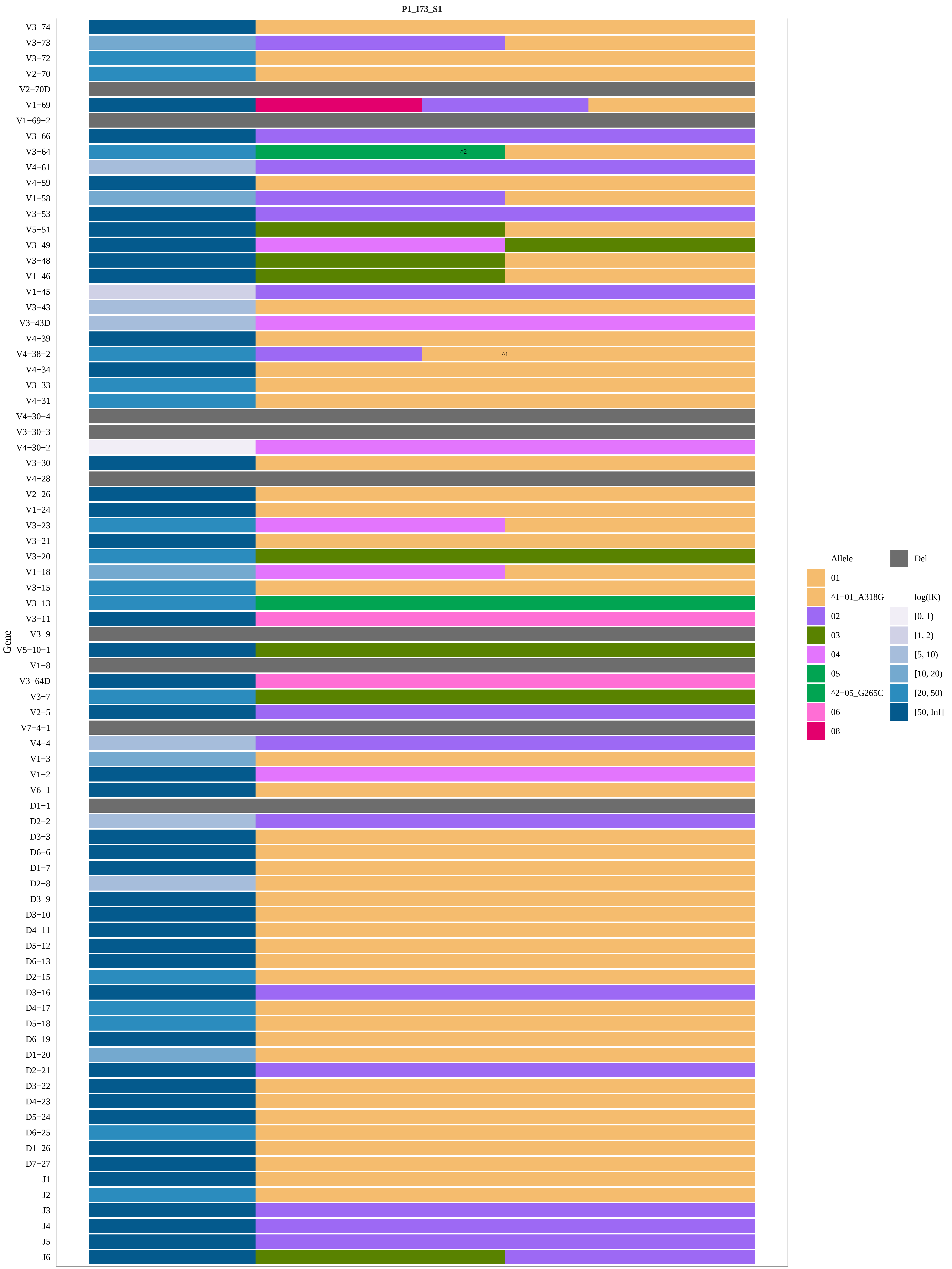

P1_I73_S1
V3−74
V3−73
V3−72
V2−70
V2−70D
V1−69
V1−69−2
V3−66
V3−64
V4−61
V4−59
V1−58
V3−53
V5−51
V3−49
V3−48
V1−46
V1−45
V3−43
V3−43D
V4−39
V4−38−2
V4−34
V3−33
V4−31
V4−30−4
V3−30−3
V4−30−2
V3−30
V4−28
V2−26
V1−24
V3−23
V3−21
V3−20
V1−18
V3−15
V3−13
V3−11
V3−9
V5−10−1
V1−8
V3−64D
V3−7
V2−5
V7−4−1
V4−4
V1−3
V1−2
V6−1
D1−1
D2−2
D3−3
D6−6
D1−7
D2−8
D3−9
D3−10
D4−11
D5−12
D6−13
D2−15
D3−16
D4−17
D5−18
D6−19
D1−20
D2−21
D3−22
D4−23
D5−24
D6−25
D1−26
D7−27
J1
J2
J3
J4
J5
J6
Gene
^2
^1
Allele
01
^1−01_A318G
02
03
04
05
^2−05_G265C
06
08
Del
log(lK)
[0, 1)
[1, 2)
[5, 10)
[10, 20)
[20, 50)
[50, Inf]

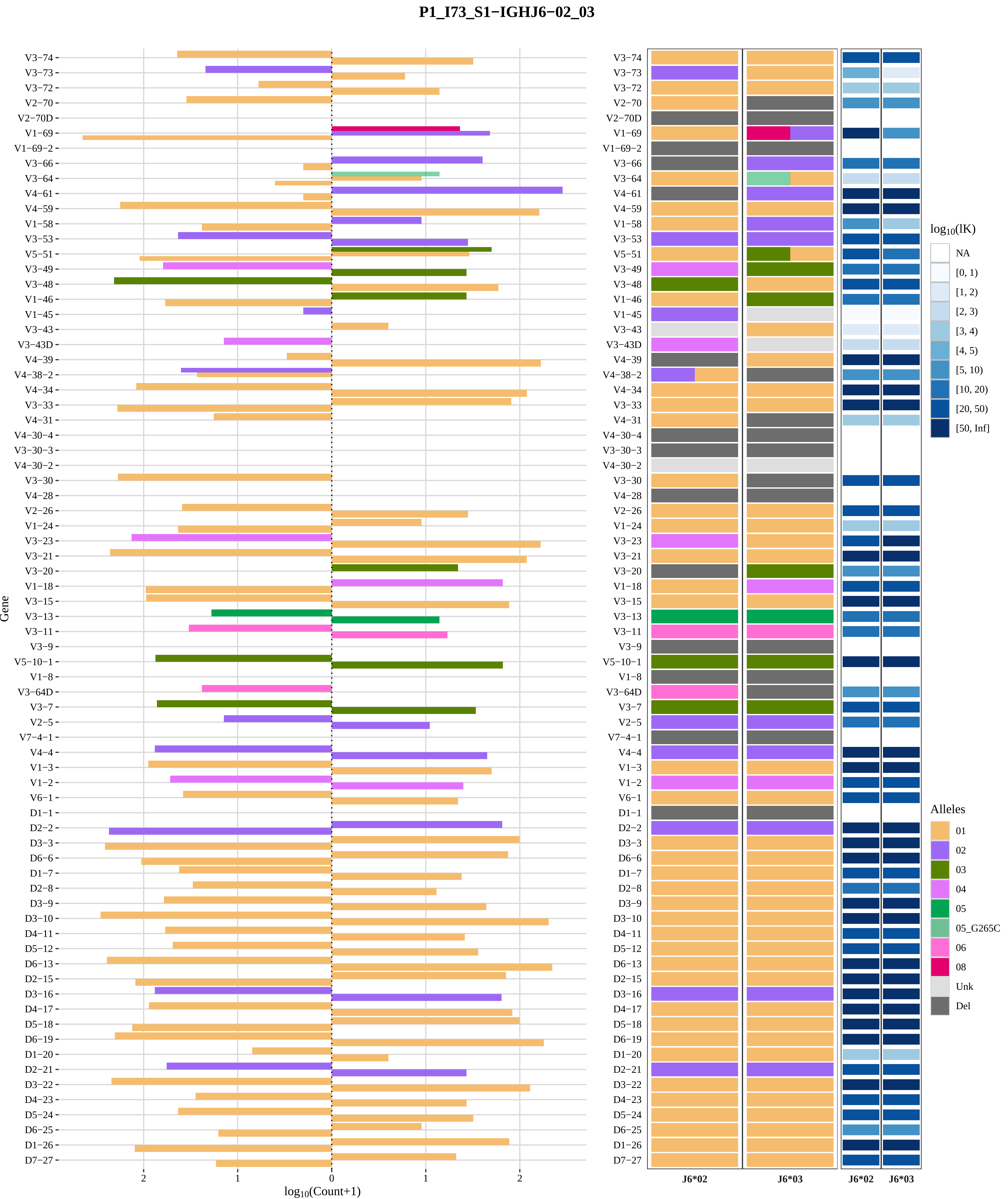

P1_I73_S1−IGHJ6−02_03
Gene
log10(Count+1)
V3−74
V3−73
V3−72
V2−70
V2−70D
V1−69
V1−69−2
V3−66
V3−64
V4−61
V4−59
V1−58
V3−53
V5−51
V3−49
V3−48
V1−46
V1−45
V3−43
V3−43D
V4−39
V4−38−2
V4−34
V3−33
V4−31
V4−30−4
V3−30−3
V4−30−2
V3−30
V4−28
V2−26
V1−24
V3−23
V3−21
V3−20
V1−18
V3−15
V3−13
V3−11
V3−9
V5−10−1
V1−8
V3−64D
V3−7
V2−5
V7−4−1
V4−4
V1−3
V1−2
V6−1
D1−1
D2−2
D3−3
D6−6
D1−7
D2−8
D3−9
D3−10
D4−11
D5−12
D6−13
D2−15
D3−16
D4−17
D5−18
D6−19
D1−20
D2−21
D3−22
D4−23
D5−24
D6−25
D1−26
D7−27
2
1
0
1
2
J6*02
J6*03
J6*02
J6*03
log10(lK)
NA
[0, 1)
[1, 2)
[2, 3)
[3, 4)
[4, 5)
[5, 10)
[10, 20)
[20, 50)
[50, Inf]
Alleles
01
02
03
04
05
05_G265C
06
08
Unk
Del

### Genotype and haplotype of VDJbase sample I76 (study P1)

(illustrations downloaded from VDJbase in July, 2020)

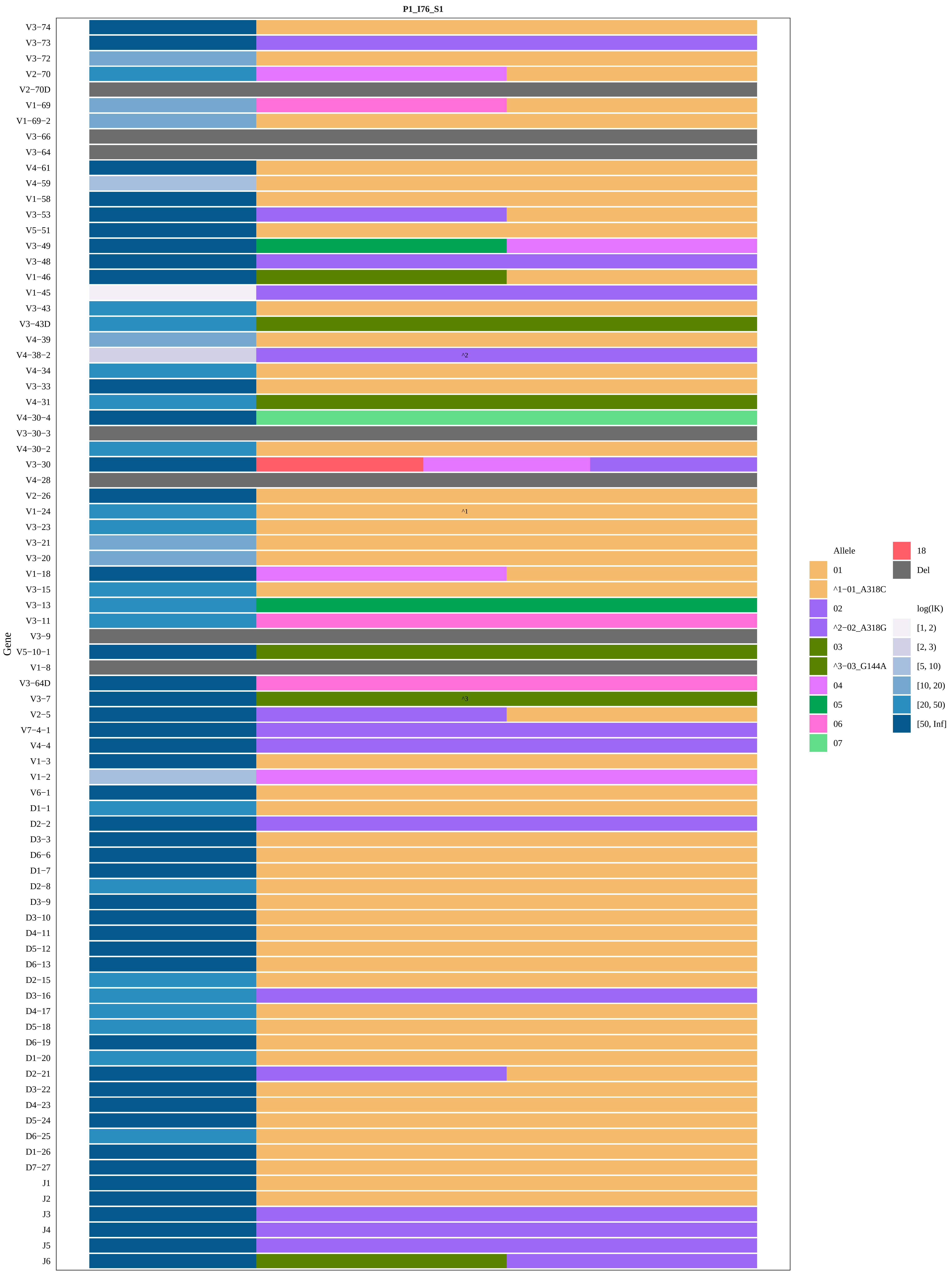
P1_I76_S1
Gene
V3−74
V3−73
V3−72
V2−70
V2−70D
V1−69
V1−69−2
V3−66
V3−64
V4−61
V4−59
V1−58
V3−53
V5−51
V3−49
V3−48
V1−46
V1−45
V3−43
V3−43D
V4−39
V4−38−2
V4−34
V3−33
V4−31
V4−30−4
V3−30−3
V4−30−2
V3−30
V4−28
V2−26
V1−24
V3−23
V3−21
V3−20
V1−18
V3−15
V3−13
V3−11
V3−9
V5−10−1
V1−8
V3−64D
V3−7
V2−5
V7−4−1
V4−4
V1−3
V1−2
V6−1
D1−1
D2−2
D3−3
D6−6
D1−7
D2−8
D3−9
D3−10
D4−11
D5−12
D6−13
D2−15
D3−16
D4−17
D5−18
D6−19
D1−20
D2−21
D3−22
D4−23
D5−24
D6−25
D1−26
D7−27
J1
J2
J3
J4
J5
J6
^2
^1
^3
Allele
01
^1−01_A318C
02
^2−02_A318G
03
^3−03_G144A
04
05
06
07
18
Del
log(lK)
[1, 2)
[2, 3)
[5, 10)
[10, 20)
[20, 50)
[50, Inf]

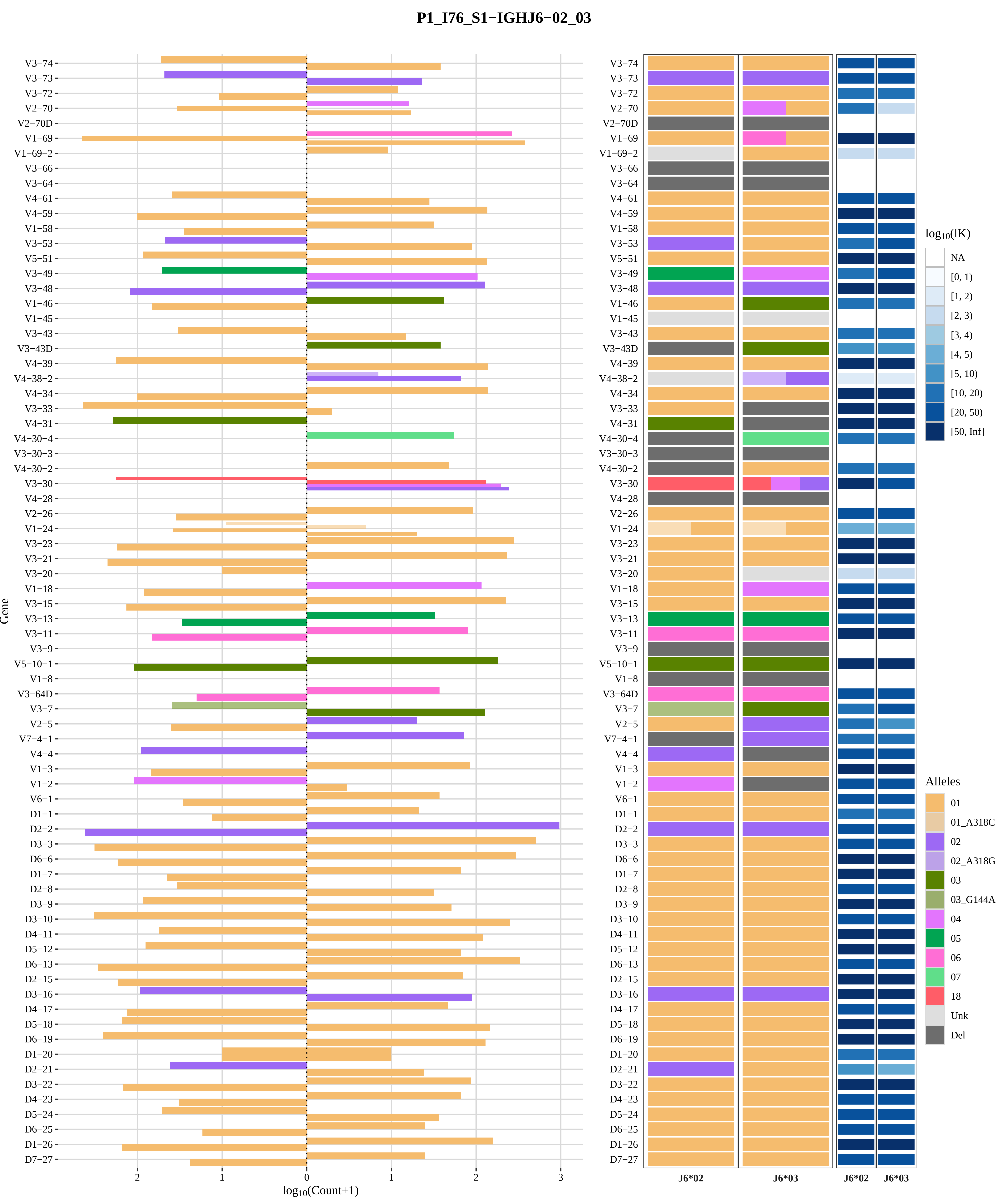
P1_I76_S1−IGHJ6−02_03
Gene
log10(Count+1)
V3−74
V3−73
V3−72
V2−70
V2−70D
V1−69
V1−69−2
V3−66
V3−64
V4−61
V4−59
V1−58
V3−53
V5−51
V3−49
V3−48
V1−46
V1−45
V3−43
V3−43D
V4−39
V4−38−2
V4−34
V3−33
V4−31
V4−30−4
V3−30−3
V4−30−2
V3−30
V4−28
V2−26
V1−24
V3−23
V3−21
V3−20
V1−18
V3−15
V3−13
V3−11
V3−9
V5−10−1
V1−8
V3−64D
V3−7
V2−5
V7−4−1
V4−4
V1−3
V1−2
V6−1
D1−1
D2−2
D3−3
D6−6
D1−7
D2−8
D3−9
D3−10
D4−11
D5−12
D6−13
D2−15
D3−16
D4−17
D5−18
D6−19
D1−20
D2−21
D3−22
D4−23
D5−24
D6−25
D1−26
D7−27
J6*02
J6*03
log10(lK)
NA
[0, 1)
[1, 2)
[2, 3)
[3, 4)
[4, 5)
[5, 10)
[10, 20)
[20, 50)
[50, Inf]
Alleles
01
01_A318C
02
02_A318G
03
03_G144A
04
05
06
07
18
Unk
Del

### Genotype and haplotype of VDJbase sample I81 (study P1)

(illustrations downloaded from VDJbase in July, 2020)

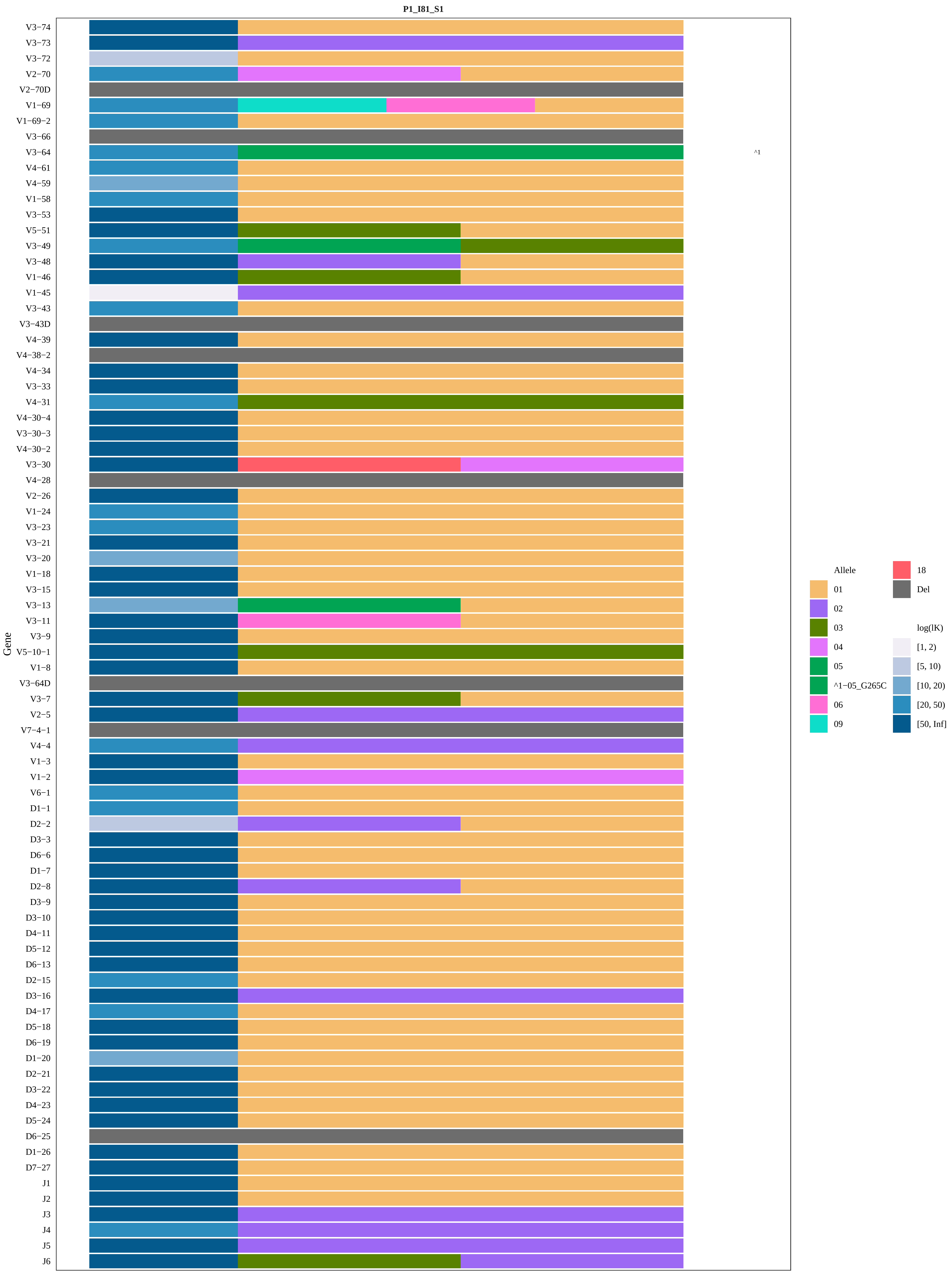

P1_I81_S1
Gene
V3−74
V3−73
V3−72
V2−70
V2−70D
V1−69
V1−69−2
V3−66
V3−64
V4−61
V4−59
V1−58
V3−53
V5−51
V3−49
V3−48
V1−46
V1−45
V3−43
V3−43D
V4−39
V4−38−2
V4−34
V3−33
V4−31
V4−30−4
V3−30−3
V4−30−2
V3−30
V4−28
V2−26
V1−24
V3−23
V3−21
V3−20
V1−18
V3−15
V3−13
V3−11
V3−9
V5−10−1
V1−8
V3−64D
V3−7
V2−5
V7−4−1
V4−4
V1−3
V1−2
V6−1
D1−1
D2−2
D3−3
D6−6
D1−7
D2−8
D3−9
D3−10
D4−11
D5−12
D6−13
D2−15
D3−16
D4−17
D5−18
D6−19
D1−20
D2−21
D3−22
D4−23
D5−24
D6−25
D1−26
D7−27
J1
J2
J3
J4
J5
J6
^1
Allele
01
02
03
04
05
^1−05_G265C
06
09
18
Del
log(lK)
[1, 2)
[5, 10)
[10, 20)
[20, 50)
[50, Inf]

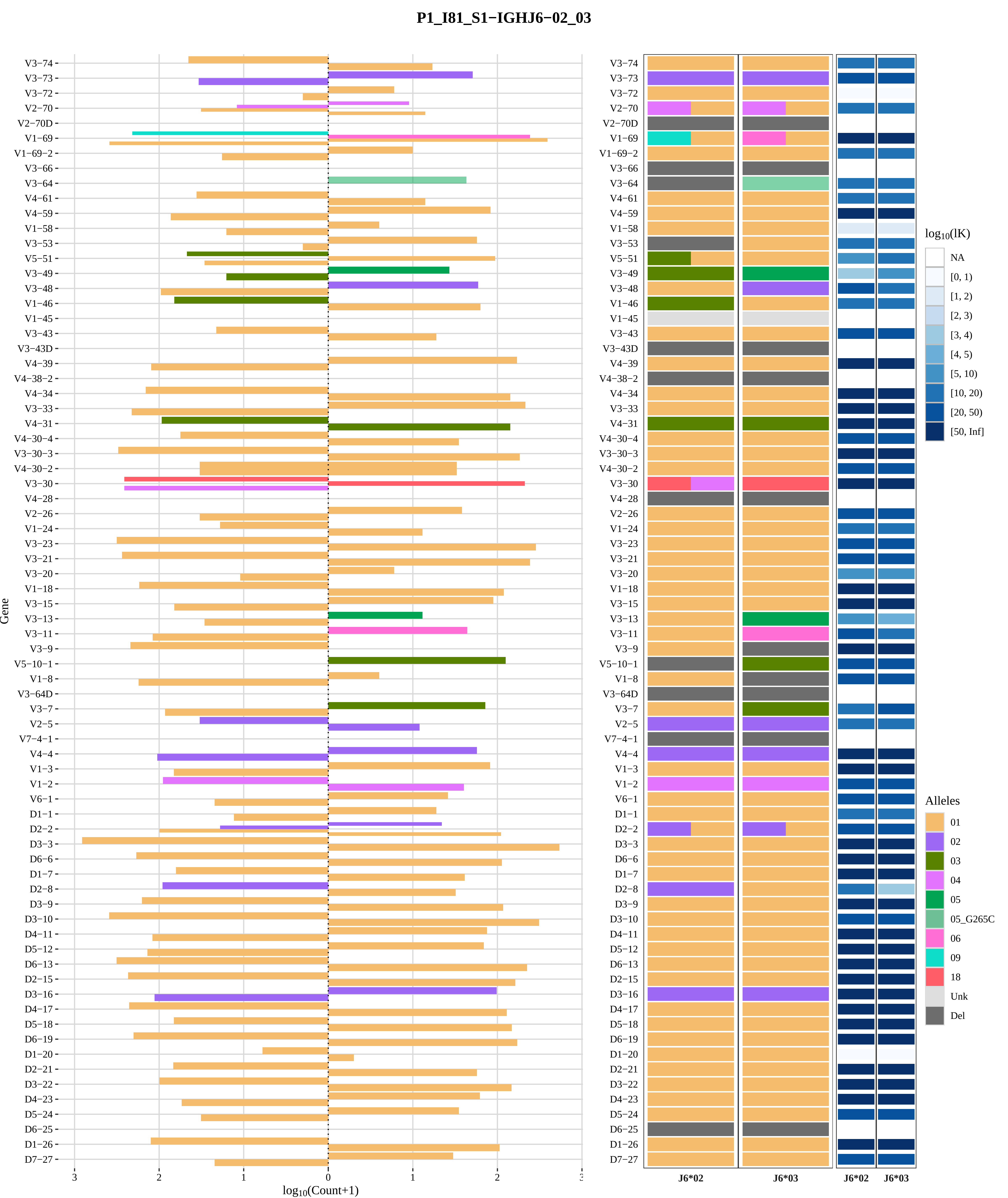
P1_I81_S1−IGHJ6−02_03
Gene
log10(Count+1)
J6*02
J6*03
log10(lK)
NA
[0, 1)
[1, 2)
[2, 3)
[3, 4)
[4, 5)
[5, 10)
[10, 20)
[20, 50)
[50, Inf]
Alleles
01
02
03
04
05
05_G265C
06
09
18
Unk
Del

### Genotype and haplotype of VDJbase sample I86 (study P1)

(illustrations downloaded from VDJbase in July, 2020)

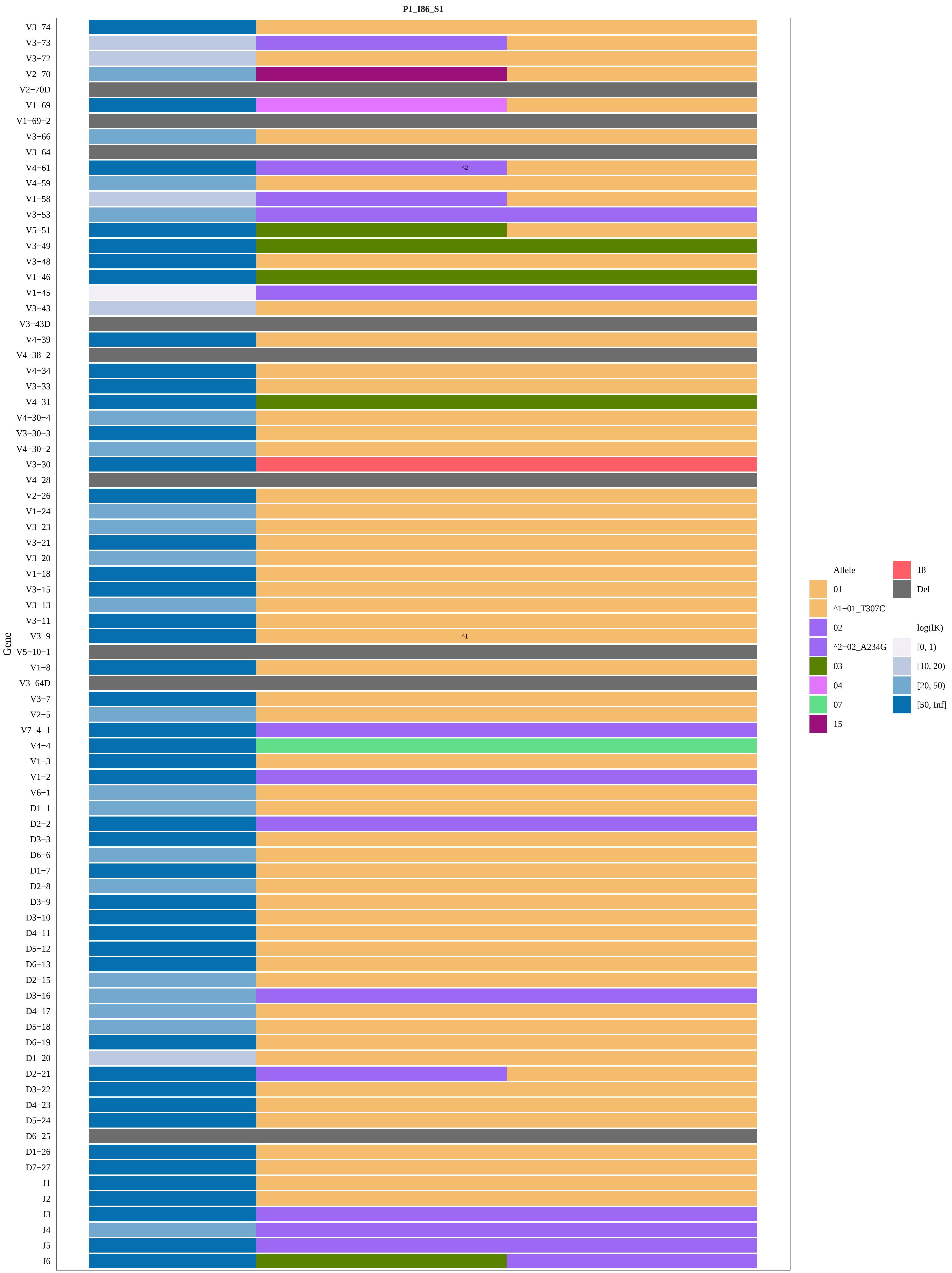

P1_I86_S1
Gene
V3−74
V3−73
V3−72
V2−70
V2−70D
V1−69
V1−69−2
V3−66
V3−64
V4−61
V4−59
V1−58
V3−53
V5−51
V3−49
V3−48
V1−46
V1−45
V3−43
V3−43D
V4−39
V4−38−2
V4−34
V3−33
V4−31
V4−30−4
V3−30−3
V4−30−2
V3−30
V4−28
V2−26
V1−24
V3−23
V3−21
V3−20
V1−18
V3−15
V3−13
V3−11
V3−9
V5−10−1
V1−8
V3−64D
V3−7
V2−5
V7−4−1
V4−4
V1−3
V1−2
V6−1
D1−1
D2−2
D3−3
D6−6
D1−7
D2−8
D3−9
D3−10
D4−11
D5−12
D6−13
D2−15
D3−16
D4−17
D5−18
D6−19
D1−20
D2−21
D3−22
D4−23
D5−24
D6−25
D1−26
D7−27
J1
J2
J3
J4
J5
J6
^2
^1
Allele
01
^1−01_T307C
02
^2−02_A234G
03
04
07
15
18
Del
log(lK)
[0, 1)
[10, 20)
[20, 50)
[50, Inf]

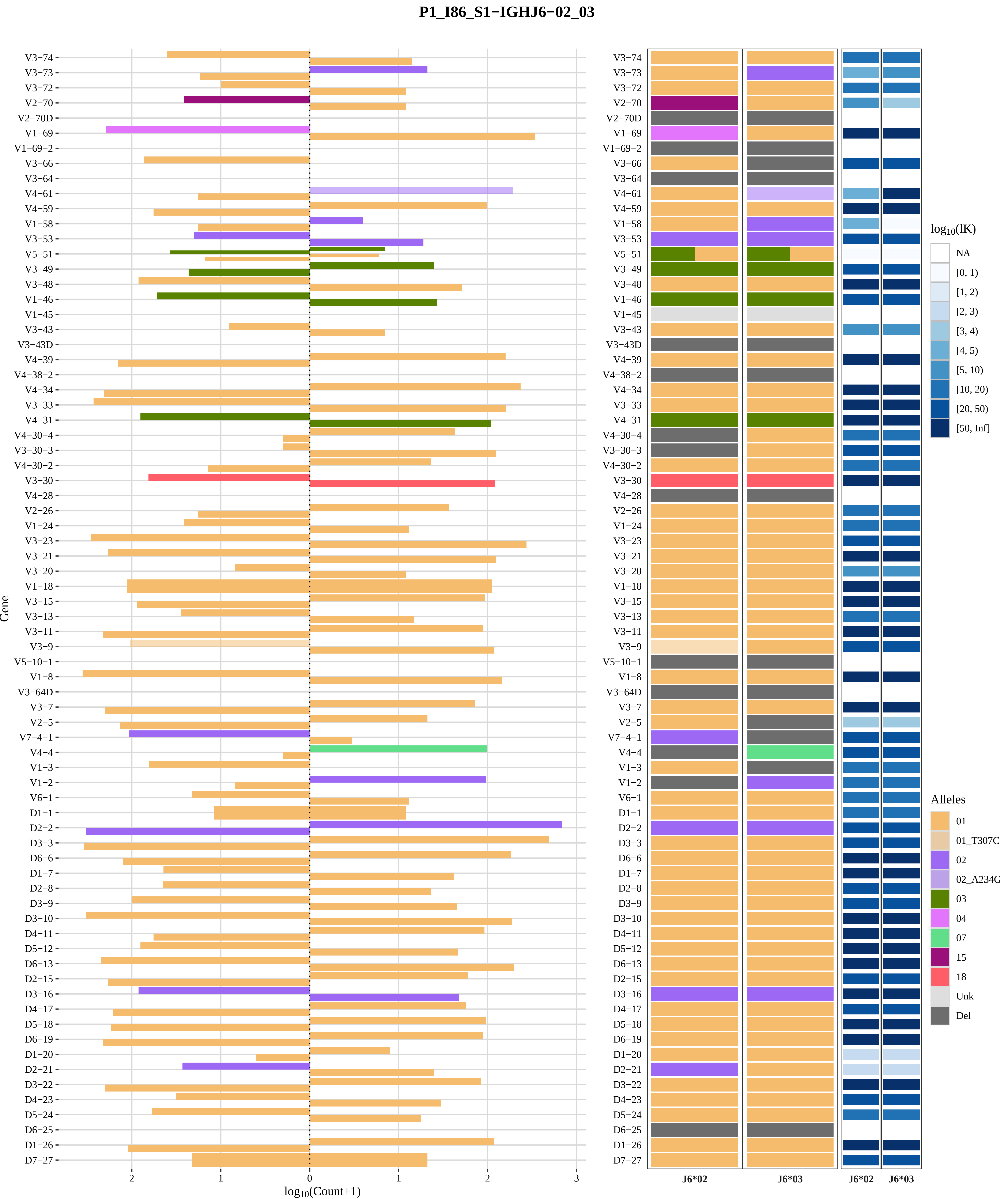

P1_I86_S1−IGHJ6−02_03
Gene
log10(Count+1)
V3−74
V3−73
V3−72
V2−70
V2−70D
V1−69
V1−69−2
V3−66
V3−64
V4−61
V4−59
V1−58
V3−53
V5−51
V3−49
V3−48
V1−46
V1−45
V3−43
V3−43D
V4−39
V4−38−2
V4−34
V3−33
V4−31
V4−30−4
V3−30−3
V4−30−2
V3−30
V4−28
V2−26
V1−24
V3−23
V3−21
V3−20
V1−18
V3−15
V3−13
V3−11
V3−9
V5−10−1
V1−8
V3−64D
V3−7
V2−5
V7−4−1
V4−4
V1−3
V1−2
V6−1
D1−1
D2−2
D3−3
D6−6
D1−7
D2−8
D3−9
D3−10
D4−11
D5−12
D6−13
D2−15
D3−16
D4−17
D5−18
D6−19
D1−20
D2−21
D3−22
D4−23
D5−24
D6−25
D1−26
D7−27
J6*02
J6*03
log10(lK)
NA
[0, 1)
[1, 2)
[2, 3)
[3, 4)
[4, 5)
[5, 10)
[10, 20)
[20, 50)
[50, Inf]
Alleles
01
01_T307C
02
02_A234G
03
04
07
15
18
Unk
Del

### Genotype and haplotype of VDJbase sample I88 (study P1)

(illustrations downloaded from VDJbase in July, 2020)

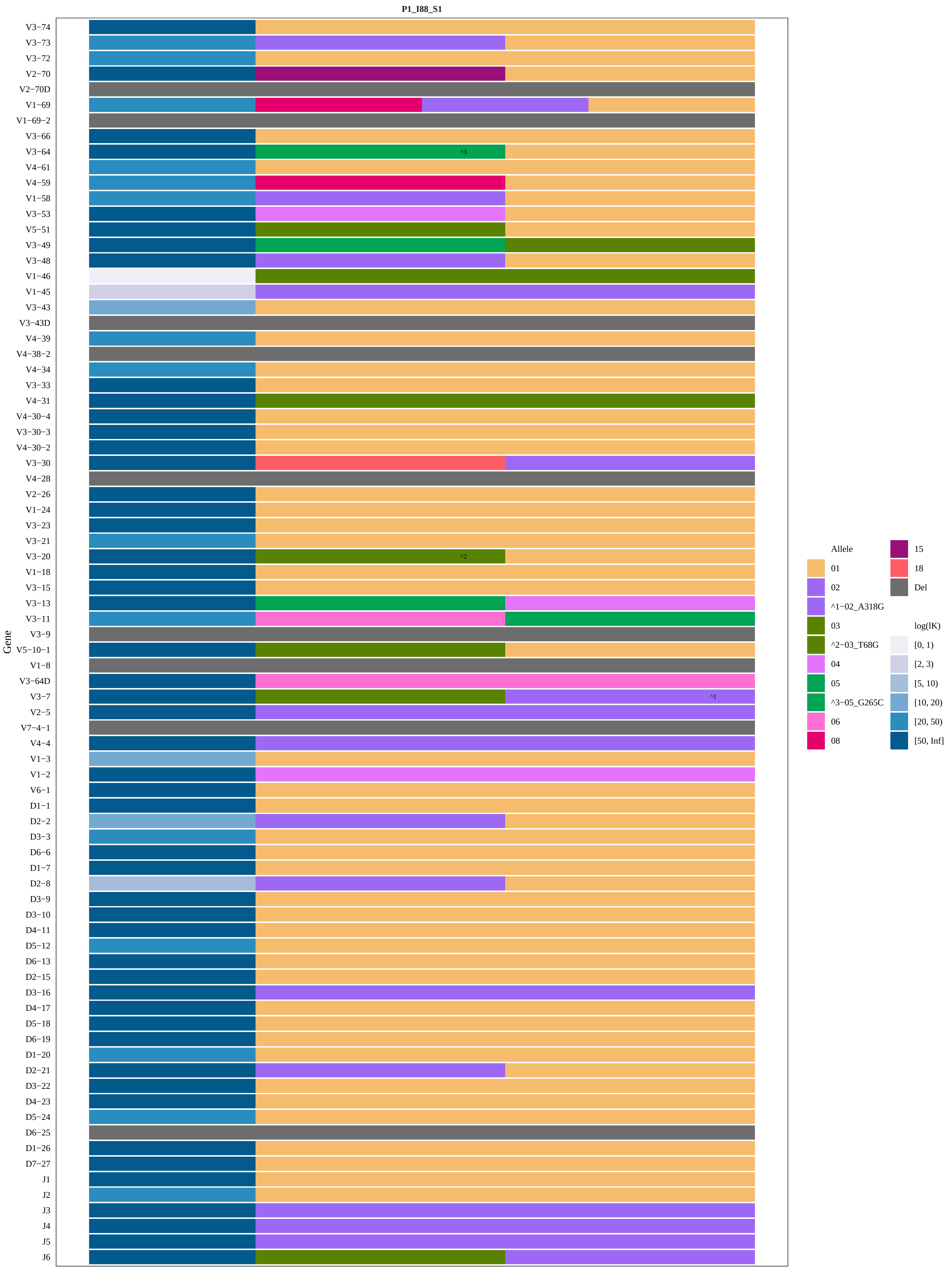

P1_I88_S1
Gene
V3−74
V3−73
V3−72
V2−70
V2−70D
V1−69
V1−69−2
V3−66
V3−64
V4−61
V4−59
V1−58
V3−53
V5−51
V3−49
V3−48
V1−46
V1−45
V3−43
V3−43D
V4−39
V4−38−2
V4−34
V3−33
V4−31
V4−30−4
V3−30−3
V4−30−2
V3−30
V4−28
V2−26
V1−24
V3−23
V3−21
V3−20
V1−18
V3−15
V3−13
V3−11
V3−9
V5−10−1
V1−8
V3−64D
V3−7
V2−5
V7−4−1
V4−4
V1−3
V1−2
V6−1
D1−1
D2−2
D3−3
D6−6
D1−7
D2−8
D3−9
D3−10
D4−11
D5−12
D6−13
D2−15
D3−16
D4−17
D5−18
D6−19
D1−20
D2−21
D3−22
D4−23
D5−24
D6−25
D1−26
D7−27
J1
J2
J3
J4
J5
J6
^3
^2
^1
Allele
01
02
^1−02_A318G
03
^2−03_T68G
04
05
^3−05_G265C
06
08
15
18
Del
log(lK)
[0, 1)
[2, 3)
[5, 10)
[10, 20)
[20, 50)
[50, Inf]

#### P1_I88_S1−IGHJ6−02_03

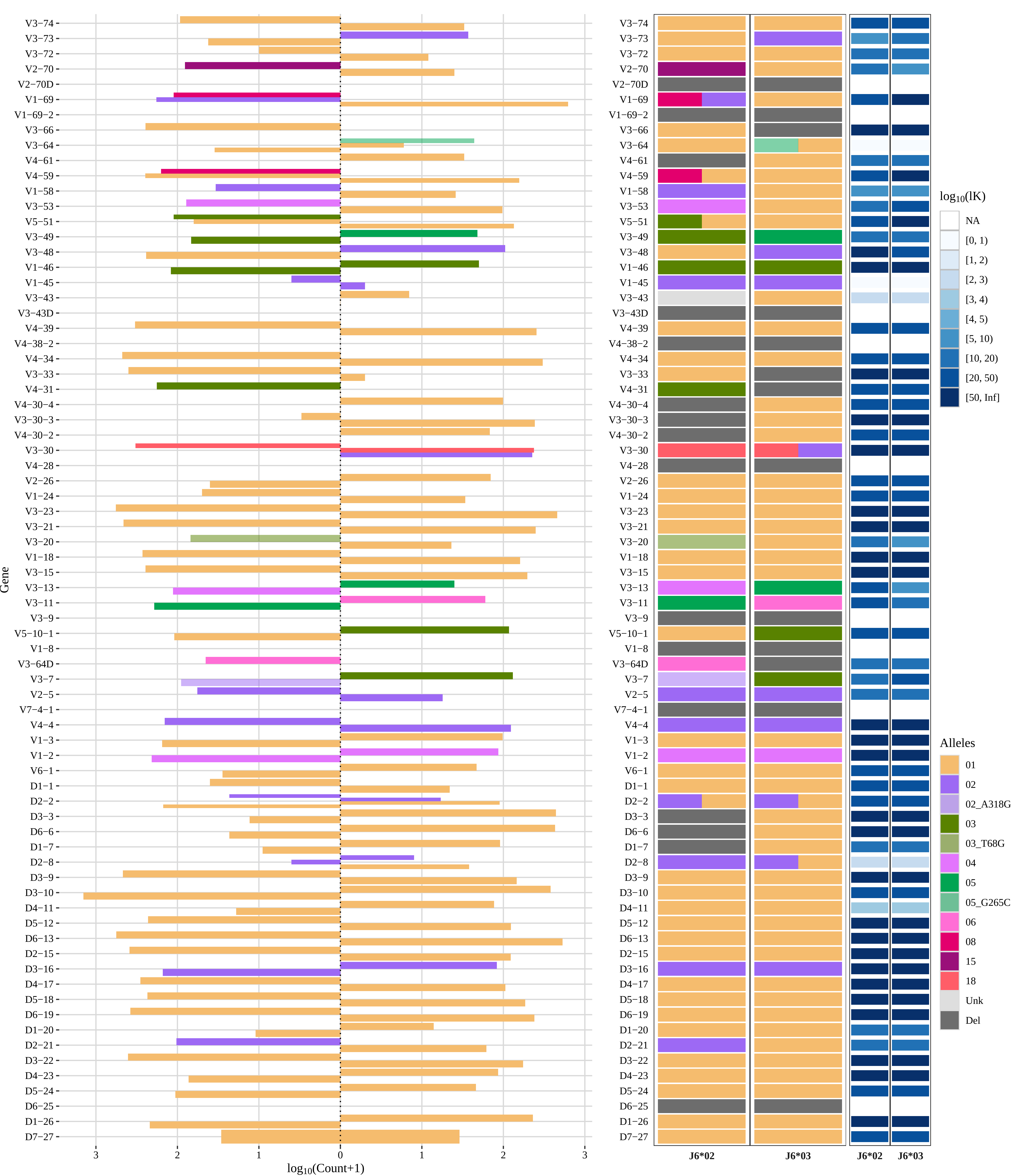

### Genotype and haplotype of VDJbase sample I90 (study P1)

(illustrations downloaded from VDJbase in July, 2020)

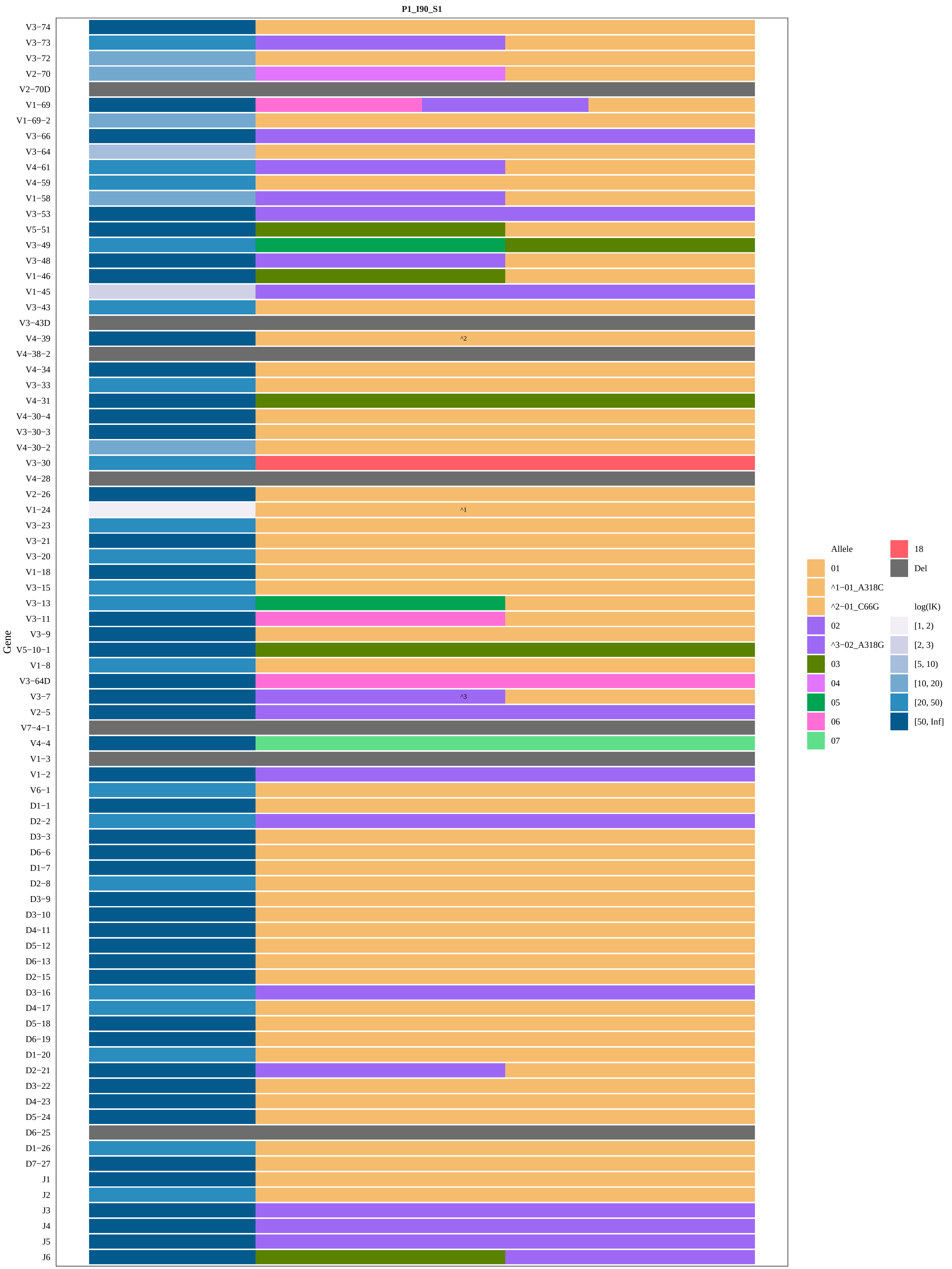
P1_I90_S1
Gene
V3−74
V3−73
V3−72
V2−70
V2−70D
V1−69
V1−69−2
V3−66
V3−64
V4−61
V4−59
V1−58
V3−53
V5−51
V3−49
V3−48
V1−46
V1−45
V3−43
V3−43D
V4−39
V4−38−2
V4−34
V3−33
V4−31
V4−30−4
V3−30−3
V4−30−2
V3−30
V4−28
V2−26
V1−24
V3−23
V3−21
V3−20
V1−18
V3−15
V3−13
V3−11
V3−9
V5−10−1
V1−8
V3−64D
V3−7
V2−5
V7−4−1
V4−4
V1−3
V1−2
V6−1
D1−1
D2−2
D3−3
D6−6
D1−7
D2−8
D3−9
D3−10
D4−11
D5−12
D6−13
D2−15
D3−16
D4−17
D5−18
D6−19
D1−20
D2−21
D3−22
D4−23
D5−24
D6−25
D1−26
D7−27
J1
J2
J3
J4
J5
J6
^2
^1
^3
Allele
01
^1−01_A318C
^2−01_C66G
02
^3−02_A318G
03
04
05
06
07
18
Del
log(lK)
[1, 2)
[2, 3)
[5, 10)
[10, 20)
[20, 50)
[50, Inf]

#### P1_I90_S1−IGHJ6−02_03

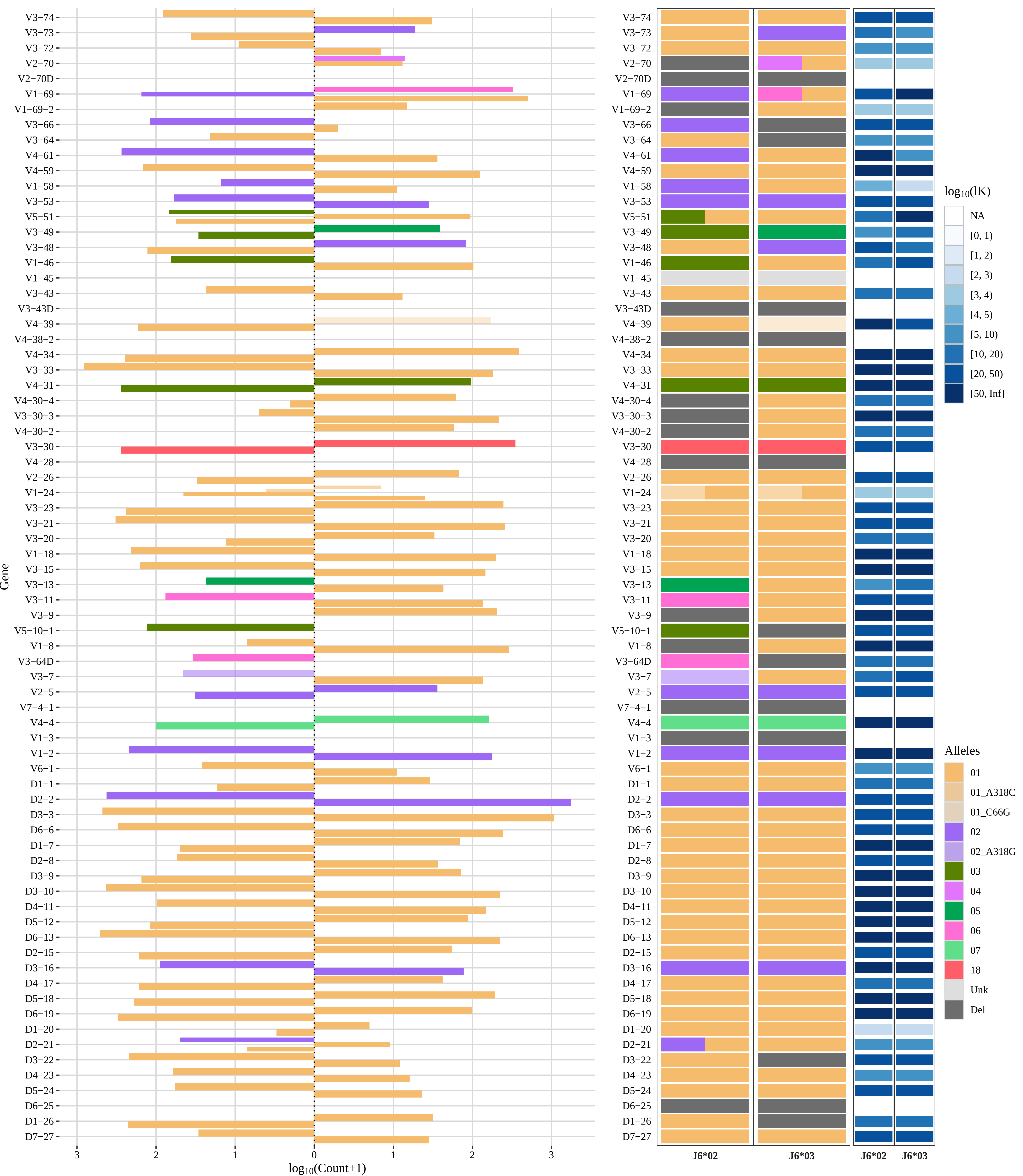

### Genotype and haplotype of VDJbase sample I91 (study P1)

(illustrations downloaded from VDJbase in July, 2020)

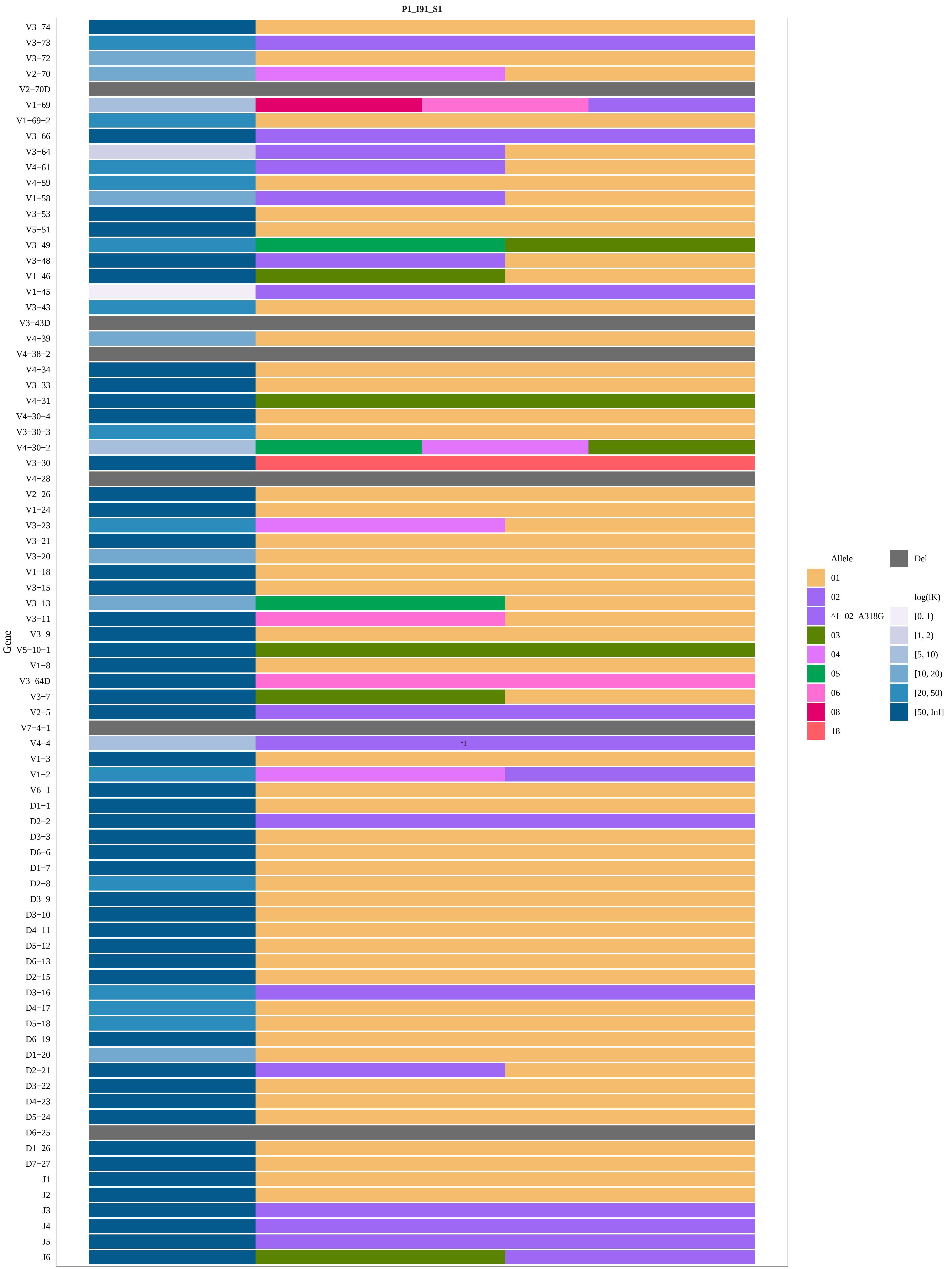

P1_I91_S1
Gene
V3−74
V3−73
V3−72
V2−70
V2−70D
V1−69
V1−69−2
V3−66
V3−64
V4−61
V4−59
V1−58
V3−53
V5−51
V3−49
V3−48
V1−46
V1−45
V3−43
V3−43D
V4−39
V4−38−2
V4−34
V3−33
V4−31
V4−30−4
V3−30−3
V4−30−2
V3−30
V4−28
V2−26
V1−24
V3−23
V3−21
V3−20
V1−18
V3−15
V3−13
V3−11
V3−9
V5−10−1
V1−8
V3−64D
V3−7
V2−5
V7−4−1
V4−4
^1
V1−3
V1−2
V6−1
D1−1
D2−2
D3−3
D6−6
D1−7
D2−8
D3−9
D3−10
D4−11
D5−12
D6−13
D2−15
D3−16
D4−17
D5−18
D6−19
D1−20
D2−21
D3−22
D4−23
D5−24
D6−25
D1−26
D7−27
J1
J2
J3
J4
J5
J6
Allele
01
02
^1−02_A318G
03
04
05
06
08
18
Del
log(lK)
[0, 1)
[1, 2)
[5, 10)
[10, 20)
[20, 50)
[50, Inf]

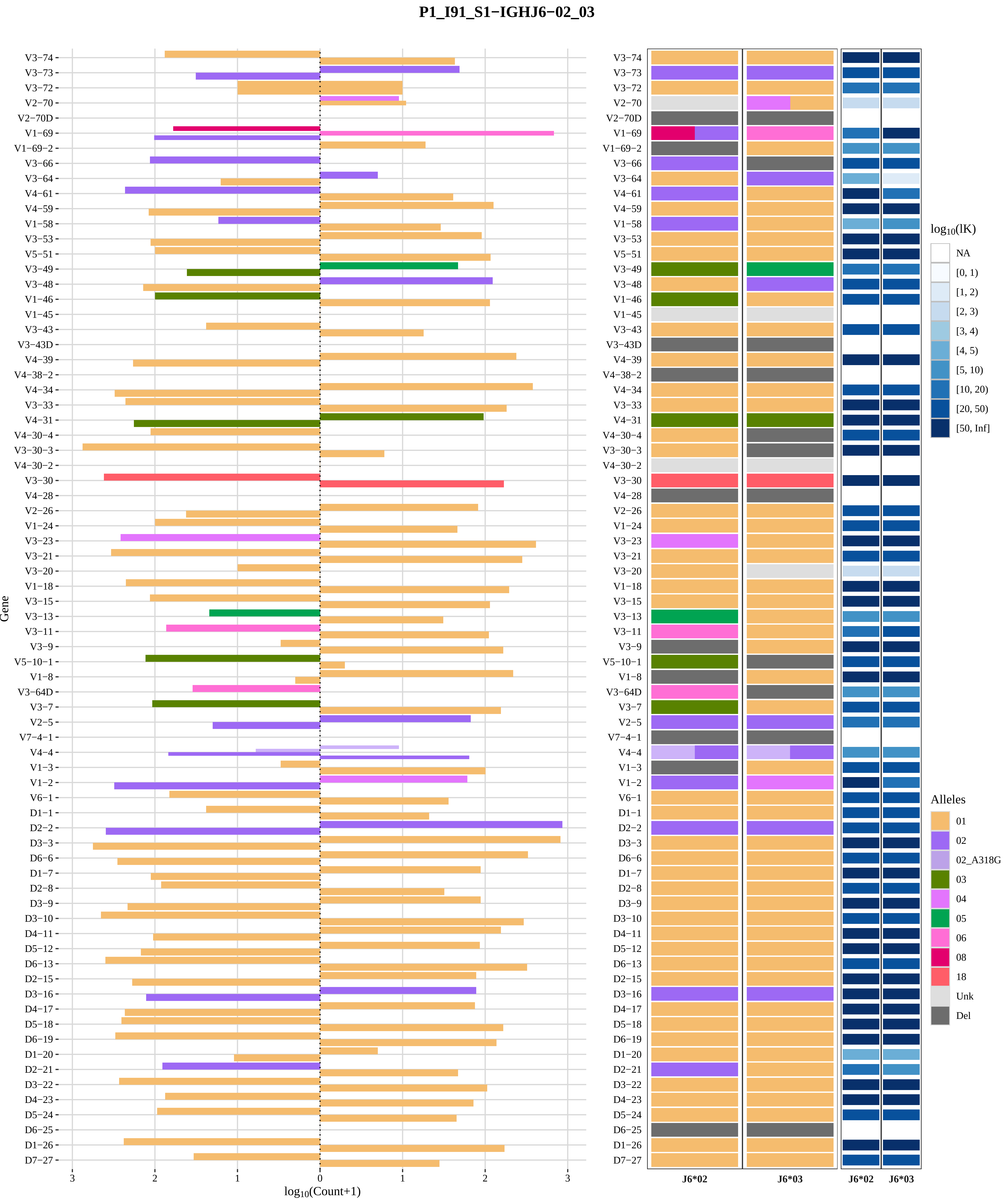

P1_I91_S1−IGHJ6−02_03
Gene
log10(Count+1)
V3−74
V3−73
V3−72
V2−70
V2−70D
V1−69
V1−69−2
V3−66
V3−64
V4−61
V4−59
V1−58
V3−53
V5−51
V3−49
V3−48
V1−46
V1−45
V3−43
V3−43D
V4−39
V4−38−2
V4−34
V3−33
V4−31
V4−30−4
V3−30−3
V4−30−2
V3−30
V4−28
V2−26
V1−24
V3−23
V3−21
V3−20
V1−18
V3−15
V3−13
V3−11
V3−9
V5−10−1
V1−8
V3−64D
V3−7
V2−5
V7−4−1
V4−4
V1−3
V1−2
V6−1
D1−1
D2−2
D3−3
D6−6
D1−7
D2−8
D3−9
D3−10
D4−11
D5−12
D6−13
D2−15
D3−16
D4−17
D5−18
D6−19
D1−20
D2−21
D3−22
D4−23
D5−24
D6−25
D1−26
D7−27
3
2
1
0
1
2
3
J6*02
J6*03
J6*02
J6*03
log10(lK)
NA
[0, 1)
[1, 2)
[2, 3)
[3, 4)
[4, 5)
[5, 10)
[10, 20)
[20, 50)
[50, Inf]
Alleles
01
02
02_A318G
03
04
05
06
08
18
Unk
Del

### Genotype and haplotype of VDJbase sample I92 (study P1)

(illustrations downloaded from VDJbase in July, 2020)

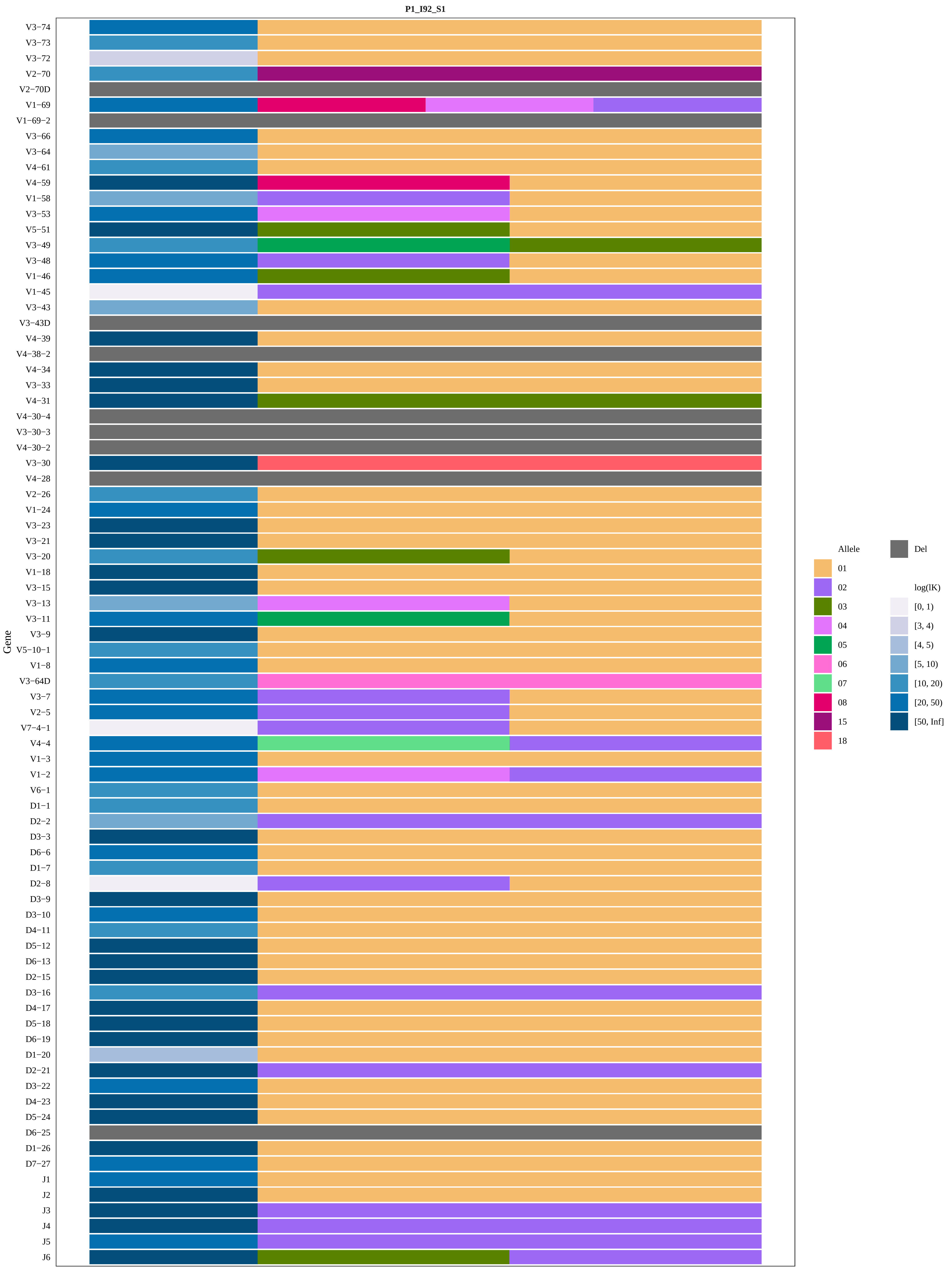
P1_I92_S1
Gene
V3−74
V3−73
V3−72
V2−70
V2−70D
V1−69
V1−69−2
V3−66
V3−64
V4−61
V4−59
V1−58
V3−53
V5−51
V3−49
V3−48
V1−46
V1−45
V3−43
V3−43D
V4−39
V4−38−2
V4−34
V3−33
V4−31
V4−30−4
V3−30−3
V4−30−2
V3−30
V4−28
V2−26
V1−24
V3−23
V3−21
V3−20
V1−18
V3−15
V3−13
V3−11
V3−9
V5−10−1
V1−8
V3−64D
V3−7
V2−5
V7−4−1
V4−4
V1−3
V1−2
V6−1
D1−1
D2−2
D3−3
D6−6
D1−7
D2−8
D3−9
D3−10
D4−11
D5−12
D6−13
D2−15
D3−16
D4−17
D5−18
D6−19
D1−20
D2−21
D3−22
D4−23
D5−24
D6−25
D1−26
D7−27
J1
J2
J3
J4
J5
J6
Allele
01
02
03
04
05
06
07
08
15
18
Del
log(lK)
[0, 1)
[3, 4)
[4, 5)
[5, 10)
[10, 20)
[20, 50)
[50, Inf]

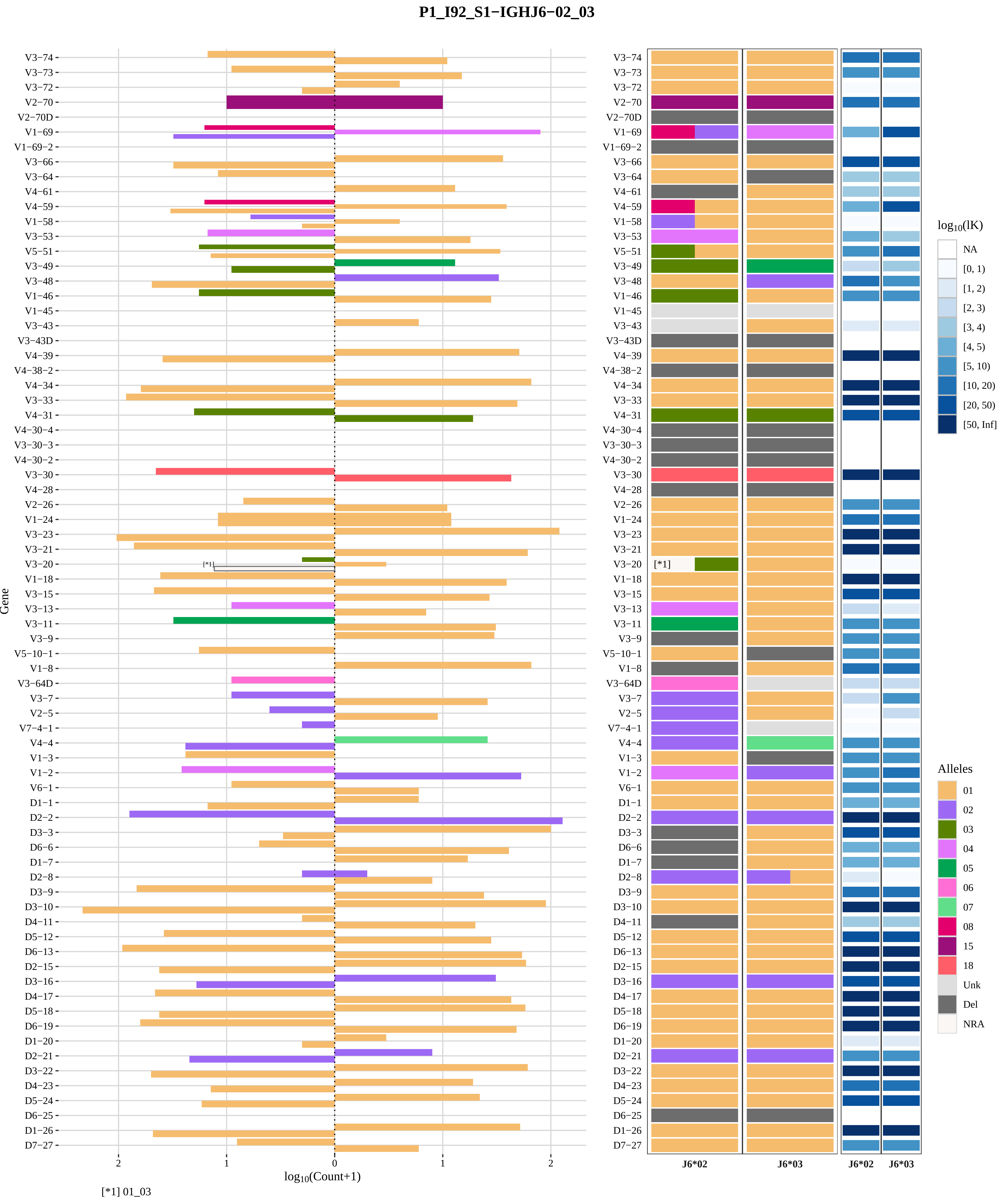
P1_I92_S1−IGHJ6−02_03
Gene
log10(Count+1)
[*1] 01_03
J6*02
J6*03
J6*02
J6*03
log10(lK)
NA
[0, 1)
[1, 2)
[2, 3)
[3, 4)
[4, 5)
[5, 10)
[10, 20)
[20, 50)
[50, Inf]
Alleles
01
02
03
04
05
06
07
08
15
18
Unk
Del
NRA

### Genotype and haplotype of VDJbase sample I93 (study P1)

(illustrations downloaded from VDJbase in July, 2020)

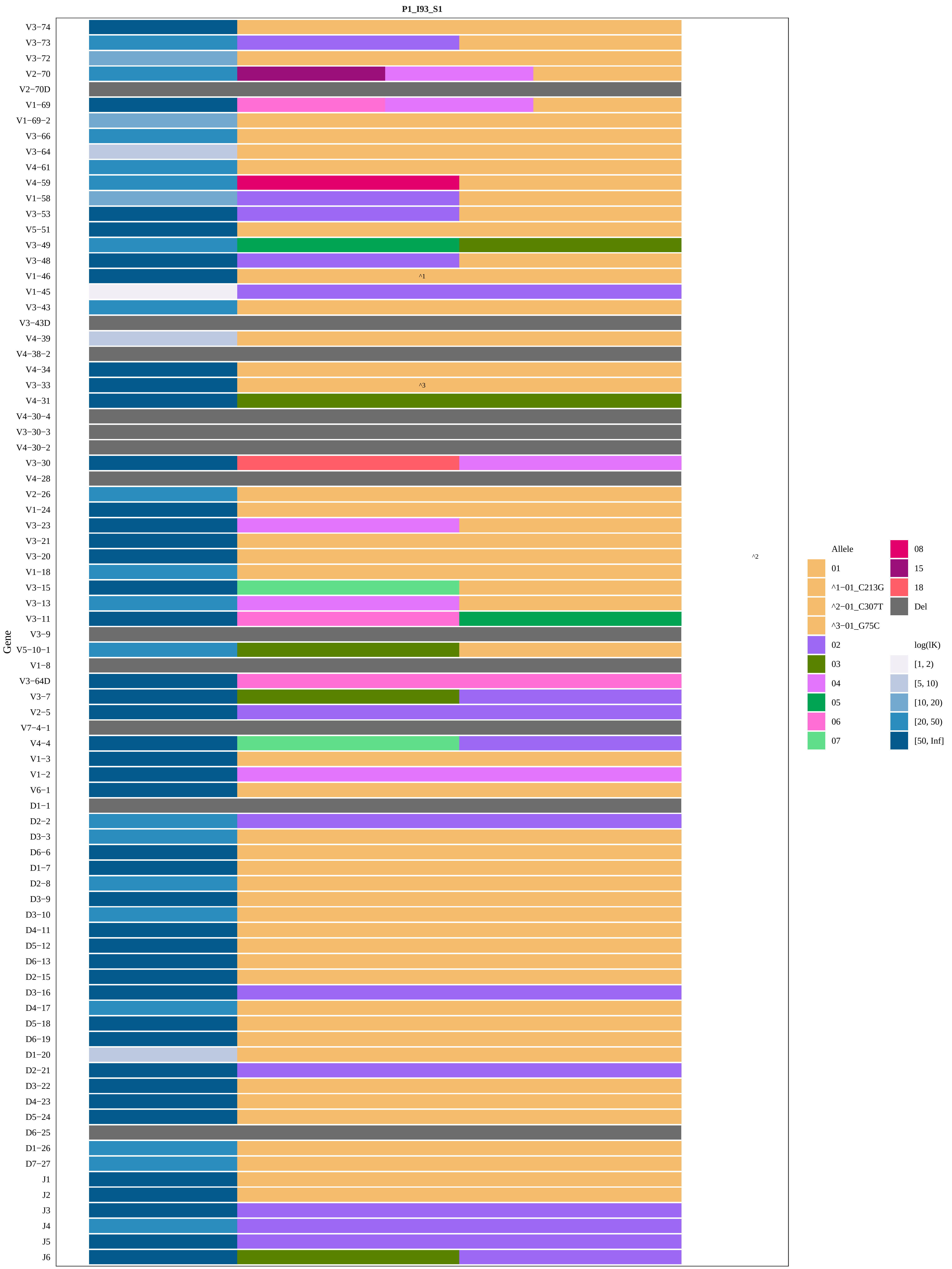
P1_I93_S1
Gene
V3−74
V3−73
V3−72
V2−70
V2−70D
V1−69
V1−69−2
V3−66
V3−64
V4−61
V4−59
V1−58
V3−53
V5−51
V3−49
V3−48
V1−46
V1−45
V3−43
V3−43D
V4−39
V4−38−2
V4−34
V3−33
V4−31
V4−30−4
V3−30−3
V4−30−2
V3−30
V4−28
V2−26
V1−24
V3−23
V3−21
V3−20
V1−18
V3−15
V3−13
V3−11
V3−9
V5−10−1
V1−8
V3−64D
V3−7
V2−5
V7−4−1
V4−4
V1−3
V1−2
V6−1
D1−1
D2−2
D3−3
D6−6
D1−7
D2−8
D3−9
D3−10
D4−11
D5−12
D6−13
D2−15
D3−16
D4−17
D5−18
D6−19
D1−20
D2−21
D3−22
D4−23
D5−24
D6−25
D1−26
D7−27
J1
J2
J3
J4
J5
J6
^1
^3
^2
Allele
01
^1−01_C213G
^2−01_C307T
^3−01_G75C
02
03
04
05
06
07
08
15
18
Del
log(lK)
[1, 2)
[5, 10)
[10, 20)
[20, 50)
[50, Inf]

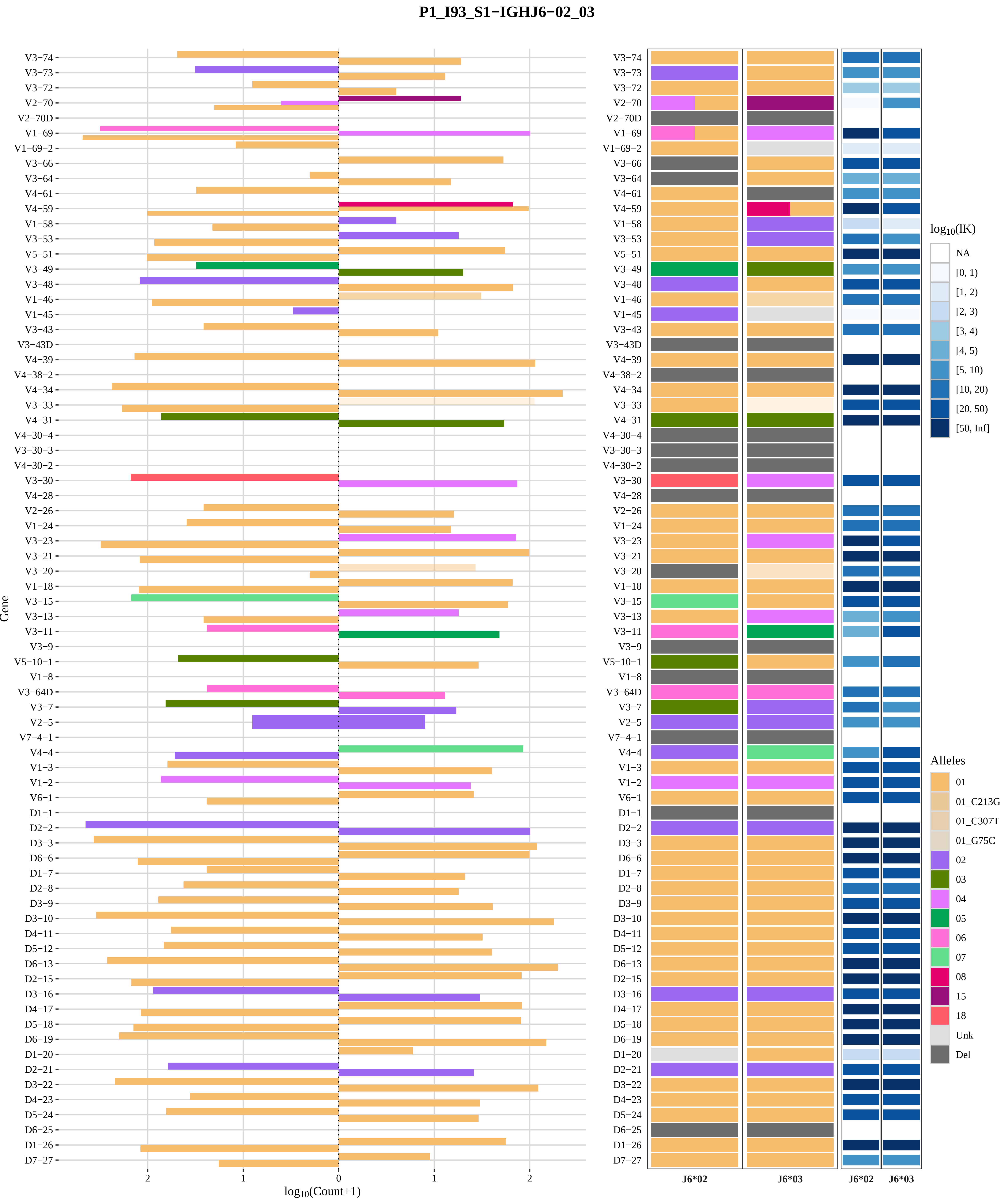
P1_I93_S1−IGHJ6−02_03
Gene
log10(Count+1)
J6*02
J6*03
log10(lK)
NA
[0, 1)
[1, 2)
[2, 3)
[3, 4)
[4, 5)
[5, 10)
[10, 20)
[20, 50)
[50, Inf]
Alleles
01
01_C213G
01_C307T
01_G75C
02
03
04
05
06
07
08
15
18
Unk
Del

### Genotype and haplotype of VDJbase sample I98 (study P1)

(illustrations downloaded from VDJbase in July, 2020)

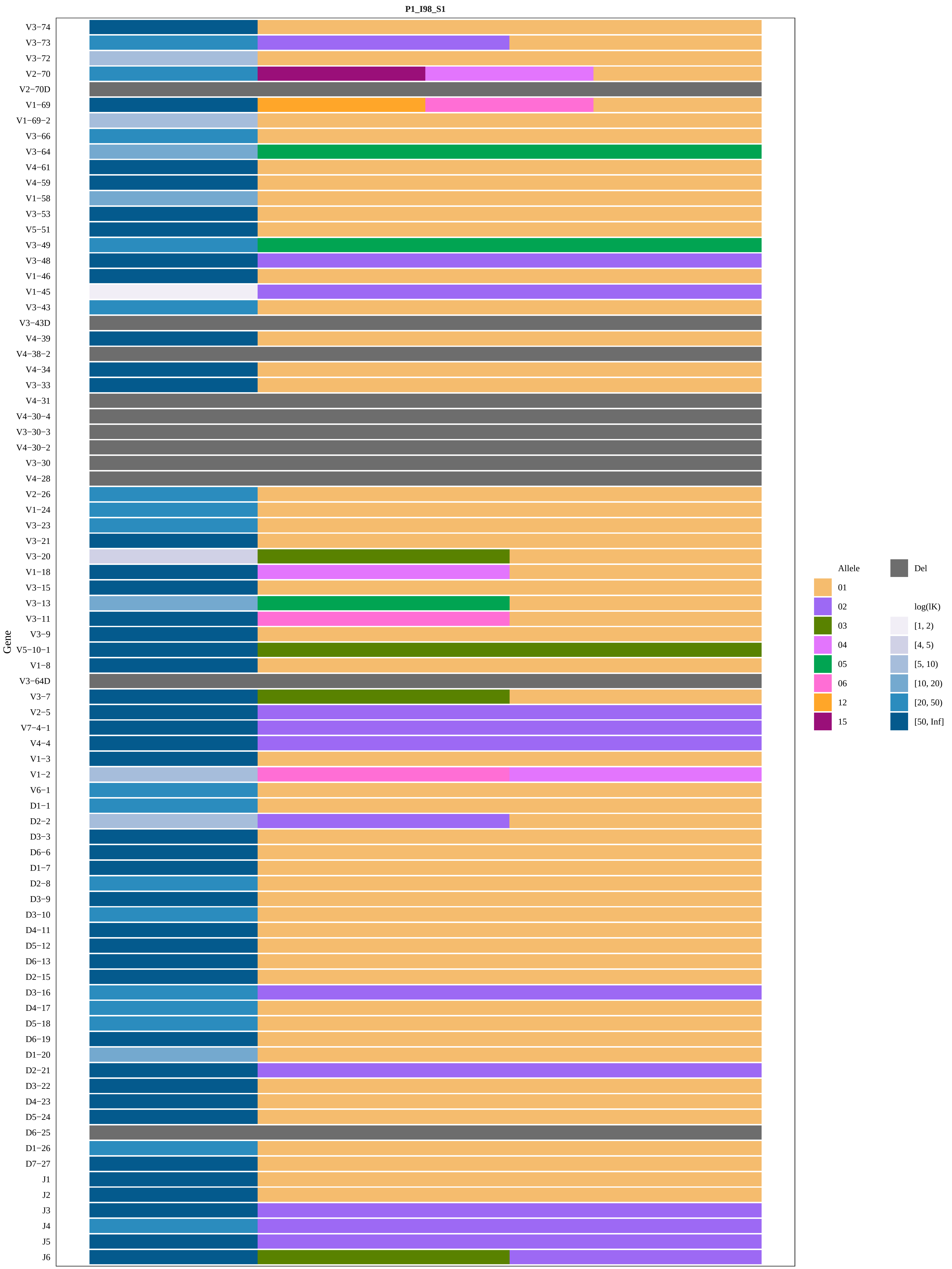
P1_I98_S1
Gene
V3−74
V3−73
V3−72
V2−70
V2−70D
V1−69
V1−69−2
V3−66
V3−64
V4−61
V4−59
V1−58
V3−53
V5−51
V3−49
V3−48
V1−46
V1−45
V3−43
V3−43D
V4−39
V4−38−2
V4−34
V3−33
V4−31
V4−30−4
V3−30−3
V4−30−2
V3−30
V4−28
V2−26
V1−24
V3−23
V3−21
V3−20
V1−18
V3−15
V3−13
V3−11
V3−9
V5−10−1
V1−8
V3−64D
V3−7
V2−5
V7−4−1
V4−4
V1−3
V1−2
V6−1
D1−1
D2−2
D3−3
D6−6
D1−7
D2−8
D3−9
D3−10
D4−11
D5−12
D6−13
D2−15
D3−16
D4−17
D5−18
D6−19
D1−20
D2−21
D3−22
D4−23
D5−24
D6−25
D1−26
D7−27
J1
J2
J3
J4
J5
J6
Allele
01
02
03
04
05
06
12
15
Del
log(lK)
[1, 2)
[4, 5)
[5, 10)
[10, 20)
[20, 50)
[50, Inf]

### P1_I98_S1−IGHJ6−02_03

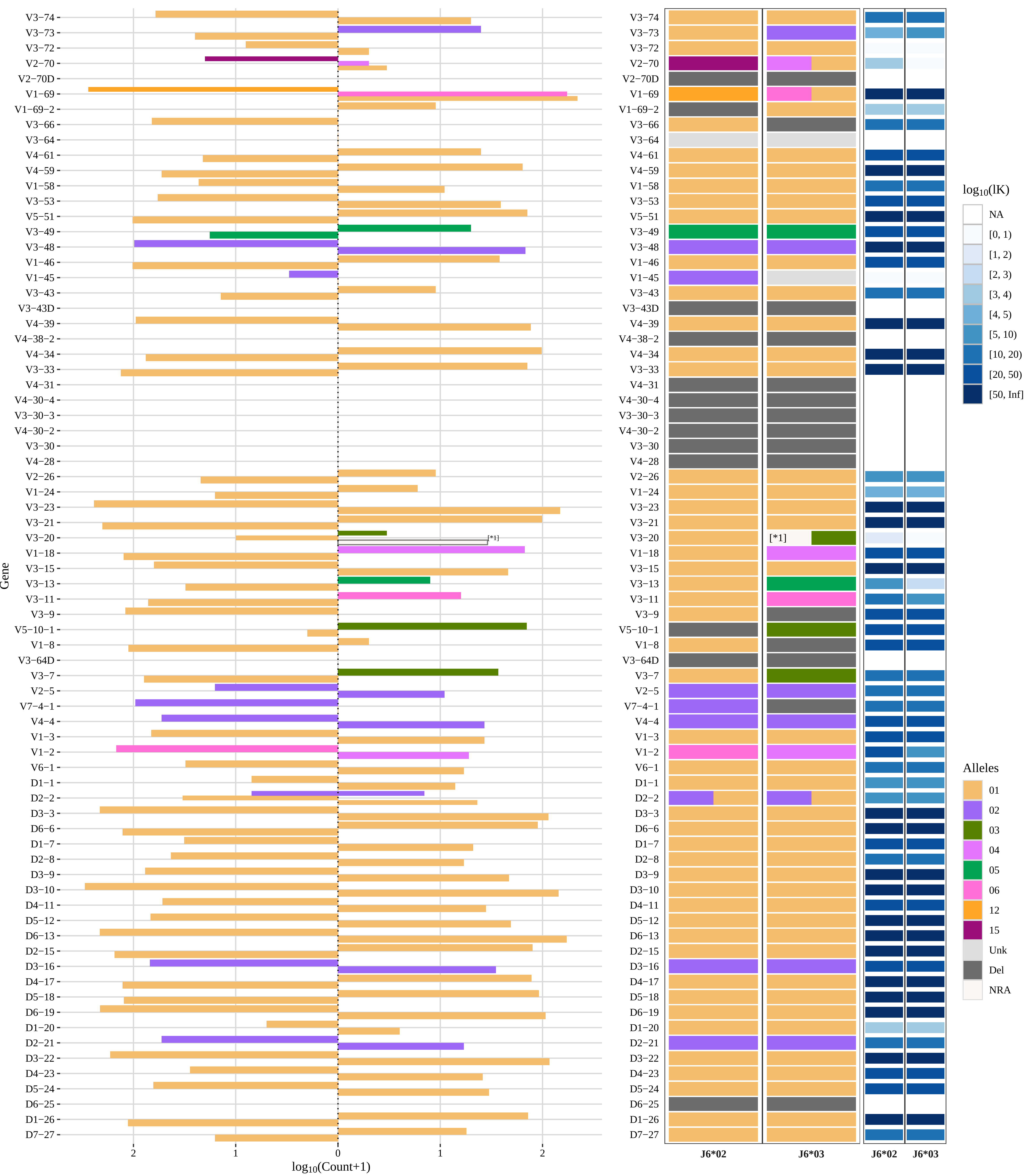


[*1] 01_03

### Genotype and haplotype of VDJbase sample I100 (study P1)

(illustrations downloaded from VDJbase in July, 2020)

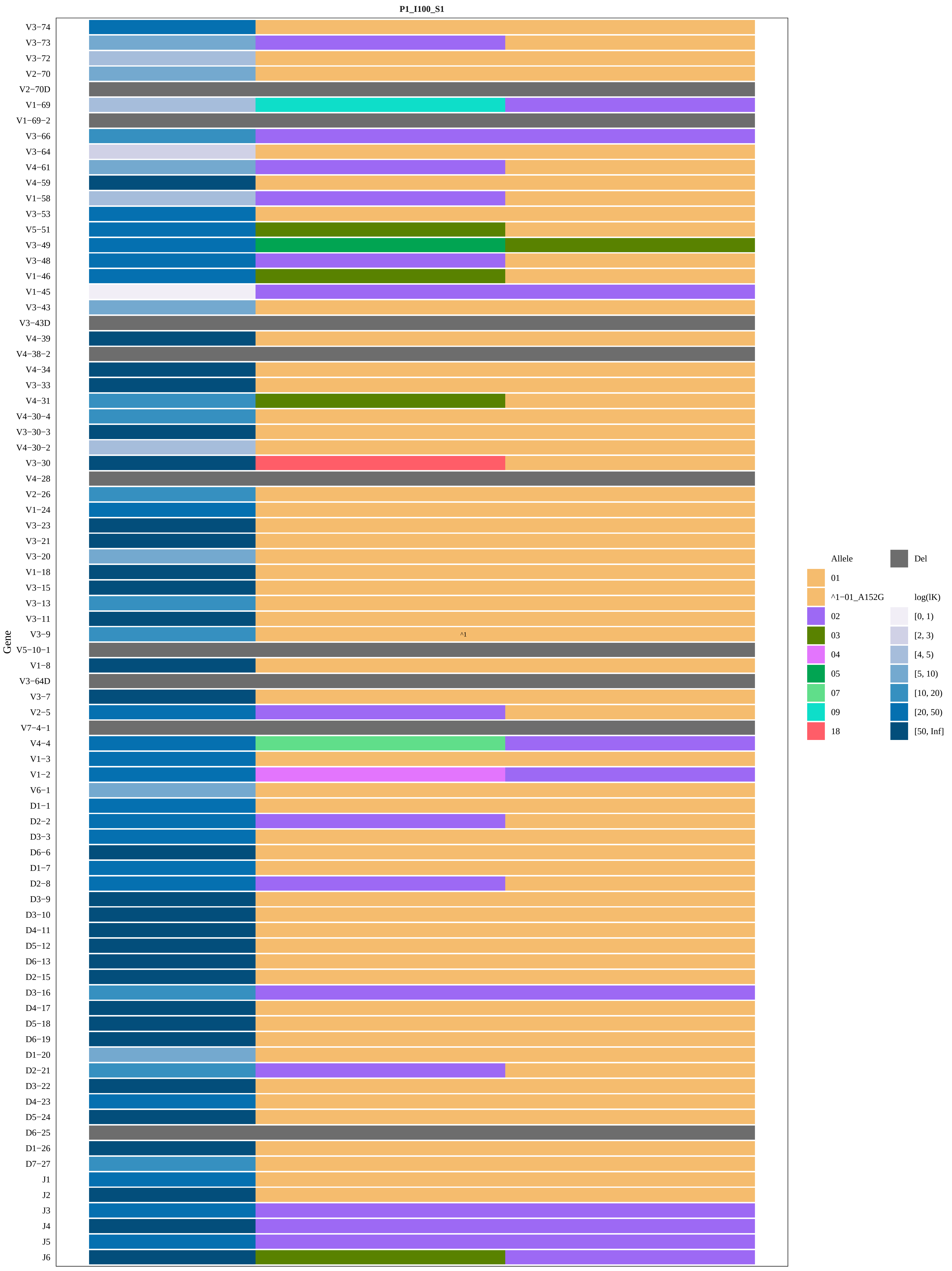

P1_I100_S1
Gene
V3−74
V3−73
V3−72
V2−70
V2−70D
V1−69
V1−69−2
V3−66
V3−64
V4−61
V4−59
V1−58
V3−53
V5−51
V3−49
V3−48
V1−46
V1−45
V3−43
V3−43D
V4−39
V4−38−2
V4−34
V3−33
V4−31
V4−30−4
V3−30−3
V4−30−2
V3−30
V4−28
V2−26
V1−24
V3−23
V3−21
V3−20
V1−18
V3−15
V3−13
V3−11
V3−9
V5−10−1
V1−8
V3−64D
V3−7
V2−5
V7−4−1
V4−4
V1−3
V1−2
V6−1
D1−1
D2−2
D3−3
D6−6
D1−7
D2−8
D3−9
D3−10
D4−11
D5−12
D6−13
D2−15
D3−16
D4−17
D5−18
D6−19
D1−20
D2−21
D3−22
D4−23
D5−24
D6−25
D1−26
D7−27
J1
J2
J3
J4
J5
J6
^1
Allele
01
^1−01_A152G
02
03
04
05
07
09
18
Del
log(lK)
[0, 1)
[2, 3)
[4, 5)
[5, 10)
[10, 20)
[20, 50)
[50, Inf]

P1_I100_S1−IGHJ6−02_03

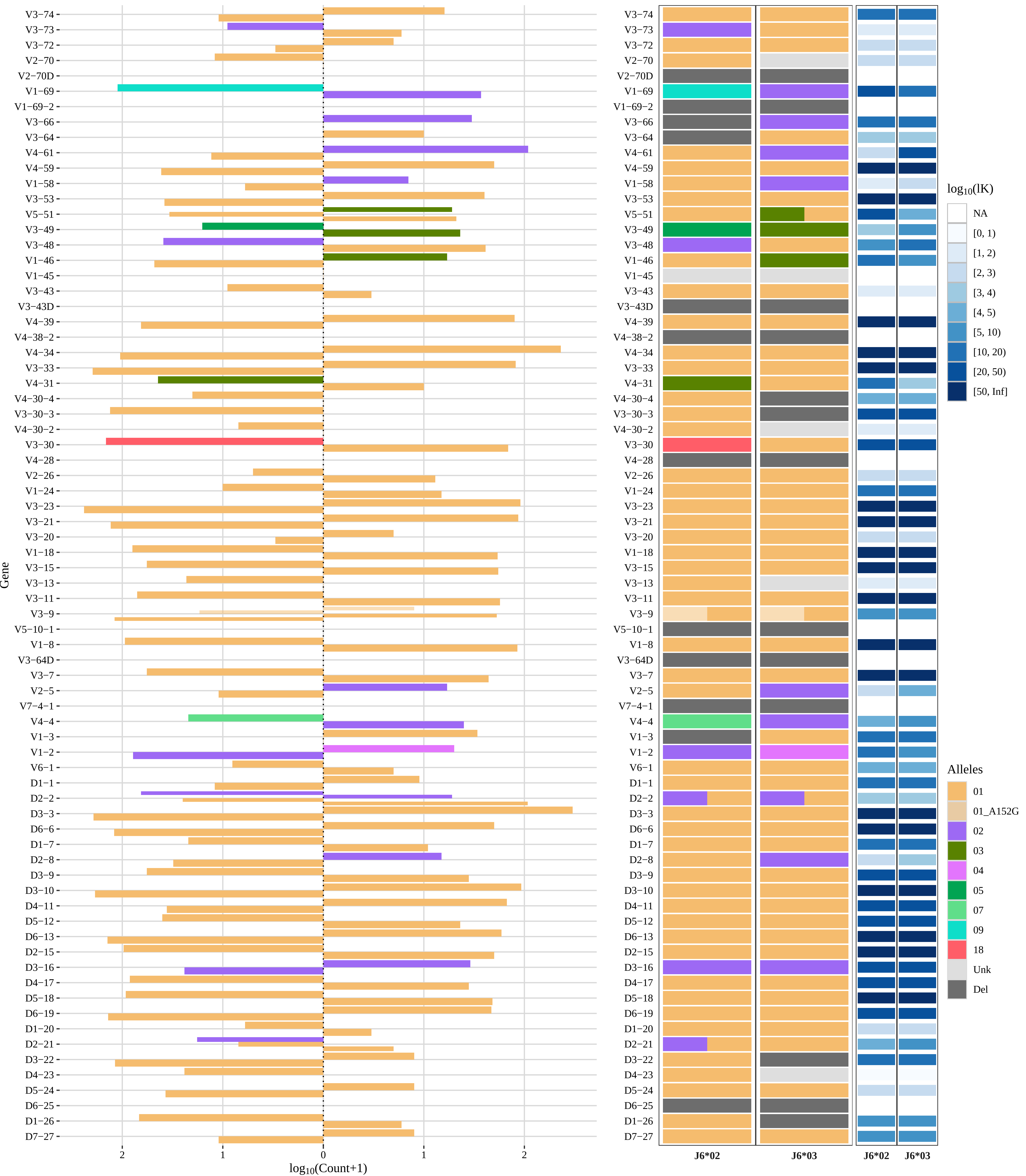
