## Supplementary Figure 2 for "Poorly expressed alleles of several human immunoglobulin heavy chain variable (IGHV) genes are common in the human population"

**Supplementary Figure 2.** Visualization of 35 genotypes of subjects for which haplotyping, based on heterozygosity of IGHJ6, is possible, as inferred by IgDiscover technology (6).

### ERR2567187

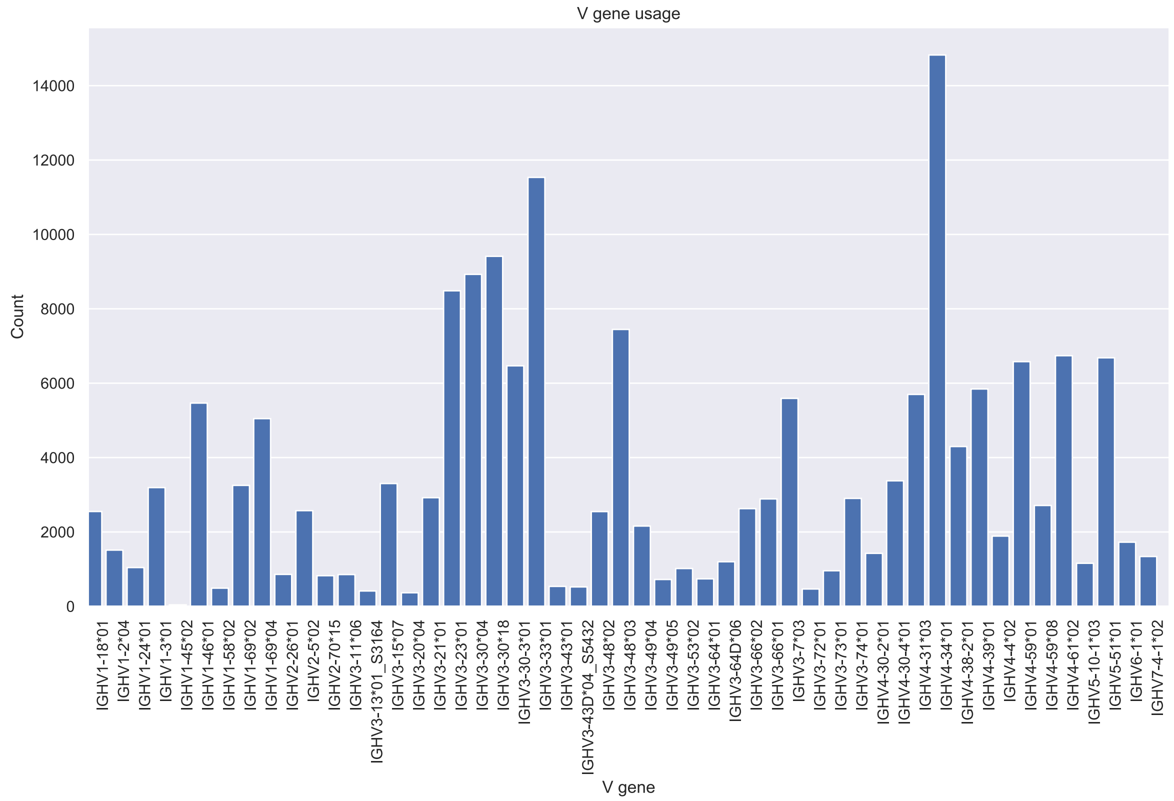
V gene usage
Count
V gene
14000
12000
10000
8000
6000
4000
2000
0
IGHV1-18*01
IGHV1-2*04
IGHV1-24*01
IGHV1-3*01
IGHV1-45*02
IGHV1-46*01
IGHV1-58*02
IGHV1-69*02
IGHV1-69*04
IGHV2-26*01
IGHV2-5*02
IGHV2-70*15
IGHV3-11*06
IGHV3-13*01_S3164
IGHV3-15*07
IGHV3-20*04
IGHV3-21*01
IGHV3-23*01
IGHV3-30*04
IGHV3-30*18
IGHV3-30-3*01
IGHV3-33*01
IGHV3-43*01
IGHV3-43D*04_S5432
IGHV3-48*02
IGHV3-48*03
IGHV3-49*04
IGHV3-49*05
IGHV3-53*02
IGHV3-64*01
IGHV3-64D*06
IGHV3-66*02
IGHV3-66*01
IGHV3-7*03
IGHV3-72*01
IGHV3-73*01
IGHV3-74*01
IGHV4-30-2*01
IGHV4-30-4*01
IGHV4-31*03
IGHV4-34*01
IGHV4-38-2*01
IGHV4-39*01
IGHV4-4*02
IGHV4-59*01
IGHV4-59*08
IGHV4-61*02
IGHV5-10-1*03
IGHV5-51*01
IGHV6-1*01
IGHV7-4-1*02

### ERR2567189

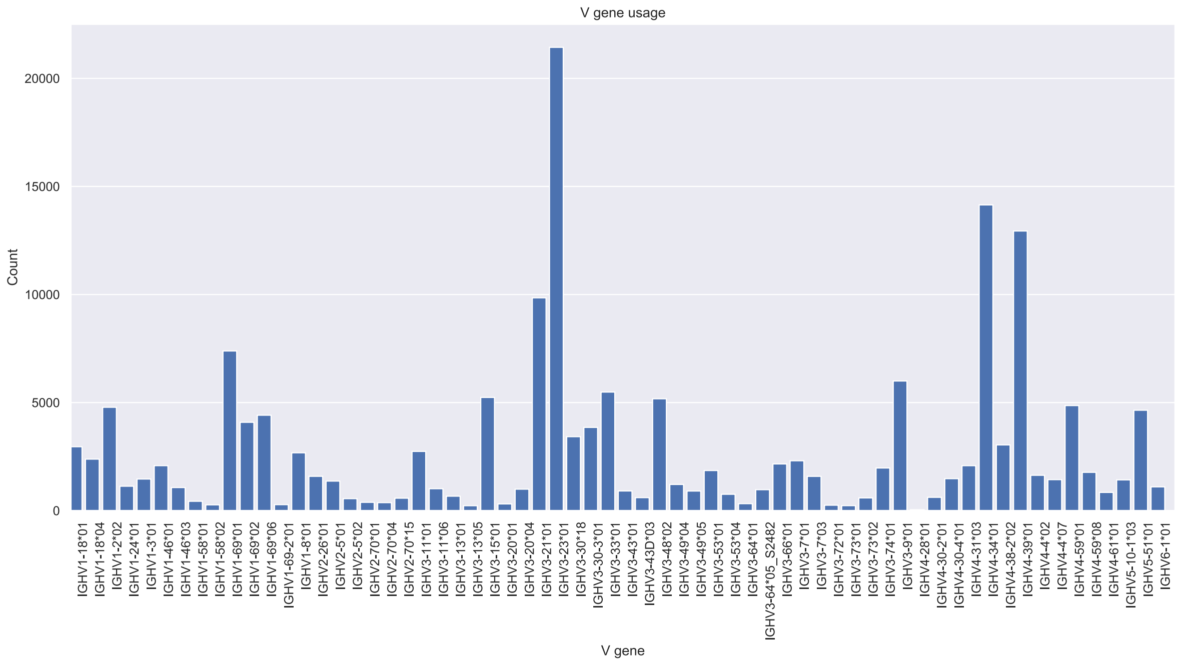
V gene usage
Count
0
5000
10000
15000
20000
IGHV1-18*01
IGHV1-18*04
IGHV1-2*02
IGHV1-24*01
IGHV1-3*01
IGHV1-46*01
IGHV1-46*03
IGHV1-58*01
IGHV1-58*02
IGHV1-69*01
IGHV1-69*02
IGHV1-69*06
IGHV1-69-2*01
IGHV1-8*01
IGHV2-26*01
IGHV2-5*01
IGHV2-5*02
IGHV2-70*01
IGHV2-70*04
IGHV2-70*15
IGHV3-11*01
IGHV3-11*06
IGHV3-13*01
IGHV3-13*05
IGHV3-15*01
IGHV3-20*01
IGHV3-20*04
IGHV3-21*01
IGHV3-23*01
IGHV3-30*18
IGHV3-30-3*01
IGHV3-33*01
IGHV3-43*01
IGHV3-43D*03
IGHV3-48*02
IGHV3-49*04
IGHV3-49*05
IGHV3-53*01
IGHV3-53*04
IGHV3-64*01
IGHV3-64*05_S2482
IGHV3-66*01
IGHV3-7*01
IGHV3-7*03
IGHV3-72*01
IGHV3-73*01
IGHV3-73*02
IGHV3-74*01
IGHV3-9*01
IGHV4-28*01
IGHV4-30-2*01
IGHV4-30-4*01
IGHV4-31*03
IGHV4-34*01
IGHV4-38-2*02
IGHV4-39*01
IGHV4-4*02
IGHV4-4*07
IGHV4-59*01
IGHV4-59*08
IGHV4-61*01
IGHV5-10-1*03
IGHV5-51*01
IGHV6-1*01
V gene

### ERR2567192

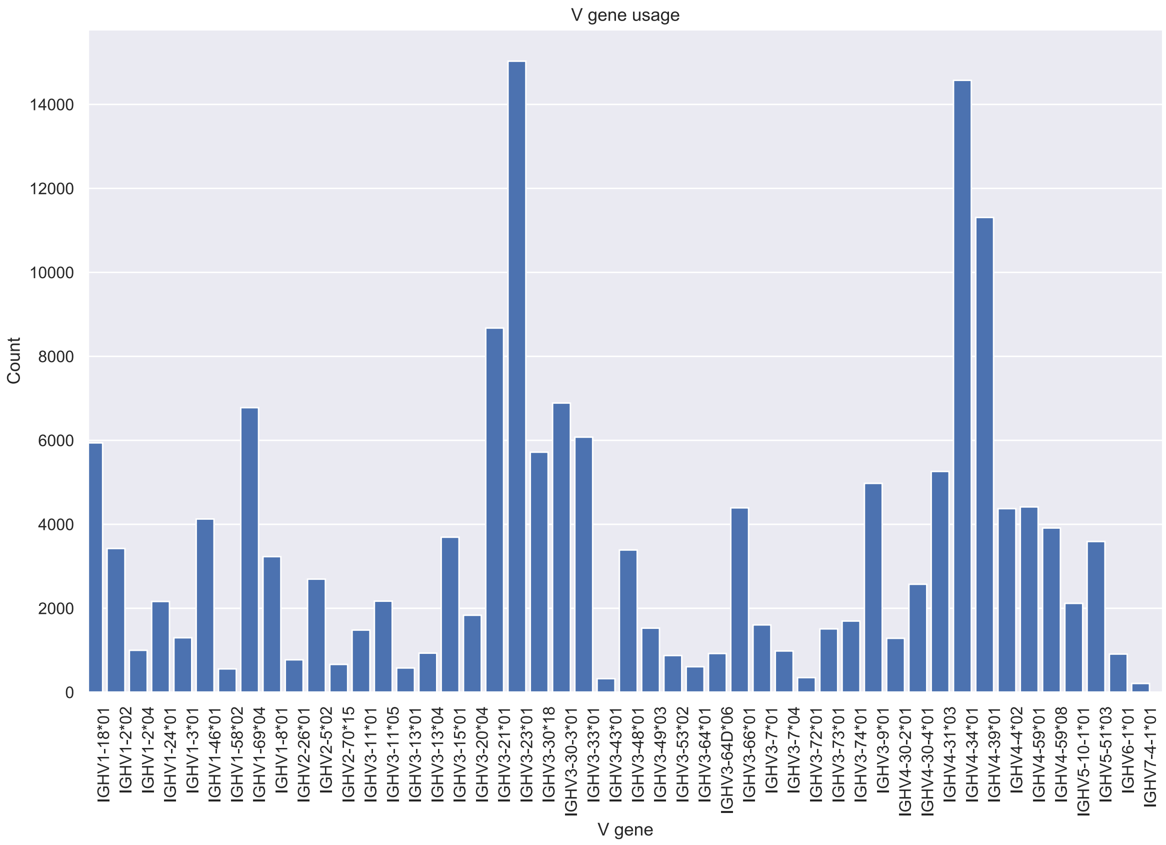
V gene usage
Count
0
2000
4000
6000
8000
10000
12000
14000
IGHV1-18*01
IGHV1-2*02
IGHV1-2*04
IGHV1-24*01
IGHV1-3*01
IGHV1-46*01
IGHV1-58*02
IGHV1-69*04
IGHV1-8*01
IGHV2-26*01
IGHV2-5*02
IGHV2-70*15
IGHV3-11*01
IGHV3-11*05
IGHV3-13*01
IGHV3-13*04
IGHV3-15*01
IGHV3-20*04
IGHV3-21*01
IGHV3-23*01
IGHV3-30*18
IGHV3-30-3*01
IGHV3-33*01
IGHV3-43*01
IGHV3-48*01
IGHV3-49*03
IGHV3-53*02
IGHV3-64*01
IGHV3-64D*06
IGHV3-66*01
IGHV3-7*01
IGHV3-7*04
IGHV3-72*01
IGHV3-73*01
IGHV3-74*01
IGHV3-9*01
IGHV4-30-2*01
IGHV4-30-4*01
IGHV4-31*03
IGHV4-34*01
IGHV4-39*01
IGHV4-4*02
IGHV4-59*01
IGHV4-59*08
IGHV5-10-1*01
IGHV5-51*03
IGHV6-1*01
IGHV7-4-1*01
V gene

### ERR2567199

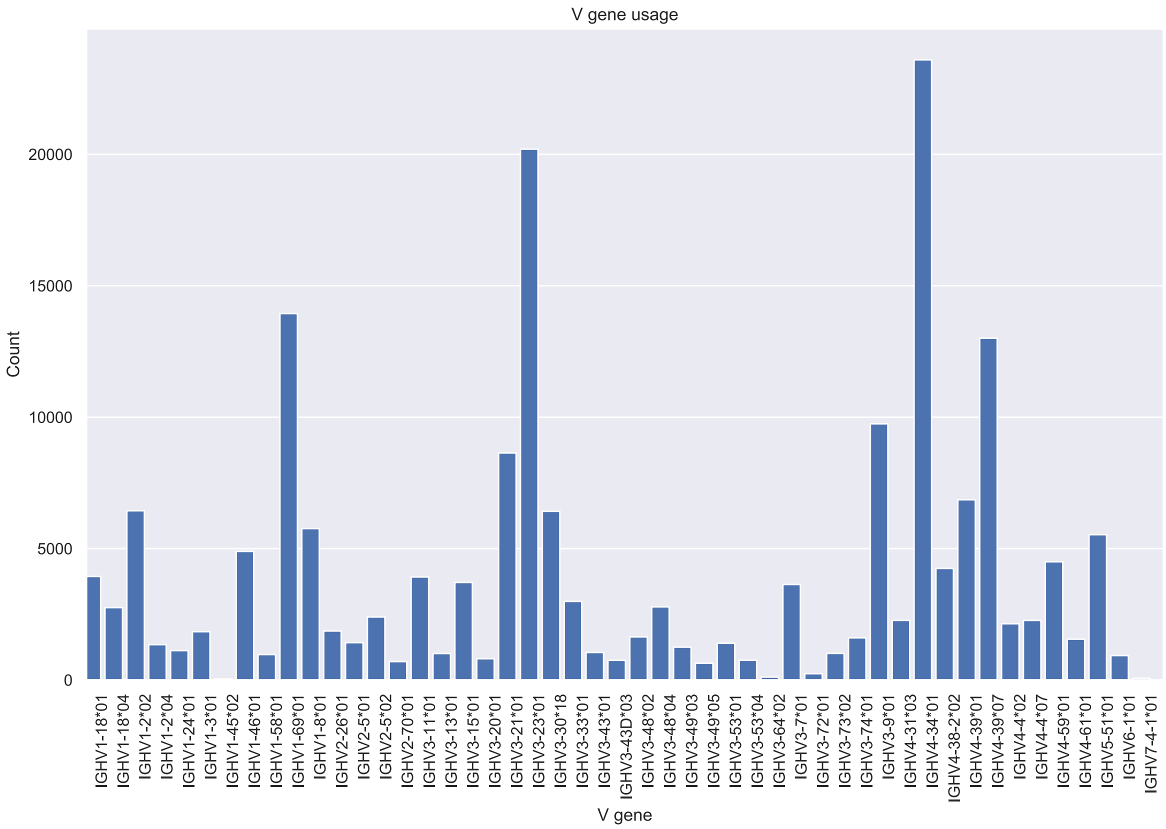

V gene usage
Count
0
5000
10000
15000
20000
V gene
IGHV1-18*01
IGHV1-18*04
IGHV1-2*02
IGHV1-2*04
IGHV1-24*01
IGHV1-3*01
IGHV1-45*02
IGHV1-46*01
IGHV1-58*01
IGHV1-69*01
IGHV1-8*01
IGHV2-26*01
IGHV2-5*01
IGHV2-5*02
IGHV2-70*01
IGHV3-11*01
IGHV3-13*01
IGHV3-15*01
IGHV3-20*01
IGHV3-21*01
IGHV3-23*01
IGHV3-30*18
IGHV3-33*01
IGHV3-43*01
IGHV3-43D*03
IGHV3-48*02
IGHV3-48*04
IGHV3-49*03
IGHV3-49*05
IGHV3-53*01
IGHV3-53*04
IGHV3-64*02
IGHV3-7*01
IGHV3-72*01
IGHV3-73*02
IGHV3-74*01
IGHV3-9*01
IGHV4-31*03
IGHV4-34*01
IGHV4-38-2*02
IGHV4-39*01
IGHV4-39*07
IGHV4-4*02
IGHV4-4*07
IGHV4-59*01
IGHV4-61*01
IGHV5-51*01
IGHV6-1*01
IGHV7-4-1*01

### ERR2567200

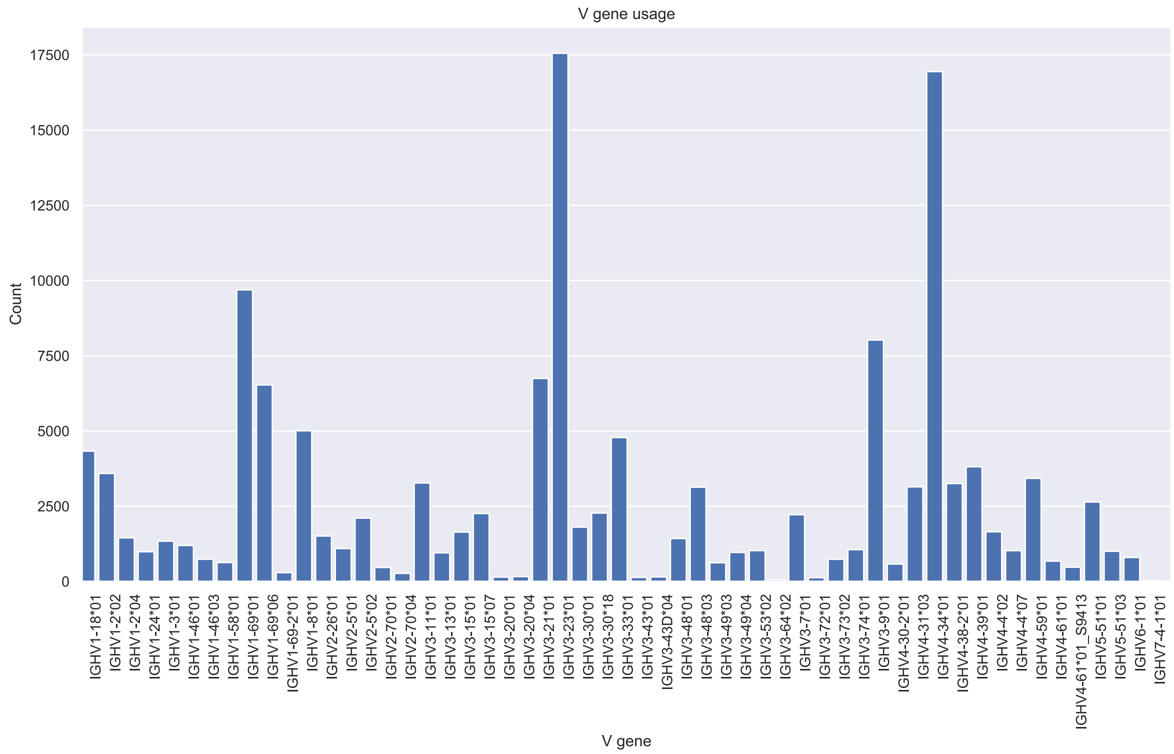

V gene usage
Count
V gene
17500
15000
12500
10000
7500
5000
2500
0
IGHV1-18*01
IGHV1-2*02
IGHV1-2*04
IGHV1-24*01
IGHV1-3*01
IGHV1-46*01
IGHV1-46*03
IGHV1-58*01
IGHV1-69*01
IGHV1-69*06
IGHV1-69-2*01
IGHV1-8*01
IGHV2-26*01
IGHV2-5*01
IGHV2-5*02
IGHV2-70*01
IGHV2-70*04
IGHV3-11*01
IGHV3-13*01
IGHV3-15*01
IGHV3-15*07
IGHV3-20*01
IGHV3-20*04
IGHV3-21*01
IGHV3-23*01
IGHV3-30*01
IGHV3-30*18
IGHV3-33*01
IGHV3-43*01
IGHV3-43D*04
IGHV3-48*01
IGHV3-48*03
IGHV3-49*03
IGHV3-49*04
IGHV3-53*02
IGHV3-64*02
IGHV3-7*01
IGHV3-72*01
IGHV3-73*02
IGHV3-74*01
IGHV3-9*01
IGHV4-30-2*01
IGHV4-31*03
IGHV4-34*01
IGHV4-38-2*01
IGHV4-39*01
IGHV4-4*02
IGHV4-4*07
IGHV4-59*01
IGHV4-61*01
IGHV4-61*01_S9413
IGHV5-51*01
IGHV5-51*03
IGHV6-1*01
IGHV7-4-1*01

### ERR2567201

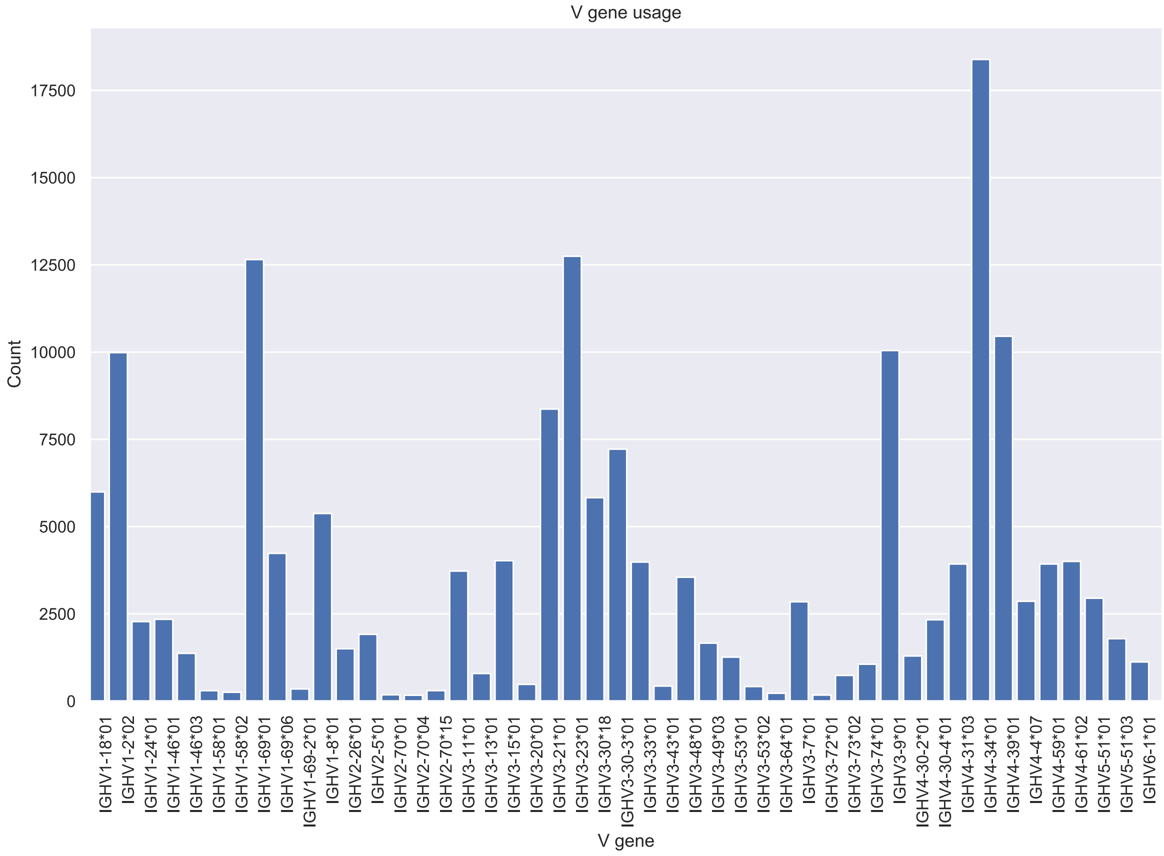

V gene usage
Count
17500
15000
12500
10000
7500
5000
2500
0
IGHV1-18*01
IGHV1-2*02
IGHV1-24*01
IGHV1-46*01
IGHV1-46*03
IGHV1-58*01
IGHV1-58*02
IGHV1-69*01
IGHV1-69*06
IGHV1-69-2*01
IGHV1-8*01
IGHV2-26*01
IGHV2-5*01
IGHV2-70*01
IGHV2-70*04
IGHV2-70*15
IGHV3-11*01
IGHV3-13*01
IGHV3-15*01
IGHV3-20*01
IGHV3-21*01
IGHV3-23*01
IGHV3-30*18
IGHV3-30-3*01
IGHV3-33*01
IGHV3-43*01
IGHV3-48*01
IGHV3-49*03
IGHV3-53*01
IGHV3-53*02
IGHV3-64*01
IGHV3-7*01
IGHV3-72*01
IGHV3-73*02
IGHV3-74*01
IGHV3-9*01
IGHV4-30-2*01
IGHV4-30-4*01
IGHV4-31*03
IGHV4-34*01
IGHV4-39*01
IGHV4-4*07
IGHV4-59*01
IGHV4-61*02
IGHV5-51*01
IGHV5-51*03
IGHV6-1*01
V gene

### ERR2567204

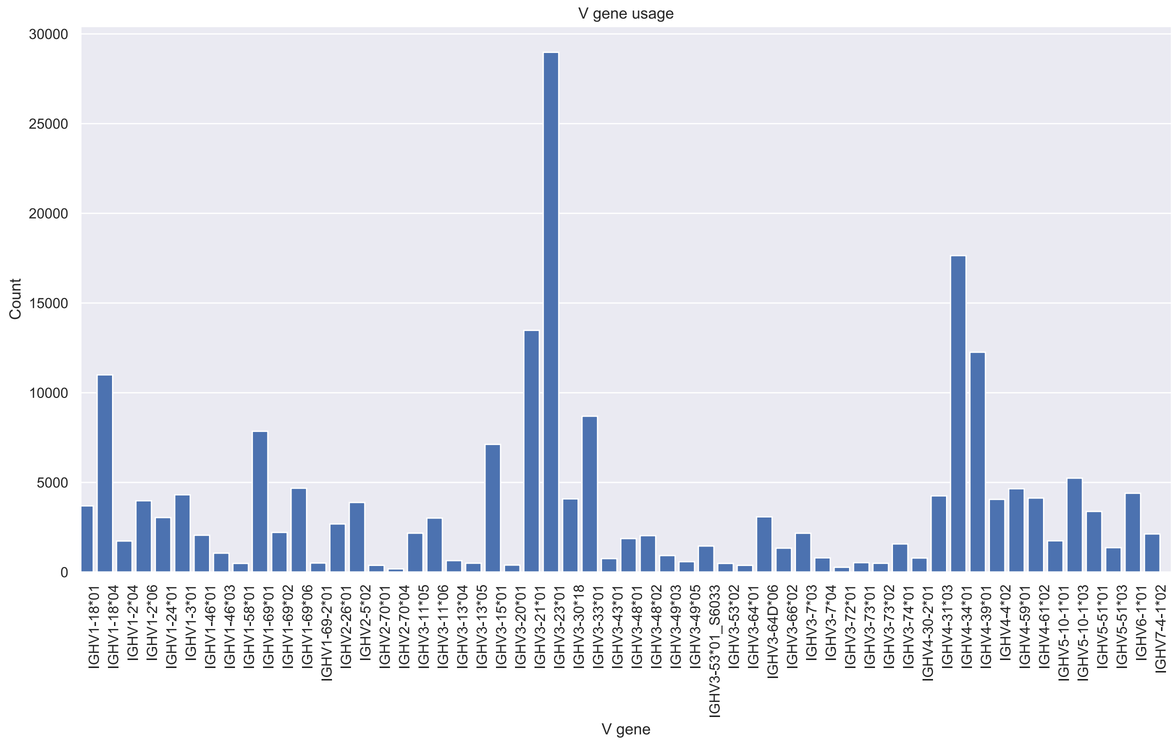

### ERR2567206

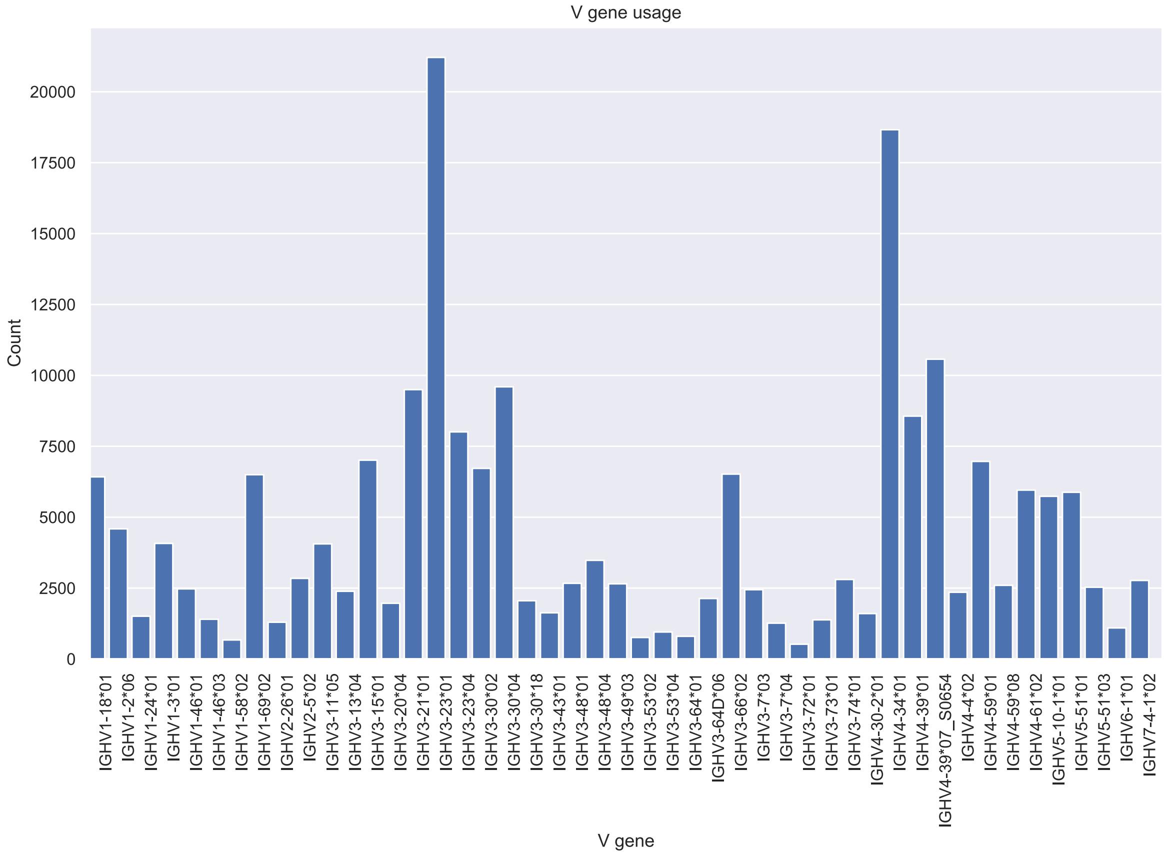

V gene usage
Count
20000
17500
15000
12500
10000
7500
5000
2500
0
IGHV1-18*01
IGHV1-2*06
IGHV1-24*01
IGHV1-3*01
IGHV1-46*01
IGHV1-46*03
IGHV1-58*02
IGHV1-69*02
IGHV2-26*01
IGHV2-5*02
IGHV3-11*05
IGHV3-13*04
IGHV3-15*01
IGHV3-20*04
IGHV3-21*01
IGHV3-23*01
IGHV3-23*04
IGHV3-30*02
IGHV3-30*04
IGHV3-30*18
IGHV3-43*01
IGHV3-48*01
IGHV3-48*04
IGHV3-49*03
IGHV3-53*02
IGHV3-53*04
IGHV3-64*01
IGHV3-64D*06
IGHV3-66*02
IGHV3-7*03
IGHV3-7*04
IGHV3-72*01
IGHV3-73*01
IGHV3-74*01
IGHV4-30-2*01
IGHV4-34*01
IGHV4-39*01
IGHV4-39*07_S0654
IGHV4-4*02
IGHV4-59*01
IGHV4-59*08
IGHV4-61*02
IGHV5-10-1*01
IGHV5-51*01
IGHV5-51*03
IGHV6-1*01
IGHV7-4-1*02
V gene

### ERR2567213

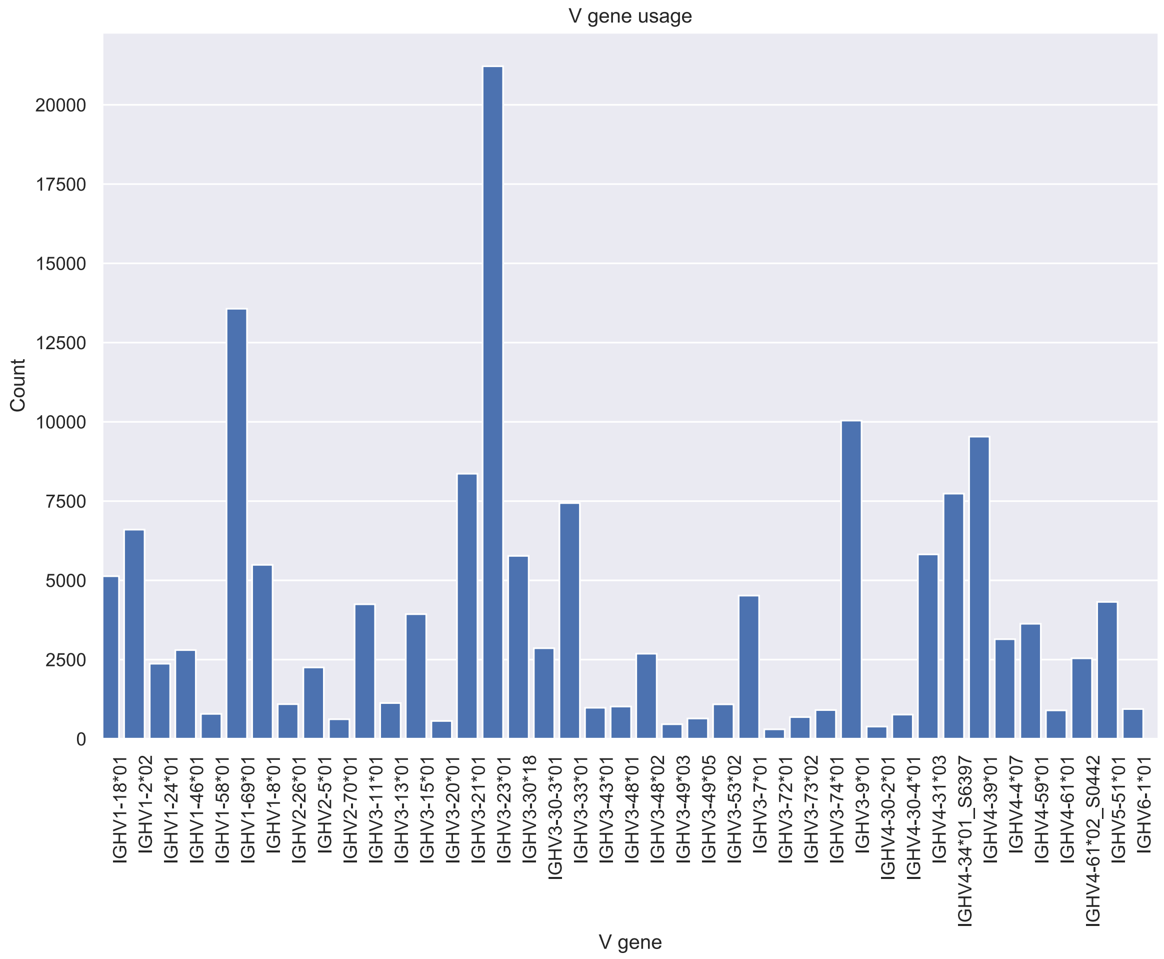

V gene usage
Count
0
2500
5000
7500
10000
12500
15000
17500
20000
IGHV1-18*01
IGHV1-2*02
IGHV1-24*01
IGHV1-46*01
IGHV1-58*01
IGHV1-69*01
IGHV1-8*01
IGHV2-26*01
IGHV2-5*01
IGHV2-70*01
IGHV3-11*01
IGHV3-13*01
IGHV3-15*01
IGHV3-20*01
IGHV3-21*01
IGHV3-23*01
IGHV3-30*18
IGHV3-30-3*01
IGHV3-33*01
IGHV3-43*01
IGHV3-48*01
IGHV3-48*02
IGHV3-49*03
IGHV3-49*05
IGHV3-53*02
IGHV3-7*01
IGHV3-72*01
IGHV3-73*02
IGHV3-74*01
IGHV3-9*01
IGHV4-30-2*01
IGHV4-30-4*01
IGHV4-31*03
IGHV4-34*01_S6397
IGHV4-39*01
IGHV4-4*07
IGHV4-59*01
IGHV4-61*01
IGHV4-61*02_S0442
IGHV5-51*01
IGHV6-1*01
V gene

### ERR2567214

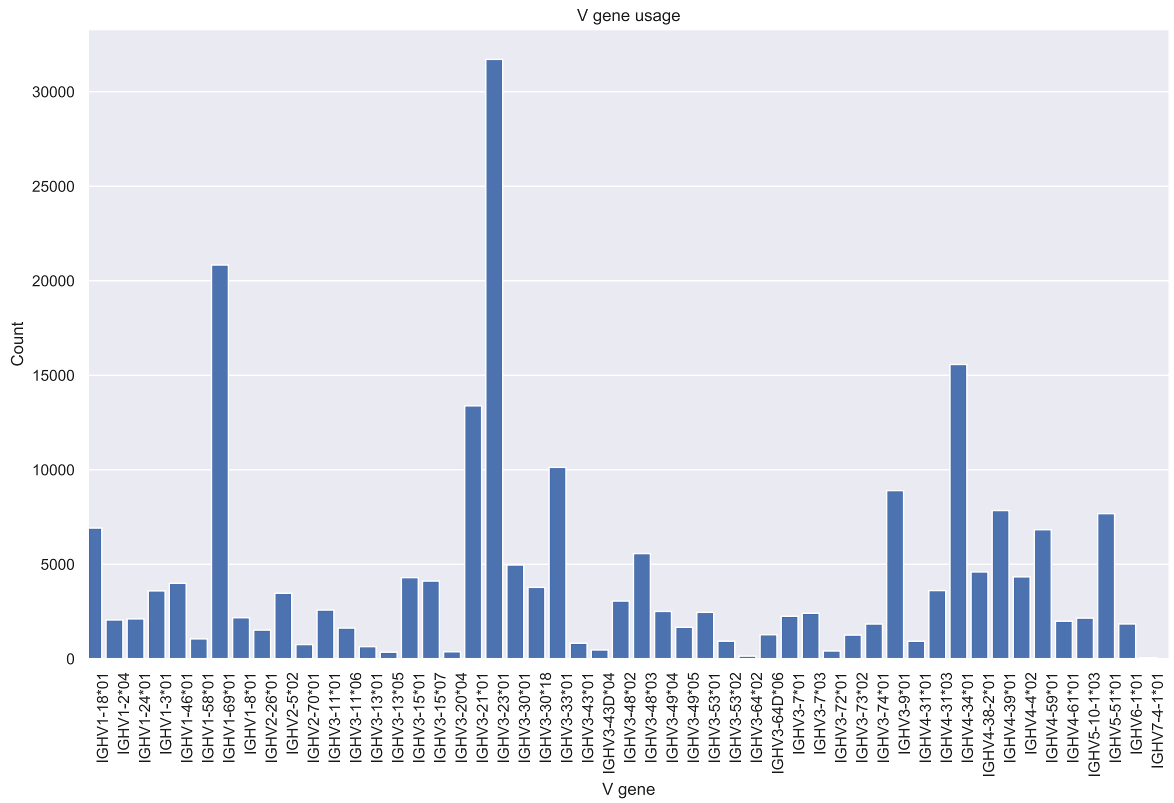
V gene usage
Count
V gene
0
5000
10000
15000
20000
25000
30000
IGHV1-18*01
IGHV1-2*04
IGHV1-24*01
IGHV1-3*01
IGHV1-46*01
IGHV1-58*01
IGHV1-69*01
IGHV1-8*01
IGHV2-26*01
IGHV2-5*02
IGHV2-70*01
IGHV3-11*01
IGHV3-11*06
IGHV3-13*01
IGHV3-13*05
IGHV3-15*01
IGHV3-15*07
IGHV3-20*04
IGHV3-21*01
IGHV3-23*01
IGHV3-30*01
IGHV3-30*18
IGHV3-33*01
IGHV3-43*01
IGHV3-43D*04
IGHV3-48*02
IGHV3-48*03
IGHV3-49*04
IGHV3-49*05
IGHV3-53*01
IGHV3-53*02
IGHV3-64*02
IGHV3-64D*06
IGHV3-7*01
IGHV3-7*03
IGHV3-72*01
IGHV3-73*02
IGHV3-74*01
IGHV3-9*01
IGHV4-31*01
IGHV4-31*03
IGHV4-34*01
IGHV4-38-2*01
IGHV4-39*01
IGHV4-4*02
IGHV4-59*01
IGHV4-61*01
IGHV5-10-1*03
IGHV5-51*01
IGHV6-1*01
IGHV7-4-1*01

### ERR2567215

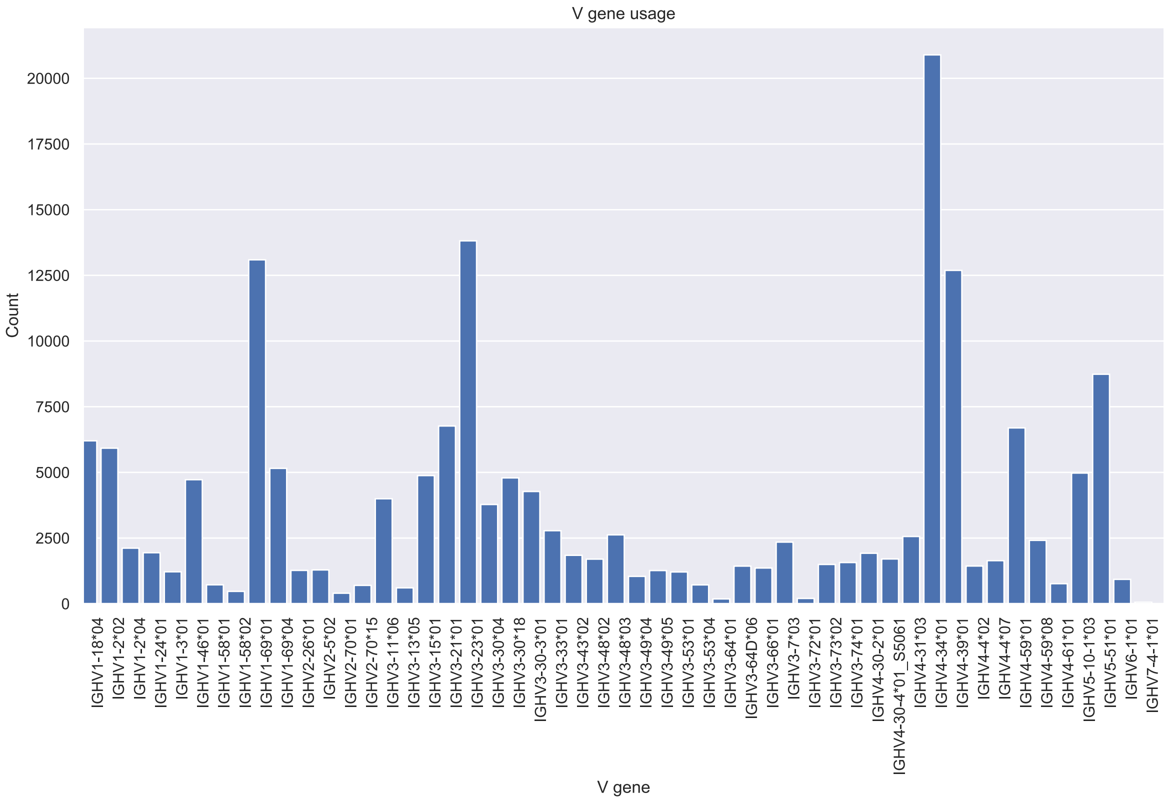
V gene usage
Count
20000
17500
15000
12500
10000
7500
5000
2500
0
V gene
IGHV1-18*04
IGHV1-2*02
IGHV1-2*04
IGHV1-24*01
IGHV1-3*01
IGHV1-46*01
IGHV1-58*01
IGHV1-58*02
IGHV1-69*01
IGHV1-69*04
IGHV2-26*01
IGHV2-5*02
IGHV2-70*01
IGHV2-70*15
IGHV3-11*06
IGHV3-13*05
IGHV3-15*01
IGHV3-21*01
IGHV3-23*01
IGHV3-30*04
IGHV3-30*18
IGHV3-30-3*01
IGHV3-33*01
IGHV3-43*02
IGHV3-48*02
IGHV3-48*03
IGHV3-49*04
IGHV3-49*05
IGHV3-53*01
IGHV3-53*04
IGHV3-64*01
IGHV3-64D*06
IGHV3-66*01
IGHV3-7*03
IGHV3-72*01
IGHV3-73*02
IGHV3-74*01
IGHV4-30-2*01
IGHV4-30-4*01_S5061
IGHV4-31*03
IGHV4-34*01
IGHV4-39*01
IGHV4-4*02
IGHV4-4*07
IGHV4-59*01
IGHV4-59*08
IGHV4-61*01
IGHV5-10-1*03
IGHV5-51*01
IGHV6-1*01
IGHV7-4-1*01

### ERR2567217

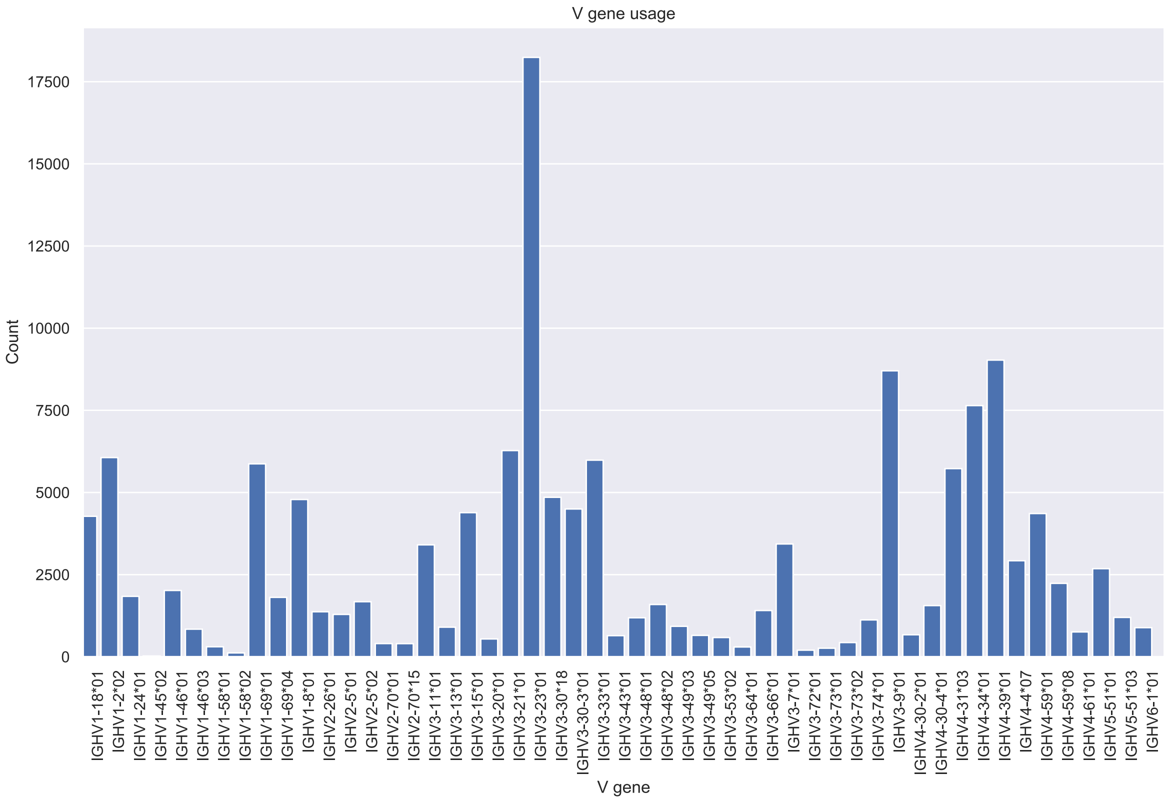
V gene usage
Count
0
2500
5000
7500
10000
12500
15000
17500
V gene
IGHV1-18*01
IGHV1-2*02
IGHV1-24*01
IGHV1-45*02
IGHV1-46*01
IGHV1-46*03
IGHV1-58*01
IGHV1-58*02
IGHV1-69*01
IGHV1-69*04
IGHV1-8*01
IGHV2-26*01
IGHV2-5*01
IGHV2-5*02
IGHV2-70*01
IGHV2-70*15
IGHV3-11*01
IGHV3-13*01
IGHV3-15*01
IGHV3-20*01
IGHV3-21*01
IGHV3-23*01
IGHV3-30*18
IGHV3-30-3*01
IGHV3-33*01
IGHV3-43*01
IGHV3-48*01
IGHV3-48*02
IGHV3-49*03
IGHV3-49*05
IGHV3-53*02
IGHV3-64*01
IGHV3-66*01
IGHV3-7*01
IGHV3-72*01
IGHV3-73*01
IGHV3-73*02
IGHV3-74*01
IGHV3-9*01
IGHV4-30-2*01
IGHV4-30-4*01
IGHV4-31*03
IGHV4-34*01
IGHV4-39*01
IGHV4-4*07
IGHV4-59*01
IGHV4-59*08
IGHV4-61*01
IGHV5-51*01
IGHV5-51*03
IGHV6-1*01

### ERR2567220

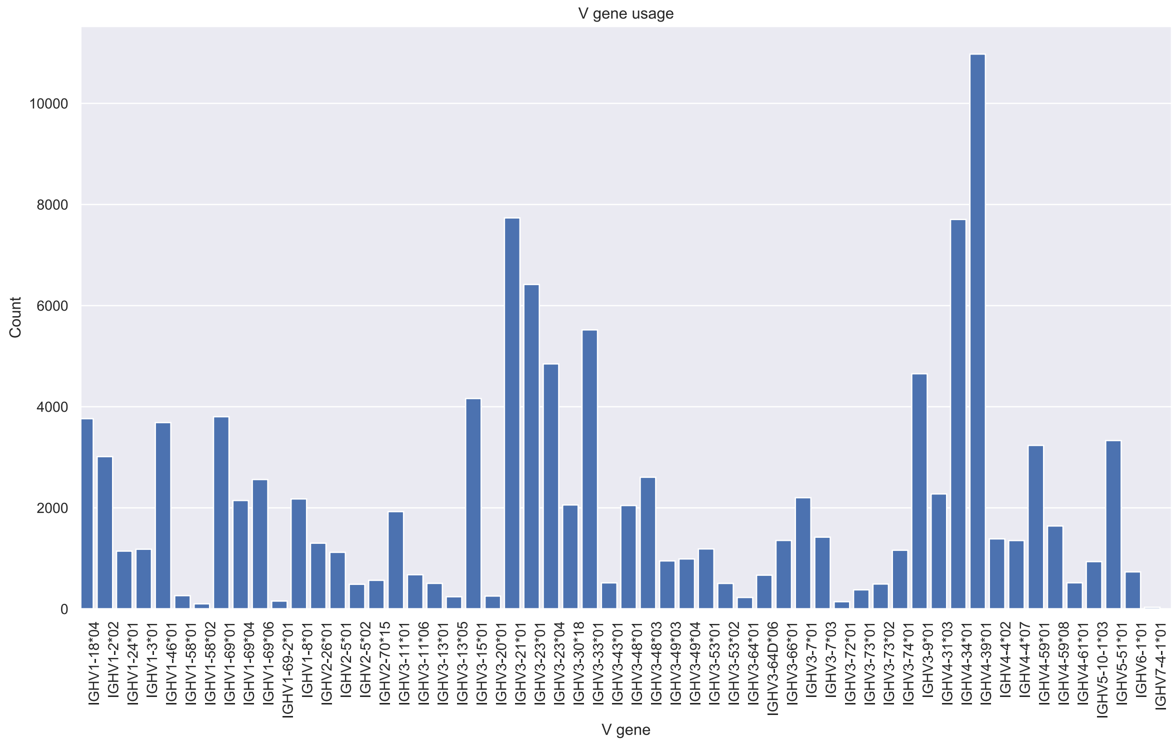

V gene usage
Count
10000
8000
6000
4000
2000
0
IGHV1-18*04
IGHV1-2*02
IGHV1-24*01
IGHV1-3*01
IGHV1-46*01
IGHV1-58*01
IGHV1-58*02
IGHV1-69*01
IGHV1-69*04
IGHV1-69*06
IGHV1-69-2*01
IGHV1-8*01
IGHV2-26*01
IGHV2-5*01
IGHV2-5*02
IGHV2-70*15
IGHV3-11*01
IGHV3-11*06
IGHV3-13*01
IGHV3-13*05
IGHV3-15*01
IGHV3-20*01
IGHV3-21*01
IGHV3-23*01
IGHV3-23*04
IGHV3-30*18
IGHV3-33*01
IGHV3-43*01
IGHV3-48*01
IGHV3-48*03
IGHV3-49*03
IGHV3-49*04
IGHV3-53*01
IGHV3-53*02
IGHV3-64*01
IGHV3-64D*06
IGHV3-66*01
IGHV3-7*01
IGHV3-7*03
IGHV3-72*01
IGHV3-73*01
IGHV3-73*02
IGHV3-74*01
IGHV3-9*01
IGHV4-31*03
IGHV4-34*01
IGHV4-39*01
IGHV4-4*02
IGHV4-4*07
IGHV4-59*01
IGHV4-59*08
IGHV4-61*01
IGHV5-10-1*03
IGHV5-51*01
IGHV6-1*01
IGHV7-4-1*01
V gene

### ERR2567221

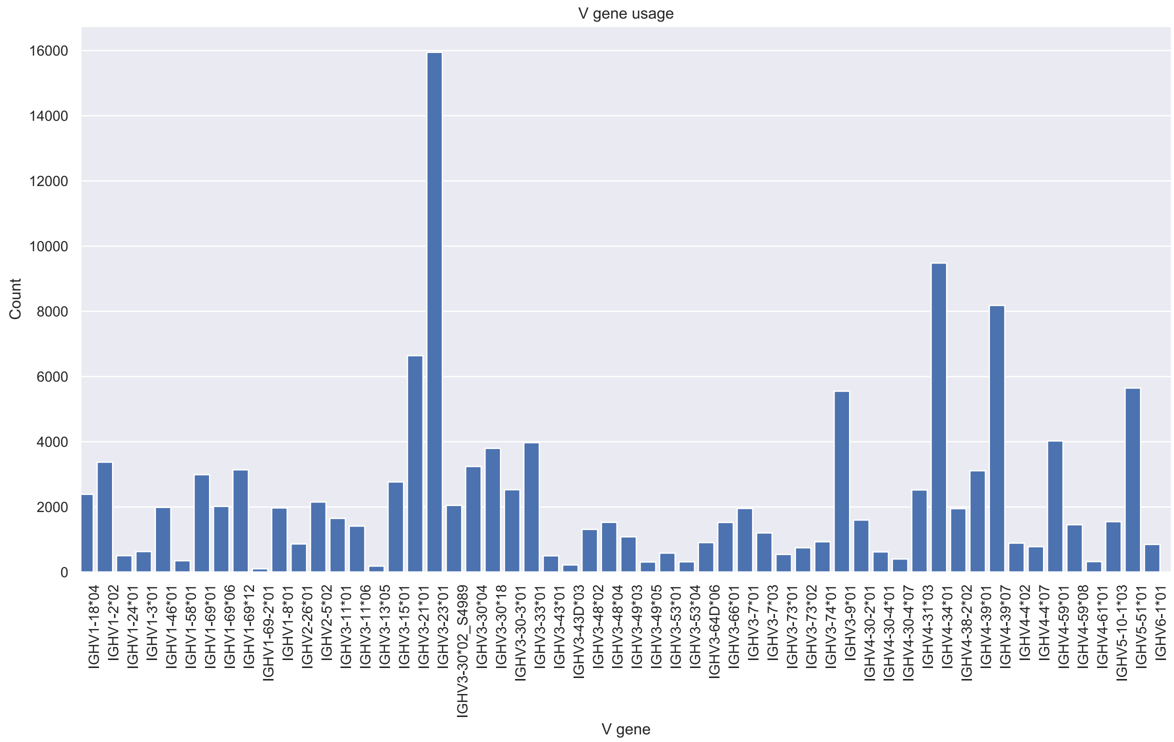
V gene usage
Count
V gene
0
2000
4000
6000
8000
10000
12000
14000
16000
IGHV1-18*04
IGHV1-2*02
IGHV1-24*01
IGHV1-3*01
IGHV1-46*01
IGHV1-58*01
IGHV1-69*01
IGHV1-69*06
IGHV1-69*12
IGHV1-69-2*01
IGHV1-8*01
IGHV2-26*01
IGHV2-5*02
IGHV3-11*01
IGHV3-11*06
IGHV3-13*05
IGHV3-15*01
IGHV3-21*01
IGHV3-23*01
IGHV3-30*02_S4989
IGHV3-30*04
IGHV3-30*18
IGHV3-30-3*01
IGHV3-33*01
IGHV3-43*01
IGHV3-43D*03
IGHV3-48*02
IGHV3-48*04
IGHV3-49*03
IGHV3-49*05
IGHV3-53*01
IGHV3-53*04
IGHV3-64D*06
IGHV3-66*01
IGHV3-7*01
IGHV3-7*03
IGHV3-73*01
IGHV3-73*02
IGHV3-74*01
IGHV3-9*01
IGHV4-30-2*01
IGHV4-30-4*01
IGHV4-30-4*07
IGHV4-31*03
IGHV4-34*01
IGHV4-38-2*02
IGHV4-39*01
IGHV4-39*07
IGHV4-4*02
IGHV4-4*07
IGHV4-59*01
IGHV4-59*08
IGHV4-61*01
IGHV5-10-1*03
IGHV5-51*01
IGHV6-1*01

### ERR2567223

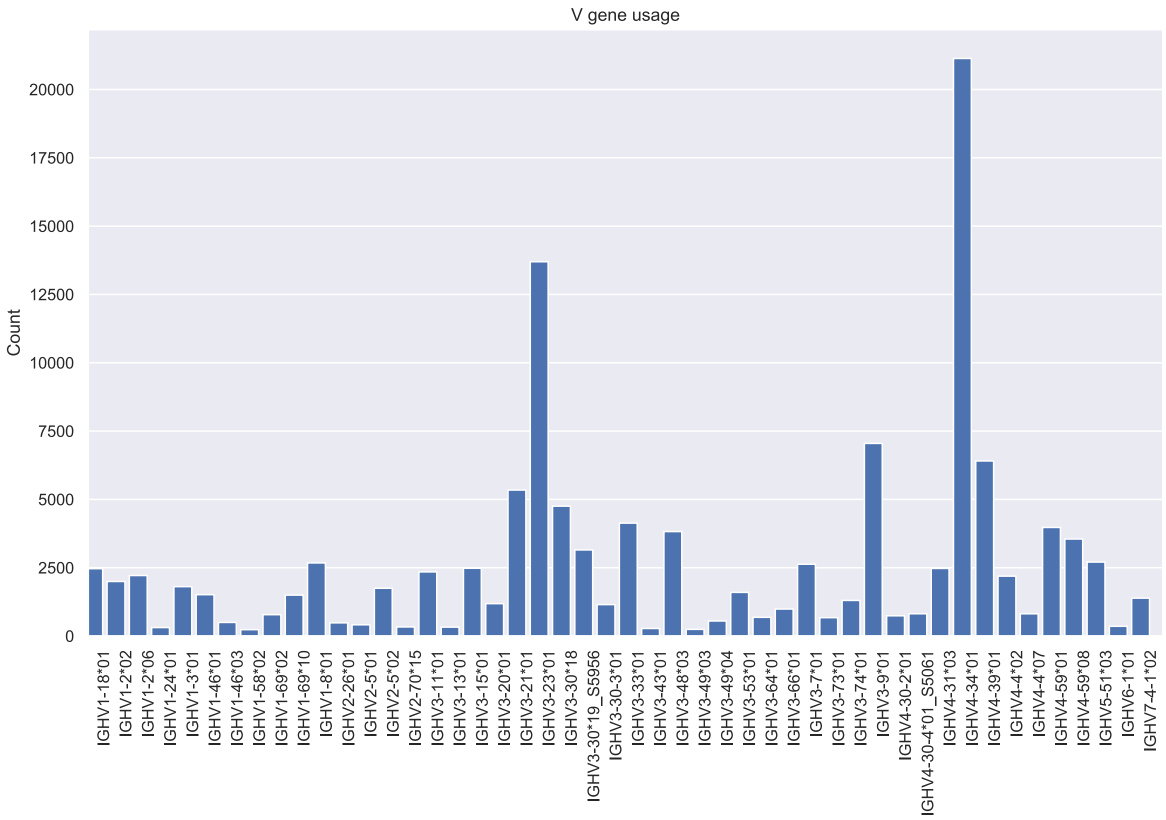
V gene usage
Count
V gene
0
2500
5000
7500
10000
12500
15000
17500
20000
IGHV1-18*01
IGHV1-2*02
IGHV1-2*06
IGHV1-24*01
IGHV1-3*01
IGHV1-46*01
IGHV1-46*03
IGHV1-58*02
IGHV1-69*02
IGHV1-69*10
IGHV1-8*01
IGHV2-26*01
IGHV2-5*01
IGHV2-5*02
IGHV2-70*15
IGHV3-11*01
IGHV3-13*01
IGHV3-15*01
IGHV3-20*01
IGHV3-21*01
IGHV3-23*01
IGHV3-30*18
IGHV3-30*19_S5956
IGHV3-30-3*01
IGHV3-33*01
IGHV3-43*01
IGHV3-48*03
IGHV3-49*03
IGHV3-49*04
IGHV3-53*01
IGHV3-64*01
IGHV3-66*01
IGHV3-7*01
IGHV3-73*01
IGHV3-74*01
IGHV3-9*01
IGHV4-30-2*01
IGHV4-30-4*01_S5061
IGHV4-31*03
IGHV4-34*01
IGHV4-39*01
IGHV4-4*02
IGHV4-4*07
IGHV4-59*01
IGHV4-59*08
IGHV5-51*03
IGHV6-1*01
IGHV7-4-1*02

### ERR2567226

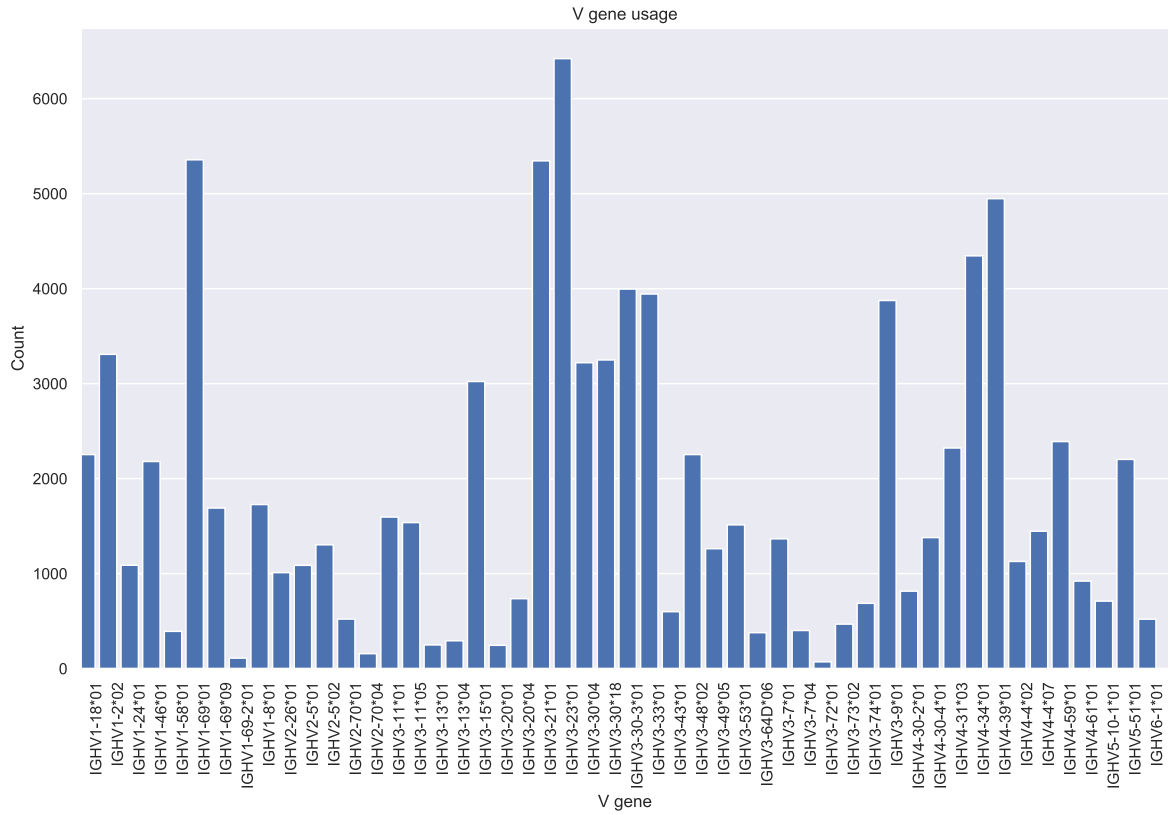
V gene usage
Count
0
1000
2000
3000
4000
5000
6000
V gene
IGHV1-18*01
IGHV1-2*02
IGHV1-24*01
IGHV1-46*01
IGHV1-58*01
IGHV1-69*01
IGHV1-69*09
IGHV1-69-2*01
IGHV1-8*01
IGHV2-26*01
IGHV2-5*01
IGHV2-5*02
IGHV2-70*01
IGHV2-70*04
IGHV3-11*01
IGHV3-11*05
IGHV3-13*01
IGHV3-13*04
IGHV3-15*01
IGHV3-20*01
IGHV3-20*04
IGHV3-21*01
IGHV3-23*01
IGHV3-30*04
IGHV3-30*18
IGHV3-30-3*01
IGHV3-33*01
IGHV3-43*01
IGHV3-48*02
IGHV3-49*05
IGHV3-53*01
IGHV3-64D*06
IGHV3-7*01
IGHV3-7*04
IGHV3-72*01
IGHV3-73*02
IGHV3-74*01
IGHV3-9*01
IGHV4-30-2*01
IGHV4-30-4*01
IGHV4-31*03
IGHV4-34*01
IGHV4-39*01
IGHV4-4*02
IGHV4-4*07
IGHV4-59*01
IGHV4-61*01
IGHV5-10-1*01
IGHV5-51*01
IGHV6-1*01

### ERR2567230

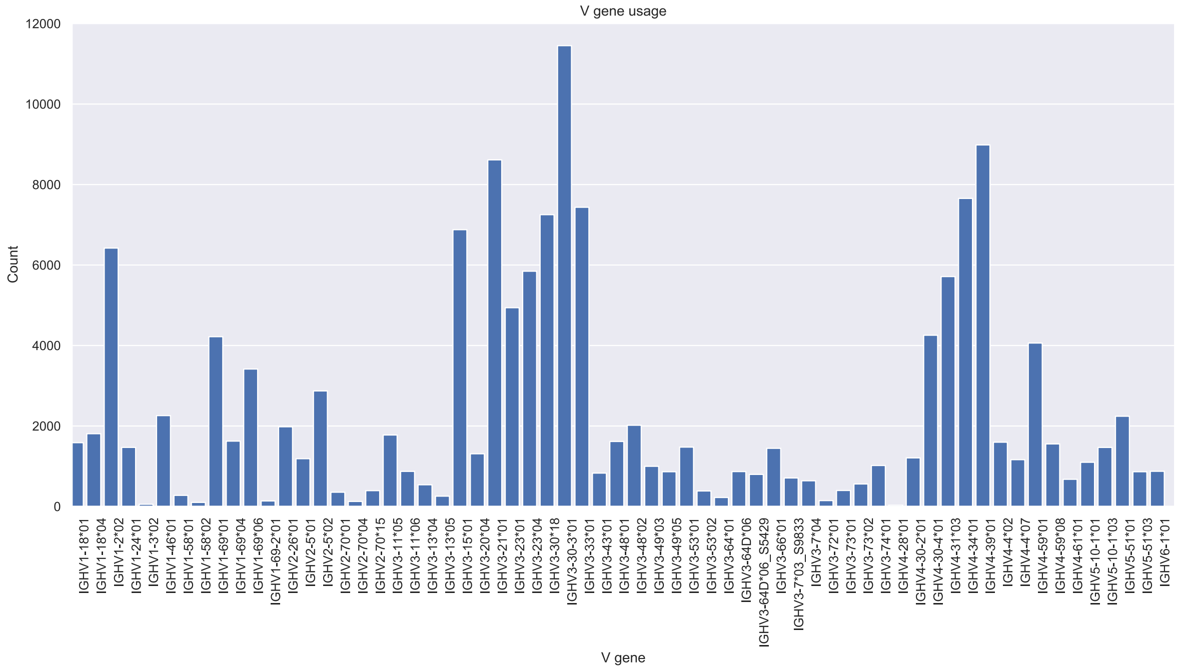

V gene usage
Count
12000
10000
8000
6000
4000
2000
0
IGHV1-18*01
IGHV1-18*04
IGHV1-2*02
IGHV1-24*01
IGHV1-3*02
IGHV1-46*01
IGHV1-58*01
IGHV1-58*02
IGHV1-69*01
IGHV1-69*04
IGHV1-69*06
IGHV1-69-2*01
IGHV2-26*01
IGHV2-5*01
IGHV2-5*02
IGHV2-70*01
IGHV2-70*04
IGHV2-70*15
IGHV3-11*05
IGHV3-11*06
IGHV3-13*04
IGHV3-13*05
IGHV3-15*01
IGHV3-20*04
IGHV3-21*01
IGHV3-23*01
IGHV3-23*04
IGHV3-30*18
IGHV3-30-3*01
IGHV3-33*01
IGHV3-43*01
IGHV3-48*01
IGHV3-48*02
IGHV3-49*03
IGHV3-49*05
IGHV3-53*01
IGHV3-53*02
IGHV3-64*01
IGHV3-64D*06
IGHV3-64D*06_S5429
IGHV3-66*01
IGHV3-7*03_S9833
IGHV3-7*04
IGHV3-72*01
IGHV3-73*01
IGHV3-73*02
IGHV3-74*01
IGHV4-28*01
IGHV4-30-2*01
IGHV4-30-4*01
IGHV4-31*03
IGHV4-34*01
IGHV4-39*01
IGHV4-4*02
IGHV4-4*07
IGHV4-59*01
IGHV4-59*08
IGHV4-61*01
IGHV5-10-1*01
IGHV5-10-1*03
IGHV5-51*01
IGHV5-51*03
IGHV6-1*01
V gene

### ERR2567231

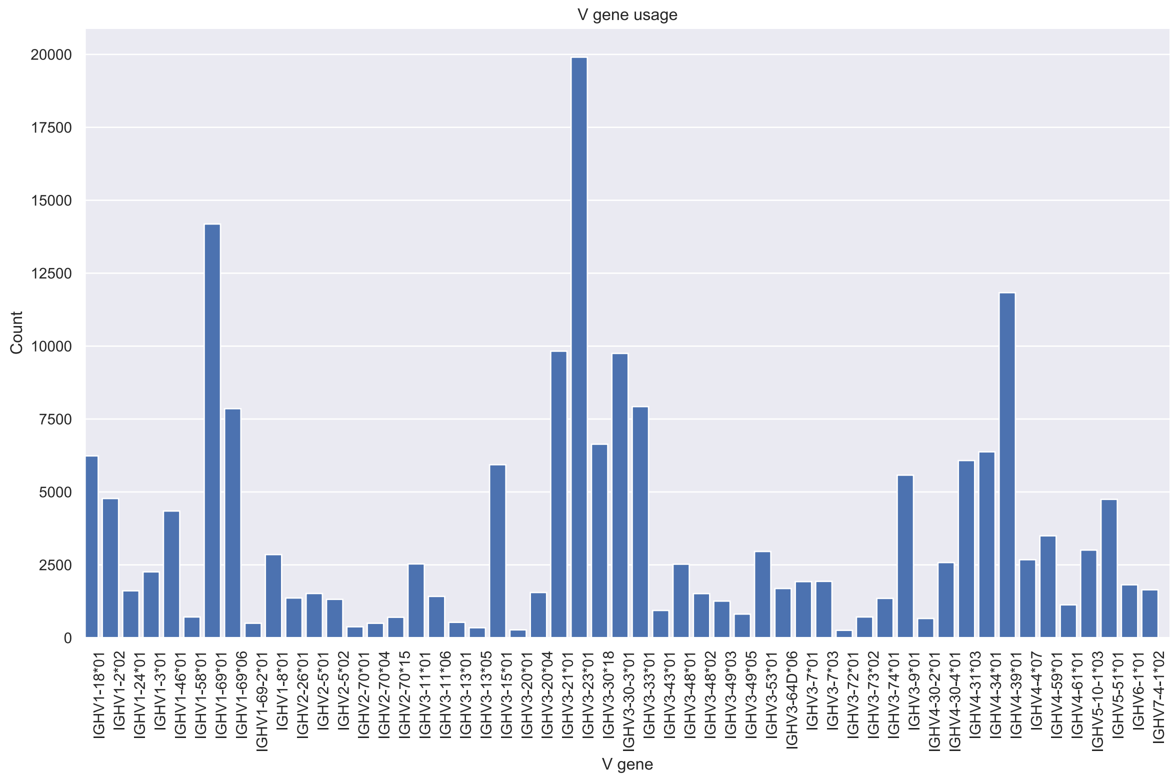

### ERR2567232

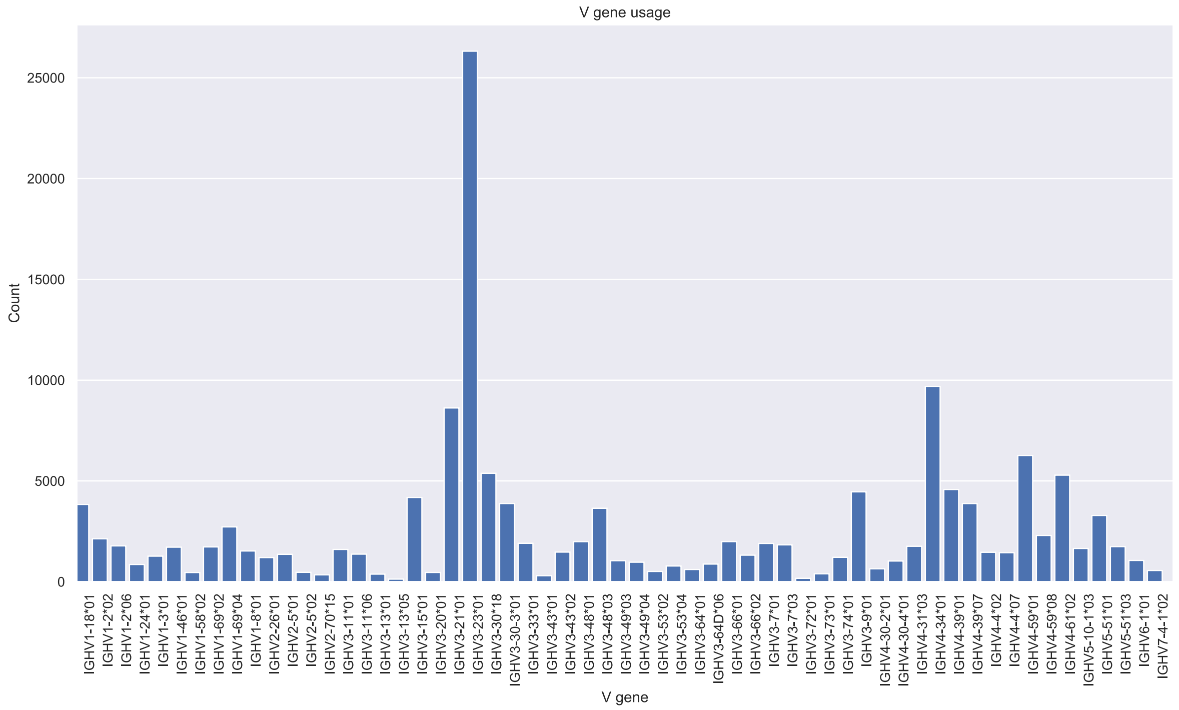
V gene usage
Count
V gene
25000
20000
15000
10000
5000
0
IGHV1-18*01
IGHV1-2*02
IGHV1-2*06
IGHV1-24*01
IGHV1-3*01
IGHV1-46*01
IGHV1-58*02
IGHV1-69*02
IGHV1-69*04
IGHV1-8*01
IGHV2-26*01
IGHV2-5*01
IGHV2-5*02
IGHV2-70*15
IGHV3-11*01
IGHV3-11*06
IGHV3-13*01
IGHV3-13*05
IGHV3-15*01
IGHV3-20*01
IGHV3-21*01
IGHV3-23*01
IGHV3-30*18
IGHV3-30-3*01
IGHV3-33*01
IGHV3-43*01
IGHV3-43*02
IGHV3-48*01
IGHV3-48*03
IGHV3-49*03
IGHV3-49*04
IGHV3-53*02
IGHV3-53*04
IGHV3-64*01
IGHV3-64D*06
IGHV3-66*01
IGHV3-66*02
IGHV3-7*01
IGHV3-7*03
IGHV3-72*01
IGHV3-73*01
IGHV3-74*01
IGHV3-9*01
IGHV4-30-2*01
IGHV4-30-4*01
IGHV4-31*03
IGHV4-34*01
IGHV4-39*01
IGHV4-39*07
IGHV4-4*02
IGHV4-4*07
IGHV4-59*01
IGHV4-59*08
IGHV4-61*02
IGHV5-10-1*03
IGHV5-51*01
IGHV5-51*03
IGHV6-1*01
IGHV7-4-1*02

### ERR2567240

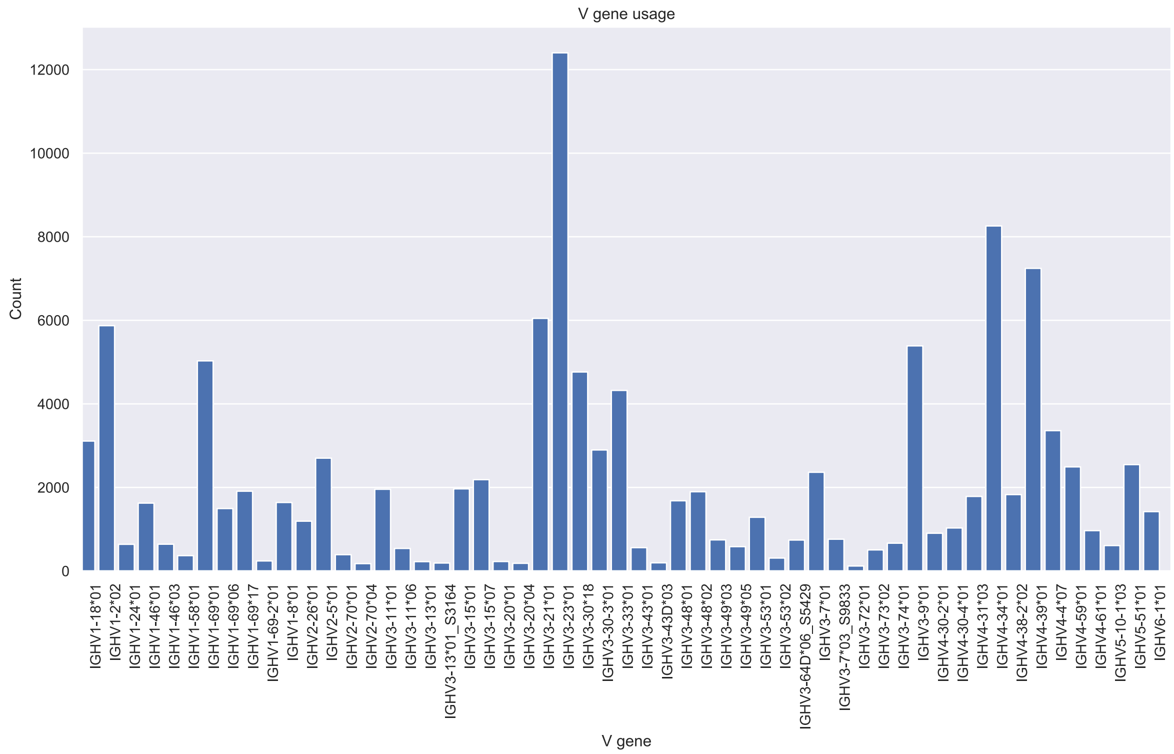

V gene usage
Count
V gene
0
2000
4000
6000
8000
10000
12000
IGHV1-18*01
IGHV1-2*02
IGHV1-24*01
IGHV1-46*01
IGHV1-46*03
IGHV1-58*01
IGHV1-69*01
IGHV1-69*06
IGHV1-69*17
IGHV1-69-2*01
IGHV1-8*01
IGHV2-26*01
IGHV2-5*01
IGHV2-70*01
IGHV2-70*04
IGHV3-11*01
IGHV3-11*06
IGHV3-13*01
IGHV3-13*01_S3164
IGHV3-15*01
IGHV3-15*07
IGHV3-20*01
IGHV3-20*04
IGHV3-21*01
IGHV3-23*01
IGHV3-30*18
IGHV3-30-3*01
IGHV3-33*01
IGHV3-43*01
IGHV3-43D*03
IGHV3-48*01
IGHV3-48*02
IGHV3-49*03
IGHV3-49*05
IGHV3-53*01
IGHV3-53*02
IGHV3-64D*06_S5429
IGHV3-7*01
IGHV3-7*03_S9833
IGHV3-72*01
IGHV3-73*02
IGHV3-74*01
IGHV3-9*01
IGHV4-30-2*01
IGHV4-30-4*01
IGHV4-31*03
IGHV4-34*01
IGHV4-38-2*02
IGHV4-39*01
IGHV4-4*07
IGHV4-59*01
IGHV4-61*01
IGHV5-10-1*03
IGHV5-51*01
IGHV6-1*01

### ERR2567242

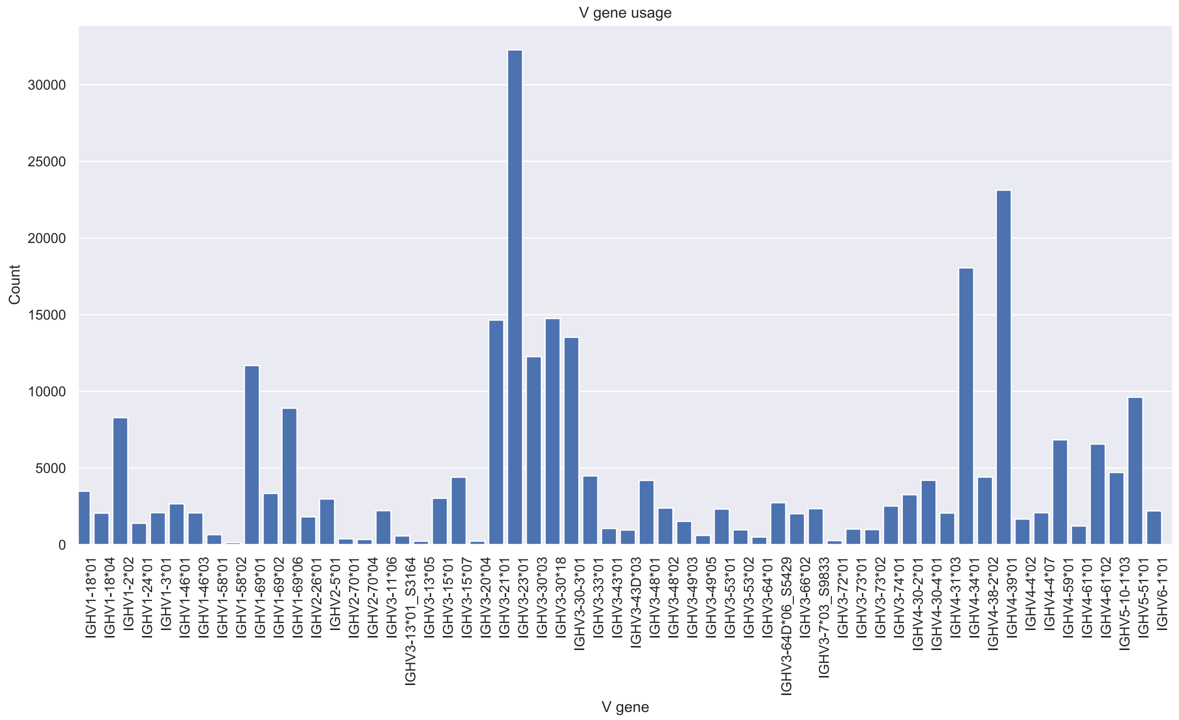

### ERR2567243

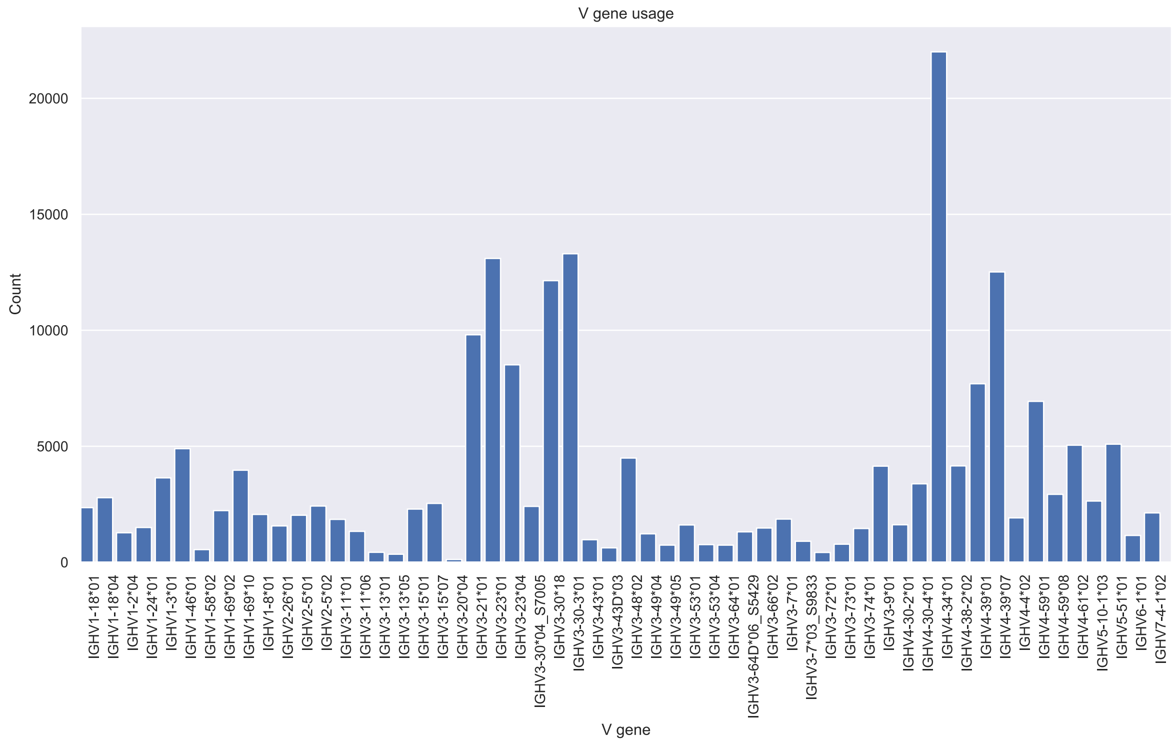
V gene usage
Count
V gene
0
5000
10000
15000
20000
IGHV1-18*01
IGHV1-18*04
IGHV1-2*04
IGHV1-24*01
IGHV1-3*01
IGHV1-46*01
IGHV1-58*02
IGHV1-69*02
IGHV1-69*10
IGHV1-8*01
IGHV2-26*01
IGHV2-5*01
IGHV2-5*02
IGHV3-11*01
IGHV3-11*06
IGHV3-13*01
IGHV3-13*05
IGHV3-15*01
IGHV3-15*07
IGHV3-20*04
IGHV3-21*01
IGHV3-23*01
IGHV3-23*04
IGHV3-30*04_S7005
IGHV3-30*18
IGHV3-30-3*01
IGHV3-43*01
IGHV3-43D*03
IGHV3-48*02
IGHV3-49*04
IGHV3-49*05
IGHV3-53*01
IGHV3-53*04
IGHV3-64*01
IGHV3-64D*06_S5429
IGHV3-66*02
IGHV3-7*01
IGHV3-7*03_S9833
IGHV3-72*01
IGHV3-73*01
IGHV3-74*01
IGHV3-9*01
IGHV4-30-2*01
IGHV4-30-4*01
IGHV4-34*01
IGHV4-38-2*02
IGHV4-39*01
IGHV4-39*07
IGHV4-4*02
IGHV4-59*01
IGHV4-59*08
IGHV4-61*02
IGHV5-10-1*03
IGHV5-51*01
IGHV6-1*01
IGHV7-4-1*02

### ERR2567246

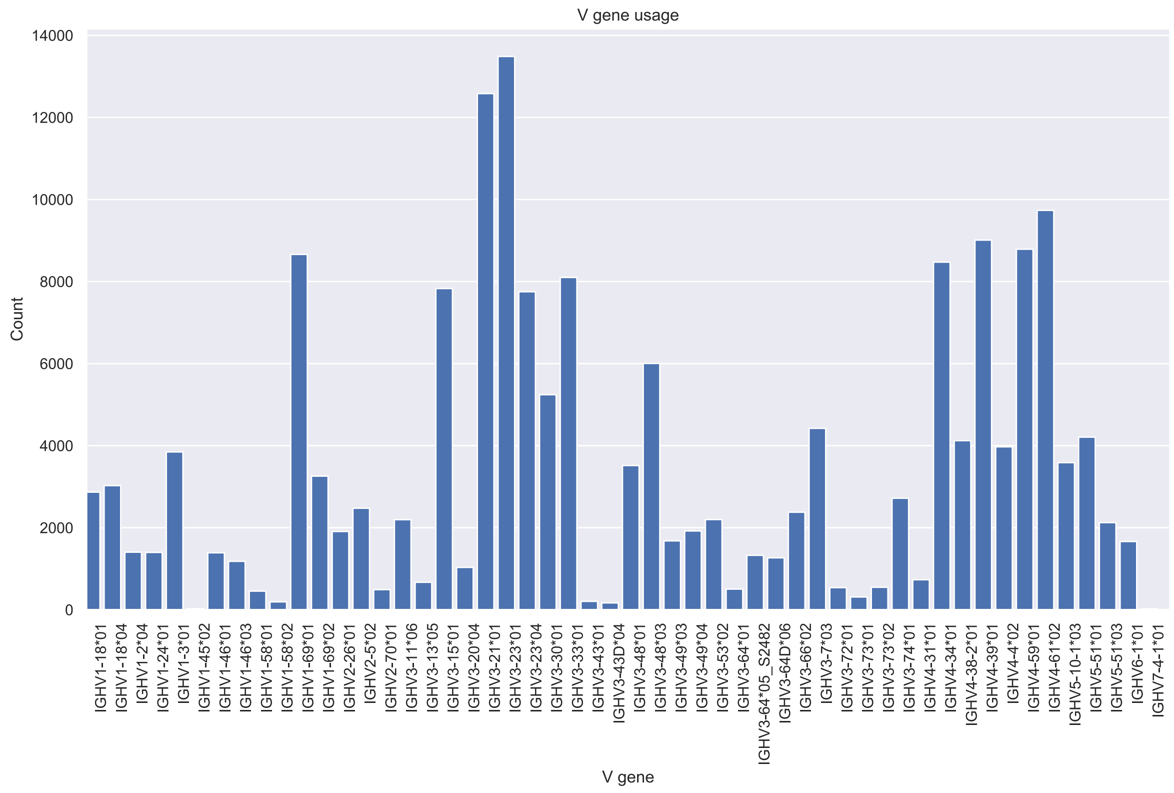

### ERR2567249

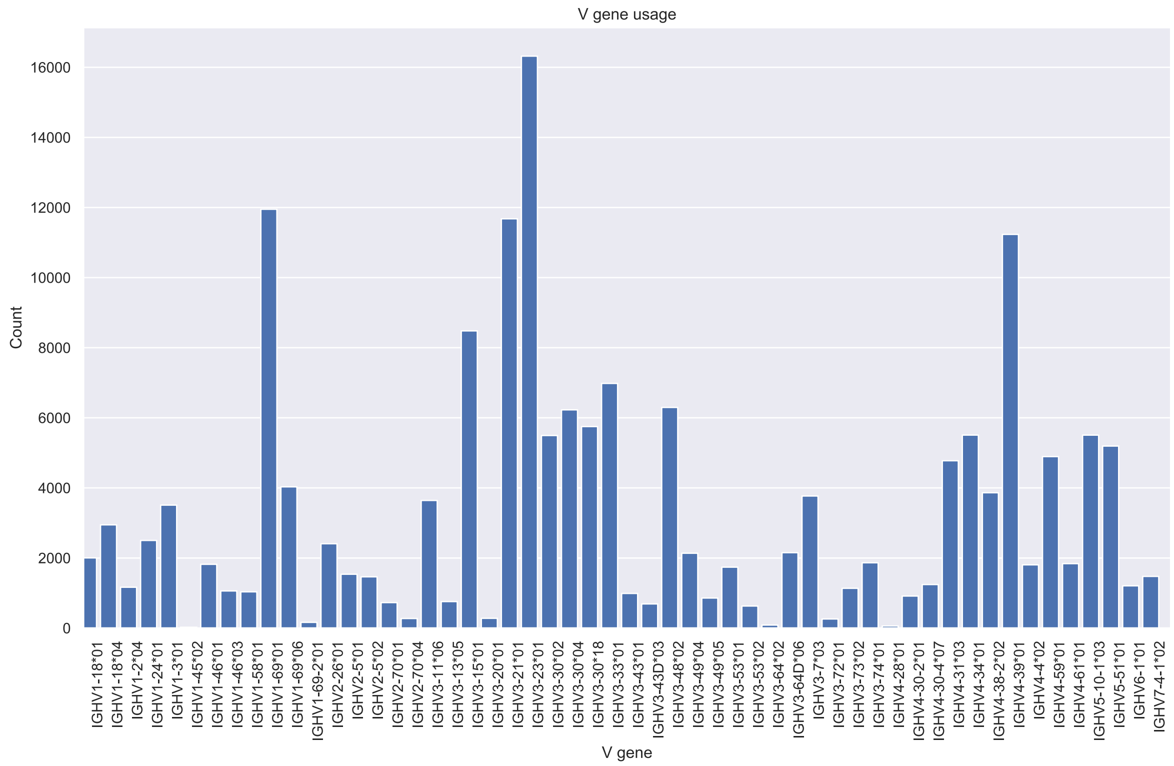
V gene usage
Count
16000
14000
12000
10000
8000
6000
4000
2000
0
IGHV1-18*01
IGHV1-18*04
IGHV1-2*04
IGHV1-24*01
IGHV1-3*01
IGHV1-45*02
IGHV1-46*01
IGHV1-46*03
IGHV1-58*01
IGHV1-69*01
IGHV1-69*06
IGHV1-69-2*01
IGHV2-26*01
IGHV2-5*01
IGHV2-5*02
IGHV2-70*01
IGHV2-70*04
IGHV3-11*06
IGHV3-13*05
IGHV3-15*01
IGHV3-20*01
IGHV3-21*01
IGHV3-23*01
IGHV3-30*02
IGHV3-30*04
IGHV3-30*18
IGHV3-33*01
IGHV3-43*01
IGHV3-43D*03
IGHV3-48*02
IGHV3-49*04
IGHV3-49*05
IGHV3-53*01
IGHV3-53*02
IGHV3-64*02
IGHV3-64D*06
IGHV3-7*03
IGHV3-72*01
IGHV3-73*02
IGHV3-74*01
IGHV4-28*01
IGHV4-30-2*01
IGHV4-30-4*07
IGHV4-31*03
IGHV4-34*01
IGHV4-38-2*02
IGHV4-39*01
IGHV4-4*02
IGHV4-59*01
IGHV4-61*01
IGHV5-10-1*03
IGHV5-51*01
IGHV6-1*01
IGHV7-4-1*02
V gene

### ERR2567254

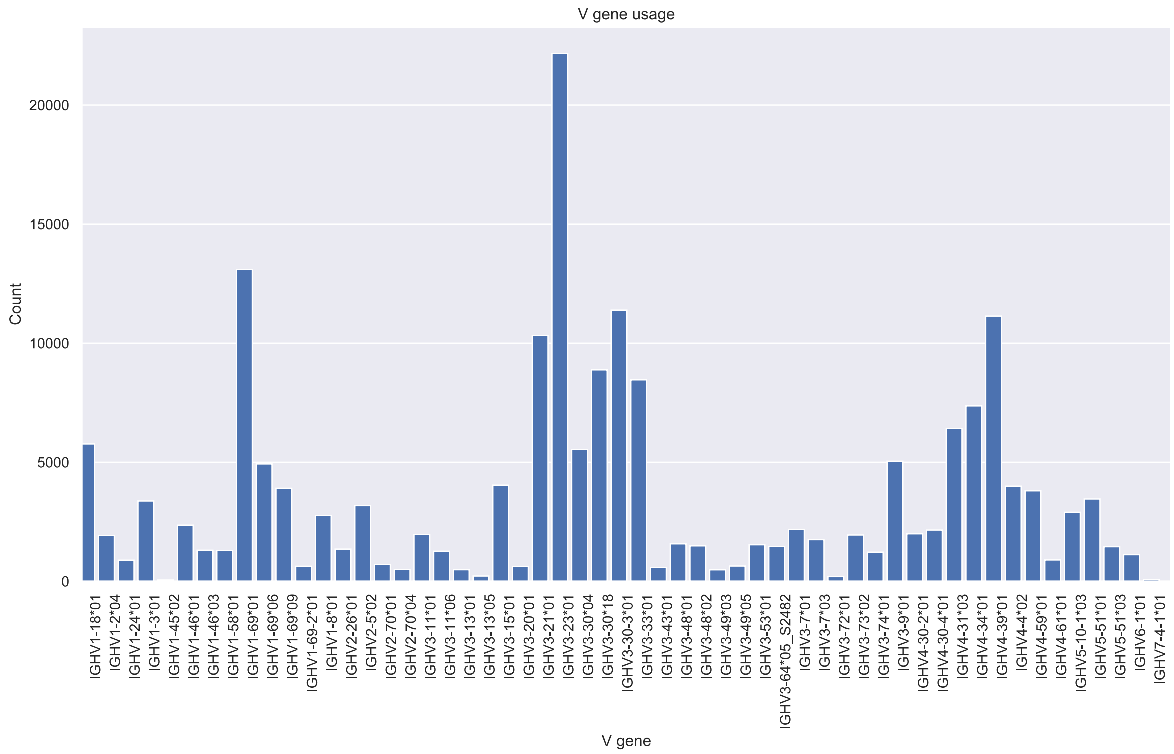

V gene usage
Count
20000
15000
10000
5000
0
V gene
IGHV1-18*01
IGHV1-2*04
IGHV1-24*01
IGHV1-3*01
IGHV1-45*02
IGHV1-46*01
IGHV1-46*03
IGHV1-58*01
IGHV1-69*01
IGHV1-69*06
IGHV1-69*09
IGHV1-69-2*01
IGHV1-8*01
IGHV2-26*01
IGHV2-5*02
IGHV2-70*01
IGHV2-70*04
IGHV3-11*01
IGHV3-11*06
IGHV3-13*01
IGHV3-13*05
IGHV3-15*01
IGHV3-20*01
IGHV3-21*01
IGHV3-23*01
IGHV3-30*04
IGHV3-30*18
IGHV3-30-3*01
IGHV3-33*01
IGHV3-43*01
IGHV3-48*01
IGHV3-48*02
IGHV3-49*03
IGHV3-49*05
IGHV3-53*01
IGHV3-64*05_S2482
IGHV3-7*01
IGHV3-7*03
IGHV3-72*01
IGHV3-73*02
IGHV3-74*01
IGHV3-9*01
IGHV4-30-2*01
IGHV4-30-4*01
IGHV4-31*03
IGHV4-34*01
IGHV4-39*01
IGHV4-4*02
IGHV4-59*01
IGHV4-61*01
IGHV5-10-1*03
IGHV5-51*01
IGHV5-51*03
IGHV6-1*01
IGHV7-4-1*01

### ERR2567259

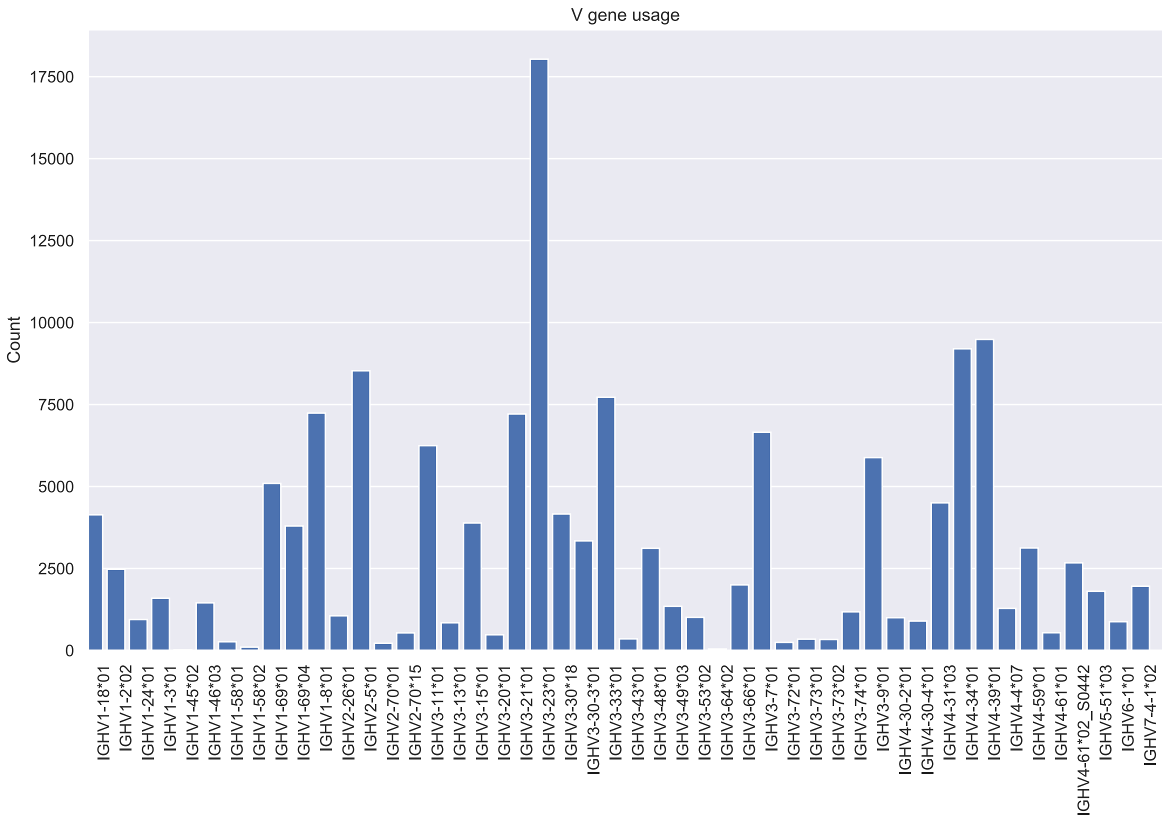

V gene usage
Count
V gene
17500
15000
12500
10000
7500
5000
2500
0
IGHV1-18*01
IGHV1-2*02
IGHV1-24*01
IGHV1-3*01
IGHV1-45*02
IGHV1-46*03
IGHV1-58*01
IGHV1-58*02
IGHV1-69*01
IGHV1-69*04
IGHV1-8*01
IGHV2-26*01
IGHV2-5*01
IGHV2-70*01
IGHV2-70*15
IGHV3-11*01
IGHV3-13*01
IGHV3-15*01
IGHV3-20*01
IGHV3-21*01
IGHV3-23*01
IGHV3-30*18
IGHV3-30-3*01
IGHV3-33*01
IGHV3-43*01
IGHV3-48*01
IGHV3-49*03
IGHV3-53*02
IGHV3-64*02
IGHV3-66*01
IGHV3-7*01
IGHV3-72*01
IGHV3-73*01
IGHV3-73*02
IGHV3-74*01
IGHV3-9*01
IGHV4-30-2*01
IGHV4-30-4*01
IGHV4-31*03
IGHV4-34*01
IGHV4-39*01
IGHV4-4*07
IGHV4-59*01
IGHV4-61*01
IGHV4-61*02_S0442
IGHV5-51*03
IGHV6-1*01
IGHV7-4-1*02

### ERR2567261

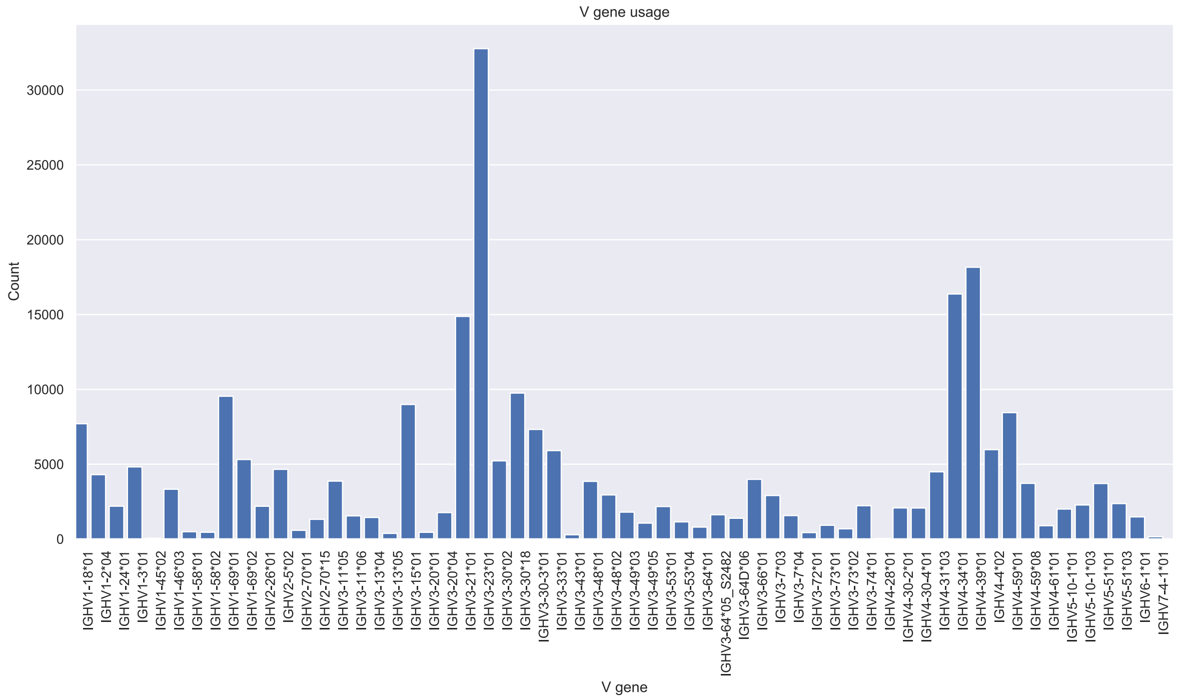
V gene usage
Count
V gene
0
5000
10000
15000
20000
25000
30000

### ERR2567263

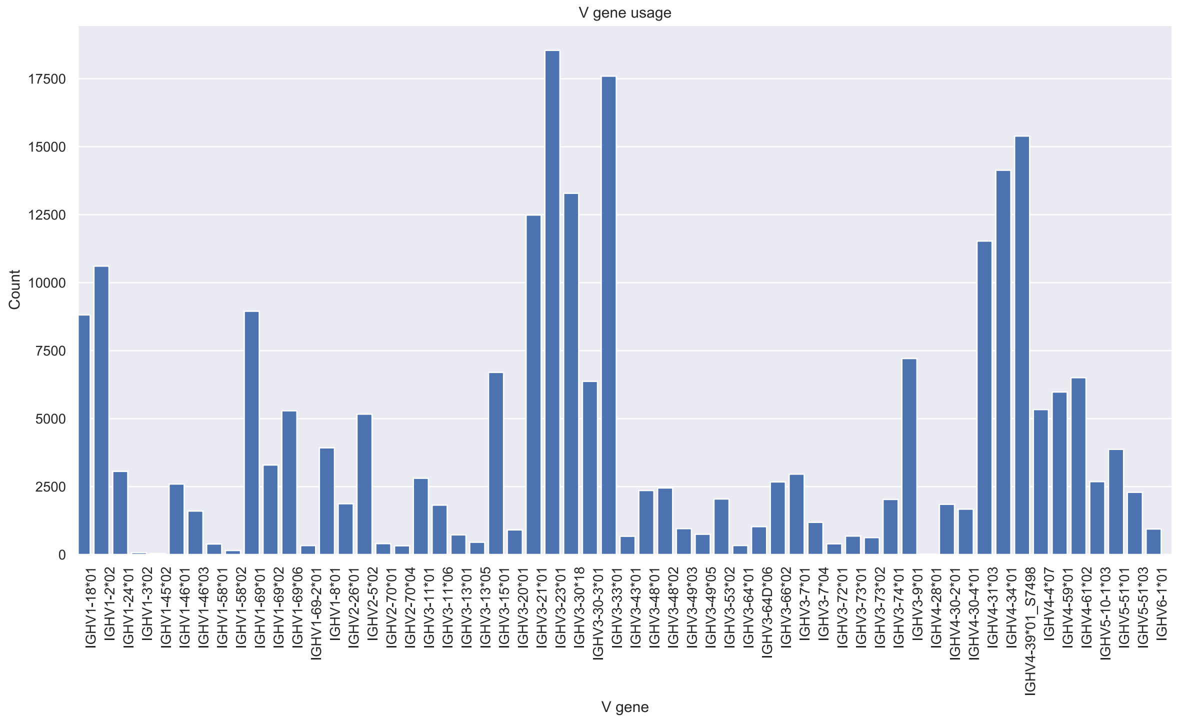
V gene usage
Count
17500
15000
12500
10000
7500
5000
2500
0
IGHV1-18*01
IGHV1-2*02
IGHV1-24*01
IGHV1-3*02
IGHV1-45*02
IGHV1-46*01
IGHV1-46*03
IGHV1-58*01
IGHV1-58*02
IGHV1-69*01
IGHV1-69*02
IGHV1-69*06
IGHV1-69-2*01
IGHV1-8*01
IGHV2-26*01
IGHV2-5*02
IGHV2-70*01
IGHV2-70*04
IGHV3-11*01
IGHV3-11*06
IGHV3-13*01
IGHV3-13*05
IGHV3-15*01
IGHV3-20*01
IGHV3-21*01
IGHV3-23*01
IGHV3-30*18
IGHV3-30-3*01
IGHV3-33*01
IGHV3-43*01
IGHV3-48*01
IGHV3-48*02
IGHV3-49*03
IGHV3-49*05
IGHV3-53*02
IGHV3-64*01
IGHV3-64D*06
IGHV3-66*02
IGHV3-7*01
IGHV3-7*04
IGHV3-72*01
IGHV3-73*01
IGHV3-73*02
IGHV3-74*01
IGHV3-9*01
IGHV4-28*01
IGHV4-30-2*01
IGHV4-30-4*01
IGHV4-31*03
IGHV4-34*01
IGHV4-39*01_S7498
IGHV4-4*07
IGHV4-59*01
IGHV4-61*02
IGHV5-10-1*03
IGHV5-51*01
IGHV5-51*03
IGHV6-1*01
V gene

### ERR2567264

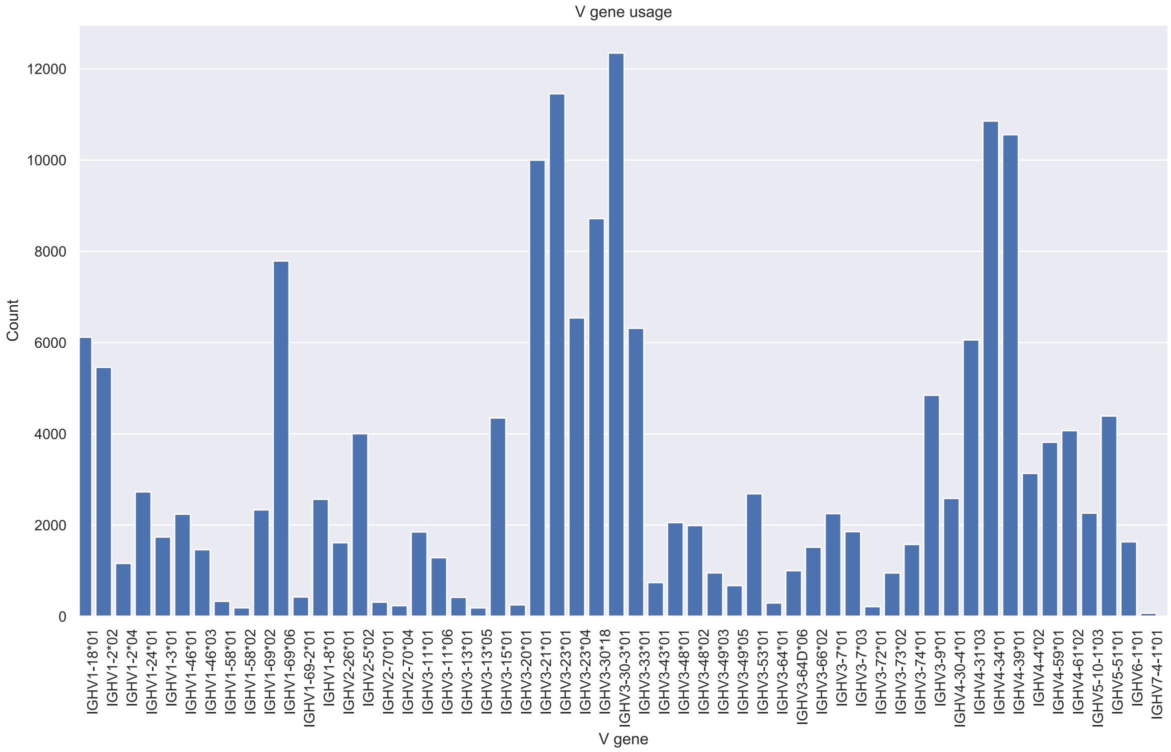
V gene usage
Count
12000
10000
8000
6000
4000
2000
0
IGHV1-18*01
IGHV1-2*02
IGHV1-2*04
IGHV1-24*01
IGHV1-3*01
IGHV1-46*01
IGHV1-46*03
IGHV1-58*01
IGHV1-58*02
IGHV1-69*02
IGHV1-69*06
IGHV1-69-2*01
IGHV1-8*01
IGHV2-26*01
IGHV2-5*02
IGHV2-70*01
IGHV2-70*04
IGHV3-11*01
IGHV3-11*06
IGHV3-13*01
IGHV3-13*05
IGHV3-15*01
IGHV3-20*01
IGHV3-21*01
IGHV3-23*01
IGHV3-23*04
IGHV3-30*18
IGHV3-30-3*01
IGHV3-33*01
IGHV3-43*01
IGHV3-48*01
IGHV3-48*02
IGHV3-49*03
IGHV3-49*05
IGHV3-53*01
IGHV3-64*01
IGHV3-64D*06
IGHV3-66*02
IGHV3-7*01
IGHV3-7*03
IGHV3-72*01
IGHV3-73*02
IGHV3-74*01
IGHV3-9*01
IGHV4-30-4*01
IGHV4-31*03
IGHV4-34*01
IGHV4-39*01
IGHV4-4*02
IGHV4-59*01
IGHV4-61*02
IGHV5-10-1*03
IGHV5-51*01
IGHV6-1*01
IGHV7-4-1*01
V gene

### ERR2567265

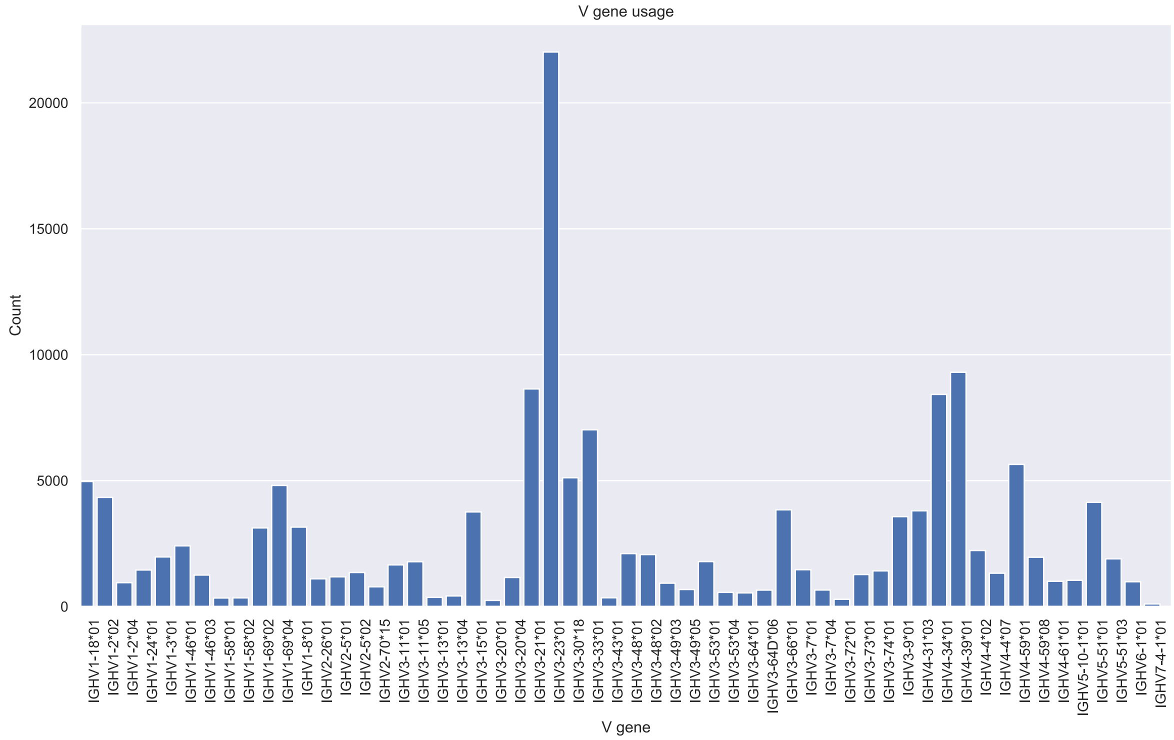
V gene usage
Count
20000
15000
10000
5000
0
V gene
IGHV1-18*01
IGHV1-2*02
IGHV1-2*04
IGHV1-24*01
IGHV1-3*01
IGHV1-46*01
IGHV1-46*03
IGHV1-58*01
IGHV1-58*02
IGHV1-69*02
IGHV1-69*04
IGHV1-8*01
IGHV2-26*01
IGHV2-5*01
IGHV2-5*02
IGHV2-70*15
IGHV3-11*01
IGHV3-11*05
IGHV3-13*01
IGHV3-13*04
IGHV3-15*01
IGHV3-20*01
IGHV3-20*04
IGHV3-21*01
IGHV3-23*01
IGHV3-30*18
IGHV3-33*01
IGHV3-43*01
IGHV3-48*01
IGHV3-48*02
IGHV3-49*03
IGHV3-49*05
IGHV3-53*01
IGHV3-53*04
IGHV3-64*01
IGHV3-64D*06
IGHV3-66*01
IGHV3-7*01
IGHV3-7*04
IGHV3-72*01
IGHV3-73*01
IGHV3-74*01
IGHV3-9*01
IGHV4-31*03
IGHV4-34*01
IGHV4-39*01
IGHV4-4*02
IGHV4-4*07
IGHV4-59*01
IGHV4-59*08
IGHV4-61*01
IGHV5-10-1*01
IGHV5-51*01
IGHV5-51*03
IGHV6-1*01
IGHV7-4-1*01

### ERR2567266

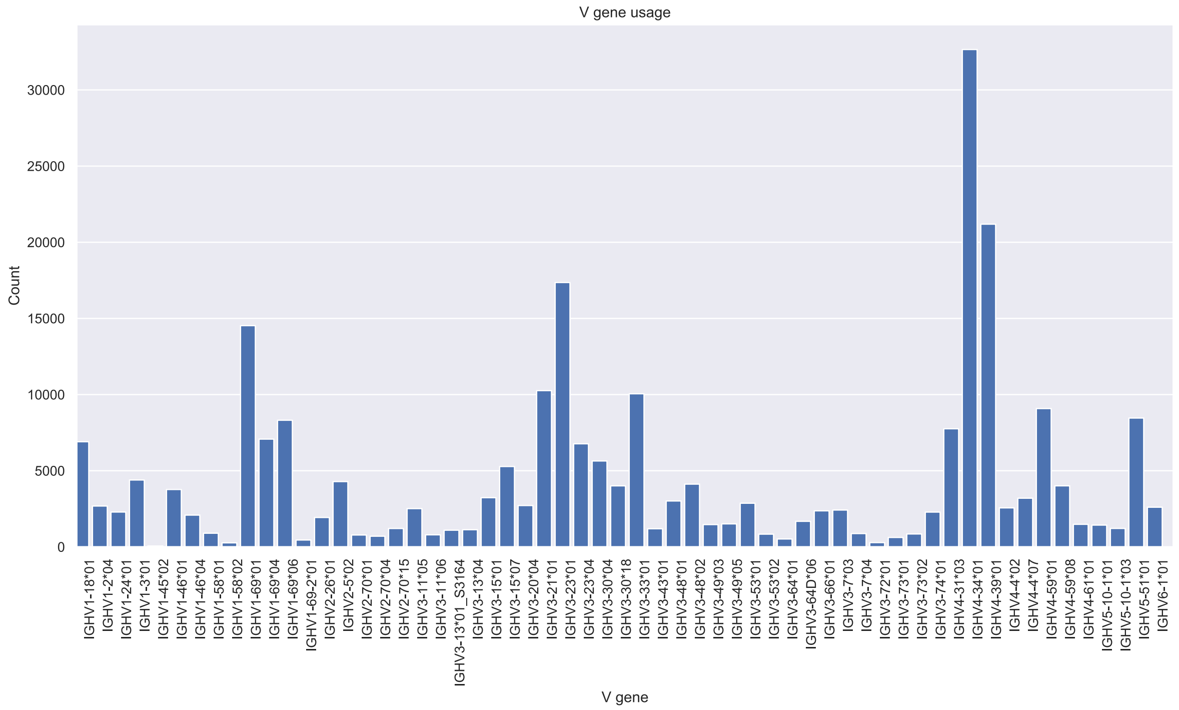
V gene usage
Count
V gene
IGHV1-18*01
IGHV1-2*04
IGHV1-24*01
IGHV1-3*01
IGHV1-45*02
IGHV1-46*01
IGHV1-46*04
IGHV1-58*01
IGHV1-58*02
IGHV1-69*01
IGHV1-69*04
IGHV1-69*06
IGHV1-69-2*01
IGHV2-26*01
IGHV2-5*02
IGHV2-70*01
IGHV2-70*04
IGHV2-70*15
IGHV3-11*05
IGHV3-11*06
IGHV3-13*01_S3164
IGHV3-13*04
IGHV3-15*01
IGHV3-15*07
IGHV3-20*04
IGHV3-21*01
IGHV3-23*01
IGHV3-23*04
IGHV3-30*04
IGHV3-30*18
IGHV3-33*01
IGHV3-43*01
IGHV3-48*01
IGHV3-48*02
IGHV3-49*03
IGHV3-49*05
IGHV3-53*01
IGHV3-53*02
IGHV3-64*01
IGHV3-64D*06
IGHV3-66*01
IGHV3-7*03
IGHV3-7*04
IGHV3-72*01
IGHV3-73*01
IGHV3-73*02
IGHV3-74*01
IGHV4-31*03
IGHV4-34*01
IGHV4-39*01
IGHV4-4*02
IGHV4-4*07
IGHV4-59*01
IGHV4-59*08
IGHV4-61*01
IGHV5-10-1*01
IGHV5-10-1*03
IGHV5-51*01
IGHV6-1*01
0
5000
10000
15000
20000
25000
30000

### ERR2567271

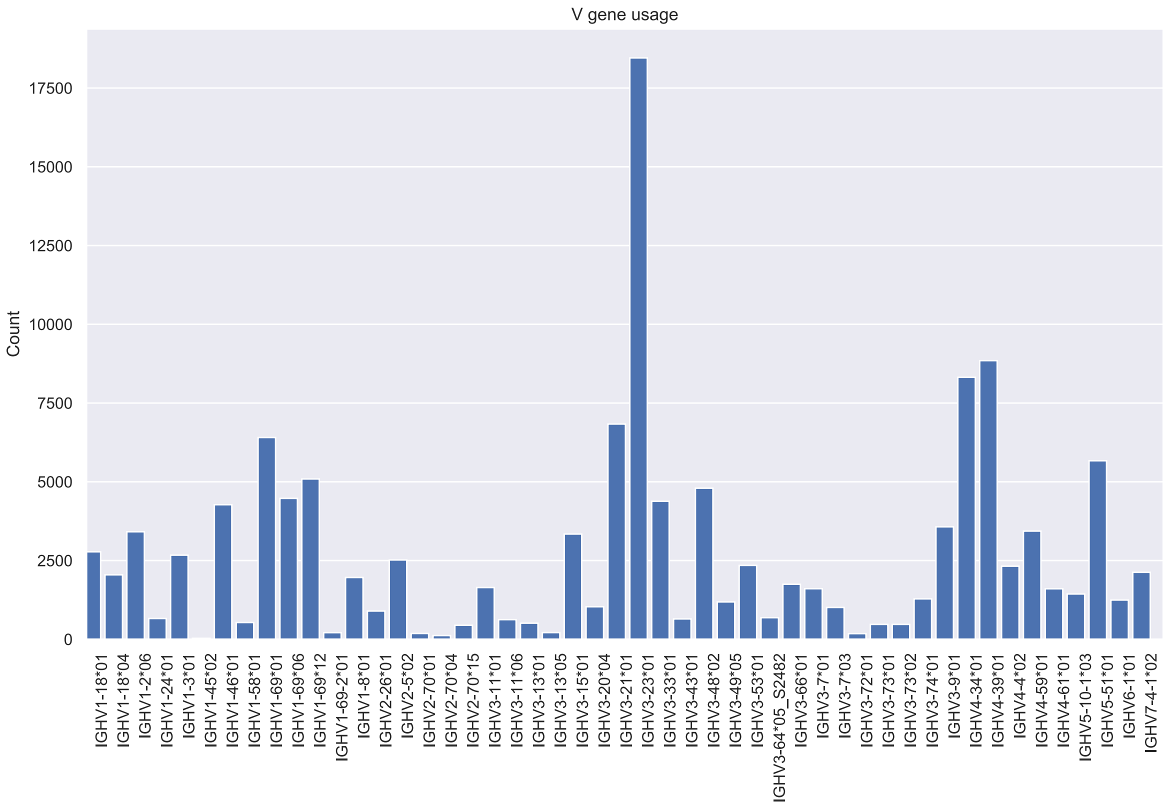

V gene usage
Count
0
2500
5000
7500
10000
12500
15000
17500
IGHV1-18*01
IGHV1-18*04
IGHV1-2*06
IGHV1-24*01
IGHV1-3*01
IGHV1-45*02
IGHV1-46*01
IGHV1-58*01
IGHV1-69*01
IGHV1-69*06
IGHV1-69*12
IGHV1-69-2*01
IGHV1-8*01
IGHV2-26*01
IGHV2-5*02
IGHV2-70*01
IGHV2-70*04
IGHV2-70*15
IGHV3-11*01
IGHV3-11*06
IGHV3-13*01
IGHV3-13*05
IGHV3-15*01
IGHV3-20*04
IGHV3-21*01
IGHV3-23*01
IGHV3-33*01
IGHV3-43*01
IGHV3-48*02
IGHV3-49*05
IGHV3-53*01
IGHV3-64*05_S2482
IGHV3-66*01
IGHV3-7*01
IGHV3-7*03
IGHV3-72*01
IGHV3-73*01
IGHV3-73*02
IGHV3-74*01
IGHV3-9*01
IGHV4-34*01
IGHV4-39*01
IGHV4-4*02
IGHV4-59*01
IGHV4-61*01
IGHV5-10-1*03
IGHV5-51*01
IGHV6-1*01
IGHV7-4-1*02
V gene

### ERR2567274

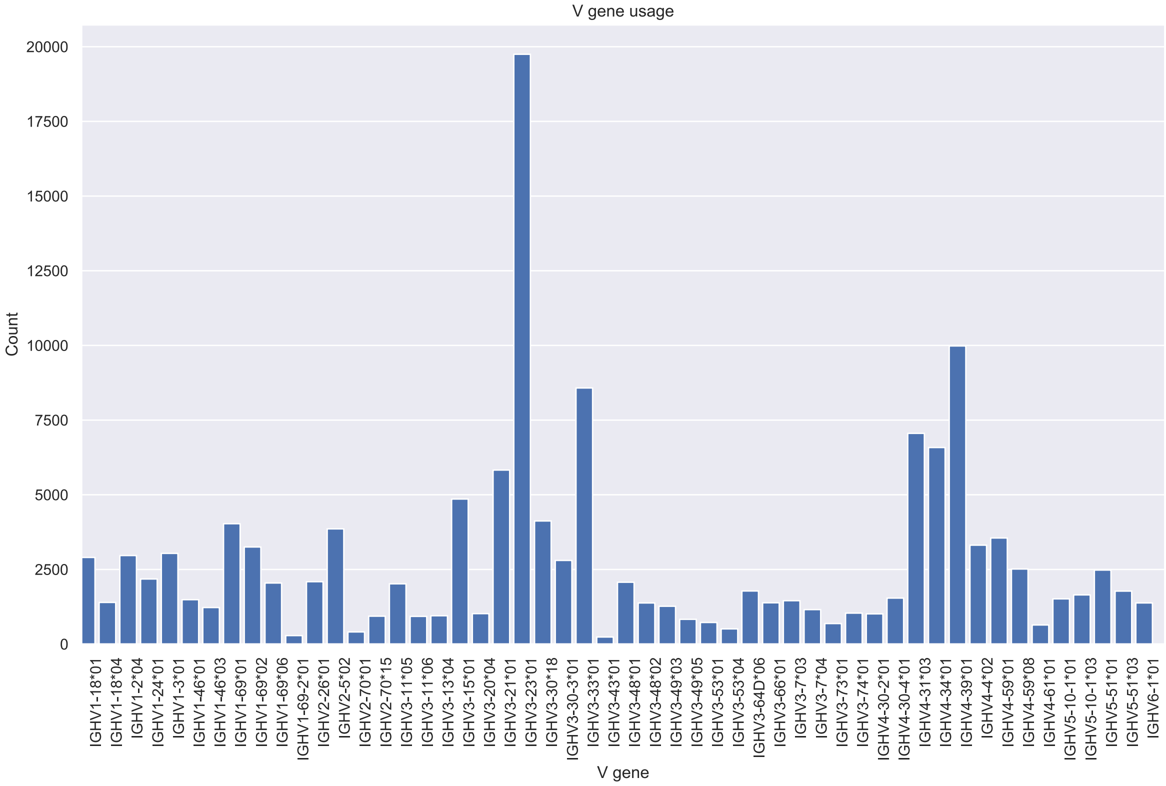

### ERR2567276

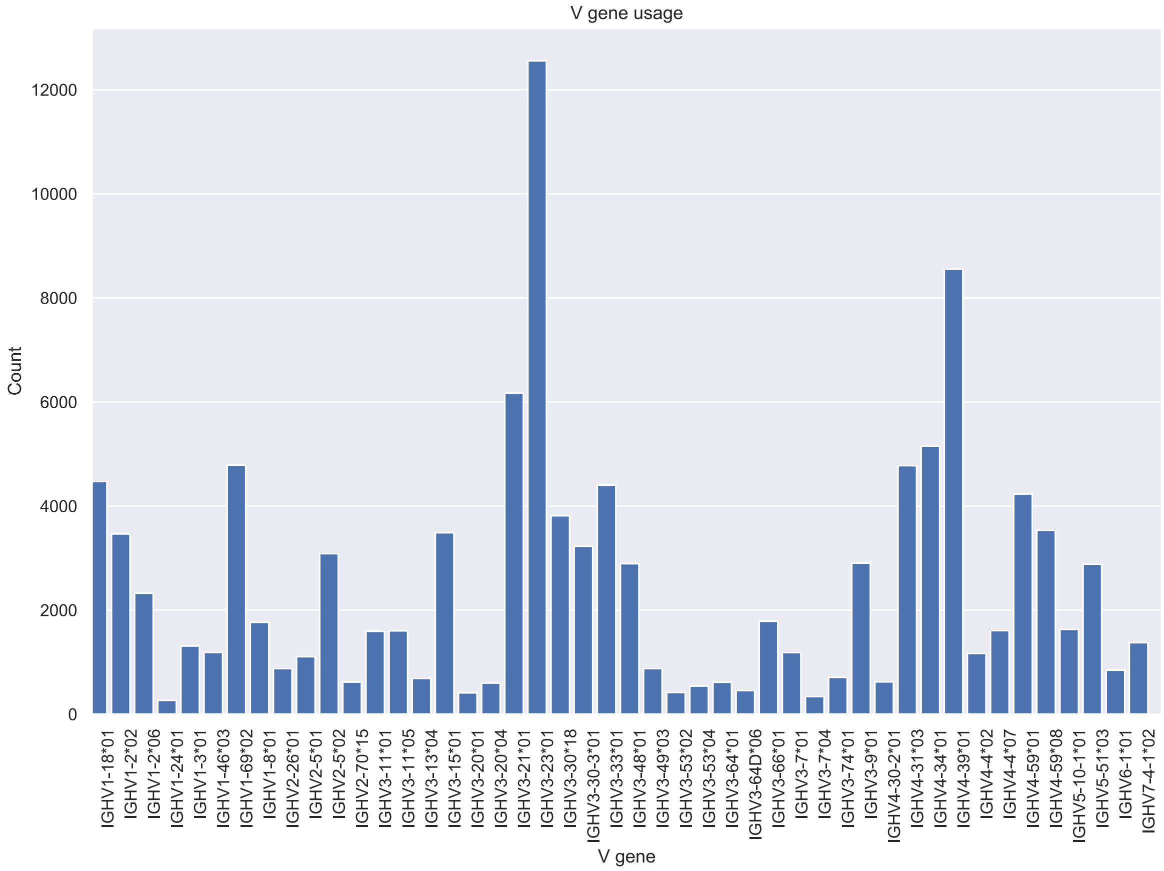

V gene usage
Count
V gene
12000
10000
8000
6000
4000
2000
0
IGHV1-18*01
IGHV1-2*02
IGHV1-2*06
IGHV1-24*01
IGHV1-3*01
IGHV1-46*03
IGHV1-69*02
IGHV1-8*01
IGHV2-26*01
IGHV2-5*01
IGHV2-5*02
IGHV2-70*15
IGHV3-11*01
IGHV3-11*05
IGHV3-13*04
IGHV3-15*01
IGHV3-20*01
IGHV3-20*04
IGHV3-21*01
IGHV3-23*01
IGHV3-30*18
IGHV3-30-3*01
IGHV3-33*01
IGHV3-48*01
IGHV3-49*03
IGHV3-53*02
IGHV3-53*04
IGHV3-64*01
IGHV3-64D*06
IGHV3-66*01
IGHV3-7*01
IGHV3-7*04
IGHV3-74*01
IGHV3-9*01
IGHV4-30-2*01
IGHV4-31*03
IGHV4-34*01
IGHV4-39*01
IGHV4-4*02
IGHV4-4*07
IGHV4-59*01
IGHV4-59*08
IGHV5-10-1*01
IGHV5-51*03
IGHV6-1*01
IGHV7-4-1*02

### ERR2567277

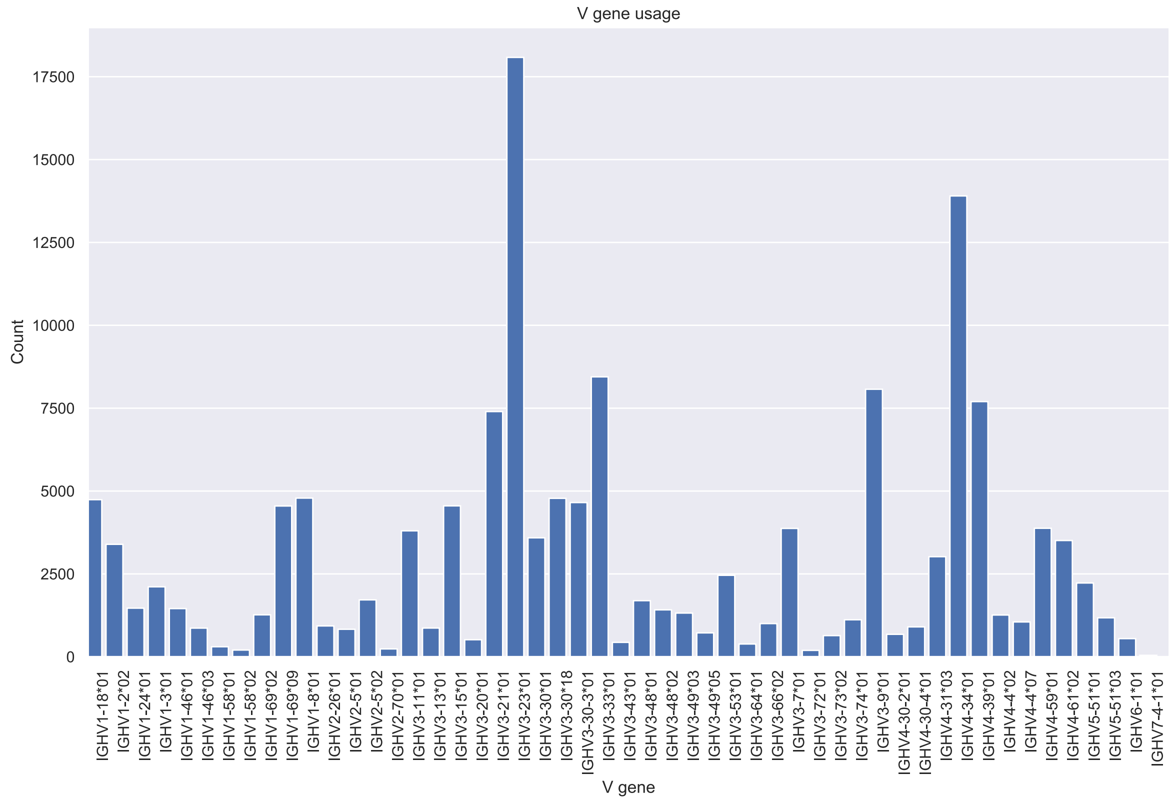
V gene usage
Count
17500
15000
12500
10000
7500
5000
2500
0
V gene
IGHV1-18*01
IGHV1-2*02
IGHV1-24*01
IGHV1-3*01
IGHV1-46*01
IGHV1-46*03
IGHV1-58*01
IGHV1-58*02
IGHV1-69*02
IGHV1-69*09
IGHV1-8*01
IGHV2-26*01
IGHV2-5*01
IGHV2-5*02
IGHV2-70*01
IGHV3-11*01
IGHV3-13*01
IGHV3-15*01
IGHV3-20*01
IGHV3-21*01
IGHV3-23*01
IGHV3-30*01
IGHV3-30*18
IGHV3-30-3*01
IGHV3-33*01
IGHV3-43*01
IGHV3-48*01
IGHV3-48*02
IGHV3-49*03
IGHV3-49*05
IGHV3-53*01
IGHV3-64*01
IGHV3-66*02
IGHV3-7*01
IGHV3-72*01
IGHV3-73*02
IGHV3-74*01
IGHV3-9*01
IGHV4-30-2*01
IGHV4-30-4*01
IGHV4-31*03
IGHV4-34*01
IGHV4-39*01
IGHV4-4*02
IGHV4-4*07
IGHV4-59*01
IGHV4-61*02
IGHV5-51*01
IGHV5-51*03
IGHV6-1*01
IGHV7-4-1*01
