## Supplementary Figures 3-8 for "Poorly expressed alleles of several human immunoglobulin heavy chain variable (IGHV) genes are common in the human population"

**Supplementary Figure 3.** Allelic variants of IGHV1-2 as defined by IMGT are illustrated. Variability of some of the positions of these genes in samples obtained in different geographical locations as illustrated by the ENSEMBL browser (release 101, August 2020) (19) is shown. Only bases 163, 223, and 299 (IMGT numbering nomenclature (16)) of this gene display frequencies of variation >1% in the 1000 Genomes Project. The variant (SNP rs12588974) at base 299, indicative of the IGHV1-2*01 or IGHV1-2*05 alleles is present at about 5% in European populations. Bases 233 and 234 (SNPs rs782139757 and rs1425538657), that separate these two alleles remains as T and G, respectively, at very high frequency in most populations suggesting that IGHV1-2*01 is not common in these populations (not shown). All sequence variants of the illustrations of SNPs are indicated as seen in the reversed strand, hence they are complementary to the base of the coding strand.

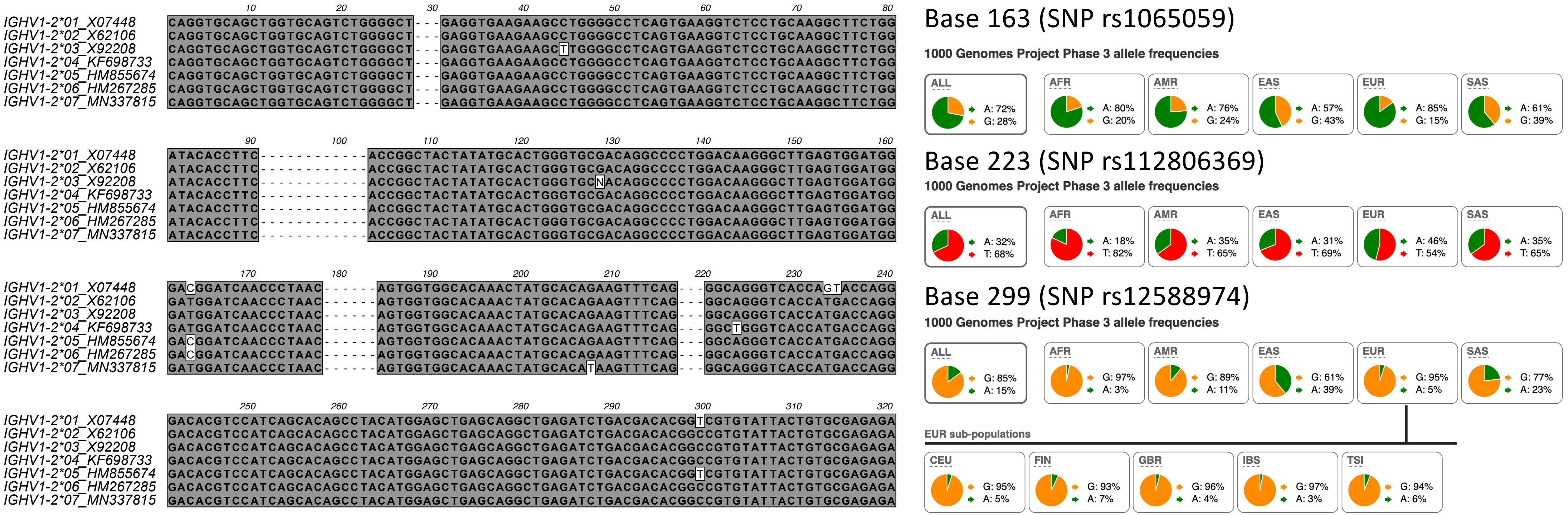

10 20 30 40 50 60 70 80
IGHV1-2*01_X07448 CAGGTGCAGCTGGTGCAGTCTGGGGCT---GAGGTGAAGAAGCCTGGGGCCTCAGTGAAGGTCTCCTGCAAGGCTTCTGG
IGHV1-2*02_X62106 CAGGTGCAGCTGGTGCAGTCTGGGGCT---GAGGTGAAGAAGCCTGGGGCCTCAGTGAAGGTCTCCTGCAAGGCTTCTGG
IGHV1-2*03_X92208 CAGGTGCAGCTGGTGCAGTCTGGGGCT---GAGGTGAAGAAGCTTGGGGCCTCAGTGAAGGTCTCCTGCAAGGCTTCTGG
IGHV1-2*04_KF698733 CAGGTGCAGCTGGTGCAGTCTGGGGCT---GAGGTGAAGAAGCCTGGGGCCTCAGTGAAGGTCTCCTGCAAGGCTTCTGG
IGHV1-2*05_HM855674 CAGGTGCAGCTGGTGCAGTCTGGGGCT---GAGGTGAAGAAGCCTGGGGCCTCAGTGAAGGTCTCCTGCAAGGCTTCTGG
IGHV1-2*06_HM267285 CAGGTGCAGCTGGTGCAGTCTGGGGCT---GAGGTGAAGAAGCCTGGGGCCTCAGTGAAGGTCTCCTGCAAGGCTTCTGG
IGHV1-2*07_MN337815 CAGGTGCAGCTGGTGCAGTCTGGGGCT---GAGGTGAAGAAGCCTGGGGCCTCAGTGAAGGTCTCCTGCAAGGCTTCTGG
90 100 110 120 130 140 150 160
IGHV1-2*01_X07448 ATACACCTTC------------ACCGGCTACTATATGCACTGGGTGCGACAGGCCCCTGGACAAGGGCTTGAGTGGATGG
IGHV1-2*02_X62106 ATACACCTTC------------ACCGGCTACTATATGCACTGGGTGCGACAGGCCCCTGGACAAGGGCTTGAGTGGATGG
IGHV1-2*03_X92208 ATACACCTTC------------ACCGGCTACTATATGCACTGGGTGCNACAGGCCCCTGGACAAGGGCTTGAGTGGATGG
IGHV1-2*04_KF698733 ATACACCTTC------------ACCGGCTACTATATGCACTGGGTGCGACAGGCCCCTGGACAAGGGCTTGAGTGGATGG
IGHV1-2*05_HM855674 ATACACCTTC------------ACCGGCTACTATATGCACTGGGTGCGACAGGCCCCTGGACAAGGGCTTGAGTGGATGG
IGHV1-2*06_HM267285 ATACACCTTC------------ACCGGCTACTATATGCACTGGGTGCGACAGGCCCCTGGACAAGGGCTTGAGTGGATGG
IGHV1-2*07_MN337815 ATACACCTTC------------ACCGGCTACTATATGCACTGGGTGCGACAGGCCCCTGGACAAGGGCTTGAGTGGATGG
170 180 190 200 210 220 230 240
IGHV1-2*01_X07448 GACGGATCAACCCTAAC------AGTGGTGGCACAAACTATGCACAGAAGTTTCAG---GGCAGGGTCACCAGTACCAGG
IGHV1-2*02_X62106 GATGGATCAACCCTAAC------AGTGGTGGCACAAACTATGCACAGAAGTTTCAG---GGCAGGGTCACCATGACCAGG
IGHV1-2*03_X92208 GATGGATCAACCCTAAC------AGTGGTGGCACAAACTATGCACAGAAGTTTCAG---GGCAGGGTCACCATGACCAGG
IGHV1-2*04_KF698733 GATGGATCAACCCTAAC------AGTGGTGGCACAAACTATGCACAGAAGTTTCAG---GGCTGGGTCACCATGACCAGG
IGHV1-2*05_HM855674 GACGGATCAACCCTAAC------AGTGGTGGCACAAACTATGCACAGAAGTTTCAG---GGCAGGGTCACCATGACCAGG
IGHV1-2*06_HM267285 GACGGATCAACCCTAAC------AGTGGTGGCACAAACTATGCACAGAAGTTTCAG---GGCAGGGTCACCATGACCAGG
IGHV1-2*07_MN337815 GATGGATCAACCCTAAC------AGTGGTGGCACAAACTATGCACATAAGTTTCAG---GGCAGGGTCACCATGACCAGG
250 260 270 280 290 300 310 320
IGHV1-2*01_X07448 GACACGTCCATCAGCACAGCCTACATGGAGCTGAGCAGGCTGAGATCTGACGACACGGTCGTGTATTACTGTGCGAGAGA
IGHV1-2*02_X62106 GACACGTCCATCAGCACAGCCTACATGGAGCTGAGCAGGCTGAGATCTGACGACACGGCCGTGTATTACTGTGCGAGAGA
IGHV1-2*03_X92208 GACACGTCCATCAGCACAGCCTACATGGAGCTGAGCAGGCTGAGATCTGACGACACGGCCGTGTATTACTGTGCGAGAGA
IGHV1-2*04_KF698733 GACACGTCCATCAGCACAGCCTACATGGAGCTGAGCAGGCTGAGATCTGACGACACGGCCGTGTATTACTGTGCGAGAGA
IGHV1-2*05_HM855674 GACACGTCCATCAGCACAGCCTACATGGAGCTGAGCAGGCTGAGATCTGACGACACGGTCGTGTATTACTGTGCGAGAGA
IGHV1-2*06_HM267285 GACACGTCCATCAGCACAGCCTACATGGAGCTGAGCAGGCTGAGATCTGACGACACGGCCGTGTATTACTGTGCGAGAGA
IGHV1-2*07_MN337815 GACACGTCCATCAGCACAGCCTACATGGAGCTGAGCAGGCTGAGATCTGACGACACGGCCGTGTATTACTGTGCGAGAGA
Base 163 (SNP rs1065059)
1000 Genomes Project Phase 3 allele frequencies
ALL A: 72% G: 28%
AFR A: 80% G: 20%
AMR A: 76% G: 24%
EAS A: 57% G: 43%
EUR A: 85% G: 15%
SAS A: 61% G: 39%
Base 223 (SNP rs112806369)
1000 Genomes Project Phase 3 allele frequencies
ALL A: 32% T: 68%
AFR A: 18% T: 82%
AMR A: 35% T: 65%
EAS A: 31% T: 69%
EUR A: 46% T: 54%
SAS A: 35% T: 65%
Base 299 (SNP rs12588974)
1000 Genomes Project Phase 3 allele frequencies
ALL G: 85% A: 15%
AFR G: 97% A: 3%
AMR G: 89% A: 11%
EAS G: 61% A: 39%
EUR G: 95% A: 5%
SAS G: 77% A: 23%
EUR sub-populations
CEU G: 95% A: 5%
FIN G: 93% A: 7%
GBR G: 96% A: 4%
IBS G: 97% A: 3%
TSI G: 94% A: 6%

**Supplementary Figure 4.** Allelic variants of IGHV1-3 as defined by IMGT are illustrated. Variability of some of the positions of these genes in samples obtained in different geographical locations as illustrated by the ENSEMBL browser (release 101, August 2020) (19) is shown. Only bases 6, 12, 167, 208, 291 and 296 (IMGT numbering nomenclature (16)) of this gene display frequencies of variation >1% in the 1000 Genomes Project. Variants indicative of the IGHV1-3*02 allele are present at about 40%. All sequence variants of the illustrations of SNPs are indicated as seen in the reversed strand, hence they are complementary to the base of the coding strand.

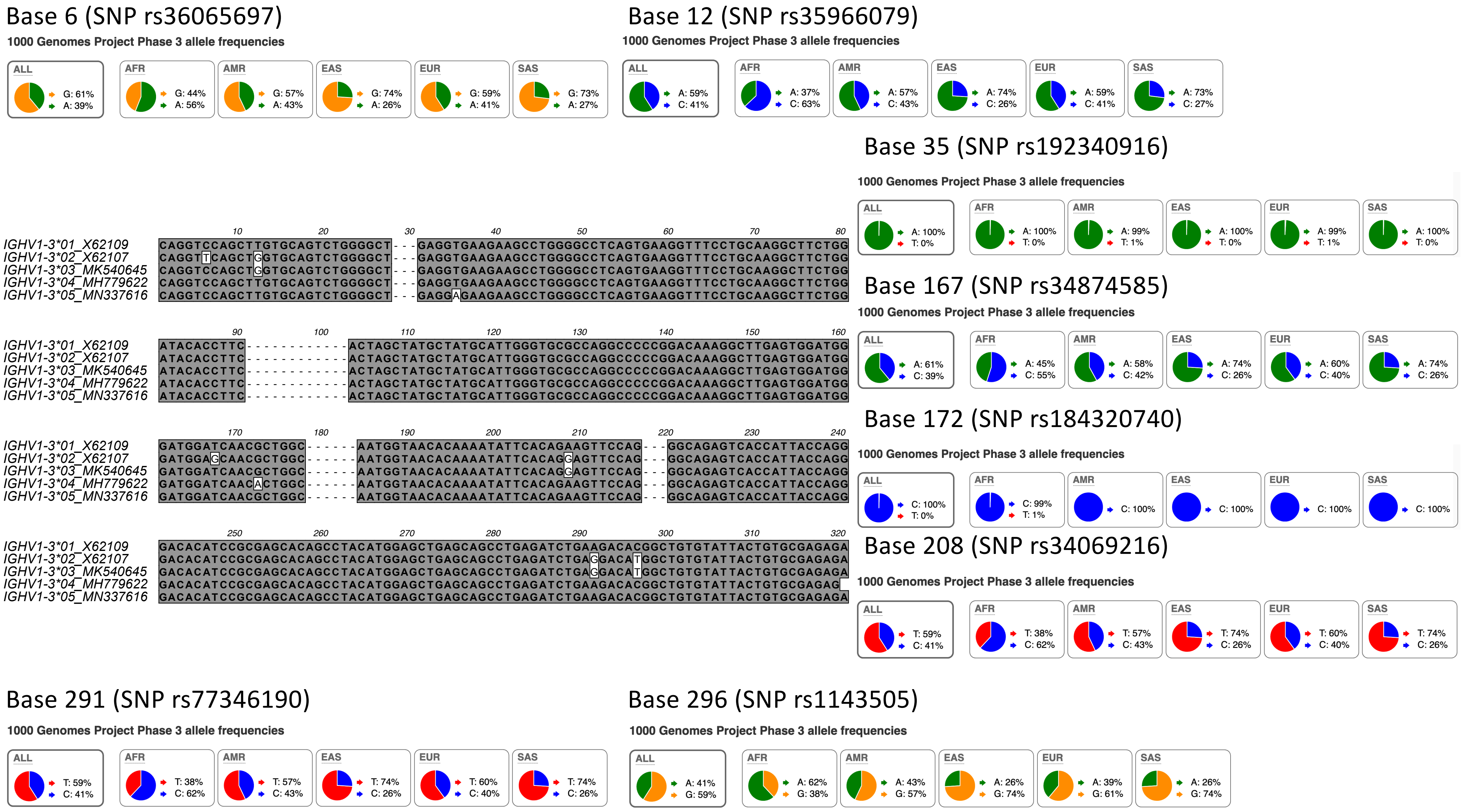

Base 6 (SNP rs36065697)
1000 Genomes Project Phase 3 allele frequencies
ALL G: 61% A: 39%
AFR G: 44% A: 56%
AMR G: 57% A: 43%
EAS G: 74% A: 26%
EUR G: 59% A: 41%
SAS G: 73% A: 27%
Base 12 (SNP rs35966079)
1000 Genomes Project Phase 3 allele frequencies
ALL A: 59% C: 41%
AFR A: 37% C: 63%
AMR A: 57% C: 43%
EAS A: 74% C: 26%
EUR A: 59% C: 41%
SAS A: 73% C: 27%
Base 35 (SNP rs192340916)
1000 Genomes Project Phase 3 allele frequencies
ALL A: 100% T: 0%
AFR A: 100% T: 0%
AMR A: 99% T: 1%
EAS A: 100% T: 0%
EUR A: 99% T: 1%
SAS A: 100% T: 0%
10 20 30 40 50 60 70 80
IGHV1-3*01_X62109 CAGGTCCAGCTTGTGCAGTCTGGGGCT---GAGGTGAAGAAGCCTGGGGCCTCAGTGAAGGTTTCCTGCAAGGCTTCTGG
IGHV1-3*02_X62107 CAGGTTCAGCTGGTGCAGTCTGGGGCT---GAGGTGAAGAAGCCTGGGGCCTCAGTGAAGGTTTCCTGCAAGGCTTCTGG
IGHV1-3*03_MK540645 CAGGTCCAGCTGGTGCAGTCTGGGGCT---GAGGTGAAGAAGCCTGGGGCCTCAGTGAAGGTTTCCTGCAAGGCTTCTGG
IGHV1-3*04_MH779622 CAGGTCCAGCTTGTGCAGTCTGGGGCT---GAGGTGAAGAAGCCTGGGGCCTCAGTGAAGGTTTCCTGCAAGGCTTCTGG
IGHV1-3*05_MN337616 CAGGTCCAGCTTGTGCAGTCTGGGGCT---GAGGAGAAGAAGCCTGGGGCCTCAGTGAAGGTTTCCTGCAAGGCTTCTGG
Base 167 (SNP rs34874585)
1000 Genomes Project Phase 3 allele frequencies
ALL A: 61% C: 39%
AFR A: 45% C: 55%
AMR A: 58% C: 42%
EAS A: 74% C: 26%
EUR A: 60% C: 40%
SAS A: 74% C: 26%
90 100 110 120 130 140 150 160
IGHV1-3*01_X62109 ATACACCTTC------------ACTAGCTATGCTATGCATTGGGTGCGCCAGGCCCCCGGACAAAGGCTTGAGTGGATGG
IGHV1-3*02_X62107 ATACACCTTC------------ACTAGCTATGCTATGCATTGGGTGCGCCAGGCCCCCGGACAAAGGCTTGAGTGGATGG
IGHV1-3*03_MK540645 ATACACCTTC------------ACTAGCTATGCTATGCATTGGGTGCGCCAGGCCCCCGGACAAAGGCTTGAGTGGATGG
IGHV1-3*04_MH779622 ATACACCTTC------------ACTAGCTATGCTATGCATTGGGTGCGCCAGGCCCCCGGACAAAGGCTTGAGTGGATGG
IGHV1-3*05_MN337616 ATACACCTTC------------ACTAGCTATGCTATGCATTGGGTGCGCCAGGCCCCCGGACAAAGGCTTGAGTGGATGG
Base 172 (SNP rs184320740)
1000 Genomes Project Phase 3 allele frequencies
ALL C: 100% T: 0%
AFR C: 99% T: 1%
AMR C: 100%
EAS C: 100%
EUR C: 100%
SAS C: 100%
170 180 190 200 210 220 230 240
IGHV1-3*01_X62109 GATGGATCAACGCTGGC------AATGGTAACACAAAATATTCACAGAAGTTCCAG---GGCAGAGTCACCATTACCAGG
IGHV1-3*02_X62107 GATGGAGCAACGCTGGC------AATGGTAACACAAAATATTCACAGGGTTCCAG---GGCAGAGTCACCATTACCAGG
IGHV1-3*03_MK540645 GATGGATCAACGCTGGC------AATGGTAACACAAAATATTCACAGGGTTCCAG---GGCAGAGTCACCATTACCAGG
IGHV1-3*04_MH779622 GATGGATCAACACTGGC------AATGGTAACACAAAATATTCACAGAAGTTCCAG---GGCAGAGTCACCATTACCAGG
IGHV1-3*05_MN337616 GATGGATCAACGCTGGC------AATGGTAACACAAAATATTCACAGAAGTTCCAG---GGCAGAGTCACCATTACCAGG
Base 208 (SNP rs34069216)
1000 Genomes Project Phase 3 allele frequencies
ALL T: 59% C: 41%
AFR T: 38% C: 62%
AMR T: 57% C: 43%
EAS T: 74% C: 26%
EUR T: 60% C: 40%
SAS T: 74% C: 26%
250 260 270 280 290 300 310 320
IGHV1-3*01_X62109 GACACATCCGCGAGCACAGCCTACATGGAGCTGAGCAGCCTGAGATCTGAAGACACGGCTGTGTATTACTGTGCGAGAGA
IGHV1-3*02_X62107 GACACATCCGCGAGCACAGCCTACATGGAGCTGAGCAGCCTGAGATCTGAGGACATGGCTGTGTATTACTGTGCGAGAGA
IGHV1-3*03_MK540645 GACACATCCGCGAGCACAGCCTACATGGAGCTGAGCAGCCTGAGATCTGAGGACATGGCTGTGTATTACTGTGCGAGAGA
IGHV1-3*04_MH779622 GACACATCCGCGAGCACAGCCTACATGGAGCTGAGCAGCCTGAGATCTGAAGACACGGCTGTGTATTACTGTGCGAGAG
IGHV1-3*05_MN337616 GACACATCCGCGAGCACAGCCTACATGGAGCTGAGCAGCCTGAGATCTGAAGACACGGCTGTGTATTACTGTGCGAGAGA
Base 291 (SNP rs77346190)
1000 Genomes Project Phase 3 allele frequencies
ALL T: 59% C: 41%
AFR T: 38% C: 62%
AMR T: 57% C: 43%
EAS T: 74% C: 26%
EUR T: 60% C: 40%
SAS T: 74% C: 26%
Base 296 (SNP rs1143505)
1000 Genomes Project Phase 3 allele frequencies
ALL A: 41% G: 59%
AFR A: 62% G: 38%
AMR A: 43% G: 57%
EAS A: 26% G: 74%
EUR A: 39% G: 61%
SAS A: 26% G: 74%

**Supplementary Figure 5.** Allelic variants of IGHV4-4 as defined by IMGT are illustrated. Variability of some of the positions of these genes in samples obtained in different geographical locations as illustrated by the ENSEMBL browser (release 101, August 2020) (19) is shown. Analysis of this gene is complicated by extensive similarity with alleles of IGHV4-59 and IGHV4-61, alleles of which are also shown. A few of the positions of IGHV4-4 that display frequencies of variation >1% in all populations in the 1000 Genomes Project are shown. Note that variants at bases 46 and 308 (IMGT numbering nomenclature (16)), indicative of the IGHV4-4*01 allele are present at about 3-4% in European populations. All sequence variants of the illustrations of SNPs are indicated as seen in the reversed strand, hence they are complementary to the base of the coding strand.

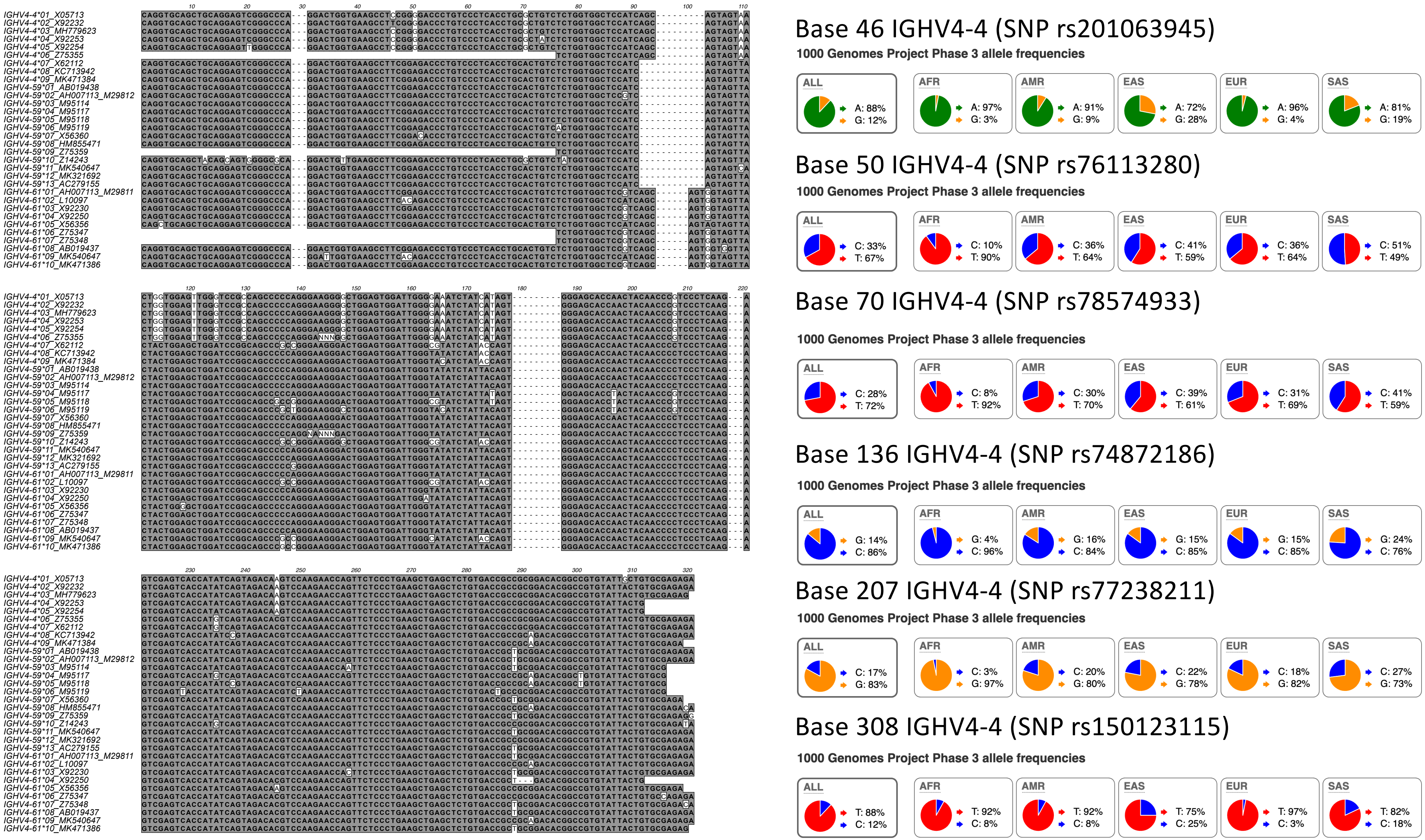
Base 46 IGHV4-4 (SNP rs201063945)
1000 Genomes Project Phase 3 allele frequencies
ALL A: 88% G: 12%
AFR A: 97% G: 3%
AMR A: 91% G: 9%
EAS A: 72% G: 28%
EUR A: 96% G: 4%
SAS A: 81% G: 19%
Base 50 IGHV4-4 (SNP rs76113280)
1000 Genomes Project Phase 3 allele frequencies
ALL C: 33% T: 67%
AFR C: 10% T: 90%
AMR C: 36% T: 64%
EAS C: 41% T: 59%
EUR C: 36% T: 64%
SAS C: 51% T: 49%
Base 70 IGHV4-4 (SNP rs78574933)
1000 Genomes Project Phase 3 allele frequencies
ALL C: 28% T: 72%
AFR C: 8% T: 92%
AMR C: 30% T: 70%
EAS C: 39% T: 61%
EUR C: 31% T: 69%
SAS C: 41% T: 59%
Base 136 IGHV4-4 (SNP rs74872186)
1000 Genomes Project Phase 3 allele frequencies
ALL G: 14% C: 86%
AFR G: 4% C: 96%
AMR G: 16% C: 84%
EAS G: 15% C: 85%
EUR G: 15% C: 85%
SAS G: 24% C: 76%
Base 207 IGHV4-4 (SNP rs77238211)
1000 Genomes Project Phase 3 allele frequencies
ALL C: 17% G: 83%
AFR C: 3% G: 97%
AMR C: 20% G: 80%
EAS C: 22% G: 78%
EUR C: 18% G: 82%
SAS C: 27% G: 73%
Base 308 IGHV4-4 (SNP rs150123115)
1000 Genomes Project Phase 3 allele frequencies
ALL T: 88% C: 12%
AFR T: 92% C: 8%
AMR T: 92% C: 8%
EAS T: 75% C: 25%
EUR T: 97% C: 3%
SAS T: 82% C: 18%

**Supplementary Figure 6.** Allelic variants of IGHV7-4-1 as defined by IMGT are illustrated. Variability of some of the positions of these genes in samples obtained in different geographical locations as illustrated by the ENSEMBL browser (release 101, August 2020) (19) is shown. Sequence variation in base 274 (IMGT numbering nomenclature (16)) suggests that the base associated to IGHV7-4-1*01 is more common than the base associated to other alleles of this gene in most populations. All sequence variants of the illustrations of SNPs are indicated as seen in the reversed strand, hence they are complementary to the base of the coding strand.

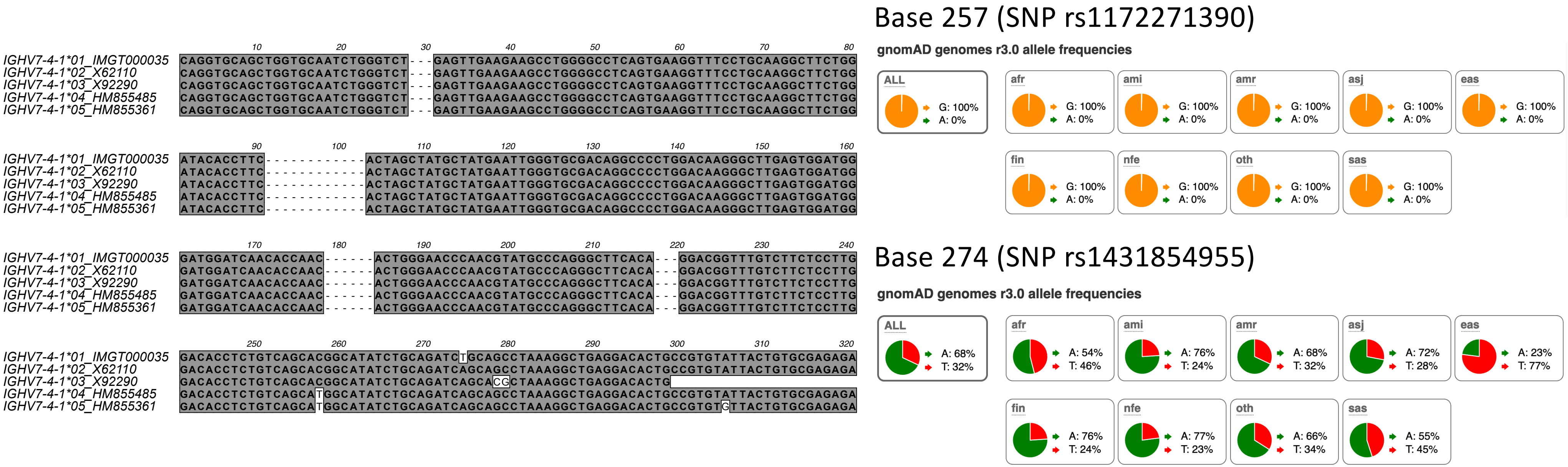

IGHV7-4-1*01_IMGT000035
IGHV7-4-1*02_X62110
IGHV7-4-1*03_X92290
IGHV7-4-1*04_HM855485
IGHV7-4-1*05_HM855361
Base 257 (SNP rs1172271390)
gnomAD genomes r3.0 allele frequencies
ALL G: 100% A: 0%
afr G: 100% A: 0%
ami G: 100% A: 0%
amr G: 100% A: 0%
asj G: 100% A: 0%
eas G: 100% A: 0%
fin G: 100% A: 0%
nfe G: 100% A: 0%
oth G: 100% A: 0%
sas G: 100% A: 0%
Base 274 (SNP rs1431854955)
gnomAD genomes r3.0 allele frequencies
ALL A: 68% T: 32%
afr A: 54% T: 46%
ami A: 76% T: 24%
amr A: 68% T: 32%
asj A: 72% T: 28%
eas A: 23% T: 77%
fin A: 76% T: 24%
nfe A: 77% T: 23%
oth A: 66% T: 34%
sas A: 55% T: 45%

**Supplementary Figure 7.** High resolution structures of five antibodies with a heavy chain variable domain encoded by a gene with an origin in IGHV7-4-1. Heavy chain CDR3 is shown at the top of each sequence in red. The side chain of residue 92 (in all these cases a serine), located far from the antibody binding site is shown in green (carbon) and red (oxygen). Structures include PDB entries 4D9Q, (A), 4EOW (B), 5CGY (C), 5ZMJ (D), and 6B5R (E).

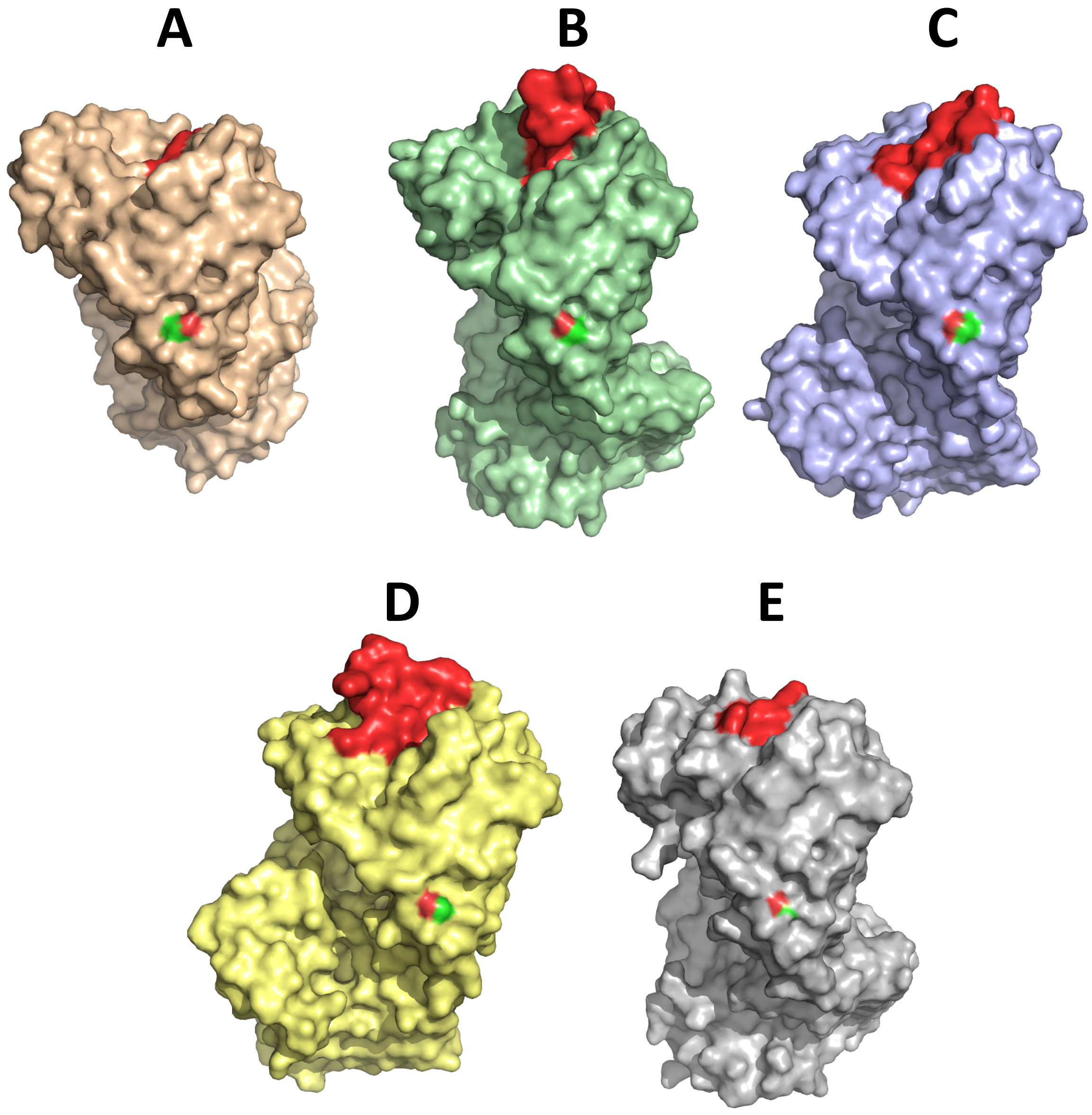
A
B
C
D
E

**Supplementary Figure 8.** Translated sequences of productive IgA and IgG-encoding reads derived from NGS data sets of two subjects (donors 2 and 4) that both have access to IGHV7-4-1*01 but not IGHV7-4-1*02 to generate an antibody repertoire (11). The sequencing protocol (15) allowed for determination of the sequence from the end of framework 1 and extended into the first constant domain of the heavy chain. The sequences encoded by IGHV7-4-1*01 and IGHV7-4-1*02 are shown on top of the figure. Residue 92 is highlighted by an arrow.

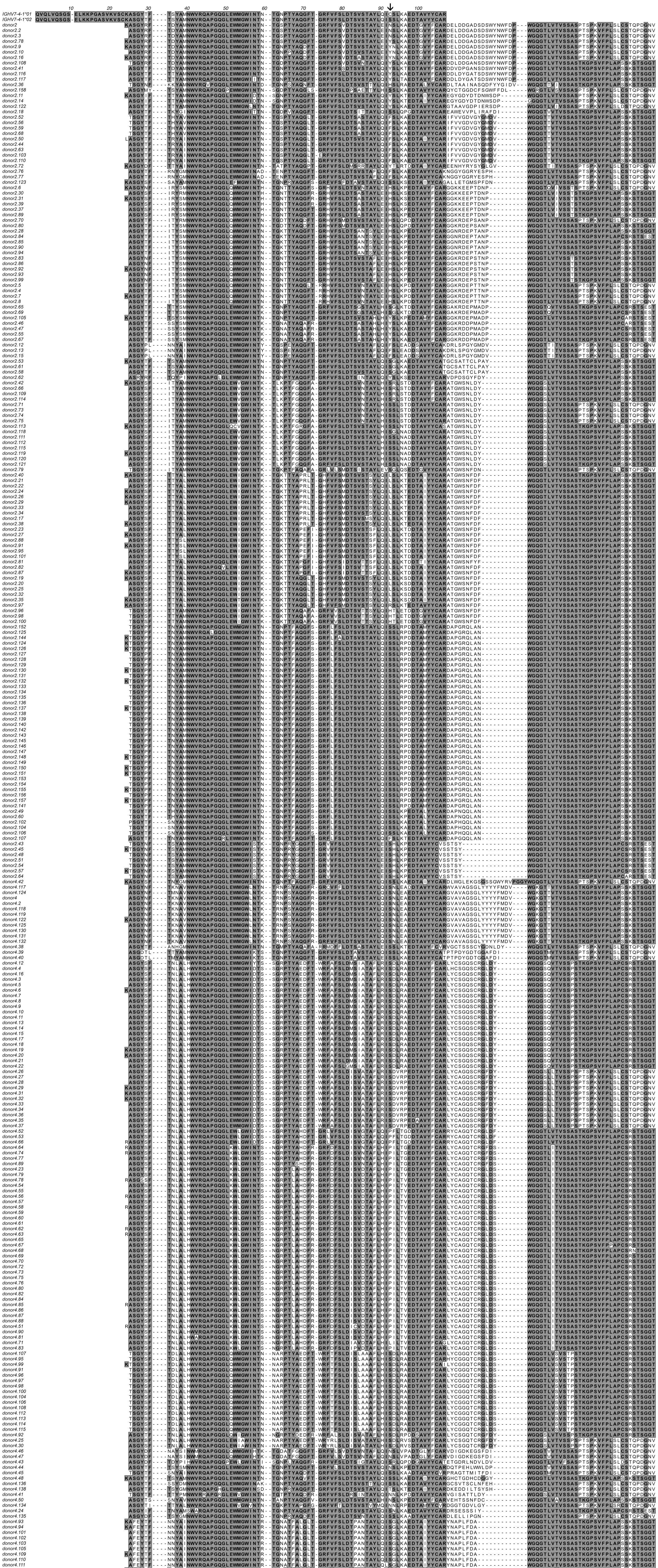
