## Supplementary Methods for "Poorly expressed alleles of several human immunoglobulin heavy chain variable (IGHV) genes are common in the human population"

Mats Ohlin

Dept. of Immunotechnology, Lund University, Lund, Sweden

Mats Ohlin, Dept. of Immunotechnology, Lund University, Medicon Village building 406, S-22381 Lund, Sweden.. Telephone: +46-462224322

**Databases of germline genes used to initiate the inference process**

**V.fasta**

**D.fasta**

**J.fasta**

**IgDiscover configuration file**

**igdiscover.yaml**

**Databases of germline genes used to initiate the inference process**

**V.fasta**

```
>IGHV1-18*01
caggttcagctggtgcagtctggagctgaggtgaagaagcctggggcctcagtgaaggtc
tcctgcaaggcttctggttacacctttaccagctatggtatcagctgggtgcgacaggcc
cctggacaagggcttgagtggatgggatggatcagcgcttacaatggtaacacaaactat
gcacagaagctccagggcagagtcaccatgaccacagacacatccacgagcacagcctac
atggagctgaggagcctgagatctgacgacacggccgtgtattactgtgcgagaga
>IGHV1-18*02
caggttcagctggtgcagtctggagctgaggtgaagaagcctggggcctcagtgaaggtc
tcctgcaaggcttctggttacacctttaccagctatggtatcagctgggtgcgacaggcc
cctggacaagggcttgagtggatgggatggatcagcgcttacaatggtaacacaaactat
gcacagaagctccagggcagagtcaccatgaccacagacacatccacgagcacagcctac
atggagctgaggagcctaagatctgacgacacggcc
>IGHV1-18*03
caggttcagctggtgcagtctggagctgaggtgaagaagcctggggcctcagtgaaggtc
tcctgcaaggcttctggttacacctttaccagctatggtatcagctgggtgcgacaggcc
cctggacaagggcttgagtggatgggatggatcagcgcttacaatggtaacacaaactat
gcacagaagctccagggcagagtcaccatgaccacagacacatccacgagcacagcctac
atggagctgaggagcctgagatctgacgacatggccgtgtattactgtgcgagaga
>IGHV1-18*04
caggttcagctggtgcagtctggagctgaggtgaagaagcctggggcctcagtgaaggtc
tcctgcaaggcttctggttacacctttaccagctacggtatcagctgggtgcgacaggcc
cctggacaagggcttgagtggatgggatggatcagcgcttacaatggtaacacaaactat
gcacagaagctccagggcagagtcaccatgaccacagacacatccacgagcacagcctac
atggagctgaggagcctgagatctgacgacacggccgtgtattactgtgcgagaga
>IGHV1-2*01
caggtgcagctggtgcagtctggggctgaggtgaagaagcctggggcctcagtgaaggtc
tcctgcaaggcttctggatacaccttcaccggctactatatgcactgggtgcgacaggcc
cctggacaagggcttgagtggatgggacggatcaaccctaacagtggtggcacaaactat
gcacagaagtttcagggcagggtcaccagtaccagggacacgtccatcagcacagcctac
atggagctgagcaggctgagatctgacgacacggtcgtgtattactgtgcgagaga
>IGHV1-2*02
```

caggtgcagctggtgcagtctggggctgaggtgaagaagcctggggcctcagtgaaggtc
tcctgcaaggcttctggatacaccttcaccggctactatatgcactgggtgcgacaggcc
cctggacaagggcttgagtggatgggatggatcaaccctaacagtggtggcacaaactat
gcacagaagtttcagggcagggtcaccatgaccagggacacgtccatcagcacagcctac
atggagctgagcaggctgagatctgacgacacggccgtgtattactgtgcgagaga
>IGHV1-2*03
caggtgcagctggtgcagtctggggctgaggtgaagaagcttggggcctcagtgaaggtc
tcctgcaaggcttctggatacaccttcaccggctactatatgcactgggtgcnacaggcc
cctggacaagggcttgagtggatgggatggatcaaccctaacagtggtggcacaaactat
gcacagaagtttcagggcagggtcaccatgaccagggacacgtccatcagcacagcctac
atggagctgagcaggctgagatctgacgacacggccgtgtattactgtgcgagaga
>IGHV1-2*04
caggtgcagctggtgcagtctggggctgaggtgaagaagcctggggcctcagtgaaggtc
tcctgcaaggcttctggatacaccttcaccggctactatatgcactgggtgcgacaggcc
cctggacaagggcttgagtggatgggatggatcaaccctaacagtggtggcacaaactat
gcacagaagtttcagggctgggtcaccatgaccagggacacgtccatcagcacagcctac
atggagctgagcaggctgagatctgacgacacggccgtgtattactgtgcgagaga
>IGHV1-2*05
caggtgcagctggtgcagtctggggctgaggtgaagaagcctggggcctcagtgaaggtc
tcctgcaaggcttctggatacaccttcaccggctactatatgcactgggtgcgacaggcc
cctggacaagggcttgagtggatgggacggatcaaccctaacagtggtggcacaaactat
gcacagaagtttcagggcagggtcaccatgaccagggacacgtccatcagcacagcctac
atggagctgagcaggctgagatctgacgacacggtcgtgtattactgtgcgagaga
>IGHV1-2*06
caggtgcagctggtgcagtctggggctgaggtgaagaagcctggggcctcagtgaaggtc
tcctgcaaggcttctggatacaccttcaccggctactatatgcactgggtgcgacaggcc
cctggacaagggcttgagtggatgggacggatcaaccctaacagtggtggcacaaactat
gcacagaagtttcagggcagggtcaccatgaccagggacacgtccatcagcacagcctac
atggagctgagcaggctgagatctgacgacacggccgtgtattactgtgcgagaga
>IGHV1-24*01
caggtccagctggtacagtctggggctgaggtgaagaagcctggggcctcagtgaaggtc
tcctgcaaggtttccggatacaccctcactgaattatccatgcactgggtgcgacaggct
cctggaaaagggcttgagtggatgggaggttttgatcctgaagatggtgaaacaatctac
gcacagaagttccagggcagagtcaccatgaccgaggacacatctacagacacagcctac

atggagctgagcagcctgagatctgaggacacggccgtgtattactgtgcaacaga
>IGHV1-3*01
caggtccagcttgtgcagtctggggctgaggtgaagaagcctggggcctcagtgaaggtt
tcctgcaaggcttctggatacaccttcactagctatgctatgcattgggtgcgccaggcc
cccggacaaaggcttgagtggatgggatggatcaacgctggcaatggtaacacaaaatat
tcacagaagttccagggcagagtcaccattaccagggacacatccgcgagcacagcctac
atggagctgagcagcctgagatctgaagacacggctgtgtattactgtgcgagaga
>IGHV1-3*02
caggttcagctggtgcagtctggggctgaggtgaagaagcctggggcctcagtgaaggtt
tcctgcaaggcttctggatacaccttcactagctatgctatgcattgggtgcgccaggcc
cccggacaaaggcttgagtggatgggatggagcaacgctggcaatggtaacacaaaatat
tcacaggagttccagggcagagtcaccattaccagggacacatccgcgagcacagcctac
atggagctgagcagcctgagatctgaggacatggctgtgtattactgtgcgagaga
>IGHV1-3*03
caggtccagctggtgcagtctggggctgaggtgaagaagcctggggcctcagtgaaggtt
tcctgcaaggcttctggatacaccttcactagctatgctatgcattgggtgcgccaggcc
cccggacaaaggcttgagtggatgggatggatcaacgctggcaatggtaacacaaaatat
tcacaggagttccagggcagagtcaccattaccagggacacatccgcgagcacagcctac
atggagctgagcagcctgagatctgaggacatggctgtgtattactgtgcgagaga
>IGHV1-3*04
caggtccagcttgtgcagtctggggctgaggtgaagaagcctggggcctcagtgaaggtt
tcctgcaaggcttctggatacaccttcactagctatgctatgcattgggtgcgccaggcc
cccggacaaaggcttgagtggatgggatggatcaacactggcaatggtaacacaaaatat
tcacagaagttccagggcagagtcaccattaccagggacacatccgcgagcacagcctac
atggagctgagcagcctgagatctgaagacacggct
>IGHV1-45*01
cagatgcagctggtgcagtctggggctgaggtgaagaagactgggtcctcagtgaaggtt
tcctgcaaggcttccggatacaccttcacctaccgctacctgcactgggtgcgacaggcc
cccggacaagcgcttgagtggatgggatggatcacacctttcaatggtaacaccaactac
gcacagaaattccaggacagagtcaccattactagggacaggtctatgagcacagcctac
atggagctgagcagcctgagatctgaggacacagccatgtattactgtgcaagana
>IGHV1-45*02
cagatgcagctggtgcagtctggggctgaggtgaagaagactgggtcctcagtgaaggtt
tcctgcaaggcttccggatacaccttcacctaccgctacctgcactgggtgcgacaggcc

cccggacaagcgcttgagtggatgggatggatcacacctttcaatggtaacaccaactac
gcacagaaattccaggacagagtcaccattaccagggacaggtctatgagcacagcctac
atggagctgagcagcctgagatctgaggacacagccatgtattactgtgcaagata
>IGHV1-45*03
cagatgcagctggtgcagtctggggctgaggtgaagaagactgggtcctcagtgaaggtt
tcctgcaaggcttccggatacaccttcacctaccgctacctgcactgggtgcgacaggcc
cccagacaagcgcttgagtggatgggatggatcacacctttcaatggtaacaccaactac
gcacagaaattccaggacagagtcaccattaccagggacaggtctatgagcacagcctac
atggagctgagcagcctgagatctgaggacacagccatgtattactgtgcaagata
>IGHV1-46*01
caggtgcagctggtgcagtctggggctgaggtgaagaagcctggggcctcagtgaaggtt
tcctgcaaggcatctggatacaccttcaccagctactatatgcactgggtgcgacaggcc
cctggacaagggcttgagtggatgggaataatcaaccctagtggtggtagcacaagctac
gcacagaagttccagggcagagtcaccatgaccagggacacgtccacgagcacagtctac
atggagctgagcagcctgagatctgaggacacggccgtgtattactgtgcgagaga
>IGHV1-46*02
caggtgcagctggtgcagtctggggctgaggtgaagaagcctggggcctcagtgaaggtt
tcctgcaaggcatctggatacaccttcaacagctactatatgcactgggtgcgacaggcc
cctggacaagggcttgagtggatgggaataatcaaccctagtggtggtagcacaagctac
gcacagaagttccagggcagagtcaccatgaccagggacacgtccacgagcacagtctac
atggagctgagcagcctgagatctgaggacacggccgtgtattactgtgcgagaga
>IGHV1-46*03
caggtgcagctggtgcagtctggggctgaggtgaagaagcctggggcctcagtgaaggtt
tcctgcaaggcatctggatacaccttcaccagctactatatgcactgggtgcgacaggcc
cctggacaagggcttgagtggatgggaataatcaaccctagtggtggtagcacaagctac
gcacagaagttccagggcagagtcaccatgaccagggacacgtccacgagcacagtctac
atggagctgagcagcctgagatctgaggacacggccgtgtattactgtgctagaga
>IGHV1-46*04
caggtgcagctggtgcagtctggggctgaggtgaagaagcctggggcctcagtgaaggtt
tcctgcaaggcatctggatacaccttcaccagctactatatgcactgggtgcgacaggcc
cctggacaagggcttgagtggatgggaataatcaaccctagtggtggtagcacaagctac
gcacagaagttgcagggcagagtcaccatgaccagggacacgtccacgagcacagtctac
atggagctgagcagcctgagatctgaggacacggccgtgtattactgtgcgagaga
>IGHV1-58*01

caaatgcagctggtgcagtctgggcctgaggtgaagaagcctgggacctcagtgaaggtc
tcctgcaaggcttctggattcacctttactagctctgctgtgcagtgggtgcgacaggct
cgtggacaacgccttgagtggataggatggatcgtcgttggcagtggtaacacaaactac
gcacagaagttccaggaaagagtcaccattaccagggacatgtccacaagcacagcctac
atggagctgagcagcctgagatccgaggacacggccgtgtattactgtgcggcaga
>IGHV1-58*02
caaatgcagctggtgcagtctgggcctgaggtgaagaagcctgggacctcagtgaaggtc
tcctgcaaggcttctggattcacctttactagctctgctatgcagtgggtgcgacaggct
cgtggacaacgccttgagtggataggatggatcgtcgttggcagtggtaacacaaactac
gcacagaagttccaggaaagagtcaccattaccagggacatgtccacaagcacagcctac
atggagctgagcagcctgagatccgaggacacggccgtgtattactgtgcggcaga
>IGHV1-58*03
caaatgcagctggtgcagtctgggcctgaagtgaagaagcctgggacctcagtgaaggtc
tcctgcaaggcttctggattcacctttactagctctgctgtgcagtgggtgcgacaggct
cgtggacaacgccttgagtggataggatggatcgtcgttggcagtggtaacacaaactac
gcacagaagttccaggaaagagtcaccattaccagggacatgtccacaagcacagcctac
atggagctgagcagcctgagatccgaggacacggccgtgtattactgtgcggcaga
>IGHV1-69*01
caggtgcagctggtgcagtctggggctgaggtgaagaagcctgggtcctcggtgaaggtc
tcctgcaaggcttctggaggcaccttcagcagctatgctatcagctgggtgcgacaggcc
cctggacaagggcttgagtggatgggagggatcatccctatctttggtacagcaaactac
gcacagaagttccagggcagagtcacgattaccgcggacgaatccacgagcacagcctac
atggagctgagcagcctgagatctgaggacacggccgtgtattactgtgcgagaga
>IGHV1-69*02
caggtccagctggtgcaatctggggctgaggtgaagaagcctgggtcctcggtgaaggtc
tcctgcaaggcttctggaggcaccttcagcagctatactatcagctgggtgcgacaggcc
cctggacaagggcttgagtggatgggaaggatcatccctatccttggtatagcaaactac
gcacagaagttccagggcagagtcacgattaccgcggacaaatccacgagcacagcctac
atggagctgagcagcctgagatctgaggacacggccgtgtattactgtgcgaga
>IGHV1-69*03
caggtgcagctggtgcagtctggggctgaggtgaagaagcctgggtcctcggtgaaggtc
tcctgcaaggcttctggaggcaccttcagcagctatgctatcagctgggtgcgacaggcc
cctggacaagggcttgagtggatgggagggatcatccctatctttggtacagcaaactac
gcacagaagttccagggcagagtcacgattaccgcggacgaatccacgagcacagcctac

atggagctgagcagcctgagatctgatgacacggc
>IGHV1-69*04
caggtccagctggtgcagtctggggctgaggtgaagaagcctgggtcctcggtgaaggtc
tcctgcaaggcttctggaggcaccttcagcagctatgctatcagctgggtgcgacaggcc
cctggacaagggcttgagtggatgggaaggatcatccctatccttggtatagcaaactac
gcacagaagttccagggcagagtcacgattaccgcggacaaatccacgagcacagcctac
atggagctgagcagcctgagatctgaggacacggccgtgtattactgtgcgagaga
>IGHV1-69*05
caggtccagctggtgcagtctggggctgaggtgaagaagcctgggtcctcggtgaaggtc
tcctgcaaggcttctggaggcaccttcagcagctatgctatcagctgggtgcgacaggcc
cctggacaagggcttgagtggatgggagggatcatccctatctttggtacagcaaactac
gcacagaagttccagggcagagtcacgattaccacggacgaatccacgagcacagcctac
atggagctgagcagcctgagatctgaggacacggccgtgtattactgtgcgaga
>IGHV1-69*06
caggtgcagctggtgcagtctggggctgaggtgaagaagcctgggtcctcggtgaaggtc
tcctgcaaggcttctggaggcaccttcagcagctatgctatcagctgggtgcgacaggcc
cctggacaagggcttgagtggatgggagggatcatccctatctttggtacagcaaactac
gcacagaagttccagggcagagtcacgattaccgcggacaaatccacgagcacagcctac
atggagctgagcagcctgagatctgaggacacggccgtgtattactgtgcgagaga
>IGHV1-69*07
agaagcctgggtcctcggtgaaggtctcctgcaaggcttctggaggcaccttcagcagct
atgctatcagctgggtgcgacaggcccctggacaagggcttgagtggatgggaaggatca
tccctatctttggtacagcaaactacgcacagaagttccagggcagagtcacgattaccg
cggacgaatccacgagcacagcctacatggagctgagcagcctgagatctgag
>IGHV1-69*08
caggtccagctggtgcaatctggggctgaggtgaagaagcctgggtcctcggtgaaggtc
tcctgcaaggcttctggaggcaccttcagcagctatactatcagctgggtgcgacaggcc
cctggacaagggcttgagtggatgggaaggatcatccctatccttggtacagcaaactac
gcacagaagttccagggcagagtcacgattaccgcggacaaatccacgagcacagcctac
atggagctgagcagcctgagatctgaggacacggccgtgtattactgtgcgagaga
>IGHV1-69*09
caggtgcagctggtgcagtctggggctgaggtgaagaagcctgggtcctcggtgaaggtc
tcctgcaaggcttctggaggcaccttcagcagctatgctatcagctgggtgcgacaggcc
cctggacaagggcttgagtggatgggaaggatcatccctatccttggtatagcaaactac

gcacagaagttccagggcagagtcacgattaccgcggacaaatccacgagcacagcctac
atggagctgagcagcctgagatctgaggacacggccgtgtattactgtgcgagaga
>IGHV1-69*10
caggtccagctggtgcagtctggggctgaggtgaagaagcctgggtcctcagtgaaggtc
tcctgcaaggcttctggaggcaccttcagcagctatgctatcagctgggtgcgacaggcc
cctggacaagggcttgagtggatgggagggatcatccctatccttggtatagcaaactac
gcacagaagttccagggcagagtcacgattaccgcggacaaatccacgagcacagcctac
atggagctgagcagcctgagatctgaggacacggccgtgtattactgtgcgagaga
>IGHV1-69*11
caggtccagctggtgcagtctggggctgaggtgaagaagcctgggtcctcggtgaaggtc
tcctgcaaggcttctggaggcaccttcagcagctatgctatcagctgggtgcgacaggcc
cctggacaagggcttgagtggatgggaaggatcatccctatccttggtacagcaaactac
gcacagaagttccagggcagagtcacgattaccgcggacgaatccacgagcacagcctac
atggagctgagcagcctgagatctgaggacacggccgtgtattactgtgcgagaga
>IGHV1-69*12
caggtccagctggtgcagtctggggctgaggtgaagaagcctgggtcctcggtgaaggtc
tcctgcaaggcttctggaggcaccttcagcagctatgctatcagctgggtgcgacaggcc
cctggacaagggcttgagtggatgggagggatcatccctatctttggtacagcaaactac
gcacagaagttccagggcagagtcacgattaccgcggacgaatccacgagcacagcctac
atggagctgagcagcctgagatctgaggacacggccgtgtattactgtgcgagaga
>IGHV1-69*13
caggtccagctggtgcagtctggggctgaggtgaagaagcctgggtcctcagtgaaggtc
tcctgcaaggcttctggaggcaccttcagcagctatgctatcagctgggtgcgacaggcc
cctggacaagggcttgagtggatgggagggatcatccctatctttggtacagcaaactac
gcacagaagttccagggcagagtcacgattaccgcggacgaatccacgagcacagcctac
atggagctgagcagcctgagatctgaggacacggccgtgtattactgtgcgagaga
>IGHV1-69*14
caggtccagctggtgcagtctggggctgaggtgaagaagcctgggtcctcggtgaaggtc
tcctgcaaggcttctggaggcaccttcagcagctatgctatcagctgggtgcgacaggcc
cctggacaagggcttgagtggatgggagggatcatccctatctttggtacagcaaactac
gcacagaagttccagggcagagtcacgattaccgcggacaaatccacgagcacagcctac
atggagctgagcagcctgagatctgaggacacggccgtgtattactgtgcgagaga
>IGHV1-69*15
caggtccagctggtgcagtctggggctgaggtgaagaagcctgggtcctcggtgaaggtc

tcctgcaaggcttctggaggcaccttcagcagctatgctatcagctgggtgcgacaggcc
cctggacaagggcttgagtggatgggaaggatcatccctatctttggtacagcaaactac
gcacagaagttccagggcagagtcacgattaccgcggacgaatccacgagcacagcctac
atggagctgagcagcctgagatctgaggacacggccgtgtattactgtgcgagaga
>IGHV1-69*16
caggtccagctggtgcagtctggggctgaggtgaagaagcctgggtcctcggtgaaggtc
tcctgcaaggcttctggaggcaccttcagcagctatactatcagctgggtgcgacaggcc
cctggacaagggcttgagtggatgggagggatcatccctatccttggtacagcaaactac
gcacagaagttccagggcagagtcacgattaccacggacgaatccacgagcacagcctac
atggagctgagcagcctgagatctgaggacacggccgtgtattactgtgcgagaga
>IGHV1-69*17
caggtgcagctggtgcagtctggggctgaggtgaagaagcctgggtcctcggtgaaggtc
tcctgcaaggcttctggaggcaccttcagcagctatgctatcagctgggtgcgacaggcc
cctggacaagggcttgagtggatgggagggatcatccctatctttggtatagcaaactac
gcacagaagttccagggcagagtcacgattaccgcggacaaatccacgagcacagcctac
atggagctgagcagcctgagatctgaggacacggccgtgtattactgtgcgagaga
>IGHV1-69-2*01
gaggtccagctggtacagtctggggctgaggtgaagaagcctggggctacagtgaaaatc
tcctgcaaggtttctggatacaccttcaccgactactacatgcactgggtgcaacaggcc
cctggaaaagggcttgagtggatgggacttgttgatcctgaagatggtgaaacaatatac
gcagagaagttccagggcagagtcaccataaccgcggacacgtctacagacacagcctac
atggagctgagcagcctgagatctgaggacacggccgtgtattactgtgcaacaga
>IGHV1-69-2*02
agaagcctggggctacagtgaaaatctcctgcaaggtttctggatacaccttcaccgact
actacatgcactgggtgcaacaggcccctggaaaagggcttgagtggatgggacttgttg
atcctgaagatggtgaaacaatatatgcagagaagttccagggcagagtcaccataaccg
cggacacgtctacagacacagcctacatggagctgagcagcctgagatctgag
>IGHV1-69D*01
caggtgcagctggtgcagtctggggctgaggtgaagaagcctgggtcctcggtgaaggtc
tcctgcaaggcttctggaggcaccttcagcagctatgctatcagctgggtgcgacaggcc
cctggacaagggcttgagtggatgggagggatcatccctatctttggtacagcaaactac
gcacagaagttccagggcagagtcacgattaccgcggacgaatccacgagcacagcctac
atggagctgagcagcctgagatctgaggacacggccgtgtattactgtgcgagaga
>IGHV1-8*01

caggtgcagctggtgcagtctggggctgaggtgaagaagcctggggcctcagtgaaggtc
tcctgcaaggcttctggatacaccttcaccagttatgatatcaactgggtgcgacaggcc
actggacaagggcttgagtggatgggatggatgaaccctaacagtggtaacacaggctat
gcacagaagttccagggcagagtcaccatgaccaggaacacctccataagcacagcctac
atggagctgagcagcctgagatctgaggacacggccgtgtattactgtgcgagagg
>IGHV1-8*02
caggtgcagctggtgcagtctggggctgaggtgaagaagcctggggcctcagtgaaggtc
tcctgcaaggcttctggatacaccttcaccagctatgatatcaactgggtgcgacaggcc
actggacaagggcttgagtggatgggatggatgaaccctaacagtggtaacacaggctat
gcacagaagttccagggcagagtcaccatgaccaggaacacctccataagcacagcctac
atggagctgagcagcctgagatctgaggacacggccgtgtattactgtgcgagagg
>IGHV1-8*03
caggtgcagctggtgcagtctggggctgaggtgaagaagcctggggcctcagtgaaggtc
tcctgcaaggcttctggatacaccttcaccagctatgatatcaactgggtgcgacaggcc
actggacaagggcttgagtggatgggatggatgaaccctaacagtggtaacacaggctat
gcacagaagttccagggcagagtcaccattaccaggaacacctccataagcacagcctac
atggagctgagcagcctgagatctgaggacacggccgtgtattactgtgcgagagg
>IGHV2-26*01
caggtcaccttgaaggagtctggtcctgtgctggtgaaacccacagagaccctcacgctg
acctgcaccgtctctgggttctcactcagcaatgctagaatgggtgtgagctggatccgt
cagcccccagggaaggccctggagtggcttgcacacatttttcgaatgacgaaaaatcc
tacagcacatctctgaagagcaggctcaccatctccaaggacacctccaaaagccaggtg
gtccttaccatgaccaacatggaccctgtggacacagccacatattactgtgcacggata
c
>IGHV2-26*02
caggtcaccttgaaggagtctggtcctgtgctggtgaaacccacagagaccctcacgctg
acctgcaccgtctctgggttctcactcagcaatgctagaatgggtgtgagctggatccgt
cagcccccagggaaggccctggagtggcttgcacacatttttcgaatgacgaaaaatcc
tacagcacatctctgaagagcaggctcaccatctccaaggacacctccaaaagccaggtg
gtccttaccatgaccaatatggaccctgtggacacagccacatattactgtgcacggata
c
>IGHV2-26*03
caggtcaccttgaaggagtctggtcctgtgctggtgaaacccacagagaccctcacgctg
acctgcaccatctctgggttctcactcagcaatgctagaatgggtgtgagctggatccgt

cagcccccagggaaggccctggagtggcttgcacacatttttcgaatgacgaaaaatcc
tacagcacatctctgaagagcaggctcaccatctccaaggacacctccaaaagccaggtg
gtccttaccatgaccaacatggaccctgtggacacagccacatattactgtgcacggata
c
>IGHV2-5*01
cagatcaccttgaaggagtctggtcctacgctggtgaaacccacacagaccctcacgctg
acctgcaccttctctgggttctcactcagcactagtggagtgggtgtgggctggatccgt
cagcccccaggaaaggccctggagtggcttgcactcatttattggaatgatgataagcgc
tacagcccatctctgaagagcaggctcaccatcaccaaggacacctccaaaaaccaggtg
gtccttacaatgaccaacatggaccctgtggacacagccacatattactgtgcacacaga
c
>IGHV2-5*02
cagatcaccttgaaggagtctggtcctacgctggtgaaacccacacagaccctcacgctg
acctgcaccttctctgggttctcactcagcactagtggagtgggtgtgggctggatccgt
cagcccccaggaaaggccctggagtggcttgcactcatttattgggatgatgataagcgc
tacagcccatctctgaagagcaggctcaccatcaccaaggacacctccaaaaaccaggtg
gtccttacaatgaccaacatggaccctgtggacacagccacatattactgtgcacacaga
c
>IGHV2-5*03
gctggtgaaacccacacagaccctcacgctgacctgcaccttctctgggttctcactcag
cactagtggagtgggtgtgggctggatccgtcagcccccaggaaaggccctggagtggct
tgcactcatttattgggatgatgataagcgctacagcccatctctgaagagcaggctcac
cattaccaaggacacctccaaaaaccaggt
>IGHV2-5*04
cagatcaccttgaaggagtctggtcctacgctggtgaaacccacacagaccctcacgctg
acctgcaccttctctgggttctcactcagcactagtggagtgggtgtgggctggatccgt
cagcccccaggaaaggccctggagtggcttgcactcatttattggaatgatgataagcgc
tacagcccatctctgaagagcaggctcaccatcaccaaggacacctccaaaaaccaggtg
gtccttacaatgaccaacatggaccctgtggacacaggcacatattactgtgtac
>IGHV2-5*05
cagatcaccttgaaggagtctggtcctacgctggtgaaacccacacagaccctcacgctg
acctgcaccttctctgggttctcactcagcactagtggagtgggtgtgggctggatccgt
cagcccccaggaaaggccctggagtggcttgcactcatttattgggatgatgataagcgc
tacggcccatctctgaagagcaggctcaccatcaccaaggacacctccaaaaaccaggtg

gtccttacaatgaccaacatggaccctgtggacacagccacatattactgtgcacacaga
c
>IGHV2-5*06
cagatcaccttgaaggagtctggtcctacgctggtaaaacccacacagaccctcacgctg
acctgcaccttctctgggttctcactcagcactagtggagtgggtgtgggctggatccgt
cagcccccaggaaaggccctggagtggcttgcactcatttattgggatgatgataagcgc
tacggcccatctctgaagagcaggctcaccatcaccaaggacacctccaaaaaccaggtg
gtccttacaatgaccaacatggaccctgtggacacagccacatattactgtgcacacaga
>IGHV2-5*08
caggtcaccttgaaggagtctggtcctgcgctggtgaaacccacacagaccctcacactg
acctgcaccttctctgggttctcactcagcactagtggaatgcgtgtgagctggatccgt
cagcccccaggaaaggccctggagtggcttgcactcatttattgggatgatgataagcgc
tacagcccatctctgaagagcaggctcaccatcaccaaggacacctccaaaaaccaggtg
gtccttacaatgaccaacatggaccctgtggacacagccacatattactgtgcacacaga
c
>IGHV2-5*09
caggtcaccttgaaggagtctggtcctacgctggtgaaacccacacagaccctcacgctg
acctgcaccttctctgggttctcactcagcactagtggagtgggtgtgggctggatccgt
cagcccccaggaaaggccctggagtggcttgcactcatttattgggatgatgataagcgc
tacggcccatctctgaagagcaggctcaccatcaccaaggacacctccaaaaaccaggtg
gtccttacaatgaccaacatggaccctgtggacacagccacatattactgtgcacacaga
c
>IGHV2-70*01
caggtcaccttgagggagtctggtcctgcgctggtgaaacccacacagaccctcacactg
acctgcaccttctctgggttctcactcagcactagtggaatgtgtgtgagctggatccgt
cagcccccagggaaggccctggagtggcttgcactcattgattgggatgatgataaatac
tacagcacatctctgaagaccaggctcaccatctccaaggacacctccaaaaaccaggtg
gtccttacaatgaccaacatggaccctgtggacacagccacgtattactgtgcacggata
c
>IGHV2-70*02
caggtcaccttgagggagtctggtcctgcgctggtgaaacccacacagaccctcacactg
acctgcaccttctctgggttctcactcagcactagtggaatgtgtgtgagctggatccgt
cagcccccagggaaggccctggagtggcttgcactcattgattgggatgatgataaatac
tacagcacatctctgaagaccaggctcaccatctccaaggacacctccaaaaaccaggtg

gtccttacaatgaccaacatggaccctgtggacacggccgtgtattactg
>IGHV2-70*03
caggtcaccttgaaggagtctggtcctgcgctggtgaaacccacacagaccctcacactg
acctgcaccttctctgggttctcactcagcactagtggaatgcgtgtgagctggatccgt
cagcccccagggaaggccctggagtggcttgcacgcattgattgggatgatgataaattc
tacagcacatctctgaagaccaggctcaccatctccaaggacacctccaaaaaccaggtg
gtccttacaatgaccaacatggaccctgtggacacggccgtgtattactg
>IGHV2-70*04
caggtcaccttgaaggagtctggtcctgcgctggtgaaacccacacagaccctcacactg
acctgcaccttctctgggttctcactcagcactagtggaatgcgtgtgagctggatccgt
cagcccccagggaaggccctggagtggcttgcacgcattgattgggatgatgataaattc
tacagcacatctctgaagaccaggctcaccatctccaaggacacctccaaaaaccaggtg
gtccttacaatgaccaacatggaccctgtggacacagccacgtattactgtgcacggata
c
>IGHV2-70*05
tgcgctggtgaaacccacacagaccctcacactgacctgcaccttctctgggttctcact
cagcactagtggaatgcgtgcgagctggatccgtcagcccccagggaaggccctggagtg
gcttgcacgcattgattgggatgatgataaattctacagcacatctctgaagaccaggct
caccatctccaaggacacctccaaaaaccaggtggtccttacaatgaccaacatgga
>IGHV2-70*06
caggtcaccttgaaggagtctggtcctgcgctggtgaaacccacacagaccctcacactg
acctgcaccttctctgggttctcactcagcactagtggaatgcgtgtgagctggatccgt
cagcccccagggaaggccctggagtggcttgcacgcattgattgggatgatgataaattc
tacagcacatccctgaagaccaggctcaccatctccaaggacacctccaaaaaccaggtg
gtccttacaatgaccaacatggaccctgtggacacggccgtgtattactg
>IGHV2-70*07
caggtcaccttgagggagtctggtcctgcgctggtgaaacccacacagaccctcacactg
acctgcaccttctctgggttctcactcagcactagtggaatgtgtgtgagctggatccgt
cagcccccggggaaggccctggagtggcttgcactcattgattgggatgatgataaatac
tacagcacatctctgaagaccaggctcaccatctccaaggacacctccaaaaaccaggtg
gtccttacaatgaccaacatggaccctgtggacacggccgtgtattactg
>IGHV2-70*08
caggtcaccttgagggagtctggtcctgcgctggtgaaacccacacagaccctcacactg
acctgcgccttctctgggttctcactcagcactagtggaatgtgtgtgagctggatccgt

cagcccccagggaaggccctggagtggcttgcacgcattgattgggatgatgataaatac
tacagcacatctctgaagaccaggctcaccatctccaaggacacctccaaaaaccaggtg
gtccttacaatgaccaacatggaccctgtggacacggccgtgtattactg
>IGHV2-70*10
caggtcaccttgaaggagtctggtcctgcgctggtgaaacccacacagaccctcacactg
acctgcaccttctctgggttctcactcagcactagtggaatgcgtgtgagctggatccgt
cagcccccagggaaggccctggagtggattgcacgcattgattgggatgatgataaatac
tacagcacatctctgaagaccaggctcaccatctccaaggacacctccaaaaaccaggtg
gtccttacaatgaccaacatggaccctgtggacacagccacgtattactgtgcacggata
c
>IGHV2-70*11
cgggtcaccttgagggagtctggtcctgcgctggtgaaacccacacagaccctcacactg
acctgcaccttctctgggttctcactcagcactagtggaatgtgtgtgagctggatccgt
cagcccccagggaaggccctggagtggcttgcacgcattgattgggatgatgataaatac
tacagcacatctctgaagaccaggctcaccatctccaaggacacctccaaaaaccaggtg
gtccttacaatgaccaacatggaccctgtggacacagccacgtattactgtgcacggata
c
>IGHV2-70*12
cagatcaccttgaaggagtctggtcctacgctggtgaaacccacacagaccctcacgctg
acctgcaccttctctgggttctcactcagcactagtggaatgtgtgtgagctggatccgt
cagcccccagggaaggccctggagtggcttgcactcattgattgggatgatgataaatac
tacagcacatctctgaagaccaggctcaccatctccaaggacacctccaaaaaccaggtg
gtccttacaatgaccaacatggaccctgtggacacagccacatattactgtgcacacaga
c
>IGHV2-70*13
caggtcaccttgagggagtctggtcctgcgctggtgaaacccacacagaccctcacactg
acctgcaccttctctgggttctcactcagcactagtggaatgtgtgtgagctggatccgt
cagcccccagggaaggccctggagtggcttgcactcattgattgggatgatgataaatac
tacagcacatctctgaagaccaggctcaccatctccaaggacacctccaaaaaccaggtg
gtccttacaatgaccaacatggaccctgtggacacagccacgtattattgtgcacggata
c
>IGHV2-70*15
caggtcaccttgagggagtctggtcctgcgctggtgaaacccacacagaccctcacactg
acctgcaccttctctgggttctcactcagcactagtggaatgtgtgtgagctggatccgt

cagcccccagggaaggccctggagtggcttgcacgcattgattgggatgatgataaatac
tacagcacatctctgaagaccaggctcaccatctccaaggacacctccaaaaaccaggtg
gtccttacaatgaccaacatggaccctgtggacacagccacgtattactgtgcacggata
c
>IGHV2-70*16
caggtcaccttgaaggagtctggtcctgtgctggtgaaacccacacagaccctcacactg
acctgcaccttctctgggttctcactcagcactagtggaatgtgtgtgagctggatccgt
cagcccccagggaaggccctggagtggcttgcacgcattgattgggatgatgataaattc
tacagcacatctctgaagaccaggctcaccatctccaaggacacctccaaaaaccaggtg
gtccttacaatgaccaacatggaccctgtggacacagccacgtattactgtgcacggata
c
>IGHV2-70*17
caggtcaccttgagggagtctggtcctgcgctggtgaaacccacacagaccctcacactg
acctgcaccttctctgggttctcactcagcactagtggaatgtgtgtgagctggatccgt
cagcccccagggaaggccctggagtggcttgcacgcattgattgggatgatgataaattc
tacagcacatctctgaagaccaggctcaccatctccaaggacacctccaaaaaccaggtg
gtccttacaatgaccaacatggaccctgtggacacagccacgtattactgtgcacggata
c
>IGHV2-70*18
caggtcaccttgagggagtctggtcctgcgctggtgaaacccacacagaccctcaccctg
acctgcaccttctctgggttctcactcagcactagtgaaatgtgtgtgagctgggtccgt
cagcccccagggaaggccctggagtggcttgcactcattgattgggatgatgataaatac
tacagcacatctctgaagaccaggctcaccatctccaaggacacctccaaaaaccaggtg
gtccttacaatgaccaacatggaccctgtggacacagccacgtattactgtgcacggata
c
>IGHV2-70*19
caggtcaccttgagggagtctggtcctgcgctggtgaaacccacacagaccctcacactg
acctgcaccttctctgggttctcactcagcactagtggaatgtgtgtgagctgggtccgt
cagcccccagggaaggccctggagtggcttgcactcattgattgggatgatgataaacac
tacagcacatctctgaagaccaggctcaccatctccaaggacacctccaaaaaccaggtg
gtccttacaatgaccaacatggaccctgtggacacagccacgtattactgtgcacggata
c
>IGHV2-70D*04
caggtcaccttgaaggagtctggtcctgcgctggtgaaacccacacagaccctcacactg

acctgcaccttctctgggttctcactcagcactagtggaatgcgtgtgagctggatccgt
cagcccccagggaaggccctggagtggcttgcacgcattgattgggatgatgataaattc
tacagcacatctctgaagaccaggctcaccatctccaaggacacctccaaaaaccaggtg
gtccttacaatgaccaacatggaccctgtggacacagccacgtattactgtgcacggata
c
>IGHV2-70D*14
caggtcaccttgaaggagtctggtcctgcgctggtgaaacccacacagaccctcacactg
acctgcaccttctctgggttctcactcagcactagtggaatgcgtgtgagctggatccgt
cagcccccaggtaaggccctggagtggcttgcacgcattgattgggatgatgataaattc
tacagcacatctctgaagaccaggctcaccatctccaaggacacctccaaaaaccaggtg
gtccttacaatgaccaacatggaccctgtggacacagccacgtattactgtgcacggata
c
>IGHV3-11*01
caggtgcagctggtggagtctgggggaggcttggtcaagcctggagggtccctgagactc
tcctgtgcagcctctggattcaccttcagtgactactacatgagctggatccgccaggct
ccagggaaggggctggagtgggtttcatacattagtagtagtggtagtaccatatactac
gcagactctgtgaagggccgattcaccatctccagggacaacgccaagaactcactgtat
ctgcaaatgaacagcctgagagccgaggacacggccgtgtattactgtgcgagaga
>IGHV3-11*03
caggtgcagctgttggagtctgggggaggcttggtcaagcctggagggtccctgagactc
tcctgtgcagcctctggattcaccttcagtgactactacatgagctggatccgccaggct
ccagggaaggggctggagtgggtttcatacattagtagtagtagtagttacacaaactac
gcagactctgtgaagggccgattcaccatctccagagacaacgccaagaactcactgtat
ctgcaaatgaacagcctgagagccgaggacacggccgtgtattactgtgcgaga
>IGHV3-11*04
caggtgcagctggtggagtctgggggaggcttggtcaagcctggagggtccctgagactc
tcctgtgcagcctctggattcaccttcagtgactactacatgagctggatccgccaggct
ccagggaaggggctggagtgggtttcatacattagtagtagtggtagtaccatatactac
gcagactctgtgaagggccgattcaccatctccagggacaacgccaagaactcactgtat
ctgcaaatgaacagcctgagagccgaggacacggctgtgtattactgtgcgagaga
>IGHV3-11*05
caggtgcagctggtggagtctgggggaggcttggtcaagcctggagggtccctgagactc
tcctgtgcagcctctggattcaccttcagtgactactacatgagctggatccgccaggct
ccagggaaggggctggagtgggtttcatacattagtagtagtagtagttacacaaactac

gcagactctgtgaagggccgattcaccatctccagagacaacgccaagaactcactgtat
ctgcaaatgaacagcctgagagccgaggacacggccgtgtattactgtgcgagaga
>IGHV3-11*06
caggtgcagctggtggagtctgggggaggcttggtcaagcctggagggtccctgagactc
tcctgtgcagcctctggattcaccttcagtgactactacatgagctggatccgccaggct
ccagggaaggggctggagtgggtttcatacattagtagtagtagtagttacacaaactac
gcagactctgtgaagggccgattcaccatctccagagacaacgccaagaactcactgtat
ctgcaaatgaacagcctgagagccgaggacacggctgtgtattactgtgcgagaga
>IGHV3-13*01
gaggtgcagctggtggagtctgggggaggcttggtacagcctggggggtccctgagactc
tcctgtgcagcctctggattcaccttcagtagctacgacatgcactgggtccgccaagct
acaggaaaaggtctggagtgggtctcagctattggtactgctggtgacacatactatcca
ggctccgtgaagggccgattcaccatctccagagaaaatgccaagaactccttgtatctt
caaatgaacagcctgagagccggggacacggctgtgtattactgtgcaagaga
>IGHV3-13*02
gaggtgcatctggtggagtctgggggaggcttggtacagcctgggggggccctgagactc
tcctgtgcagcctctggattcaccttcagtaactacgacatgcactgggtccgccaagct
acaggaaaaggtctggagtgggtctcagccaatggtactgctggtgacacatactatcca
ggctccgtgaaggggcgattcaccatctccagagaaaatgccaagaactccttgtatctt
caaatgaacagcctgagagccggggacacggctgtgtattactgtgcaagaga
>IGHV3-13*03
gaggtgcagctggtggagtctgggggaggcttggtacagcctggggggtccctgagactc
tcctgtgcagcctgtggattcaccttcagtagctacgacatgcactgggtccgccaagct
acaggaaaaggtctggagtgggtctcagctattggtactgctggtgacacatactatcca
ggctccgtgaagggccaattcaccatctccagagaaaatgccaagaactccttgtatctt
caaatgaacagcctgagagccggggacacggctgtgtattactgtgcaaga
>IGHV3-13*04
gaggtgcagctggtggagtctgggggaggcttggtacagcctggggggtccctgagactc
tcctgtgcagcctctggattcaccttcagtagctacgacatgcactgggtccgccaagct
acaggaaaaggtctggaatgggtctcagctattggtactgctggtgacacatactatcca
ggctccgtgaagggccgattcaccatctccagagaaaatgccaagaactccttgtatctt
caaatgaacagcctgagagccggggacacggctgtgtattactgtgcaagaga
>IGHV3-13*05
gaggtgcagctggtggagtctgggggaggcttggtacagcctggggggtccctgagactc

tcctgtgcagcctctggattcaccttcagtagctacgacatgcactgggtccgccaagct
acaggaaaaggtctggagtgggtctcagctattggtactgctggtgacccatactatcca
ggctccgtgaagggccgattcaccatctccagagaaaatgccaagaactccttgtatctt
caaatgaacagcctgagagccggggacacggctgtgtattactgtgcaagaga
>IGHV3-15*01
gaggtgcagctggtggagtctgggggaggcttggtaaagcctggggggtcccttagactc
tcctgtgcagcctctggattcactttcagtaacgcctggatgagctgggtccgccaggct
ccagggaaggggctggagtgggttggccgtattaaaagcaaaactgatggtgggacaaca
gactacgctgcacccgtgaaaggcagattcaccatctcaagagatgattcaaaaaacacg
ctgtatctgcaaatgaacagcctgaaaaccgaggacacagccgtgtattactgtaccaca
ga
>IGHV3-15*02
gaggtgcagctggtggagtctgggggagccttggtaaagcctggggggtcccttagactc
tcctgtgcagcctctggattcactttcagtaacgcctggatgagctgggtccgccaggct
ccagggaaggggctggagtgggttggccgtattaaaagcaaaactgatggtgggacaaca
gactacgctgcacccgtgaaaggcagattcaccatctcaagagatgattcaaaaaacacg
ctgtatctgcaaatgaacagcctgaaaaccgaggacacagccgtgtattactgtaccaca
ga
>IGHV3-15*03
gaggtgcagctggtggagtctgccggagccttggtacagcctggggggtcccttagactc
tcctgtgcagcctctggattcacttgcagtaacgcctggatgagctgggtccgccaggct
ccagggaaggggctggagtgggttggccgtattaaaagcaaagctaatggtgggacaaca
gactacgctgcacctgtgaaaggcagattcaccatctcaagagttgattcaaaaaacacg
ctgtatctgcaaatgaacagcctgaaaaccgaggacacagccgtgtattactgtaccaca
ga
>IGHV3-15*04
gaggtgcagctggtggagtctgggggaggcttggtaaagcctggggggtcccttagactc
tcctgtgcagcctctggattcactttcagtaacgcctggatgagctgggtccgccaggct
ccagggaaggggctggagtgggttggccgtattgaaagcaaaactgatggtgggacaaca
gactacgctgcacccgtgaaaggcagattcaccatctcaagagatgattcaaaaaacacg
ctgtatctgcaaatgaacagcctgaaaaccgaggacacagccgtgtattactgtaccaca
ga
>IGHV3-15*05
gaggtgcagctggtggagtctgggggaggcttggtaaagcctggggggtcccttagactc

tcctgtgcagcctctggattcactttcagtaacgcctggatgagctgggtccgccaggct
ccagggaaggggctggagtgggttggccgtattaaaagcaaaactgatggtgggacaaca
gactacgctgcacccgtgaaaggcagattcaccatctcaagagatgattcaaaaaacacg
ctgtatctgcaaatgaacagtctgaaaaccgaggacacagccgtgtattactgtaccaca
ga
>IGHV3-15*06
gaggtgcagctggtggagtctgggggaggcttggtaaagcctggggggtcccttagactc
tcctgtgcagcctctggattcactttcagtaacgcctggatgagctgggtccgccaggct
ccagggaaggggctggagtgggtcggccgtattaaaagcaaaactgatggtgggacaaca
aactacgctgcacccgtgaaaggcagattcaccatctcaagagatgattcaaaaaacacg
ctgtatctgcaaatgaacagcctgaaaaccgaggacacagccgtgtattactgtaccaca
ga
>IGHV3-15*07
gaggtgcagctggtggagtctgggggaggcttggtaaagcctggggggtcccttagactc
tcctgtgcagcctctggtttcactttcagtaacgcctggatgaactgggtccgccaggct
ccagggaaggggctggagtgggtcggccgtattaaaagcaaaactgatggtgggacaaca
gactacgctgcacccgtgaaaggcagattcaccatctcaagagatgattcaaaaaacacg
ctgtatctgcaaatgaacagcctgaaaaccgaggacacagccgtgtattactgtaccaca
ga
>IGHV3-15*08
gaggtgcagctggtggagtctgcgggaggcttggtacagcctggggggtcccttagactc
tcctgtgcagcctctggattcacttgcagtaacgcctggatgagctgggtccgccaggct
ccagggaaggggctggagtgggttggctgtattaaaagcaaagctaatggtgggacaaca
gactacgctgcacctgtgaaaggcagattcaccatctcaagagatgattcaaaaaacacg
ctgtatctgcaaatgatcagcctgaaaaccgaggacacggccgtgtattactgtaccaca
gg
>IGHV3-20*01
gaggtgcagctggtggagtctgggggaggtgtggtacggcctggggggtccctgagactc
tcctgtgcagcctctggattcacctttgatgattatggcatgagctgggtccgccaagct
ccagggaaggggctggagtgggtctctggtattaattggaatggtggtagcacaggttat
gcagactctgtgaagggccgattcaccatctccagagacaacgccaagaactccctgtat
ctgcaaatgaacagtctgagagccgaggacacggccttgtatcactgtgcgagaga
>IGHV3-20*04
gaggtgcagctggtggagtctgggggaggtgtggtacggcctggggggtccctgagactc

tcctgtgcagcctctggattcacctttgatgattatggcatgagctgggtccgccaagct
ccagggaaggggctggagtgggtctctggtattaattggaatggtggtagcacaggttat
gcagactctgtgaagggccgattcaccatctccagagacaacgccaagaactccctgtat
ctgcaaatgaacagtctgagagccgaggacacggccttgtattactgtgcgagaga
>IGHV3-21*01
gaggtgcagctggtggagtctgggggaggcctggtcaagcctggggggtccctgagactc
tcctgtgcagcctctggattcaccttcagtagctatagcatgaactgggtccgccaggct
ccagggaaggggctggagtgggtctcatccattagtagtagtagtagttacatatactac
gcagactcagtgaagggccgattcaccatctccagagacaacgccaagaactcactgtat
ctgcaaatgaacagcctgagagccgaggacacggctgtgtattactgtgcgagaga
>IGHV3-21*02
gaggtgcaactggtggagtctgggggaggcctggtcaagcctggggggtccctgagactc
tcctgtgcagcctctggattcaccttcagtagctatagcatgaactgggtccgccaggct
ccagggaaggggctggagtgggtctcatccattagtagtagtagtagttacatatactac
gcagactcagtgaagggccgattcaccatctccagagacaacgccaagaactcactgtat
ctgcaaatgaacagcctgagagccgaggacacggctgtgtattactgtgcgagaga
>IGHV3-21*03
gaggtgcagctggtggagtctgggggaggcctggtcaagcctggggggtccctgagactc
tcctgtgcagcctctggattcaccttcagtagctatagcatgaactgggtccgccaggct
ccagggaaggggctggagtgggtctcatccattagtagtagtagtagttacatatactac
gcagactcagtgaagggccgattcaccatctccagagacaacgccaagaactcactgtat
ctgcaaatgaacagcctgagagccgaggacacagctgtgtattactgtgcgagaga
>IGHV3-21*04
gaggtgcagctggtggagtctgggggaggcctggtcaagcctggggggtccctgagactc
tcctgtgcagcctctggattcaccttcagtagctatagcatgaactgggtccgccaggct
ccagggaaggggctggagtgggtctcatccattagtagtagtagtagttacatatactac
gcagactcagtgaagggccgattcaccatctccagagacaacgccaagaactcactgtat
ctgcaaatgaacagcctgagagccgaggacacggccgtgtattactgtgcgagaga
>IGHV3-21*05
gaggtgcagctggtggagtctgggggaggcctggtcaagcctggggggtccctgagactc
tcctgtgcagcctctggattcaccttcagtagctatagcatgaactgggtccgccaggct
ccagggaaggggctggagtgggtttcatacattagtagtagtagtagttacatatactac
gcagactcagtgaagggccgattcaccatctccagagacaacgccaagaactcactgtat
ctgcaaatgaacagcctgagagccgaggacacggctgtgtattactgtgcgagaga

>IGHV3-23*01
gaggtgcagctgttggagtctgggggaggcttggtacagcctggggggtccctgagactc
tcctgtgcagcctctggattcacctttagcagctatgccatgagctgggtccgccaggct
ccagggaaggggctggagtgggtctcagctattagtggtagtggtggtagcacatactac
gcagactccgtgaagggccggttcaccatctccagagacaattccaagaacacgctgtat
ctgcaaatgaacagcctgagagccgaggacacggccgtatattactgtgcgaaaga
>IGHV3-23*02
gaggtgcagctgttggagtctgggggaggcttggtacagcctggggggtccctgagactc
tcctgtgcagcctctggattcacctttagcagctatgccatgagctgggtccgccaggct
ccagggaaggggctggagtgggtctcagctattagtggtagtggtggtagcacatactac
ggagactccgtgaagggccggttcaccatctcaagagacaattccaagaacacgctgtat
ctgcaaatgaacagcctgagagccgaggacacggccgtatattactgtgcgaaaga
>IGHV3-23*03
gaggtgcagctgttggagtctgggggaggcttggtacagcctggggggtccctgagactc
tcctgtgcagcctctggattcacctttagcagctatgccatgagctgggtccgccaggct
ccagggaaggggctggagtgggtctcagttatttatagcggtggtagtagcacatactat
gcagactccgtgaagggccggttcaccatctccagagataattccaagaacacgctgtat
ctgcaaatgaacagcctgagagccgaggacacggccgtatattactgtgcgaaaga
>IGHV3-23*04
gaggtgcagctggtggagtctgggggaggcttggtacagcctggggggtccctgagactc
tcctgtgcagcctctggattcacctttagcagctatgccatgagctgggtccgccaggct
ccagggaaggggctggagtgggtctcagctattagtggtagtggtggtagcacatactac
gcagactccgtgaagggccggttcaccatctccagagacaattccaagaacacgctgtat
ctgcaaatgaacagcctgagagccgaggacacggccgtatattactgtgcgaaaga
>IGHV3-23*05
gaggtgcagctgttggagtctgggggaggcttggtacagcctggggggtccctgagactc
tcctgtgcagcctctggattcacctttagcagctatgccatgagctgggtccgccaggct
ccagggaaggggctggagtgggtctcagctatttatagcagtggtagtagcacatactat
gcagactccgtgaagggccggttcaccatctccagagacaattccaagaacacgctgtat
ctgcaaatgaacagcctgagagccgaggacacggccgtatattactgtgcgaaa
>IGHV3-23D*01
gaggtgcagctgttggagtctgggggaggcttggtacagcctggggggtccctgagactc
tcctgtgcagcctctggattcacctttagcagctatgccatgagctgggtccgccaggct
ccagggaaggggctggagtgggtctcagctattagtggtagtggtggtagcacatactac

gcagactccgtgaagggccggttcaccatctccagagacaattccaagaacacgctgtat
ctgcaaatgaacagcctgagagccgaggacacggccgtatattactgtgcgaaaga
>IGHV3-30*01
caggtgcagctggtggagtctgggggaggcgtggtccagcctgggaggtccctgagactc
tcctgtgcagcctctggattcaccttcagtagctatgctatgcactgggtccgccaggct
ccaggcaaggggctagagtgggtggcagttatatcatatgatggaagtaataaatactac
gcagactccgtgaagggccgattcaccatctccagagacaattccaagaacacgctgtat
ctgcaaatgaacagcctgagagctgaggacacggctgtgtattactgtgcgagaga
>IGHV3-30*02
caggtgcagctggtggagtctgggggaggcgtggtccagcctgggggtccctgagactc
tcctgtgcagcgtctggattcaccttcagtagctatggcatgcactgggtccgccaggct
ccaggcaaggggctggagtgggtggcatttatacggtatgatggaagtaataaatactat
gcagactccgtgaagggccgattcaccatctccagagacaattccaagaacacgctgtat
ctgcaaatgaacagcctgagagctgaggacacggctgtgtattactgtgcgaaaga
>IGHV3-30*03
caggtgcagctggtggagtctgggggaggcgtggtccagcctgggaggtccctgagactc
tcctgtgcagcctctggattcaccttcagtagctatggcatgcactgggtccgccaggct
ccaggcaaggggctggagtgggtggcagttatatcatatgatggaagtaataaatactat
gcagactccgtgaagggccgattcaccatctccagagacaattccaagaacacgctgtat
ctgcaaatgaacagcctgagagctgaggacacggctgtgtattactgtgcgagaga
>IGHV3-30*04
caggtgcagctggtggagtctgggggaggcgtggtccagcctgggaggtccctgagactc
tcctgtgcagcctctggattcaccttcagtagctatgctatgcactgggtccgccaggct
ccaggcaaggggctggagtgggtggcagttatatcatatgatggaagtaataaatactac
gcagactccgtgaagggccgattcaccatctccagagacaattccaagaacacgctgtat
ctgcaaatgaacagcctgagagctgaggacacggctgtgtattactgtgcgagaga
>IGHV3-30*05
caggtgcagctggtggagtctgggggaggcgtggtccagcctgggaggtccctgagactc
tcctgtgcagcctctggattcaccttcagtagctatggcatgcactgggtccgccaggct
ccaggcaaggggctagagtgggtggcagttatatcatatgatggaagtaataaatactac
gcagactccgtgaagggccgattcaccatctccagagacaattccaagaacacgctgtat
ctgcaaatgaacagcctgagactgagggcacggctgtgtattactgtgcgagaga
>IGHV3-30*06
caggtgcagctggtggagtctgggggaggcgtggtccagcctgggaggtccctgagactc

tcctgtgcagcgtctggattcaccttcagtagctatggcatgcactgggtccgccaggct
ccaggcaaggggctagagtgggtggcagttatatcatatgatggaagtaataaatactac
gcagactccgtgaagggccgattcaccatctccagagacaattccaagaacacgctgtat
ctgcaaatgaacagcctgagagctgaggacacggctgtgtattactgtgcgagaga
>IGHV3-30*07
caggtgcagctggtggagtctgggggaggcgtggtccagcctgggaggtccctgagactc
tcctgtgcagcctctggattcaccttcagtagctatgctatgcactgggtccgccaggct
ccaggcaaggggctagagtgggtggcagttatatcatatgatggaagtaataaatactac
gcagactccgtgaagggccgattcaccatctccagagacaattccaagaacacgctgtat
ctgcaaatgaacagcctgagagccgaggacacggctgtgtattactgtgcgagaga
>IGHV3-30*08
caggtgcagctggtggactctgggggaggcgtggtccagcctgggaggtccctgagactc
tcctgtgcagcctctgcattcaccttcagtagctatgctatgcactgggtccgccaggct
ccaggcaaggggctagagtgggtggcagttatatcatatgatggaagtaataaatactac
gcagactccgtgaagggccgattcaccatctccagagacaattccaagaacacgctgtat
ctgcaaatgaacagcctgagagctgaggacacggctgtgtattactgtgcgaga
>IGHV3-30*09
caggtgcagctggtggagtctgggggaggcgtggtccagcctgggaggtccctgagactc
tcctgtgcagcctctggattcaccttcagtagctatgctatgcactgggtccgccaggct
ccaggcaaggggctggagtgggtggcagttatatcatatgatggaagtaataaatactac
gcagactccgtgaagggccgattcgccatctccagagacaattccaagaacacgctgtat
ctgcaaatgaacagcctgagagctgaggacacggctgtgtattactgtgcgagaga
>IGHV3-30*10
caggtgcagctggtggagtctgggggaggcgtggtccagcctgggaggtccctgagactc
tcctgtgcagcctctggattcaccttcagtagctatgctatgcactgggtccgccaggct
ccaggcaaggggctagagtgggtggcagttatatcatatgatggaagtaataaatactac
acagactccgtgaagggccgattcaccatctccagagacaattccaagaacacgctgtat
ctgcaaatgaacagcctgagagctgaggacacggctgtgtattactgtgcgagaga
>IGHV3-30*11
caggtgcagctggtggagtctgggggaggcgtggtccagcctgggaggtccctgagactc
tcctgtgcagcgtctggattcaccttcagtagctatgctatgcactgggtccgccaggct
ccaggcaaggggctagagtgggtggcagttatatcatatgatggaagtaataaatactac
gcagactccgtgaagggccgattcaccatctccagagacaattccaagaacacgctgtat
ctgcaaatgaacagcctgagagctgaggacacggctgtgtattactgtgcgagaga

>IGHV3-30*12
caggtgcagctggtggagtctgggggggcgtggtccagcctgggaggtccctgagactc
tcctgtgcagcgtctggattcaccttcagtagctatggcatgcactgggtccgccaggct
ccaggcaaggggctagagtgggtggcagttatatcatatgatggaagtaataaatactac
gcagactccgtgaagggccgattcaccatctccagagacaattccaagaacacgctgtat
ctgcaaatgaacagcctgagagccgaggacacggctgtgtattactgtgcgagaga
>IGHV3-30*13
caggtgcagctggtggagtctgggggaggcgtggtccagcctgggaggtccctgagactc
tcctgtgcagcctctggattcaccttcagtagctatggcatgcactgggtccgccaggct
ccaggcaaggggctagagtgggtggcagttatatcatatgatggaagtaataaatactac
gcagactccgtgaagggccgattcaccatctccagagacaattccaagaacaggctgtat
ctgcaaatgaacagcctgagagctgaggacacggctgtgtattactgtgcgagaga
>IGHV3-30*14
caggtgcagctggtggagtctgggggaggcgtggtccagcctgggaggtccctgagactc
tcctgtgcagcctctggattcaccttcagtagctatgctatgcactgggtccgccaggct
ccaggcaaggggctggagtgggtggcagttatatcatatgatggaagtaataaatactac
gcagactccgtgaagggccgattcaccatctccagagacaattccaagaacacgctgtat
cttcaaatgaacagcctgagagctgaggacacggctgtgtattactgtgcgagaga
>IGHV3-30*15
caggtgcagctggtggagtctgggggaggcgtggtccagcctgggaggtccctgagactc
tcctgtgcagcctctggattcaccttcagtagctatgctatgcactgggtccgccaggct
ccaggcaaggggctagagtgggtggcagttatatcatatgatggaagtaataaatactac
gcagactccgtgaagggccgattcaccatctccagagacaattccaagaacacgctgtat
ctgcaaatgagcagcctgagagctgaggacacggctgtgtattactgtgcgagaga
>IGHV3-30*16
caggtgcagctggtggagtctgggggaggcgtggtccagcctgggaggtccctgagactc
tcctgtgcagcctctggattcaccttcagtagctatgctatgcactgggtccgccaggcc
ccaggcaaggggctagagtgggtggcagttatatcatatgatggaagtaataaatactac
gcagactccgtgaagggccgattcaccatctccagagacaattccaagaacacgctgtat
ctgcaaatgaacagcctgagagctgaggacacggctgtgtattactgtgcgagaga
>IGHV3-30*17
caggtgcagctggtggagtctgggggaggcgtggtccagcctgggaggtccctgagactc
tcctgtgcagcctctggattcaccttcagtagctatgctatgcactgggtccgccaggct
ccgggcaaggggctagagtgggtggcagttatatcatatgatggaagtaataaatactac

```
gcagactccgtgaagggccgattcaccatctccagagacaattccaagaacacgctgtat
ctgcaaatgaacagcctgagagctgaggacacggctgtgtattactgtgcgagaga
>IGHV3-30*18
caggtgcagctggtggagtctgggggaggcgtggtccagcctgggaggtccctgagactc
tcctgtgcagcctctggattcaccttcagtagctatggcatgcactgggtccgccaggct
ccaggcaaggggctggagtgggtggcagttatatcatatgatggaagtaataaatactat
gcagactccgtgaagggccgattcaccatctccagagacaattccaagaacacgctgtat
ctgcaaatgaacagcctgagagctgaggacacggctgtgtattactgtgcgaaaga
>IGHV3-30*19
caggtgcagctggtggagtctgggggaggcgtggtccagcctgggaggtccctgagactc
tcctgtgcagcgtctggattcaccttcagtagctatggcatgcactgggtccgccaggct
ccaggcaaggggctggagtgggtggcagttatatcatatgatggaagtaataaatactac
gcagactccgtgaagggccgattcaccatctccagagacaattccaagaacacgctgtat
ctgcaaatgaacagcctgagagctgaggacacggctgtgtattactgtgcgagaga
>IGHV3-30-3*01
caggtgcagctggtggagtctgggggaggcgtggtccagcctgggaggtccctgagactc
tcctgtgcagcctctggattcaccttcagtagctatgctatgcactgggtccgccaggct
ccaggcaaggggctggagtgggtggcagttatatcatatgatggaagcaataaatactac
gcagactccgtgaagggccgattcaccatctccagagacaattccaagaacacgctgtat
ctgcaaatgaacagcctgagagctgaggacacggctgtgtattactgtgcgagaga
>IGHV3-30-3*02
caggtgcagctggtggagtctgggggaggcgtggtccagcctgggaggtccctgagactc
tcctgtgcagcgtctggattcaccttcagtagctatgctatgcactgggtccgccaggct
ccaggcaaggggctggagtgggtggcagttatatcatatgatggaagcaataaatactac
gcagactccgtgaagggccgattcaccatctccagagacaattccaagaacacgctgtat
ctgcaaatgaacagcctgagagctgaggacacggctgtgtattactgtgcgaaaga
>IGHV3-30-3*03
caggtgcagctggtggagtctgggggaggcgtggtccagcctgggaggtccctgagactc
tcctgtgcagcctctggattcaccttcagtagctatgctatgcactgggtccgccaggct
ccaggcaaggggctggagtgggtggcagttatatcatatgatggaagtaataaatactac
gcagactccgtgaagggccgattcaccatctccagagacaattccaagaacacgctgtat
ctgcaaatgaacagcctgagagctgaggacacggctgtgtattactgtgcgagaga
>IGHV3-30-5*01
caggtgcagctggtggagtctgggggaggcgtggtccagcctgggaggtccctgagactc
```

tcctgtgcagcctctggattcaccttcagtagctatggcatgcactgggtccgccaggct
ccaggcaaggggctggagtgggtggcagttatatcatatgatggaagtaataaatactat
gcagactccgtgaagggccgattcaccatctccagagacaattccaagaacacgctgtat
ctgcaaatgaacagcctgagagctgaggacacggctgtgtattactgtgcgaaaga
>IGHV3-30-5*02
caggtgcagctggtggagtctgggggaggcgtggtccagcctgggggtccctgagactc
tcctgtgcagcgtctggattcaccttcagtagctatggcatgcactgggtccgccaggct
ccaggcaaggggctggagtgggtggcatttatacggtatgatggaagtaataaatactat
gcagactccgtgaagggccgattcaccatctccagagacaattccaagaacacgctgtat
ctgcaaatgaacagcctgagagctgaggacacggctgtgtattactgtgcgaaaga
>IGHV3-33*01
caggtgcagctggtggagtctgggggaggcgtggtccagcctgggaggtccctgagactc
tcctgtgcagcgtctggattcaccttcagtagctatggcatgcactgggtccgccaggct
ccaggcaaggggctggagtgggtggcagttatatggtatgatggaagtaataaatactat
gcagactccgtgaagggccgattcaccatctccagagacaattccaagaacacgctgtat
ctgcaaatgaacagcctgagagccgaggacacggctgtgtattactgtgcgagaga
>IGHV3-33*02
caggtacagctggtggagtctgggggaggcgtggtccagcctgggaggtccctgagactc
tcctgtgcagcgtctggattcaccttcagtagctatggcatgcactgggtccgccaggct
ccaggcaaggggctggagtgggtggcagttatatggtatgatggaagtaataaatactat
gcagactccgcgaagggccgattcaccatctccagagacaattccacgaacacgctgttt
ctgcaaatgaacagcctgagagccgaggacacggctgtgtattactgtgcgagaga
>IGHV3-33*03
caggtgcagctggtggagtctgggggaggcgtggtccagcctgggaggtccctgagactc
tcctgtgcagcgtctggattcaccttcagtagctatggcatgcactgggtccgccaggct
ccaggcaaggggctggagtgggtggcagttatatggtatgatggaagtaataaatactat
gcagactccgtgaagggccgattcaccatctccagagacaactccaagaacacgctgtat
ctgcaaatgaacagcctgagagccgaggacacggctgtgtattactgtgcgaaaga
>IGHV3-33*04
caggtgcagctggtggagtctgggggaggcgtggtccagcctgggaggtccctgagactc
tcctgtgcagcgtctggattcaccttcagtagctatggcatgcactgggtccgccaggct
ccaggcaaggggctagagtgggtggcagttatatggtatgacggaagtaataaatactat
gcagactccgtgaagggccgattcaccatctccagagacaattccaagaacacgctgtat
ctgcaaatgaacagcctgagagccgaggacacggctgtgtattactgtgcgagaga

>IGHV3-33*05
caggtgcagctggtggagtctgggggaggcgtggtccagcctgggaggtccctgagactc
tcctgtgcagcgtctggattcaccttcagtagctatggcatgcactgggtccgccaggct
ccaggcaaggggctggagtgggtggcagttatatcatatgatggaagtaataaatactat
gcagactccgtgaagggccgattcaccatctccagagacaattccaagaacacgctgtat
ctgcaaatgaacagcctgagagccgaggacacggctgtgtattactgtgcgagaga
>IGHV3-33*06
caggtgcagctggtggagtctgggggaggcgtggtccagcctgggaggtccctgagactc
tcctgtgcagcgtctggattcaccttcagtagctatggcatgcactgggtccgccaggct
ccaggcaaggggctggagtgggtggcagttatatggtatgatggaagtaataaatactat
gcagactccgtgaagggccgattcaccatctccagagacaattccaagaacacgctgtat
ctgcaaatgaacagcctgagagccgaggacacggctgtgtattactgtgcgaaaga
>IIGHV3-33*07
caggtgcagctggtggagtctgggggacgcgtggtccagcctgggaggtccctgagactc
tcctgtgcagcgtctggattcaccttcagtaggtatggcatgtactgggtccgccaggct
ccaggcaaggggctggagtgggtggcagttatatggtatgatggaagtaataaatactat
gcagactccgtgaagggccgattcaccatctccagagacaattccaagaacacgctgtat
ctgcaaatgaacagcctgagagccgaggacacggctgtgtattactgtgcgagaga
>IGHV3-43*01
gaagtgcagctggtggagtctgggggagtcgtggtacagcctggggggtccctgagactc
tcctgtgcagcctctggattcacctttgatgattataccatgcactgggtccgtcaagct
ccgggaagggtctggagtgggtctctcttattagttgggatggtggtagcacatactat
gcagactctgtgaagggccgattcaccatctccagagacaacagcaaaaactccctgtat
ctgcaaatgaacagtctgagaactgaggacaccgccttgtattactgtgcaaaagata
>IGHV3-43*02
gaagtgcagctggtggagtctgggggaggcgtggtacagcctggggggtccctgagactc
tcctgtgcagcctctggattcacctttgatgattatgccatgcactgggtccgtcaagct
ccagggaagggtctggagtgggtctctcttattagtggggatggtggtagcacatactat
gcagactctgtgaagggccgattcaccatctccagagacaacagcaaaaactccctgtat
ctgcaaatgaacagtctgagaactgaggacaccgccttgtattactgtgcaaaagata
>IGHV3-43D*03
gaagtgcagctggtggagtctgggggagtcgtggtacagcctggggggtccctgagactc
tcctgtgcagcctctggattcacctttgatgattatgccatgcactgggtccgtcaagct
ccggggaagggtctggagtgggtctctcttattagttgggatggtggtagcacctactat

gcagactctgtgaagggtcgattcaccatctccagagacaacagcaaaaactccctgtat
ctgcaaatgaacagtctgagagctgaggacaccgccttgtattactgtgcaaaagata
>IGHV3-43D*04
gaagtgcagctggtggagtctgggggagtcgtggtacagcctggggggtccctgagactc
tcctgtgcagcctctggattcacctttgatgattatgccatgcactgggtccgtcaagct
ccggggaagggtctggagtgggtctctcttattagttgggatggtggtagcacatactat
gcagactctgtgaagggtcgattcaccatctccagagacaacagcaaaaactccctgtat
ctgcaaatgaacagtctgagagctgaggacaccgccttgtattactgtgcaaaagata
>IGHV3-48*01
gaggtgcagctggtggagtctgggggaggcttggtacagcctggggggtccctgagactc
tcctgtgcagcctctggattcaccttcagtagctatagcatgaactgggtccgccaggct
ccagggaaggggctggagtgggtttcatacattagtagtagtagtagtaccatatactac
gcagactctgtgaagggccgattcaccatctccagagacaatgccaagaactcactgtat
ctgcaaatgaacagcctgagagccgaggacacggctgtgtattactgtgcgagaga
>IGHV3-48*02
gaggtgcagctggtggagtctgggggaggcttggtacagcctggggggtccctgagactc
tcctgtgcagcctctggattcaccttcagtagctatagcatgaactgggtccgccaggct
ccagggaaggggctggagtgggtttcatacattagtagtagtagtagtaccatatactac
gcagactctgtgaagggccgattcaccatctccagagacaatgccaagaactcactgtat
ctgcaaatgaacagcctgagagacgaggacacggctgtgtattactgtgcgagaga
>IGHV3-48*03
gaggtgcagctggtggagtctgggggaggcttggtacagcctggagggtccctgagactc
tcctgtgcagcctctggattcaccttcagtagttatgaaatgaactgggtccgccaggct
ccagggaaggggctggagtgggtttcatacattagtagtagtggtagtaccatatactac
gcagactctgtgaagggccgattcaccatctccagagacaacgccaagaactcactgtat
ctgcaaatgaacagcctgagagccgaggacacggctgtttattactgtgcgagaga
>IGHV3-48*04
gaggtgcagctggtggagtctgggggaggcttggtacagcctggggggtccctgagactc
tcctgtgcagcctctggattcaccttcagtagctatagcatgaactgggtccgccaggct
ccagggaaggggctggagtgggtttcatacattagtagtagtagtagtaccatatactac
gcagactctgtgaagggccgattcaccatctccagagacaacgccaagaactcactgtat
ctgcaaatgaacagcctgagagccgaggacacggctgtgtattactgtgcgagaga
>IGHV3-49*01
gaggtgcagctggtggagtctgggggaggcttggtacagccagggcggtccctgagactc

tcctgtacagcttctggattcacctttggtgattatgctatgagctggttccgccaggct
ccagggaaggggctggagtgggtaggtttcattagaagcaaagcttatggtgggacaaca
gaatacaccgcgtctgtgaaaggcagattcaccatctcaagagatggttccaaaagcatc
gcctatctgcaaatgaacagcctgaaaaccgaggacacagccgtgtattactgtactaga
ga
>IGHV3-49*02
gaggtgcagctggtggagtctgggggaggcttggtacagccagggccgtccctgagactc
tcctgtacagcttctggattcacctttgggtattatcctatgagctgggtccgccaggct
ccagggaaggggctggagtgggtaggtttcattagaagcaaagcttatggtgggacaaca
gaatacgccgcgtctgtgaaaggcagattcaccatctcaagagatgattccaaaagcatc
gcctatctgcaaatgaacagcctgaaaaccgaggacacagccgtgtattactgtactaga
ga
>IGHV3-49*03
gaggtgcagctggtggagtctgggggaggcttggtacagccagggcggtccctgagactc
tcctgtacagcttctggattcacctttggtgattatgctatgagctggttccgccaggct
ccagggaaggggctggagtgggtaggtttcattagaagcaaagcttatggtgggacaaca
gaatacgccgcgtctgtgaaaggcagattcaccatctcaagagatgattccaaaagcatc
gcctatctgcaaatgaacagcctgaaaaccgaggacacagccgtgtattactgtactaga
ga
>IGHV3-49*04
gaggtgcagctggtggagtctgggggaggcttggtacagccagggcggtccctgagactc
tcctgtacagcttctggattcacctttggtgattatgctatgagctgggtccgccaggct
ccagggaaggggctggagtgggtaggtttcattagaagcaaagcttatggtgggacaaca
gaatacgccgcgtctgtgaaaggcagattcaccatctcaagagatgattccaaaagcatc
gcctatctgcaaatgaacagcctgaaaaccgaggacacagccgtgtattactgtactaga
ga
>IGHV3-49*05
gaggtgcagctggtggagtctgggggaggcttggtaaagccagggcggtccctgagactc
tcctgtacagcttctggattcacctttggtgattatgctatgagctggttccgccaggct
ccagggaaggggctggagtgggtaggtttcattagaagcaaagcttatggtgggacaaca
gaatacgccgcgtctgtgaaaggcagattcaccatctcaagagatgattccaaaagcatc
gcctatctgcaaatgaacagcctgaaaaccgaggacacagccgtgtattactgtactaga
ga
>IGHV3-53*01

gaggtgcagctggtggagtctggaggaggcttgatccagcctggggggtccctgagactc
tcctgtgcagcctctgggttcaccgtcagtagcaactacatgagctgggtccgccaggct
ccagggaaggggctggagtgggtctcagttatttatagcggtggtagcacatactacgca
gactccgtgaagggccgattcaccatctccagagacaattccaagaacacgctgtatctt
caaatgaacagcctgagagccgaggacacggccgtgtattactgtgcgagaga
>IGHV3-53*02
gaggtgcagctggtggagactggaggaggcttgatccagcctggggggtccctgagactc
tcctgtgcagcctctgggttcaccgtcagtagcaactacatgagctgggtccgccaggct
ccagggaaggggctggagtgggtctcagttatttatagcggtggtagcacatactacgca
gactccgtgaagggccgattcaccatctccagagacaattccaagaacacgctgtatctt
caaatgaacagcctgagagccgaggacacggccgtgtattactgtgcgagaga
>IGHV3-53*03
gaggtgcagctggtggagtctggaggaggcttgatccagcctggggggtccctgagactc
tcctgtgcagcctctgggttcaccgtcagtagcaactacatgagctgggtccgccagcct
ccagggaaggggctggagtgggtctcagttatttatagcggtggtagcacatactacgca
gactctgtgaagggccgattcaccatctccagagacaattccaagaacacgctgtatctt
caaatgaacagcctgagagccgaggacacggccgtgtattactgtgctaggga
>IGHV3-53*04
gaggtgcagctggtggagtctggaggaggcttggtccagcctggggggtccctgagactc
tcctgtgcagcctctgggttcaccgtcagtagcaactacatgagctgggtccgccaggct
ccagggaaggggctggagtgggtctcagttatttatagcggtggtagcacatactacgca
gactccgtgaagggccgattcaccatctccagacacaattccaagaacacgctgtatctt
caaatgaacagcctgagagctgaggacacggccgtgtattactgtgcgagaga
>IGHV3-53*05
gaggtgcagctggtggagactggaggaggcttgatccagcctggggggtccctgagactc
tcctgtgcagcctctgggttcaccgtcagtagcaactacatgagctgggtccgccaggct
ccagggaaggggctggagtgggtctcagttatttatagcggtggtagcacatactacgca
gactccgtgaagggccgattcaccatctccagagacaattccaagaacacgctgtatctt
caaatgaacagcctgagagctgaggacacggccgtgtattactgtgcgagaga
>IGHV3-62*04
gaggtgcagctggtgaagtctggaggaggcttggtacagcctggggggtccctgagactc
tcctgtgcagcctctggattcaccttcagtagctctgctatgcactgggtccgccaggct
ccaagaaagggtttggagtgggtctcagttattagtacaagtggtgataccgtactctac
acagactctgtgaagggccgattcaccatctccagagacaatgcccagaattcactgtct

ctgcaaatgaacagcctgagagccgaggacatggctgtgtattactgtgtgaaaga
>IGHV3-64*01
gaggtgcagctggtggagtctgggggaggcttggtccagcctggggggtccctgagactc
tcctgtgcagcctctggattcaccttcagtagctatgctatgcactgggtccgccaggct
ccagggaagggactggaatatgtttcagctattagtagtaatgggggtagcacatattat
gcaaactctgtgaagggcagattcaccatctccagagacaattccaagaacacgctgtat
cttcaaatgggcagcctgagagctgaggacatggctgtgtattactgtgcgagaga
>IGHV3-64*02
gaggtgcagctggtggagtctggggaaggcttggtccagcctggggggtccctgagactc
tcctgtgcagcctctggattcaccttcagtagctatgctatgcactgggtccgccaggct
ccagggaagggactggaatatgtttcagctattagtagtaatgggggtagcacatattat
gcagactctgtgaagggcagattcaccatctccagagacaattccaagaacacgctgtat
cttcaaatgggcagcctgagagctgaggacatggctgtgtattactgtgcgagaga
>IGHV3-64*03
gaggtgcagctggtggagtctgggggaggcttggtccagcctggggggtccctgagactc
tcctgttcagcctctggattcaccttcagtagctatgctatgcactgggtccgccaggct
ccagggaagggactggaatatgtttcagctattagtagtaatgggggtagcacatactac
gcagactcagtgaagggcagattcaccatctccagagacaattccaagaacacgctgtat
gtccaaatgagcagtctgagagctgaggacacggctgtgtattactgtgtgaaaga
>IGHV3-64*04
caggtgcagctggtggagtctgggggaggcttggtccagcctggggggtccctgagactc
tcctgttcagcctctggattcaccttcagtagctatgctatgcactgggtccgccaggct
ccagggaagggactggaatatgtttcagctattagtagtaatgggggtagcacatactac
gcagactcagtgaagggcagattcaccatctccagagacaattccaagaacacgctgtat
ctgcaaatgaacagcctgagagctgaggacacggctgtgtattactgtgcgagaga
>IGHV3-64*05
gaggtgcagctggtggagtctgggggaggcttggtccagcctggggggtccctgagactc
tcctgttcagcctctggattcaccttcagtagctatgctatgcactgggtccgccaggct
ccagggaagggactggaatatgtttcagctattagtagtaatgggggtagcacatactac
gcagactcagtgaagggcagattcaccatctccagagacaattccaagaacacgctgtat
gttcaaatgagcagtctgagagctgaggacacggctgtgtattactgtgtgaaaga
>IGHV3-64*07
gaggtgcagctggtggagtctgggggaggcttggtccagcctggggggtccctgagactc
tcctgtgcagcctctggattcaccttcagtagctatgctatgcactgggtccgccaggct

ccagggaagggactggaatatgtttcagctattagtagtaatgggggtagcacatattat
gcagactctgtgaagggcagattcaccatctccagagacaattccaagaacacgctgtat
cttcaaatgggcagcctgagagctgaggacatggctgtgtattactgtgcgagaga
>IGHV3-64D*06
gaggtgcagctggtggagtctgggggaggcttggtccagcctggggggtccctgagactc
tcctgttcagcctctggattcaccttcagtagctatgctatgcactgggtccgccaggct
ccagggaagggactggaatatgtttcagctattagtagtaatgggggtagcacatactac
gcagactccgtgaagggcagattcaccatctccagagacaattccaagaacacgctgtat
cttcaaatgagcagtctgagagctgaggacacggctgtgtattactgtgtgaaaga
>IGHV3-66*01
gaggtgcagctggtggagtctgggggaggcttggtccagcctggggggtccctgagactc
tcctgtgcagcctctggattcaccgtcagtagcaactacatgagctgggtccgccaggct
ccagggaaggggctggagtgggtctcagttatttatagcggtggtagcacatactacgca
gactccgtgaagggcagattcaccatctccagagacaattccaagaacacgctgtatctt
caaatgaacagcctgagagccgaggacacggctgtgtattactgtgcgagaga
>IGHV3-66*02
gaggtgcagctggtggagtctgggggaggcttggtccagcctggggggtccctgagactc
tcctgtgcagcctctggattcaccgtcagtagcaactacatgagctgggtccgccaggct
ccagggaaggggctggagtgggtctcagttatttatagcggtggtagcacatactacgca
gactccgtgaagggccgattcaccatctccagagacaattccaagaacacgctgtatctt
caaatgaacagcctgagagctgaggacacggctgtgtattactgtgcgaga
>IGHV3-66*03
gaggtgcagctggtggagtctggaggaggcttgatccagcctggggggtccctgagactc
tcctgtgcagcctctgggttcaccgtcagtagcaactacatgagctgggtccgccaggct
ccagggaaggggctggagtgggtctcagttatttatagctgtggtagcacatactacgca
gactccgtgaagggccgattcaccatctccagagacaattccaagaacacgctgtatctt
caaatgaacagcctgagagctgaggacacggctgtgtattactgtgcgagaga
>IGHV3-66*04
gaggtgcagctggtggagtctgggggaggcttggtccagcctggggggtccctgagactc
tcctgtgcagcctctggattcaccgtcagtagcaactacatgagctgggtccgccaggct
ccagggaaggggctggagtgggtctcagttatttatagcggtggtagcacatactacgca
gactccgtgaagggcagattcaccatctccagagacaattccaagaacacgctgtatctt
caaatgaacagcctgagagccgaggacacggctgtgtattactgtgcgagaca
>IGHV3-7*01

gaggtgcagctggtggagtctgggggaggcttggtccagcctggggggtccctgagactc
tcctgtgcagcctctggattcacctttagtagctattggatgagctgggtccgccaggct
ccagggaaggggctggagtgggtggccaacataaagcaagatggaagtgagaaatactat
gtggactctgtgaagggccgattcaccatctccagagacaacgccaagaactcactgtat
ctgcaaatgaacagcctgagagccgaggacacggctgtgtattactgtgcgagaga
>IGHV3-7*02
gaggtgcagctggtggagtctgggggaggcttggtccagcctggggggtccctgagactc
tcctgtgcagcctctggattcacctttagtagctattggatgagctgggtccgccaggct
ccagggaaagggctggagtgggtggccaacataaagcaagatggaagtgagaaatactat
gtggactctgtgaagggccgattcaccatctccagagacaacgccaagaactcactgtat
ctgcaaatgaacagcctgagagccgaggacacggctgtgtattactgtgcgaga
>IGHV3-7*03
gaggtgcagctggtggagtctgggggaggcttggtccagcctggggggtccctgagactc
tcctgtgcagcctctggattcacctttagtagctattggatgagctgggtccgccaggct
ccagggaaggggctggagtgggtggccaacataaagcaagatggaagtgagaaatactat
gtggactctgtgaagggccgattcaccatctccagagacaacgccaagaactcactgtat
ctgcaaatgaacagcctgagagccgaggacacggccgtgtattactgtgcgagaga
>IGHV3-7*04
gaggtgcagctggtggagtctgggggaggcttggtccagcctggggggtccctgagactc
tcctgtgcagcctctggattcacctttagtagctattggatgagctgggtccgccaggct
ccagggaaagggctggagtgggtggccaacataaagcaagatggaagtgagaaatactat
gtggactctgtgaagggccgattcaccatctccagagacaacgccaagaactcactgtat
ctgcaaatgaacagcctgagagccgaggacacggctgtgtattactgtgcgaggga
>IGHV3-72*01
gaggtgcagctggtggagtctgggggaggcttggtccagcctggagggtccctgagactc
tcctgtgcagcctctggattcaccttcagtgaccactacatggactgggtccgccaggct
ccagggaaggggctggagtgggttggccgtactagaaacaaagctaacagttacaccaca
gaatacgccgcgtctgtgaaaggcagattcaccatctcaagagatgattcaaagaactca
ctgtatctgcaaatgaacagcctgaaaaccgaggacacggccgtgtattactgtgctaga
ga
>IGHV3-72*02
accttcagtgaccactacatggactgggtccgccaggctccagggaaggggctggagtgg
gttggccgtactagaaacaaagctaacagctacaccacagaatacgccgcgtctgtgaaa
ggcagattcaccatctcaagagatgattcaaagaactcactgtat

>IGHV3-73*01
gaggtgcagctggtggagtctgggggaggcttggtccagcctggggggtccctgaaactc
tcctgtgcagcctctgggttcaccttcagtggctctgctatgcactgggtccgccaggct
tccgggaaagggctggagtgggttggccgtattagaagcaaagctaacagttacgcgaca
gcatatgctgcgtcggtgaaaggcaggttcaccatctccagagatgattcaaagaacacg
gcgtatctgcaaatgaacagcctgaaaaccgaggacacggccgtgtattactgtactaga
ca
>IGHV3-73*02
gaggtgcagctggtggagtccgggggaggcttggtccagcctggggggtccctgaaactc
tcctgtgcagcctctgggttcaccttcagtggctctgctatgcactgggtccgccaggct
tccgggaaagggctggagtgggttggccgtattagaagcaaagctaacagttacgcgaca
gcatatgctgcgtcggtgaaaggcaggttcaccatctccagagatgattcaaagaacacg
gcgtatctgcaaatgaacagcctgaaaaccgaggacacggccgtgtattactgtactaga
ca
>IGHV3-74*01
gaggtgcagctggtggagtccgggggaggcttagttcagcctggggggtccctgagactc
tcctgtgcagcctctggattcaccttcagtagctactggatgcactgggtccgccaagct
ccagggaaggggctggtgtgggtctcacgtattaatagtgatgggagtagcacaagctac
gcggactccgtgaagggccgattcaccatctccagagacaacgccaagaacacgctgtat
ctgcaaatgaacagtctgagagccgaggacacggctgtgtattactgtgcaagaga
>IGHV3-74*02
gaggtgcagctggtggagtctgggggaggcttagttcagcctggggggtccctgagactc
tcctgtgcagcctctggattcaccttcagtagctactggatgcactgggtccgccaagct
ccagggaaggggctggtgtgggtctcacgtattaatagtgatgggagtagcacaagctac
gcggactccgtgaagggccgattcaccatctccagagacaacgccaagaacacgctgtat
ctgcaaatgaacagtctgagagccgaggacacggctgtgtattactgtgcaaga
>IGHV3-74*03
gaggtgcagctggtggagtccgggggaggcttagttcagcctggggggtccctgagactc
tcctgtgcagcctctggattcaccttcagtagctactggatgcactgggtccgccaagct
ccagggaaggggctggtgtgggtctcacgtattaatagtgatgggagtagcacaacgtac
gcggactccgtgaagggccgattcaccatctccagagacaacgccaagaacacgctgtat
ctgcaaatgaacagtctgagagccgaggacacggctgtgtattactgtgcaagaga
>IGHV3-9*01
gaagtgcagctggtggagtctgggggaggcttggtacagcctggcaggtccctgagactc

tcctgtgcagcctctggattcacctttgatgattatgccatgcactgggtccggcaagct
ccagggaagggcctggagtgggtctcaggtattagttggaatagtggtagcataggctat
gcggactctgtgaagggccgattcaccatctccagagacaacgccaagaactccctgtat
ctgcaaatgaacagtctgagagctgaggacacggccttgtattactgtgcaaaagata
>IGHV3-9*02
gaagtgcagctggtggagtctgggggaggcttggtacagcctggcaggtccctgagactc
tcctgtgcagcctctggattcacctctgatgattatgccatgcactgggtccggcaagct
ccagggaagggcctggagtgggtctcaggtattagttggaatagtggtagcataggctat
gcggactctgtgaagggccgattcaccatctccagagacaacgccaagaactccctgtat
ctgcaaatgaacagtctgagagctgaggacacggccttgtattactgtgcaaaagata
>IGHV3-9*03
gaagtgcagctggtggagtctgggggaggcttggtacagcctggcaggtccctgagactc
tcctgtgcagcctctggattcacctttgatgattatgccatgcactgggtccggcaagct
ccagggaagggcctggagtgggtctcaggtattagttggaatagtggtagcataggctat
gcggactctgtgaagggccgattcaccatctccagagacaacgccaagaactccctgtat
ctgcaaatgaacagtctgagagctgaggacatggccttgtattactgtgcaaaagata
>IGHV3-NL1*01
caggtgcagctggtggagtctgggggaggcgtggtccagcctgggggtccctgagactc
tcctgtgcagcgtctggattcaccttcagtagctatggcatgcactgggtccgccaggct
ccaggcaaggggctggagtgggtctcagttatttatagcggtggtagtagcacatactat
gcagactccgtgaagggccgattcaccatctccagagacaattccaagaacacgctgtat
ctgcaaatgaacagcctgagagctgaggacacggctgtgtattactgtgcgaaaga
>IGHV4-28*01
caggtgcagctgcaggagtcgggcccaggactggtgaagccttcggacaccctgtccctc
acctgcgctgtctctggttactccatcagcagtagtaactggtggggctggatccggcag
cccccagggaagggactggagtggattgggtacatctattatagtgggagcacctactac
aacccgtccctcaagagtcgagtcaccatgtcagtagacacgtccaagaaccagttctcc
ctgaagctgagctctgtgaccgccgtggacacggccgtgtattactgtgcgagaaa
>IGHV4-28*02
caggtgcagctgcaggagtcgggcccaggactggtgaagccttcacagaccctgtccctc
acctgcgctgtctctggttactccatcagcagtagtaactggtggggctggatccggcag
cccccagggaagggactggagtggattgggtacatctattatagtgggagcatctactac
aacccgtccctcaagagtcgagtcaccatgtcagtagacacgtccaagaaccagttctcc
ctgaagctgagctctgtgaccgccgtggacacggccgtgtattactgtgcgagaaa

>IGHV4-28*03
caggtgcagctgcaggagtcgggcccaggactggtgaagccttcggacaccctgtccctc
acctgcgctgtctctggttactccatcagcagtagtaactggtggggctggatccggcag
cccccagggaagggactggagtggattgggtacatctattatagtgggagcacctactac
aacccgtccctcaagagtcgagtcaccatgtcagtagacacgtccaagaaccagttctcc
ctgaagctgagctctgtgaccgccgtggacacggccgtgtattactgtgcgagaga
>IGHV4-28*04
caggtgcagctgcaggagtcgggcccaggactggtgaagccttcggacaccctgtccctc
acctgcgctgtctctggttactccatcagcagtagtaactggtggggctggatccggcag
cccccagggaagggactggagtggattgggtacatctattatagtgggagcacctactac
aacccgtccctcaagagtcgagtcaccatgtcagtagacacgtccaagaaccagttctcc
ctgaagctgagctctgtgaccgccgtggacaccggcgtgtattactgtgcgaga
>IGHV4-28*05
caggtgcagctgcaggagtcgggcccaggactggtgaagccttcggacaccctgtccctc
acctgcgctgtctctggttactccatcagcagtagtaactggtggggctggatccggcag
cccccagggaagggactggagtggattgggtacatctattatagtgggagcatctactac
aacccgtccctcaagagtcgagtcaccatgtcagtagacacgtccaagaaccagttctcc
ctgaagctgagctctgtgaccgccgtggacacggccgtgtattactgtgcgagaaa
>IGHV4-28*06
caggtgcagctacaggagtcgggcccaggactggtgaagccttcggacaccctgtccctc
acctgcgctgtctctggttactccatcagcagtagtaactggtggggctggatccggcag
cccccagggaagggactggagtggattgggtacatctattatagtgggagcaccaactac
aacccgtccctcaagagtcgagtcaccatgtcagtagacacgtccaagaaccagttctcc
ctgaagctgagctctgtgaccgccttggacacggccgtgtattactgtgcgagaaa
>IGHV4-28*07
caggtacagctgcaggagtcgggcccaggactggtgaagccttcggacaccctgtccctc
acctgcgctgtctctggttactccatcagcagtagtaactggtggggctggatccggcag
cccccagggaagggactggagtggattgggtacatctattatagtgggagcacctactac
aacccgtccctcaagagtcgagtcaccatgtcagtagacacgtccaagaaccagttctcc
ctgaagctgagctctgtgaccgccgtggacacggccgtgtattactgtgcgagaaa
>IGHV4-30-2*01
cagctgcagctgcaggagtccggctcaggactggtgaagccttcacagaccctgtccctc
acctgcgctgtctctggtggctccatcagcagtggtggttactcctggagctggatccgg
cagccaccagggaagggcctggagtggattgggtacatctatcatagtgggagcacctac

tacaacccgtccctcaagagtcgagtcaccatatcagtagacaggtccaagaaccagttc
tccctgaagctgagctctgtgaccgccgcggacacggccgtgtattactgtgccagaga
>IGHV4-30-2*02
cagctgcagctgcaggagtccggctcaggactggtgaagccttcacagaccctgtccctc
acctgcgctgtctctggtggctccatcagcagtggtggttactcctggagctggatccgg
cagccaccagggaagggcctggagtggattgggtacatctatcatagtgggagcacctac
tacaacccgtccctcaagagtcgagtcaccatatcagtagacaggtccaagaaccagttc
tccctgaagctgagctctgtgaccgctgcggacacggccgtgtattactgtgcg
>IGHV4-30-2*03
cagctgcagctgcaggagtccggctcaggactggtgaagccttcacagaccctgtccctc
acctgcgctgtctctggtggctccatcagcagtggtggttactcctggagctggatccgg
cagccaccagggaagggcctggagtggattgggagtatctattatagtgggagcacctac
tacaacccgtccctcaagagtcgagtcaccatatccgtagacacgtccaagaaccagttc
tccctgaagctgagctctgtgaccgctgcagacacggctgtgtattactgtgcgagaca
>IGHV4-30-2*04
tctggtggctccatcagcagtggtggttactcctggagctggatccggcagccaccaggg
aagggcctggagtggattgggtacatctatcatagtgggagcacctactacaacccgtcc
ctcaagagtcgagtcaccatatcagtagacacgtccaagaaccagttctccctgaagctg
agctctgtgaccgccgcagacacggccgtgtattactgtgcgagaga
>IGHV4-30-2*05
cagctgcagctgcaggagtccggctcaggactggtgaagccttcacagaccctgtccctc
acctgcgctgtctctggtggctccatcagcagtggtggttactcctggagctggatccgg
cagccaccagggaagggcctggagtggattgggtacatctatcatagtgggagcacctac
tacaacccgtccctcaagagtcgagttaccatatcagtagacacgtccaagaaccagttc
tccctgaagctgagctctgtgactgccgcagacacggccgtgtattactgtgccagaga
>IGHV4-30-2*06
cagctgcagctgcaggagtccggctcaggactggtgaagccttcacagaccctgtccctc
acctgcgctgtctctggtggctccatcagcagtggtggttactcctggagctggatccgg
cagtcaccagggaagggcctggagtggattgggtacatctatcatagtgggagcacctac
tacaacccgtccctcaagagtcgagtcaccatatcagtagacaggtccaagaaccagttc
tccctgaagctgagctctgtgaccgccgcggacacggccgtgtattactgtgccagaga
>IGHV4-30-4*01
caggtgcagctgcaggagtcgggcccaggactggtgaagccttcacagaccctgtccctc
acctgcactgtctctggtggctccatcagcagtggtgattactactggagttggatccgc

cagcccccagggaagggcctggagtggattgggtacatctattacagtgggagcacctac
tacaacccgtccctcaagagtcgagttaccatatcagtagacacgtccaagaaccagttc
tccctgaagctgagctctgtgactgccgcagacacggccgtgtattactgtgccagaga
>IGHV4-30-4*02
caggtgcagctgcaggagtcgggcccaggactggtgaagccttcggacaccctgtccctc
acctgcactgtctctggtggctccatcagcagtggtgattactactggagttggatccgc
cagcccccagggaagggcctggagtggattgggtacatctattacagtgggagcacctac
tacaacccgtccctcaagagtcgagttaccatatcagtagacacgtccaagaaccagttc
tccctgaagctgagctctgtgactgcagcagacacggccgtgtattactgtgccagaga
>IGHV4-30-4*03
caggtgcagctgcaggagtcgggcccaggactggtgaagccttcacagaccctgtccctc
acctgcactgtctctggtggctccatcagcagtggtgattactactggagttggatccgc
cagcccccagggaagggcctggagtggattgggtacatctattacagtgggagcacctac
tacaacccgtccctcaagagtcgagttaccatatcagtagacacgtccaagaaccagttc
tccctgaagctgagctctgtgactgccgcggacacggccgtgtattactg
>IGHV4-30-4*04
caggtgcagctgcaggactcgggcccaggactggtgaagccttcacagaccctgtccctc
acctgcactgtctctggtggctccatcagcagtggtgattactactggagttggatccgc
cagcccccagggaagggcctggagtggattgggtacttctattacagtgggagcacctac
tacaacccgtccctcaagagtcgagttaccatatcagtagacacgtccaagaaccagttc
tccctgaagctgagctctgtgactgccgcagacacggccgtgtattactg
>IGHV4-30-4*05
ctctggtggctccatcagcagtggtgattactactggagttggatccgccagcncccagg
gaagggcctggagtggattgggtacatctattacagtgggagcacctactacaacccgtc
cctcaagagtcgagtcaccatatcagtagacacgtccaagaaccagttctccctgaagct
gagctctgtgactgccgcagacacggccgtgtattactgtgccagaga
>IGHV4-30-4*06
tctggtggctccatcagcagtggtgattactactggagttggatccgccagcacccaggg
aagggcctggagtggattgggtacatctattacagtgggagcacctactacaacccgtcc
ctcaagagtcgagttaccatatcagtagacacgtccaagaaccagttctccctgaagctg
agctctgtgactgccgcagacacggccgtgtattactgtgccagaga
>IGHV4-30-4*07
caggtgcagctgcaggagtcgggcccaggactggtgaagccttcacagaccctgtccctc
acctgcgctgtctctggtggctccatcagcagtggtggttactcctggagctggatccgg

cagccaccagggaagggactggagtggattgggtatatctattacagtgggagcacctac
tacaacccgtccctcaagagtcgagttaccatatcagtagacacgtccaagaaccagttc
tccctgaagctgagctctgtgaccgccgcggacacggccgtgtattactgtgccagaga
>IGHV4-30-4*08
caggtgcagctgcaggagtcgggcccaggactggtgaagccttcacagaccctgtccctc
acctgcactgtctctggtggctccatcagcagtggtgattactactggagctggatccgc
cagcccccagggaagggcctggagtggattgggtacatctattacagtgggagcacctac
tacaacccgtccctcaagagtcgagttaccatatcagtagacacgtccaagaaccagttc
tccctgaagctgagctctgtgactgccgcagacacggccgtgtattactgtgccagag
>IGHV4-31*01
caggtgcagctgcaggagtcgggcccaggactggtgaagccttcacagaccctgtccctc
acctgcactgtctctggtggctccatcagcagtggtggttactactggagctggatccgc
cagcacccagggaagggcctggagtggattgggtacatctattacagtgggagcacctac
tacaacccgtccctcaagagtctagttaccatatcagtagacacgtctaagaaccagttc
tccctgaagctgagctctgtgactgccgcggacacggccgtgtattactgtgcgagaga
>IGHV4-31*02
caggtgcagctgcaggagtcgggcccaggactggtgaagccttcacagaccctgtccctc
acctgtactgtctctggtggctccatcagcagtggtggttactactggagctggatccgc
cagcacccagggaagggcctggagtggattgggtacatctattacagtgggagcacctac
tacaacccgtccctcaagagtcgagttaccatatcagtagacacgtctaagaaccagttc
tccctgaagctgagctctgtgactgccgcggacacggccgtgtattactgtgcgagaga
>IGHV4-31*03
caggtgcagctgcaggagtcgggcccaggactggtgaagccttcacagaccctgtccctc
acctgcactgtctctggtggctccatcagcagtggtggttactactggagctggatccgc
cagcacccagggaagggcctggagtggattgggtacatctattacagtgggagcacctac
tacaacccgtccctcaagagtcgagttaccatatcagtagacacgtctaagaaccagttc
tccctgaagctgagctctgtgactgccgcggacacggccgtgtattactgtgcgagaga
>IGHV4-31*04
caggtgcggctgcaggagtcgggcccaggactggtgaagccttcacagaccctgtccctc
acctgcactgtctctggtggctccatcagcagtggtggttactactggagctggatccgc
cagcacccagggaagggcctggagtggattgggtacatctattacagtgggagcacctac
tacaacccgtccctcaagagtcgagttaccatatcagtagacacgtctaagaaccagttc
tccctgaagctgagctctgtgactgccgcggacacggccgtgtattactgtgcg
>IGHV4-31*05

caggtgcagctgcaggagtcgggcccaggactggtgaagccttcacagaccctgtccctc
acctgcactgtctctggtggctccatcagcagtggtggttactactggagctggatccgc
cagcacccagggaagggcctggagtggattgggtacatctattacagtgggagcacctac
tacaacccgtccctcaagagtcgagttaccatatcagtagacacgtctaagaaccagttc
tccctgaagctgagctctgtgaccgcggacgcggccgtgtattactgtgcg
>IGHV4-31*06
caggtgcagctgcaggagtcgggcccaggactggtgaagccttcacagaccctgtccctc
acctgcactgtctctggtggctccatcagcagtggtagttactactggagctggatccgc
cagcacccagggaagggcctggagtggattgggtacatctattacagtgggagcacctac
tacaacccgtccctcaagagtcgagttaccatatcagtagacacgtctaagaaccagttc
tccctgaagctgagctctgtgactgccgcggacacggccgtgtattactg
>IGHV4-31*07
caggtgcagctgcaggagtcgggcccaggactggtgaagccttcacagaccctgtccctc
acctgcactgtctctggtggatccatcagcagtggtggttactactggagctggatccgc
cagcacccagggaagggcctggagtggattgggtacatctattacagtgggagcacctac
tacaacccgtccctcaagagtcgagttaccatatcagtagacacgtctaagaaccagttc
tccctgaagctgagctctgtgactgccgcggacacggccgtgtattactg
>IGHV4-31*08
caggtgcagctgcaggagtcgggcccaggactggtgaagccttcacagaccctgtccctc
acctgcactgtctctggtggctccatcagcagtggtggttactactggagctggatccgc
cagcacccagggaagggcctggagtggattgggtacatctattacagtgggagcacctac
tacaacccgtccctcaagagtcgagttaccatatccgtagacacgtccaagaaccagttc
tccctgaagctgagctctgtgactgccgcggacacggccgtgtattactg
>IGHV4-31*09
caggtgcagctgcaggagtcgggcccaggactggtgaagccttcacagaccctgtccctc
acctgcactgtctctggtggctccatcagcagtggtggttactactggagctggatccgc
cagcacccagggaagggcctggagtggattgggtacatctattacagtgggagcacctac
tacaacccgtccctcaagagtcgagttaccatatcagtagacaagtccaagaaccagttc
tccctgaagctgagctctgtgaccgccgcggacacggccgtgtattactg
>IGHV4-31*10
caggtgcagctgcaggagtcgggcccaggactgttgaagccttcacagaccctgtccctc
acctgcactgtctctggtggctccatcagcagtggtggttactactggagctggatccgc
cagcacccagggaagggcctggagtggattgggtgcatctattacagtgggagcacctac
tacaacccgtccctcaagagtcgagttaccatatcagtagacccgtccaagaaccagttc

tccctgaagccgagctctgtgactgccgcggacacggccgtggattactgtgcgagaga
>IGHV4-31*11
caggtgcagctgcaggagtcgggcccaggactggtgaagccttcacagaccctgtccctc
acctgcgctgtctctggtggctccatcagcagtggtggttactactggagctggatccgc
cagcacccagggaagggcctggagtggattgggtacatctattacagtgggagcacctac
tacaacccgtccctcaagagtcgagttaccatatcagtagacacgtctaagaaccagttc
tccctgaagctgagctctgtgactgccgcggacacggccgtgtattactgtgcgagaga
>IGHV4-34*01
caggtgcagctacagcagtggggcgcaggactgttgaagccttcggagaccctgtccctc
acctgcgctgtctatggtgggtccttcagtggttactactggagctggatccgccagccc
ccagggaaggggctggagtggattggggaaatcaatcatagtggaagcaccaactacaac
ccgtccctcaagagtcgagtcaccatatcagtagacacgtccaagaaccagttctccctg
aagctgagctctgtgaccgccgcggacacggctgtgtattactgtgcgagagg
>IGHV4-34*02
caggtgcagctacaacagtggggcgcaggactgttgaagccttcggagaccctgtccctc
acctgcgctgtctatggtgggtccttcagtggttactactggagctggatccgccagccc
ccagggaaggggctggagtggattggggaaatcaatcatagtggaagcaccaactacaac
ccgtccctcaagagtcgagtcaccatatcagtagacacgtccaagaaccagttctccctg
aagctgagctctgtgaccgccgcggacacggctgtgtattactgtgcgagagg
>IGHV4-34*03
caggtgcagctacagcagtggggcgcaggactgttgaagccttcggagaccctgtccctc
acctgcgctgtctatggtgggtccttcagtggttactactggagctggatccgccagccc
ccagggaaggggctggagtggattggggaaatcaatcatagtggaagcaccaactacaac
ccgtccctcaagagtcgagtcaccatatcagtagacacgtccaagaaccagttctccctg
aagctgagctctgtgaccgccgcggacacggccgtgtattactg
>IGHV4-34*04
caggtgcagctacagcagtggggcgcaggactgttgaagccttcggagaccctgtccctc
acctgcgctgtctatggtgggtccttcagtggttactactggagctggatccgccagccc
ccagggaaggggctggagtggattggggaaatcaatcatagtggaagcaccaacaacaac
ccgtccctcaagagtcgagccaccatatcagtagacacgtccaagaaccagttctccctg
aagctgagctctgtgaccgccgcggacacggctgtgtattactgtgcgagagg
>IGHV4-34*05
caggtgcagctacagcagtggggcgcaggactgttgaagccttcggagaccctgtccctc
acctgcgctgtctatggtgggtccttcagtggttactactggtgctggatccgccagccc

ctagggaaggggctggagtggattggggaaatcaatcatagtggaagcaccaacaacaac
ccgtccctcaagagtcgagccaccatatcagtagacacgtccaagaaccagttctccctg
aagctgagctctgtgaccgccgcggacacggctgtgtattactgtgcgagagg
>IGHV4-34*06
caggtgcagctacagcagtggggcgcaggactgttgaagccttcggagaccctgtccctc
acctgcgctgtctatggtgggtccttcagtggttactactggagctggatccgccagccc
ccagggaaggggctggagtggattggggaaatcaatcatagtggaagcaccaactacaac
ccgtccctcaagagtcgagtcaccatatcagtagacacgtccaagaaccagttctccctg
aagctgggctctgtgaccgccgcggacacggccgtgtattactg
>IGHV4-34*07
caggtgcagctacagcagtggggcgcaggactgttgaagccttcggagaccctgtccctc
acctgcgctgtctatggtgggtccttcagtggttactactggagctggatccgccagccc
ccagggaaggggctggagtggattggggaaatcaaccatagtggaagcaccaactacaac
ccgtccctcaagagtcgagtcaccatatcagtagacacgtccaagaaccagttctccctg
aagctgagctctgtgaccgccgcggacacggccgtgtattactg
>IGHV4-34*08
caggtgcagctacagcagtggggcgcaggactgttgaagccttcggagaccctgtccctc
acctgcgctgtctatggtgggaccttcagtggttactactggagctggatccgccagccc
ccagggaaggggctggagtggattggggaaatcaatcatagtggaagcaccaactacaac
ccgtccctcaagagtcgagtcaccatatcagtagacacgtccaagaaccagttctccctg
aagctgagctctgtgaccgccgcggacacggctgtgtattactgtgcg
>IGHV4-34*09
caggtgcagctgcaggagtcgggcccaggactggtgaagccttcacagaccctgtccctc
acctgcgctgtctatggtgggtccttcagtggttactactggagctggatccgccagccc
ccagggaagggactggagtggattggggaaatcaatcatagtggaagcaccaactacaac
ccgtccctcaagagtcgagttaccatatcagtagacacgtctaagaaccagttctccctg
aagctgagctctgtgactgccgcggacacggccgtgtattactgtgcgagaga
>IGHV4-34*10
caggtgcagctgcaggagtcgggcccaggactggtgaagccttcggagaccctgtccctc
acctgcgctgtctatggtgggtccttcagtggttactactggagctggatccgccagccc
ccagggaagggactggagtggattggggaaatcaatcatagtggaagcaccaactacaac
ccgtccctcaagagtcgaatcaccatgtcagtagacacgtccaagaaccagttctacctg
aagctgagctctgtgaccgccgcggacacggccgtgtattactgtgcgagata
>IGHV4-34*11

caggtgcagctacagcagtggggcgcaggactgttgaagccttcggagaccctgtccctc
acctgcgctgtctatggtgggtccgtcagtggttactactggagctggatccggcagccc
ccagggaaggggctggagtggattgggtatatctattatagtgggagcaccaacaacaac
ccctccctcaagagtcgagccaccatatcagtagacacgtccaagaaccagttctccctg
aacctgagctctgtgaccgccgcggacacggccgtgtattgctgtgcgagaga
>IGHV4-34*12
caggtgcagctacagcagtggggcgcaggactgttgaagccttcggagaccctgtccctc
acctgcgctgtctatggtgggtccttcagtggttactactggagctggatccgccagccc
ccagggaaggggctggagtggattggggaaatcattcatagtggaagcaccaactacaac
ccgtccctcaagagtcgagtcaccatatcagtagacacgtccaagaaccagttctccctg
aagctgagctctgtgaccgccgcggacacggctgtgtattactgtgcgaga
>IGHV4-34*13
tatggtgggtccttcagtggttactactggagctggatccgccagcccccagggaagggg
ctggagtggattggggaaatcaatcatagtggaagcaccaactacaacccctccctcaag
agtcgagtcaccatatcagtagacacgtccaagaaccagttctccctgaagctgagctct
gtgaccgccgcggacacggctgtgtattactgtgcgagagg
>IGHV4-38-2*01
caggtgcagctgcaggagtcgggcccaggactggtgaagccttcggagaccctgtccctc
acctgcgctgtctctggttactccatcagcagtggttactactggggctggatccggcag
cccccagggaaggggctggagtggattgggagtatctatcatagtgggagcacctactac
aacccgtccctcaagagtcgagtcaccatatcagtagacacgtccaagaaccagttctcc
ctgaagctgagctctgtgaccgccgcagacacggccgtgtattactgtgcgaga
>IGHV4-38-2*02
caggtgcagctgcaggagtcgggcccaggactggtgaagccttcggagaccctgtccctc
acctgcactgtctctggttactccatcagcagtggttactactggggctggatccggcag
cccccagggaaggggctggagtggattgggagtatctatcatagtgggagcacctactac
aacccgtccctcaagagtcgagtcaccatatcagtagacacgtccaagaaccagttctcc
ctgaagctgagctctgtgaccgccgcagacacggccgtgtattactgtgcgagaga
>IGHV4-39*01
cagctgcagctgcaggagtcgggcccaggactggtgaagccttcggagaccctgtccctc
acctgcactgtctctggtggctccatcagcagtagtagttactactggggctggatccgc
cagcccccagggaaggggctggagtggattgggagtatctattatagtgggagcacctac
tacaacccgtccctcaagagtcgagtcaccatatccgtagacacgtccaagaaccagttc
tccctgaagctgagctctgtgaccgccgcagacacggctgtgtattactgtgcgagaca

```
>IGHV4-39*02
cagctgcagctgcaggagtcgggcccaggactggtgaagccttcggagaccctgtccctc
acctgcactgtctctggtggctccatcagcagtagtagttactactggggctggatccgc
cagcccccagggaaggggctggagtggattgggagtatctattatagtgggagcacctac
tacaacccgtccctcaagagtcgagtcaccatatccgtagacacgtccaagaaccacttc
tccctgaagctgagctctgtgaccgccgcagacacggctgtgtattactgtgcgagaga
>IGHV4-39*03
cagctgcagctgcaggagtcgggcccaggactggtgaagccttcggagaccctgtccctc
acctgcactgtctctggtggctccatcagcagtagtagttactactggggctggatccgc
cagcccccagggaaggggctggagtggattgggagtatctattatagtgggagcacctac
tacaacccgtccctcaagagtcgagtcaccatatccgtagacacgtccaagaaccagttc
tccctgaagctgagctctgtgaccgccgcagacacggccgtgtattactg
>IGHV4-39*04
gctccatcagcagtagtagttactactggggctggatccgccagcccccagggaaggggc
tggagtggattgggagtatctattatagtgggagcacctactacaacccgtccctcaaga
gtcgagtcaccatatccgtagacacgtccaagaaccagttctccctgaagctgagctctg
tgaccgccgcggacac
>IGHV4-39*05
cagctgcagctgcaggagtcgggcccaggactggtgaagccttcggagaccccgtccctc
acctgcactgtctctggtggctccatcagcagtagtagttactactggggctggatccgc
cagcccccagggaaggggctggagtggattgggagtatctattatagtgggagcacctac
tacaacccgtccctcaagagtcgagtcaccatatccgtagacacgtccaagaaccagttc
tccctgaagctgagctctgtgaccgccgcagacacggctgtgtattactgtgcg
>IGHV4-39*06
cggctgcagctgcaggagtcgggcccaggactggtgaagccttcggagaccctgtccctc
acctgcactgtctctggtggctccatcagcagtagtagttactactggggctggatccgc
cagcccccagggaaggggctggagtggattgggagtatctattatagtgggagcacctac
tacaacccgtccctcaagagtcgagtcaccatatcagtagacacgtccaagaaccagttc
cccctgaagctgagctctgtgaccgccgcggacacggccgtgtattactgtgcgagaga
>IGHV4-39*07
cagctgcagctgcaggagtcgggcccaggactggtgaagccttcggagaccctgtccctc
acctgcactgtctctggtggctccatcagcagtagtagttactactggggctggatccgc
cagcccccagggaaggggctggagtggattgggagtatctattatagtgggagcacctac
tacaacccgtccctcaagagtcgagtcaccatatcagtagacacgtccaagaaccagttc
```

```
tccctgaagctgagctctgtgaccgccgcggacacggccgtgtattactgtgcgagaga
>IGHV4-4*01
caggtgcagctgcaggagtcgggcccaggactggtgaagcctccggggaccctgtccctc
acctgcgctgtctctggtggctccatcagcagtagtaactggtggagttgggtccgccag
cccccagggaaggggctggagtggattggggaaatctatcatagtgggagcaccaactac
aacccgtccctcaagagtcgagtcaccatatcagtagacaagtccaagaaccagttctcc
ctgaagctgagctctgtgaccgccgcggacacggccgtgtattgctgtgcgagaga
>IGHV4-4*02
caggtgcagctgcaggagtcgggcccaggactggtgaagccttcggggaccctgtccctc
acctgcgctgtctctggtggctccatcagcagtagtaactggtggagttgggtccgccag
cccccagggaaggggctggagtggattggggaaatctatcatagtgggagcaccaactac
aacccgtccctcaagagtcgagtcaccatatcagtagacaagtccaagaaccagttctcc
ctgaagctgagctctgtgaccgccgcggacacggccgtgtattactgtgcgagaga
>IGHV4-4*03
caggtgcagctgcaggagtcgggcccaggactggtgaagcctccggggaccctgtccctc
acctgcgctgtctctggtggctccatcagcagtagtaactggtggagttgggtccgccag
cccccagggaaggggctggagtggattggggaaatctatcatagtgggagcaccaactac
aacccgtccctcaagagtcgagtcaccatatcagtagacaagtccaagaaccagttctcc
ctgaagctgagctctgtgaccgccgcggacacggccgtgtattactgtgcgagag
>IGHV4-4*04
caggtgcagctgcaggagtcgggcccaggactggtgaagcctccggggaccctgtccctc
acctgcgctatctctggtggctccatcagcagtagtaactggtggagttgggtccgccag
cccccagggaaggggctggagtggattggggaaatctatcatagtgggagcaccaactac
aacccgtccctcaagagtcgagtcaccatatcagtagacaagtccaagaaccagttctcc
ctgaagctgagctctgtgaccgccgcggacacggccgtgtattactg
>IGHV4-4*05
caggtgcagctgcaggagttgggcccaggactggtgaagcctccggggaccctgtccctc
acctgcgctgtctctggtggctccatcagcagtagtaactggtggagttgggtccgccag
cccccagggaaggggctggagtggattggggaaatctatcatagtgggagcaccaactac
aacccgtccctcaagagtcgagtcaccatatcagtagacaagtccaagaaccagttctcc
ctgaagctgagctctgtgaccgccgcggacacggccgtgtattactg
>IGHV4-4*06
tctggtggctccatcagcagtagtaactggtggagttgggtccgccagcccccagggann
nggctggagtggattggggaaatctatcatagtgggagcaccaactacaacccgtccctc
```

aagagtcgagtcaccatgtcagtagacacgtccaagaaccagttctccctgaagctgagc
tctgtgaccgccgcggacacggccgtgtattactgtgcgagaga
>IGHV4-4*07
caggtgcagctgcaggagtcgggcccaggactggtgaagccttcggagaccctgtccctc
acctgcactgtctctggtggctccatcagtagttactactggagctggatccggcagccc
gccgggaagggactggagtggattgggcgtatctataccagtgggagcaccaactacaac
ccctccctcaagagtcgagtcaccatgtcagtagacacgtccaagaaccagttctccctg
aagctgagctctgtgaccgccgcggacacggccgtgtattactgtgcgagaga
>IGHV4-4*08
caggtgcagctgcaggagtcgggcccaggactggtgaagccttcggagaccctgtccctc
acctgcactgtctctggtggctccatcagtagttactactggagctggatccggcagccc
ccagggaagggactggagtggattgggtatatctataccagtgggagcaccaactacaac
ccctccctcaagagtcgagtcaccatatccgtagacacgtccaagaaccagttctccctg
aagctgagctctgtgaccgccgcagacacggccgtgtattactgtgcgagaga
>IGHV4-59*01
caggtgcagctgcaggagtcgggcccaggactggtgaagccttcggagaccctgtccctc
acctgcactgtctctggtggctccatcagtagttactactggagctggatccggcagccc
ccagggaagggactggagtggattgggtatatctattacagtgggagcaccaactacaac
ccctccctcaagagtcgagtcaccatatcagtagacacgtccaagaaccagttctccctg
aagctgagctctgtgaccgctgcggacacggccgtgtattactgtgcgagaga
>IGHV4-59*02
caggtgcagctgcaggagtcgggcccaggactggtgaagccttcggagaccctgtccctc
acctgcactgtctctggtggctccgtcagtagttactactggagctggatccggcagccc
ccagggaagggactggagtggattgggtatatctattacagtgggagcaccaactacaac
ccctccctcaagagtcgagtcaccatatcagtagacacgtccaagaaccagttctccctg
aagctgagctctgtgaccgctgcggacacggccgtgtattactgtgcgagaga
>IGHV4-59*03
caggtgcagctgcaggagtcgggcccaggactggtgaagccttcggagaccctgtccctc
acctgcactgtctctggtggctccatcagtagttactactggagctggatccggcagccc
ccagggaagggactggagtggattgggtatatctattacagtgggagcaccaactacaac
ccctccctcaagagtcgagtcaccatatcagtagacacgtccaagaaccaattctccctg
aagctgagctctgtgaccgctgcggacacggccgtgtattactgtgcg
>IGHV4-59*04
caggtgcagctgcaggagtcgggcccaggactggtgaagccttcggagaccctgtccctc

acctgcactgtctctggtggctccatcagtagttactactggagctggatccggcagccc
ccagggaagggactggagtggattgggtatatctattatagtgggagcacctactacaac
ccgtccctcaagagtcgagtcaccatgtcagtagacacgtccaagaaccagttctccctg
aagctgagctctgtgaccgccgcagacacggctgtgtattactgtgcg
>IGHV4-59*05
caggtgcagctgcaggagtcgggcccaggactggtgaagccttcggagaccctgtccctc
acctgcactgtctctggtggctccatcagtagttactactggagctggatccggcagccg
ccggggaagggactggagtggattgggcgtatctattatagtgggagcacctactacaac
ccgtccctcaagagtcgagtcaccatatccgtagacacgtccaagaaccagttctccctg
aagctgagctctgtgaccgccgcagacacggctgtgtattactgtgcg
>IGHV4-59*06
caggtgcagctgcaggagtcgggcccaggactggtgaagccttcggagaccctgtccctc
acctgcactgtcactggtggctccatcagtagttactactggagctggatccggcagccc
gctgggaagggcctggagtggattgggtacatctattacagtgggagcacctactacaac
ccgtccctcaagagtcgagttaccatatcagtagacacgtctaagaaccagttctccctg
aagctgagctctgtgactgccgcggacacggccgtgtattactgtgcg
>IGHV4-59*07
caggtgcagctgcaggagtcgggcccaggactggtgaagccttcggacaccctgtccctc
acctgcactgtctctggtggctccatcagtagttactactggagctggatccggcagccc
ccagggaagggactggagtggattgggtatatctattacagtgggagcaccaactacaac
ccctccctcaagagtcgagtcaccatatcagtagacacgtccaagaaccagttctccctg
aagctgagctctgtgaccgctgcggacacggccgtgtattactgtgcgaga
>IGHV4-59*08
caggtgcagctgcaggagtcgggcccaggactggtgaagccttcggagaccctgtccctc
acctgcactgtctctggtggctccatcagtagttactactggagctggatccggcagccc
ccagggaagggactggagtggattgggtatatctattacagtgggagcaccaactacaac
ccctccctcaagagtcgagtcaccatatcagtagacacgtccaagaaccagttctccctg
aagctgagctctgtgaccgccgcagacacggccgtgtattactgtgcgagaca
>IGHV4-59*09
tctggtggctccatcagtagttactactggagctggatccggcagcccccaggnannnga
ctggagtggattgggtatatctattacagtgggagcaccaactacaacccctccctcaag
agtcgagtcaccatatcagtagacacgtccaagaaccagttctccctgaagctgagctct
gtgaccgctgcggacacggccgtgtattactgtgcgagagg
>IGHV4-59*10

caggtgcagctacagcagtggggcgcaggactgttgaagccttcggagaccctgtccctc
acctgcgctgtctatggtggctccatcagtagttactactggagctggatccggcagccc
gccgggaaggggctggagtggattgggcgtatctataccagtgggagcaccaactacaac
ccctccctcaagagtcgagtcaccatgtcagtagacacgtccaagaaccagttctccctg
aagctgagctctgtgaccgccgcggacacggccgtgtattactgtgcgagata
>IGHV4-59*11
caggtgcagctgcaggagtcgggcccaggactggtgaagccttcggagaccctgtccctc
acctgcactgtctctggtggctccatcagtagtcactactggagctggatccggcagccc
ccagggaagggactggagtggattgggtatatctattacagtgggagcaccaactacaac
ccctccctcaagagtcgagtcaccatatcagtagacacgtccaagaaccagttctccctg
aagctgagctctgtgaccgctgcggacacggccgtgtattactgtgcgagaga
>IGHV4-59*12
caggtgcagctgcaggagtcgggcccaggactggtgaagccttcggagaccctgtccctc
acctgcactgtctctggtggctccatcagtagttactactggagctggatccggcagccc
ccagggaagggactggagtggattgggtatatctattacagtgggagcaccaactacaac
ccctccctcaagagtcgagtcaccatatcagtagacacgtccaagaaccagttctccctg
aagctgagctctgtgaccgccgcggacacggccgtgtattactgtgcgagaga
>IGHV4-59*13
caggtgcagctgcaggagtcgggcccaggactggtgaagccttcggagaccctgtccctc
acctgcactgtctctggtggctccatcagtagttactactggagctggatccggcagccc
ccggggaagggactggagtggattgggtatatctattacagtgggagcaccaactacaac
ccctccctcaagagtcgagtcaccatatcagtagacacgtccaagaaccagttctccctg
aagctgagctctgtgaccgctgcggacacggccgtgtattactgtgcgagaga
>IGHV4-61*01
caggtgcagctgcaggagtcgggcccaggactggtgaagccttcggagaccctgtccctc
acctgcactgtctctggtggctccgtcagcagtggtagttactactggagctggatccgg
cagcccccagggaagggactggagtggattgggtatatctattacagtgggagcaccaac
tacaacccctccctcaagagtcgagtcaccatatcagtagacacgtccaagaaccagttc
tccctgaagctgagctctgtgaccgctgcggacacggccgtgtattactgtgcgagaga
>IGHV4-61*02
caggtgcagctgcaggagtcgggcccaggactggtgaagccttcacagaccctgtccctc
acctgcactgtctctggtggctccatcagcagtggtagttactactggagctggatccgg
cagcccgccgggaagggactggagtggattgggcgtatctataccagtgggagcaccaac
tacaacccctccctcaagagtcgagtcaccatatcagtagacacgtccaagaaccagttc

tccctgaagctgagctctgtgaccgccgcagacacggccgtgtattactgtgcgagaga
>IGHV4-61*03
caggtgcagctgcaggagtcgggcccaggactggtgaagccttcggagaccctgtccctc
acctgcactgtctctggtggctccgtcagcagtggtagttactactggagctggatccgg
cagcccccagggaagggactggagtggattgggtatatctattacagtgggagcaccaac
tacaacccctccctcaagagtcgagtcaccatatcagtagacacgtccaagaaccacttc
tccctgaagctgagctctgtgaccgctgcggacacggccgtgtattactgtgcgagaga
>IGHV4-61*04
caggtgcagctgcaggagtcgggcccaggactggtgaagccttcggagaccctgtccctc
acctgcactgtctctggtggctccgtcagcagtggtagttactactggagctggatccgg
cagcccccagggaagggactggagtggattggatatatctattacagtgggagcaccaac
tacaacccctccctcaagagtcgagtcaccatatcagtagacacgtccaagaaccagttc
tccctgaagctgagctctgtgaccgctgacacggccgtgtattactg
>IGHV4-61*05
cagctgcagctgcaggagtcgggcccaggactggtgaagccttcggagaccctgtccctc
acctgcactgtctctggtggctccatcagcagtagtagttactactggggctggatccgg
cagcccccagggaagggactggagtggattgggtatatctattacagtgggagcaccaac
tacaacccctccctcaagagtcgagtcaccatatcagtagacaagtccaagaaccagttc
tccctgaagctgagctctgtgaccgccgcggacacggccgtgtattactgtgcgaga
>IGHV4-61*06
tctggtggctccgtcagcagtggtagttactactggagctggatccggcagcccccaggg
aagggactggagtggattgggtatatctattacagtgggagcaccaactacaacccctcc
ctcaagagtcgagtcaccatatcagtagacacgtccaagaaccagttctccctgaagctg
agctctgtgaccgccgcggacacggccgtgtattactgtgccagaga
>IGHV4-61*07
tctggtggctccgtcagcagtggtagttactactggagctggatccggcagcccccaggg
aagggactggagtggattgggtatatctattacagtgggagcaccaactacaacccctcc
ctcaagagtcgagtcaccatatcagtagacacgtccaagaaccagttctccctgaagctg
agctctgtgaccgctgcggacacggccgtgtattactgtgcgagaca
>IGHV4-61*08
caggtgcagctgcaggagtcgggcccaggactggtgaagccttcggagaccctgtccctc
acctgcactgtctctggtggctccgtcagcagtggtggttactactggagctggatccgg
cagcccccagggaagggactggagtggattgggtatatctattacagtgggagcaccaac
tacaacccctccctcaagagtcgagtcaccatatcagtagacacgtccaagaaccagttc

tccctgaagctgagctctgtgaccgctgcggacacggccgtgtattactgtgcgagaga
>IGHV4-61*09
caggtgcagctgcaggagtcgggcccaggattggtgaagccttcacagaccctgtccctc
acctgcactgtctctggtggctccatcagcagtggtagttactactggagctggatccgg
cagcccgccgggaagggactggagtggattgggcatatctataccagtgggagcaccaac
tacaacccctccctcaagagtcgagtcaccatatcagtagacacgtccaagaaccagttc
tccctgaagctgagctctgtgaccgccgcagacacggccgtgtattactgtgcgagaga
>IGHV5-10-1*01
gaagtgcagctggtgcagtctggagcagaggtgaaaaagcccggggagtctctgaggatc
tcctgtaagggttctggatacagctttaccagctactggatcagctgggtgcgccagatg
cccgggaaaggcctggagtggatggggaggattgatcctagtgactcttataccaactac
agcccgtccttccaaggccacgtcaccatctcagctgacaagtccatcagcactgcctac
ctgcagtggagcagcctgaaggcctcggacaccgccatgtattactgtgcgaga
>IGHV5-10-1*02
gaagtgcagctggtgcagtctggagcagaggtgaaaaagcccggggagtctctgaggatc
tcctgtaagggttctggatacagctttaccagctactggatcagctgggtgcgccagatg
cccgggaaaggcttggagtggatggggaggattgatcctagtgactcttataccaactac
agcccgtccttccaaggccacgtcaccatctcagctgacaagtccatcagcactgcctac
ctgcagtggagcagcctgaaggctcggacaccgccatgtattactgtgcgagaca
>IGHV5-10-1*03
gaagtgcagctggtgcagtccggagcagaggtgaaaaagcccggggagtctctgaggatc
tcctgtaagggttctggatacagctttaccagctactggatcagctgggtgcgccagatg
cccgggaaaggcctggagtggatggggaggattgatcctagtgactcttataccaactac
agcccgtccttccaaggccacgtcaccatctcagctgacaagtccatcagcactgcctac
ctgcagtggagcagcctgaaggcctcggacaccgccatgtattactgtgcgaga
>IGHV5-10-1*04
gaagtgcagctggtgcagtctggagcagaggtgaaaaagcccggggagtctctgaggatc
tcctgtaagggttctggatacagctttaccagctactggatcagctgggtgcgccagatg
cccgggaaaggcctggagtggatggggaggattgatcctagtgactcttataccaactac
agcccgtccttccaaggccaggtcaccatctcagctgacaagtccatcagcactgcctac
ctgcagtggagcagcctgaaggcctcggacaccgccatgtattactgtgcgaga
>IGHV5-51*01
gaggtgcagctggtgcagtctggagcagaggtgaaaaagcccggggagtctctgaagatc
tcctgtaagggttctggatacagctttaccagctactggatcggctgggtgcgccagatg

cccgggaaaggcctggagtggatggggatcatctatcctggtgactctgataccagatac
agcccgtccttccaaggccaggtcaccatctcagccgacaagtccatcagcaccgcctac
ctgcagtggagcagcctgaaggcctcggacaccgccatgtattactgtgcgagaca
>IGHV5-51*02
gaggtgcagctggtgcagtctggagcagaggtgaaaaagcccggggagtctctgaagatc
tcctgtaagggttctggatacagctttaccagctactggaccggctgggtgcgccagatg
cccgggaaaggcttggagtggatggggatcatctatcctggtgactctgataccagatac
agcccgtccttccaaggccaggtcaccatctcagccgacaagtccatcagcaccgcctac
ctgcagtggagcagcctgaaggcctcggacaccgccatgtattactgtgcgagaca
>IGHV5-51*03
gaggtgcagctggtgcagtctggagcagaggtgaaaaagccggggagtctctgaagatc
tcctgtaagggttctggatacagctttaccagctactggatcggctgggtgcgccagatg
cccgggaaaggcctggagtggatggggatcatctatcctggtgactctgataccagatac
agcccgtccttccaaggccaggtcaccatctcagccgacaagtccatcagcaccgcctac
ctgcagtggagcagcctgaaggcctcggacaccgccatgtattactgtgcgaga
>IGHV5-51*04
gaggtgcagctggtgcagtctggagcagaggtgaaaaagccggggagtctctgaagatc
tcctgtaagggttctggatacagctttaccagctactggatcggctgggtgcgccagatg
cccgggaaaggcctggagtggatggggatcatctatcctggtgactctgataccagatac
agcccgtccttccaaggccaggtcaccatctcagccgacaagcccatcagcaccgcctac
ctgcagtggagcagcctgaaggcctcggacaccgccatgtattactgtgcgaga
>IGHV5-51*05
aaaagcccggggagtctctgaagatctcctgtaagggttctggatacagctttaccagct
actggatcggctgggtgcgccagatgcccaggaaaggcctggagtggatggggatcatct
atcctggtgactctgataccagatacagcccgtccttccaaggccaggtcaccatctcag
ccgacaagtccatcagcaccgcctacctgcagtggagcagcctgaaggcctcggacaccg
ccatg
>IGHV5-51*06
gaggtgcagctggtgcagtctggagcagaggtgaaaaagccggggagtctctgaagatc
tcctgtaagggttctggatacagctttaccagctactggatcggctgggtgcgccagatg
cccgggaaaggcctggagtggatggggatcatctatcctggtgactctgataccagatac
agcccgtccttccaaggccaggttaccatctcagccgacaagtccatcagcaccgcctac
ctgcagtggagcagcctgaaggcctcggacaccgccatgtattactgtgcgaga
>IGHV5-51*07

gaggtgcagctggtgcagtctggagcagaggtgaaaaagcccggggagtctctgaagatc
tcctgtaagggttctggatacagctttaccagctactggatcggctgggtgcaccagatg
cccgggaaaggcctggagtggatggggatcatctatcctggtgactctgataccagatac
agcccgtccttccaaggccaggtcaccatctcagccgacaagtccatcagcaccgcctac
ctgcagtggagcagcctgaaggcctcggacaccgccatgtattactgtgcgagaca
>IGHV6-1*01
caggtacagctgcagcagtcaggtccaggactggtgaagccctcgcagaccctctcactc
acctgtgccatctccggggacagtgtctctagcaacagtgctgcttggaactggatcagg
cagtccccatcgagaggccttgagtggctgggaaggacatactacaggtccaagtggtat
aatgattatgcagtatctgtgaaaagtcgaataaccatcaacccagacacatccaagaac
cagttctccctgcagctgaactctgtgactcccgaggacacggctgtgtattactgtgca
agaga
>IGHV6-1*02
caggtacagctgcagcagtcaggtccgggactggtgaagccctcgcagaccctctcactc
acctgtgccatctccggggacagtgtctctagcaacagtgctgcttggaactggatcagg
cagtccccatcgagaggccttgagtggctgggaaggacatactacaggtccaagtggtat
aatgattatgcagtatctgtgaaaagtcgaataaccatcaacccagacacatccaagaac
cagttctccctgcagctgaactctgtgactcccgaggacacggctgtgtattactgtgca
agaga
>IGHV6-1*03
caggtacagctgcagcagtcaggtccaggactggtgaagccctcgcagaccctctcactc
acctgtgccatctccggggacagtgtctctagcaacagtgctgcttggaactggatcagg
cagtccccatcgagaggccttgagtggctgggaaggacatactacaggtccaagtggtat
aatgattatgcagtatctgtgaaaagttgaataaccatcaacccagacacatccaagaac
cagttctccctgcagctgaactctgtgactcccgaggacacggctgtgtattactgtgca
agaga
>IGHV7-4-1*01
caggtgcagctggtgcaatctgggtctgagttgaagaagcctggggcctcagtgaaggtt
tcctgcaaggcttctggatacaccttcactagctatgctatgaattgggtgcgacaggcc
cctggacaagggcttgagtggatgggatggatcaacaccaacactgggaacccaacgtat
gcccagggcttcacaggacggtttgtcttctccttggacacctctgtcagcacggcatat
ctgcagatctgcagcctaaaggctgaggacactgccgtgtattactgtgcgagaga
>IGHV7-4-1*02
caggtgcagctggtgcaatctgggtctgagttgaagaagcctggggcctcagtgaaggtt

tcctgcaaggcttctggatacaccttcactagctatgctatgaattgggtgcgacaggcc
cctggacaagggcttgagtggatgggatggatcaacaccaacactgggaacccaacgtat
gcccagggcttcacaggacggtttgtcttctccttggacacctctgtcagcacggcatat
ctgcagatcagcagcctaaaggctgaggacactgccgtgtattactgtgcgagaga
>IGHV7-4-1*03
caggtgcagctggtgcaatctgggtctgagttgaagaagcctggggcctcagtgaaggtt
tcctgcaaggcttctggatacaccttcactagctatgctatgaattgggtgcgacaggcc
cctggacaagggcttgagtggatgggatggatcaacaccaacactgggaacccaacgtat
gcccagggcttcacaggacggtttgtcttctccttggacacctctgtcagcacggcatat
ctgcagatcagcacgctaaaggctgaggacactg
>IGHV7-4-1*04
caggtgcagctggtgcaatctgggtctgagttgaagaagcctggggcctcagtgaaggtt
tcctgcaaggcttctggatacaccttcactagctatgctatgaattgggtgcgacaggcc
cctggacaagggcttgagtggatgggatggatcaacaccaacactgggaacccaacgtat
gcccagggcttcacaggacggtttgtcttctccttggacacctctgtcagcatggcatat
ctgcagatcagcagcctaaaggctgaggacactgccgtgtattactgtgcgagaga
>IGHV7-4-1*05
caggtgcagctggtgcaatctgggtctgagttgaagaagcctggggcctcagtgaaggtt
tcctgcaaggcttctggatacaccttcactagctatgctatgaattgggtgcgacaggcc
cctggacaagggcttgagtggatgggatggatcaacaccaacactgggaacccaacgtat
gcccagggcttcacaggacggtttgtcttctccttggacacctctgtcagcatggcatat
ctgcagatcagcagcctaaaggctgaggacactgccgtgtgttactgtgcgagaga

**D.fasta**

```
>IGHD1-1*01
ggtacaactggaacgac
>IGHD1-14*01
ggtataaccggaaccac
>IGHD1-20*01
ggtataactggaacgac
>IGHD1-26*01
ggtatagtgggagctactac
>IGHD1-7*01
ggtataactggaactac
>IGHD1/OR15-1a*01
ggtataactggaacaac
>IGHD1/OR15-1b*01
ggtataactggaacaac
>IGHD2-15*01
aggatattgtagtggtggtagctgctactcc
>IGHD2-2*01
aggatattgtagtagtaccagctgctatgcc
>IGHD2-2*02
aggatattgtagtagtaccagctgctatacc
>IGHD2-2*03
tggatattgtagtagtaccagctgctatgcc
>IGHD2-21*01
agcatattgtggtggtgattgctattcc
>IGHD2-21*02
agcatattgtggtggtgactgctattcc
>IGHD2-8*01
aggatattgtactaatggtgtatgctatacc
>IGHD2-8*02
aggatattgtactggtggtgtatgctatacc
>IGHD2/OR15-2a*01
agaatattgtaatagtactactttctatgcc
>IGHD2/OR15-2b*01
```

agaatattgtaatagtactactttctatgcc
>IGHD3-10*01
gtattactatggttcggggagttattataac
>IGHD3-10*02
gtattactatgttcggggagttattataac
>IGHD3-16*01
gtattatgattacgtttgggggagttatgcttatacc
>IGHD3-16*02
gtattatgattacgtttgggggagttatcgttatacc
>IGHD3-22*01
gtattactatgatagtagtggttattactac
>IGHD3-3*01
gtattacgatttttggagtggttattatacc
>IGHD3-3*02
gtattagcatttttggagtggttattatacc
>IGHD3-9*01
gtattacgatattttgactggttattataac
>IGHD3/OR15-3a*01
gtattatgatttttggactggttattatacc
>IGHD3/OR15-3b*01
gtattatgatttttggactggttattatacc
>IGHD4-11*01
tgactacagtaactac
>IGHD4-17*01
tgactacggtgactac
>IGHD4-23*01
tgactacggtggtaactcc
>IGHD4-4*01
tgactacagtaactac
>IGHD4/OR15-4a*01
tgactatggtgctaactac
>IGHD4/OR15-4b*01
tgactatggtgctaactac
>IGHD5-12*01

gtggatatagtggctacgattac
>IGHD5-18*01
gtggatacagctatggttac
>IGHD5-24*01
gtagagatggctacaattac
>IGHD5-5*01
gtggatacagctatggttac
>IGHD5/OR15-5a*01
gtggatatagtgtctacgattac
>IGHD5/OR15-5b*01
gtggatatagtgtctacgattac
>IGHD6-13*01
gggtatagcagcagctggtac
>IGHD6-19*01
gggtatagcagtggctggtac
>IGHD6-25*01
gggtatagcagcggctac
>IGHD6-6*01
gagtatagcagctcgtcc
>IGHD7-27*01
ctaactgggga

**J.fasta**

```
>IGHJ1*01
gctgaatacttccagcactggggccagggcaccctggtcaccgtctcctcag
>IGHJ2*01
ctactggtacttcgatctctggggccgtggcaccctggtcactgtctcctcag
>IGHJ3*01
tgatgcttttgatgtctggggccaagggacaatggtcaccgtctcttcag
>IGHJ3*02
tgatgcttttgatatctggggccaagggacaatggtcaccgtctcttcag
>IGHJ4*01
actactttgactactggggccaaggaaccctggtcaccgtctcctcag
>IGHJ4*02
actactttgactactggggccagggaaccctggtcaccgtctcctcag
>IGHJ4*03
gctactttgactactggggccaagggaccctggtcaccgtctcctcag
>IGHJ5*01
acaactggttcgactcctggggccaaggaaccctggtcaccgtctcctcag
>IGHJ5*02
acaactggttcgacccctggggccagggaaccctggtcaccgtctcctcag
>IGHJ6*01
attactactactactacggtatggacgtctgggggcaagggaccacggtcaccgtctcct
cag
>IGHJ6*02
attactactactactacggtatggacgtctggggccaagggaccacggtcaccgtctcct
ca
>IGHJ6*03
attactactactactactacatggacgtctggggcaaagggaccacggtcaccgtctcct
ca
>IGHJ6*04
attactactactactacggtatggacgtctggggcaaagggaccacggtcaccgtctcct
cag
```

**IgDiscover configuration file (igdiscover.yaml)**


```yaml
## IgDiscover configuration


# How many discovery iterations to run. If 0, no updated database is created,
# but expression profiles are still computed. Unless working with a highly
# incomplete starting database, a single iteration is usually sufficient.
#
iterations: 1


# Type of sequences: Choose 'Ig' or 'TCR'.
#
sequence_type: Ig


## Barcoding settings


# If you have a random barcode sequence (unique molecular identifier) at the 5' end,
# set this to its length. Leave at 0 when you have no 5' barcode.
#
barcode_length_5prime: 17


# Same as above, but for the 3' end of the sequence. Leave at 0 when you have no 3' barcode.
# Currently, you cannot have a barcode in both ends, so at least one of the two settings
# must be zero.
#
barcode_length_3prime: 0


# When barcoding is enabled, sequences that have identical barcode and CDR3 are
```

```
# collapsed into a single consensus sequence.
# If you set this to false, no collapsing and consensus taking is done and
# only the barcode is removed from each sequence.
#
barcode_consensus: false


# When grouping by barcode and CDR3, the CDR3 location is either detected with a
# regular expressions or a 'pseudo' CDR3 sequence is used, which is at a
# pre-defined position within the sequence.
#
# Set this configuration option to a region like [-80, -60] to use a pseudo
# CDR3 located at bases 80 to 60 counted from the 3' end. (Use negative numbers to
# count from the 3' end, positive ones to count from the 5' end. The most 5'
# base has index 0.)
#
# Set this to 'detect' (with quotation marks) in order to use CDR3s
# detected by regular expression. This assumes that the input contains
# VH sequences!
#
# Set this to false (no quotation marks) in order to *only* group by barcode, not by CDR3.
#
cdr3_location: 'detect'  # Works only with VH sequences!


# When you use a RACE protocol, then the sequences have a run of G nucleotides in the beginning
# which need to be removed when barcodes are used. If you use RACE, set this to true.
# The G nucleotides are assumed to be in the 5' end (but after the barcode if it exists).
#
race_g: true


## Primer-related settings
```

```
# If set to true, it is assumed that the forward primer is always at the 5' end
# of the first read and that the reverse primer is always at the 5' end of the
# second read. If it can also be the other way, set this to false.
# This setting has no effect if no primer sequences are defined below.
#
stranded: false

# List of 5' primers
#
forward_primers:


# List of 3' primers
#
reverse_primers:


# Work only on this number of reads (for quick test runs). Set to false to
# process all reads.
#
#limit: false


# Filter out merged reads that are shorter than this length.
#
minimum_merged_read_length: 300


# Read merging program. Choose either 'pear' or 'flash'.
# pear merges more reads, but is slower.
#
merge_program: pear
```

```
# Maximum overlap (-M) for the flash read merger.
# If you use pear, this is ignored.
#
flash_maximum_overlap: 300


# Do not mention the original FASTA or FASTQ sequence names in the
# assigned.tab files, but instead use names <analysis_directory_name>_seq<number>,
# where <number> is a running number starting at 1.
#  true: yes, rename
#  false: no, do not rename
#
rename: false


# Whether debugging is enabled or not. Currently, if this is set to true,
# some large intermediate files that would otherwise be deleted will be
# kept.
#
debug: false


# The "seed value" is in arbitrary number used to get reproducible
# runs. Two runs that use the same software version, the same seed
# and otherwise the same configuration will give identical results.
#
# Set this to false in order to use a different seed each run.
# The results will then be not exactly reproducible.
#
seed: 1


# The preprocessing filter is always applied directly after running IgBLAST,
# even if no gene discovery is requested.
```

```
#
preprocessing_filter:
  v_coverage: 90   # Match must cover V gene by at least this percentage
  j_coverage: 60   # Match must cover J gene by at least this percentage
  v_evalue: 0.001  # Highest allowed V gene match E-value


## Candidate discovery settings

# When discovering new V genes, ignore whether a J gene has been assigned
# and also ignore its %SHM.
#  true: yes, ignore the J
#  false: do not ignore J assignment, do not ignore its %SHM
#
ignore_j: false


# When clustering sequences to discover new genes, subsample to this number of
# sequences. Higher is slower.
#
subsample: 1000


# When computing the Ds_exact column, consider only D hits that
# cover the reference D gene sequence by at least this percentage.
#
#d_coverage: 70


## V candidate filtering (germline filtering) settings


# Filtering criteria applied to candidate sequences in all iterations except the last.
#
```

```
pre_germline_filter:
  unique_cdr3s: 2            # Minimum number of unique CDR3s (within exact matches)
  unique_js: 2               # Minimum number of unique J genes (within exact matches)
  whitelist: true            # Add database sequences to the whitelist
  cluster_size: 0            # Minimum number of sequences assigned to cluster
  allow_stop: true           # Whether to allow non-productive sequences containing stop codons
  cross_mapping_ratio: 0.02  # Threshold for removal of cross-mapping artifacts (set to 0 to
disable)
  clonotype_ratio: 0.12      # Required minimum ratio of clonotype counts between alleles of the
same gene
  exact_ratio: 0.12          # Required minimum ratio of "exact" counts between alleles of the
same gene
  cdr3_shared_ratio: 0.8      # Maximum allowed CDR3_shared_ratio
#  unique_d_ratio: 0.3        # Minimum Ds_exact ratio between alleles
#  unique_d_threshold: 10     # Check Ds_exact ratio only if highest-expressed allele has at
least this Ds_exact count


# Filtering criteria applied to candidate sequences in the last iteration.
# These should be more strict than the pre_germline_filter criteria.
#
germline_filter:
  unique_cdr3s: 5            # Minimum number of unique CDR3s (within exact matches)
  unique_js: 3               # Minimum number of unique J genes (within exact matches)
  whitelist: true            # Add database sequences to the whitelist
  cluster_size: 100          # Minimum number of sequences assigned to cluster
  allow_stop: false          # Whether to allow non-productive sequences containing stop codons
  cross_mapping_ratio: 0.02  # Threshold for removal of cross-mapping artifacts (set to 0 to
disable)
  clonotype_ratio: 0.12      # Required minimum ratio of clonotype counts between alleles of the
same gene
  exact_ratio: 0.12          # Required minimum ratio of "exact" counts between alleles of the
same gene
  cdr3_shared_ratio: 0.8     # Maximum allowed CDR3_shared_ratio
```

```
#  unique_d_ratio: 0.3        # Minimum Ds_exact ratio between alleles
#  unique_d_threshold: 10     # Check Ds_exact ratio only if highest-expressed allele has at
least this Ds_exact count


## J discovery settings

j_discovery:
  allele_ratio: 0.2         # Required minimum ratio between alleles of a single gene
  cross_mapping_ratio: 0.1  # Threshold for removal of cross-mapping artifacts.
  propagate: true           # Use J genes discovered in iteration 1 in subsequent ones
```
